# Regulated Induced Proximity Targeting Chimeras (RIPTACs): a Novel Heterobifunctional Small Molecule Therapeutic Strategy for Killing Cancer Cells Selectively

**DOI:** 10.1101/2023.01.01.522436

**Authors:** Kanak Raina, Chris D. Forbes, Rebecca Stronk, Jonathan P. Rappi, Kyle J. Eastman, Samuel W. Gerritz, Xinheng Yu, Hao Li, Amit Bhardwaj, Mia Forgione, Abigail Hundt, Madeline P. King, Zoe M. Posner, Allison Denny, Andrew McGovern, David E. Puleo, Ethan Garvin, Rebekka Chenard, Nilesh Zaware, James J. Mousseau, Jennifer Macaluso, Michael Martin, Kyle Bassoli, Kelli Jones, Marco Garcia, Katia Howard, Levi M. Smith, Jinshan M. Chen, Cesar A. De Leon, John Hines, Katherine J. Kayser-Bricker, Craig M. Crews

## Abstract

While specific cell signaling pathway inhibitors have yielded great success in oncology, directly triggering cancer cell death is one of the great drug discovery challenges facing biomedical research in the era of precision oncology. Attempts to eradicate cancer cells expressing unique target proteins, such as antibody-drug conjugates (ADCs), T-cell engaging therapies, and radiopharmaceuticals have been successful in the clinic, but they are limited by the number of targets given the inability to target intracellular proteins. More recently, heterobifunctional small molecules such as Proteolysis Targeting Chimera (PROTACs) have paved the way for protein proximity inducing therapeutic modalities.

Here, we describe a proof-of-concept study using novel heterobifunctional small molecules called **<u>R</u>**egulated **I**nduced **<u>P</u>**roximity **<u>Ta</u>**rgeting **<u>C</u>**himeras or RIPTACs, which elicit a stable ternary complex between a target protein selectively expressed in cancer tissue and a pan-expressed protein essential for cell survival. The resulting cooperative protein:protein interaction (PPI) abrogates the function of the essential protein, thus leading to cell death selectively in cells expressing the target protein. This approach not only opens new target space by leveraging differentially expressed intracellular proteins but also has the advantage of not requiring the target to be a driver of disease. Thus, RIPTACs can address non-target mechanisms of resistance given that cell killing is driven by inactivation of the essential protein.

Using the HaloTag7-FKBP model system as a target protein, we describe RIPTACs that incorporate a covalent or non-covalent target ligand connected via a linker to effector ligands such as JQ1 (BRD4), BI2536 (PLK1), or multi-CDK inhibitors such as TMX3013 or dinaciclib. We show that these RIPTACs exhibit positive co-operativity, accumulate selectively in cells expressing HaloTag7-FKBP, form stable target:RIPTAC:effector trimers in cells, and induce an anti-proliferative response in target-expressing cells. We propose that RIPTACs are a novel heterobifunctional therapeutic modality to treat cancers that are known to selectively express a specific intracellular protein.

## Introduction

A classical targeted therapy is typically designed to inhibit a cancer driver protein (e.g., AR, mEGFR, KRASG12C) or a pathway on which the cancer cell is selectively dependent (e.g., proteasome, PARP). While it has been estimated that there are about 300 cancer driver proteins, only about 50 have been successfully targeted with an FDA-approved drug[1], [2]. Since the advent of the genomics revolution, many targeted therapies have advanced into Phase I clinical trials in oncology. However, the overall likelihood of success for cancer drugs entering the clinic range remains low[3], [4]. Targeted therapies fail for two principal and often related reasons: lack of efficacy and toxicity[5]. For a variety of reasons, the drug discovery process has historically identified genes essential for cell survival as cancer-selective targets[6]. As a result, several selective agents that have been tested in patients target essential proteins such as mitotic kinases like PLK1 and AURKA/B and epigenetic regulators like BRD4 and the HDACs. Many of these agents have failed in the clinic due to the lack of a Therapeutic Index (TI), while a few have received FDA approval albeit with a narrow TI and significant side effects.

Alternatively, novel methods to selectively trigger cancer cell death have emerged in the past few decades. Antibody-drug conjugates (ADCs), T-cell engagers, and radiopharmaceuticals all deploy a multifunctional agent to engage a cancer cell and then to selectively cause its death. In the case of an ADC, a toxic payload is delivered to the cancer cell. T-cell recruiters or CAR-T cells activate T cells to secrete perforin and granzyme in the vicinity of a cancer cell, thus killing it. Radiopharmaceuticals also selectively bind cancer cells and their ability to trigger cell death relies on emitting radiation over a short distance. The target protein/antigen for these approaches does not need to be a driver of the disease, thus greatly expanding the number of cancer targets in oncology. In particular, lineage-specific antigens, such as CD19, have been successfully employed as targets for ADCs and CAR-T therapy in hematological malignancies[7], [8].

Unfortunately, the above modalities rely on engaging cancer cells extracellularly, and as a result are unable to take advantage of most cancer-selective proteins. In addition, their utility is limited by their complexity and cost. More recently, heterobifunctional protein degrading small molecules like Proteolysis Targeting Chimera (PROTACs) have paved the way for beyond rule of five (bRo5) drugs as novel therapeutics[9]–[11]. Initial concerns about the potential drawbacks of the physicochemical properties of such molecules and the expectations surrounding poor oral bioavailability and pharmacokinetics have been allayed by recent positive clinical trial data with oral PROTACs reported by companies like Arvinas, Kymera Therapeutics, and Nurix Therapeutics. PROTACs rely on the induction of novel protein-protein interactions (PPIs) in a trimeric complex comprising the drug, its target and an E3 ubiquitin ligase. The PPIs impart co-operativity to this protein complex, resulting in high binding affinity, and selective degradation of the target protein[12]–[14].

The concept of ligand-induced PPIs has a long history in natural products prior to its use in the engineering of heterobifunctional small molecules. Examples include the macrolides rapamycin, which recruits the prolyl isomerase FKBP12 to inhibit the kinase activity of mTOR, and cyclosporin A, which forms a complex with cyclophilin to inhibit the phosphatase calcineurin[15]–[17]. Recruitment of immunophilins using small molecules has also been attempted in a more targeted manner and the first KRas inhibitor from Revolution Medicines that works via this mechanism entered clinical trials in 2022[18].

We hypothesized that it is possible to selectively induce death in cells expressing a particular intracellular Target Protein (TP) using a heterobifunctional small molecule called a RIPTAC via formation of a co-operative ternary complex with the TP and a pan-essential Effector Protein (EP) required for survival. Such a ternary complex would abrogate EP function selectively in target-expressing cells, causing cell death. Herein we describe how positive co-operativity, a well-known feature of ternary complexes formed by biologically active heterobifunctional small molecules, allows for selective accumulation inside the cell via TP binding and /or concurrent engagement of the pan-essential protein. This unique mechanism is leveraged into a novel treatment modality that delivers a therapeutic window relative to cells lacking the TP (Fig. 1a).

**Fig. 1.**
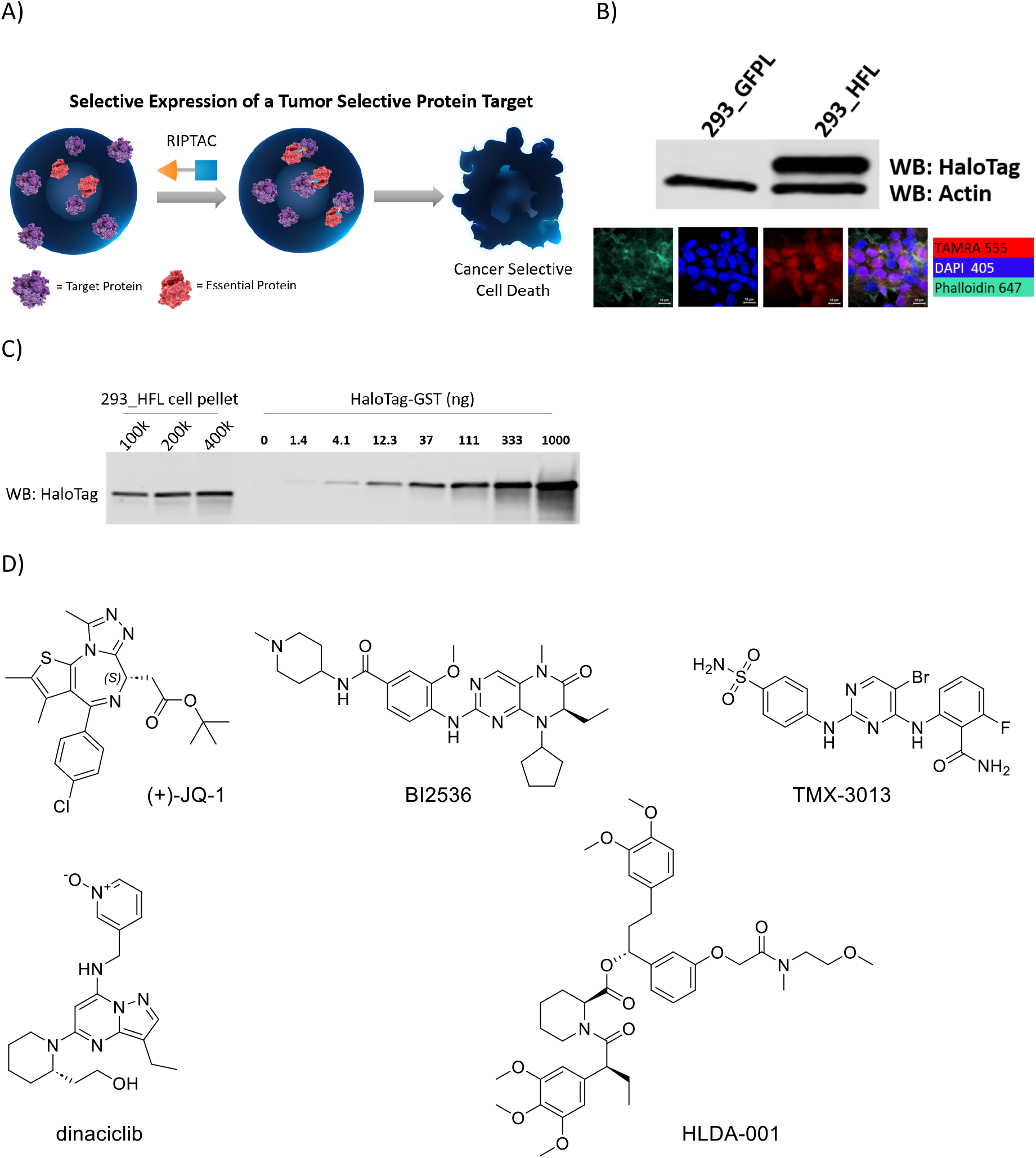
The RIPTAC hypothesis and the HaloTag-FKBP model system. a) RIPTACs leverage target-dependent intracellular accumulation and positive co-operativity in ternary complex formation to selectively inhibit the proliferation of target protein expressing cells. b) The HaloTag-FKBP fusion protein is selectively expressed in 293_HFL cells and not in the control 293_GFPL cells. c) Quantitation of cellular HaloTag-FKBP concentration using recombinant HaloTag-GST standard curve. d) RIPTAC Effector Ligands (ELs) JQ-1, BI2536, TMX-3013, Dinaciclib, and the FKBP Target Ligand (TL) HLDA-001.

To reduce RIPTACs to practice, we designed a proof-of-concept study using chemical biology systems with well-characterized target proteins and ligands. Employing a HaloTag-FKBP12^F36V^ fusion protein as a TP, we demonstrate ternary complex formation with various RIPTACs and their cognate EPs, ultimately resulting in the selective killing of cells expressing the HaloTag-FKBP12^F36V^ fusion.

## Results

### The HaloTag-FKBP model system

To generate a model system, we engineered a HEK293-derived cell line (293_HFL) with lentiviral overexpression of Flag-tagged HaloTag7-FKBP^F36V^ (hereafter HaloTag-FKBP) with a C-terminal P2A-EGFP sequence as target protein. This allowed access to both covalent (via HaloTag) and non-covalent (via FKBP) means of recruiting the same TP. A cell line overexpressing only EGFP (hereafter 293_GFPL) was generated as a control. GFP expression-based sorting by flow cytometry allowed selection of a high expressing clone, in which HaloTag-FKBP expression was confirmed by western blotting (Fig. 1b). Confirmation that the fusion protein expression was not restricted to any one compartment of the cell was determined with confocal microscopy using the HaloTag binding TAMRA-CA fluorescent reagent (Fig. 1b). In order to more thoroughly characterize the cell line, the concentration of the HaloTag-FKBP protein inside the cell was determined. Whole cell lysates from a fixed number of 293_HFL cells were probed using a HaloTag antibody alongside a purified recombinant HaloTag-GST protein standard curve (Fig. 1c). Assuming a HEK293 cell radius of 6.5μm, this yielded an intracellular target protein concentration of 1μM-3μM. Suitable essential effector proteins were chosen based on published CRISPR dropout screening data and the potency, selectivity, and extent of validation of the chemical matter available to bind them. The final Effector Ligands (ELs) chosen were JQ1 (BET inhibitor), BI2536 (PLK1 inhibitor), TMX3013 (multi-CDK inhibitor), and dinaciclib (multi-CDK inhibitor). The chemical structures of these molecules as well as the FKBP binding ligand are shown in Fig. 1d.

### RIPTACs show enhanced anti-proliferative activity in HaloTag-FKBP expressing cells

We synthesized RIPTACs of varying linker lengths with the above effector ligands at one end, and either a HaloTag-binding chloroalkane (CA), or an FKBP-binding ligand at the other (Fig. 2a). Dinaciclib, for which only chloroalkane-containing RIPTACs were synthesized, was the exception. Binding of select RIPTACs to their respective EPs was confirmed in biochemical assays (Supplementary Table 1). Next, we treated 293_HFL and 293_GFPL cells with these RIPTACs in CellTiter-Glo viability assays in two different formats. First, we tested these compounds in a 7-day assay with continuous RIPTAC exposure. Second, RIPTACs were washed out following a 4h treatment and cellular viability determined after 7 days. A clear pattern of enhanced anti-proliferative RIPTAC activity in 293_HFL cells was observed for every RIPTAC series under both treatment conditions, with the fold-shift in 293_HFL cells showing enhancement for RIPTACs incorporating BI-2536 or dinaciclib under washout conditions relative to continuous treatment (Supplementary Fig. 1a). The best performing linker lengths for each EL are shown in Fig.2b (covalent) & Fig.2c (non-covalent). Importantly, neither the ELs themselves nor the FKBP target ligand showed a significant difference in potency between the two cell lines (Supplementary Fig. 1b). The GI_50_ values for the optimal linker length for each series, along with the *in vitro* fold-selectivity over the non-target expressing 293_GFPL cell line are summarized in Fig. 2d. Next, we confirmed that the differential anti-proliferative activity seen with RIPTACs in presence of the target protein was not an artifact of the cell line background. We generated a HT29 derived cell line stably expressing the HaloTag-FKBP protein and tested the JQ1-CA HLDA-124 in a three day CellTiter-Glo assay. We observed a clear, albeit modest, leftward shift in the TP-expressing HT29_HF cell line (Supplementary Fig. 1c). Similarly, we confirmed that the observed leftward shifts in TP-expressing lines were not an artifact of the FKBP immunophilin domain of the fusion protein. We generated 293 cells expressing either Akt-HaloTag (293_AktH), or HaloTag-BTK (293_HBTKL), and in both instances observed leftward shifts in RIPTAC potency in CellTiter-Glo assays, although not necessarily with all effector ligands (Supplementary Fig. 1d).

**Fig. 2.**
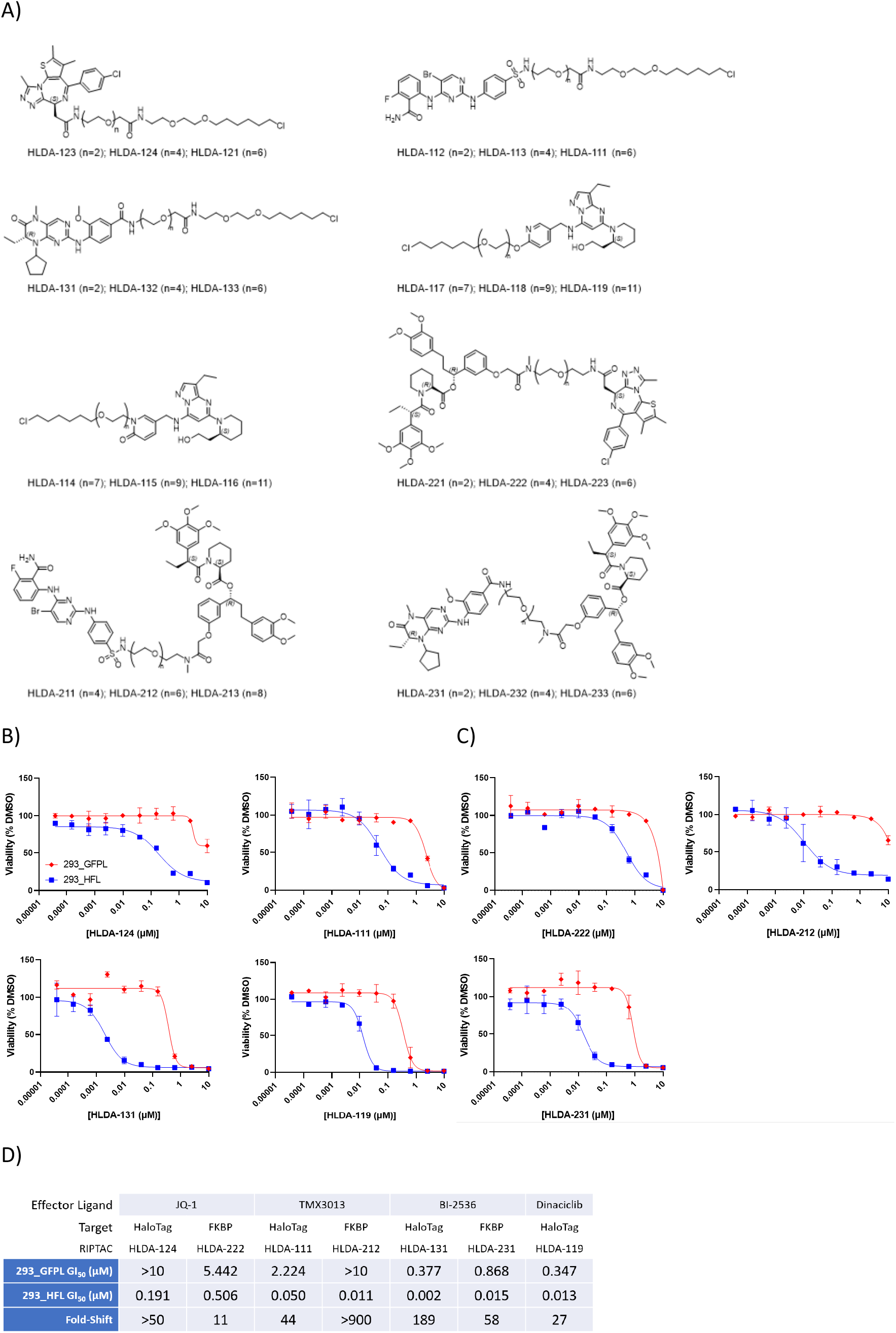
RIPTAC differential biology in 293_HFL model system. a) Structures of HaloTag-FKBP fusion protein targeting RIPTACs with JQ1, TMX3013, BI-2536, or dinaciclib as effector ligands b) Differential anti-proliferative activity of select covalent RIPTACs in target protein expressing 293_HFL cells in a 7-day Cell TiterGlo assay c) Differential anti-proliferative activity of select non-covalent RIPTACs in a 7-day Cell TiterGlo assay. d) GI_50_ values and fold-shift over 293_GFPL cells, obtained under continuous 7-day treatment with the optimal linker length from each RIPTAC series. All data representative of 3 independent experiments (N=3)

### Mechanisms underlying RIPTAC differential biology

We next sought to determine at a molecular level whether our hypothesis regarding RIPTAC accumulation via ternary complex formation was sufficient to explain the observed leftward shifts in RIPTAC activity in TP-containing cell lines. We first tested whether the FKBP ligand HLDA-001 showed any target-dependent accumulation in cells. We treated 293_GFPL and 293_HFL cells with 10nM or 100nM HLDA-001 for 4h, trypsinized the cells, and analyzed the cell pellet by LCMS (see methods). We observed >1000x enrichment of the ligand in 293_HFL cells compared to the control cell line lacking the fusion protein (Fig. 3a). Next, we tested select non-covalent RIPTACs using the same method and observed significant accumulation of HLDA-222 (JQ1), HLDA-212 (TMX3013), and HLDA-231 (PLK1) in the 293_HFL cell line, although the values were smaller than those observed for HLDA-001 (Fig. 3b). RIPTAC accumulation was shown to be target-dependent as evidenced by the loss of HLDA-212 accumulation upon pre-treatment with excess of the FKBP ligand HLDA-001 (Supplementary Fig. 2a).

**Fig. 3:**
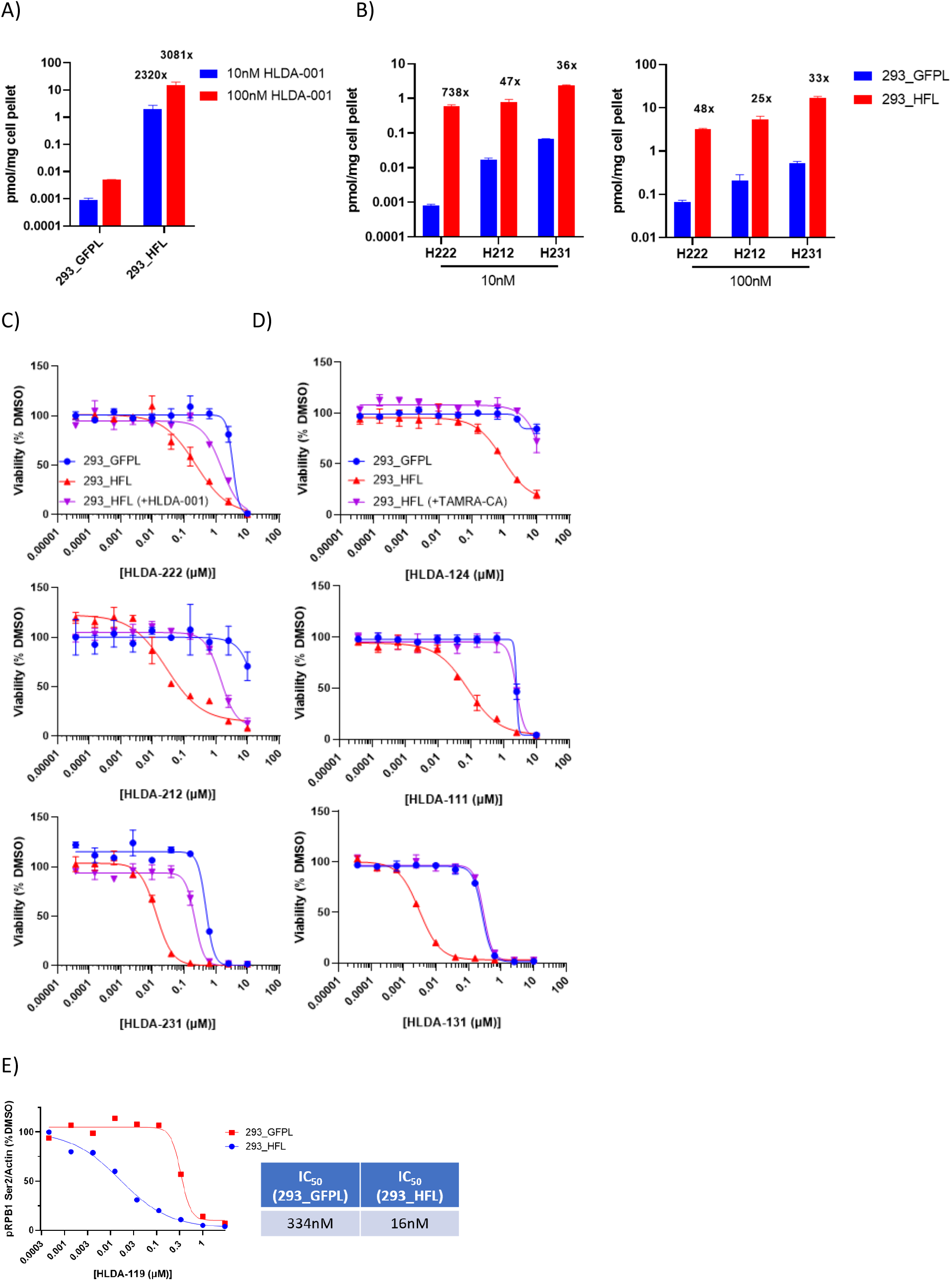
Mechanisms underlying RIPTAC differential biology. a) The FKBP ligand accumulates selectively in 293_HFL cells b) Non-covalent RIPTACs also accumulate selectively in 293_HFL cells c&d) RIPTAC activity in 7 day viability assays can be competed off by pre-treatment with 10μM HLDA-001 in the case of non-covalent RIPTACs and 300nM TAMRA-CA in the case of covalent RIPTACs e) RIPTAC HLDA-119 is a 20-fold more potent CDK9 inhibitor in 293_HFL cells than in control 293_GFPL cells.

Similarly, the differential biology observed in viability assays was shown to be target-dependent as shown by the loss of a leftward shift in 293_HFL viability upon competition with TAMRA-CA (Fig. 3c), and HLDA-001 (Fig 3d). This data also confirmed that the observed differential biology was not a result of the combined cytoxicities of the target and effector ligands. The effector ligands by themselves showed no shift under competition conditions (Supplementary Fig. 2b). As an additional control, we confirmed the role of the TP in the observed viability effects by testing des-chloro RIPTAC negative controls (HLDA-120 and HLDA-110) that cannot bind HaloTag (Supplementary Fig. 2c, Supplementary Fig. 4a) and the role of the effector with a JQ1 epimer negative control RIPTAC HLDA-125 that cannot bind BRD2/3/4 (Supplementary Fig. 2d, Supplementary Fig. 4a). As expected, none of these controls showed any differential antiproliferative activity in the 293_HFL cell line.

We next asked whether the leftward shift in cell viability seen with a RIPTAC could be recapitulated by a leftward shift in the cellular inactivation of the EP. For this purpose, we chose the dinaciclib-CA RIPTAC HLDA-119. We treated 293_GFPL and 293_HFL cells for 4h with increasing concentrations of either HLDA-119 or dinaciclib and measured the levels of phosphorylation at Ser2 of RPB1-CTD. The putative kinase for this site is CDK9, which is a known cellular target of dinaciclib. The IC_50_ of HLDA-119 in 293_HFL cells was 16nM compared to 334nM in 293_GFPL cells, a shift of ∼20-fold (Fig. 3e and Supplementary Fig. 2e). These values are highly congruous with the GI_50_ values of 13nM and 347nM observed for the same RIPTAC in viability assays in the two cell lines. In comparison, dinaciclib itself showed no significant difference in IC_50_ between the cell lines (Supplementary Fig. 2e).

### Ternary Complex Formation with RIPTACs

We next confirmed that the observed differential biology with RIPTACs was due to the formation of TP:RIPTAC:EP ternary complexes. We treated 293_HFL cells with JQ1-based non-covalent RIPTAC HLDA-222 and covalent RIPTAC HLDA-121 for 3h, after which we immunoprecipitated the HaloTag protein with Halo-Trap Agarose beads. We observed co-immunoprecipitation of BRD4 with both molecules, but not with the negative control HLDA-125 (Fig. 4a). We next showed that the BRD4 coimmunoprecipitation seen with HLDA-121 and HLDA-222 could be prevented by pre-treating the 293_HFL cells for 30min with TAMRA-CA in the case of HLDA-121, and HLDA-001 in the case of HLDA-222 (Fig. 4b). In the same experiment, pre-treatment with JQ1 also prevented complex formation with both RIPTACs, but only partially. Using the same Halo-Trap based pulldown protocol, we showed that cellular ternary complexes could also be obtained with treatment of 293_HFL cells with 100nM of the PLK1 binding non-covalent RIPTAC HLDA-231, 100nM of the CDK9 binding covalent RIPTAC HLDA-115, and 100nM of the CDK1/2/5 binding covalent RIPTAC HLDA-111, but not with negative control ELs, the FKBP TL HLDA-001, or the TAMRA-CA HaloTag ligand (Supplementary Fig 3a). We also established that the cellular ternary complexes formed by RIPTACs were profoundly stable by conducting a washout experiment. We treated 293_HFL cells with 1μM HLDA-121 or 100nM HLDA-222 for 2h, followed by washout, and tracked complex formation over a period of 72h. Incredibly, the cellular ternary complexes, even with the non-covalent RIPTAC HLDA-222, were readily detectable even 72h following washout (Fig. 4c).

**Fig. 4.**
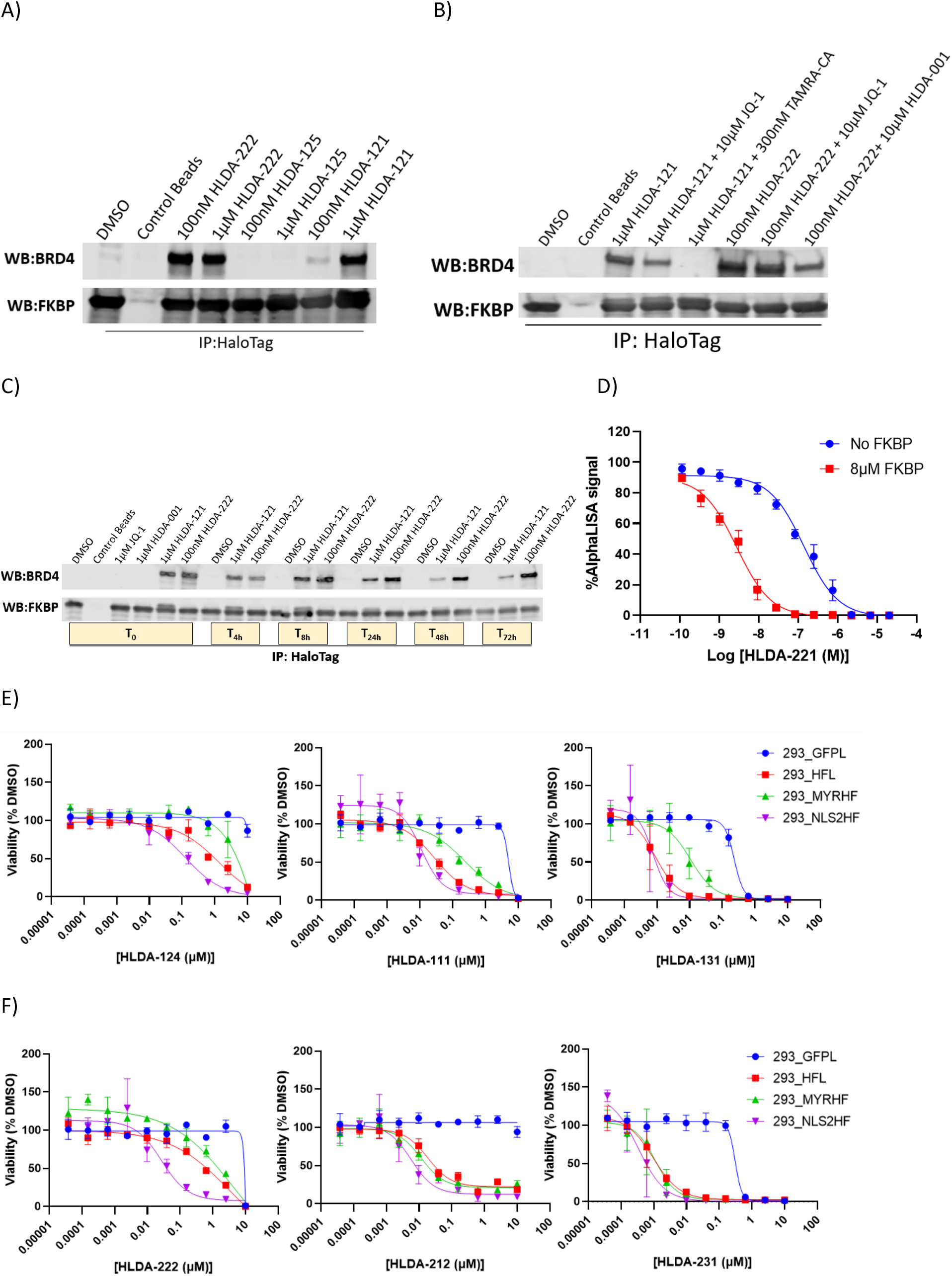
Ternary Complex Formation with RIPTACs. a) 3h treatment of 293_HFL cells with indicated compounds followed by HaloTag immunoprecipitation using HaloTrap beads demonstrates cellular ternary complex formation with both the non-covalent RIPTAC HLDA-222 and the covalent RIPTAC HLDA-121, but not with the control molecule HLDA-125 that does not bind BRD4. b) Ternary complex formation with HLDA-121 and HLDA-222 can be competed away with 30min pre-treatment with TAMRA-CA and HLDA-001, respectively. 30min JQ1 pre-treatment competes away complex formation with both RIPTACs partially. c) 293_HFL cells were treated with the indicated compounds for 3h (T_0_), following which the compound treated medium was washout out and replaced with normal growth medium for up to 72h. The HaloTag-FKBP fusion protein was immunoprecipitated at the timepoints shown and the BRD4 protein levels in the complex were detected by immunoblotting.d) AlphaLISA assay measuring BRD4-BD1 inhibition demonstrates positive co-operativity in biochemical RIPTAC ternary complex formation in presence of the non-covalent RIPTAC HLDA-221 and FKBP. e) 7 day CellTiter Glo viability assay with indicated covalent RIPTACs in cell lines expressing the HaloTag-FKBP fusion protein selectively in the nucleus (293_NLS2HF), plasma membrane (293_MYRHF), or both (293_HFL). f) 7 day CellTiter Glo viability assay in the same cell lines using non-covalent RIPTACs.

Given that ternary complex formation is an underlying mechanistic feature of all RIPTACs, we next tested whether positive co-operativity in complex formation could in part account for the enhanced potency seen with RIPTACs in 293_HFL cells. For this purpose, we developed an AlphaLISA-based assay, in which we measured the binding of JQ1 containing non-covalent RIPTAC HLDA-221 to BRD4-BD1 in the presence or absence of purified recombinant FKBP. We observed a dramatic (∼35-fold, range 25-50 fold) increase in BRD4 BD1 binding of HLDA-221 upon pre-incubation with FKBP (Fig. 4d). No such potency shift was observed for the EL JQ1, or the target ligand HLDA-001 (Supplementary Fig. 3b). This suggests that, as with other heterobifunctional molecules and molecular glues, protein-protein interactions (PPI) play an important role in the pharmacology of RIPTACs, and likely partially explain the selective antiproliferative activity seen in 293_HFL cells.

Having established that RIPTAC differential biology likely benefits from the ability of the TP and EP to form ternary complexes, we postulated that the relative subcellular localization of the two proteins in question likely contributes significantly to the RIPTAC differential biology observed in viability assays. To test this hypothesis, cell lines expressing either a nuclear localized (293_NLS2HF) or a plasma membrane localized (293_MYRHF) version of the HaloTag-FKBP fusion protein (Supplementary Fig. 3c) were tested with select RIPTACs in parallel with 293_HFL cells (Figs 4e and 4f). Interestingly, while covalent RIPTACs incorporating JQ1, TMX3013, and BI2536 showed reduced potency in the 293_MYRHF cell line, the FKBP-BRD4 RIPTAC HLDA-222 was the only non-covalent RIPTAC that showed a clear preference for the nuclear localized target protein compared to the plasma membrane localized one. Non-covalent RIPTACs may therefore be more tolerant towards disparate TP-EP localization than covalent RIPTACs. The activity of control compounds, including all three ELs and the FKBP ligand HLDA-001, did not differ significantly in all four cell lines tested (Supplementary Fig. 3d).

## Discussion

We have described herein a chemical biology proof-of-concept study for a novel heterobifunctional small molecule modality called RIPTACs, which has potential as an anticancer therapeutic strategy. Our hypothesis stemmed from two previously reported observations.

Firstly, a large number of Fluorine-18 and Carbon-11 Positron Emission Tomography (PET) radiotracers have been used in preclinical *in vivo* studies to monitor protein expression in tissues that express their cognate receptor, and many have made their way into clinical use[19]–[21]. Examples of FDA-approved PET ligands include Piflufolastat F-18 (PYLARIFY), used to detect PSMA+ prostate cancer, [^18^F] fluoroestradiol (CERIANNA) for detection of ER-positive breast cancer, and Ga-68 dotatate (NETSPOT) for detection of somatostatin receptor positive neuroendocrine tumors, among others. This suggests that some subset of high-affinity, high selectivity ligands may be prone to accumulation in cells that express their protein target. The biochemical, physicochemical, and pharmacokinetic rules that enable the identification of such ligands remain to be established.

Secondly, we know from the field of heterobifunctional small molecules and molecular glues that the apparent K_d_ of a ligand:protein interaction can be dramatically enhanced by the presence of beneficial PPIs in a ternary complex[22], [23]. As elucidated by a number of landmark structural studies, this is because the formation of the first ligand:protein interaction results in the creation of a composite surface that has higher affinity for the second protein than the small molecule or the first protein by itself[24]– [26]. Our work using the HaloTag model system suggests that the ability to form a stable trimer is not restricted to proteins with a chaperone function, like immunophilins, and enzymes with a relatively promiscuous substrate recognition domain, like some E3 ligases.

Our hypothesis relies on the expression of a target protein that we use to block the function of a pan-essential effector protein in a target-expressing, cell-selective manner. There are several additional factors to consider in developing the RIPTAC technology as a platform.

Firstly, the relative abundance of the target protein and the pan-essential effector protein will likely play a role in RIPTAC activity. The HaloTag-FKBP model target in our work is highly expressed by a CMV promoter and attains micromolar cellular concentrations at steady state. High concentrations in tumor cells have been reported for certain highly amplified oncogenes such as HER2[27], and the relationship between the level of target expression and the abundance of effector protein will be interrogated when this work is extended to cancer targets.

Secondly, as we have shown with chemical biology tools, the relative localization of the target and effector proteins can have a significant impact on the activity of the RIPTAC. In the case of BRD4 as an effector, for example, nuclear target proteins will clearly be the most amenable to the RIPTAC approach, while other effector proteins suggest greater compatibility with non-nuclear targets.

Thirdly, we have not examined here whether the RIPTAC mechanism of action involves more than mere inhibition of the active site of the effector protein. It is reasonable to speculate that in some instances RIPTAC activity may derive from partial or complete mislocalization of the effector protein, as a result of its incorporation into a stable ternary complex with the target bound RIPTAC. This may result in a phenotype more akin to genetic loss-of-function (LoF) of the effector protein rather than its inhibition with a small molecule. In fact, the greatest therapeutic index may be achieved with a RIPTAC that shows only weak binding to the effector protein in healthy cells lacking the target protein, but owing to ternary complex formation and novel PPIs, results in LoF of the effector protein in tumor cells. Such phenomena are well-documented in the field of molecular glues. Cereblon-binding imid drugs, for example, have little or no binding to the transcription factors degraded through formation of the ternary complex[28], and rapamycin by itself is only a weak micromolar binder of mTOR[29].

Fourthly, an interesting feature of RIPTACs is the enhancement of their selectivity for target-expressing cells under washout conditions in viability assays. This is likely a result of both aspects of the underlying pharmacology, namely, target dependent intracellular accumulation, and the stability of ternary complexes. Given the appropriate pharmacokinetics, it may be possible to leverage the rapid washout of RIPTACs from non-target expressing cells to further enhance the therapeutic index observed in whole organisms.

Finally, an important, salient feature of the RIPTAC modality is that it is agnostic to the identity of the oncogenic driver of the disease. Conventional targeted therapy relies on the inhibition of the cellular signaling pathway responsible for tumor growth. In response to such a drug, cancer cells frequently acquire resistance by a myriad of different mechanisms. For example, increasing expression of the target or splice variants such as in the case of AR in prostate cancer[30]–[32], increasing signaling flux via RTK upregulation in response to BRAF inhibitors in colorectal cancer[33]–[35], or turning on parallel growth pathways, such as MET amplification, NRas mutation among others, in the case of the newly FDA-approved KRas^G12C^ inhibitors in NSCLC and CRC[36]. Identifying the relevant resistance mechanism at play for each patient in the clinic remains a challenge. This leads to a need for further target validation and new therapies for each segment of patients in the relapsed/refractory population. RIPTACs, which rely only on the presence of a tumor-specific protein and not on its role in tumor biology, potentially offer a means to address this problem. We speculate that a KRas^G12C-^targeting RIPTAC, for example, would continue to be active regardless of the acquisition of parallel oncogenic signaling pathway alterations, and greatly reduce the number of possible mechanisms of resistance.

Taken together, our data support further study of RIPTACs as a novel heterobifunctional small molecule modality for the treatment of cancer. Future work will address several outstanding questions described in this chemical biology proof-of-concept study and focus on advancing bona fide drug-like RIPTACs towards the clinic.

## Acknowledgements

This study was supported in part by NIH R35 CA197589 to C.M.C., who is also an American Cancer Society Professor. C.A.D. was supported by an HHMI Hanna H. Gray Fellowship.

## Materials and Methods

### Cell lines and reagents

The HaloTag7-FKBP sequence was fused to a P2A-EGFP and cloned into a lentiviral expression vector and introduced into the HEK293 cell using polybrene (8μg/mL) mediated lentiviral transduction. Clonal selection was performed using 1μg/mL Puromycin. A GFP-infected HEK293 line was used as a negative control. In order to induce membrane localization of the fusion protein, an MGSSKSKPK sequence was added to the N-terminus, and an N-terminal PKKKRKV sequence used for nuclear localization. The 293_AktH and 293_HBTKL cell lines were generated similarly. All cell lines were maintained in DMEM (Gibco) supplemented with 10% heat inactivated FBS (Gibco), 1% penicillin-streptomycin (Gibco), and 1ug/mL Puromycin as needed. Cells were cultured in a 37°C incubator with 5% CO_2_. Compounds were dissolved in DMSO and all treatments performed in complete medium unless specified otherwise. TAMRA-CA was purchased from Promega (G8251). JQ-1 (HY-13030), BI-2536 (HY-50698), Dinaciclib (HY-10492) were purchased from MedChemExpress.

### Chemical Synthesis

RIPTAC synthesis and characterization provided in Supplementary Methods.

### Recombinant protein production

Protein expression and purification experiments were performed at Selvita S.A. *BRD4-BD1*: A DNA construct encompassing genes encoding for human BRD4 bromodomain 1 (BD1, aa 42-168) was codon-optimized for expression in *E*.*coli*, synthesized, and cloned into an appropriate expression vector together with the N-terminal 6xHis-tag and TEV cleavage site. The construct was transformed into BL21(DE3) using a standard protocol. The preculture was prepared by inoculation of several colonies into 20 mL of LB medium supplemented with 100 μg/mL kanamycin and incubated overnight at 37 °C at 200 rpm. This culture was used to seed 500 mL LB-autoinduction medium (LB-AIM, Formedium) supplemented with 100 μg/mL kanamycin and 8 g/L glycerol in 2-L Erlenmeyer flasks and grown at 30 °C shaking at 200 rpm until OD600 reached 0.6-0.8. Then the incubation temperature was lowered to 18 °C and flasks were incubated shaking at 200 rpm overnight. The cell pellets were harvested by centrifugation at 8,000 x g, for 10 minutes, and stored frozen at -20 °C. Pellets were resuspended in the lysis buffer, 25 mM HEPES, pH 7.5, 500 mM NaCl, 5% (w/v) glycerol, 0.5 mM TCEP, 5 mM imidazole, supplemented with protease inhibitor cocktail (Sigma), 0.1 μg/mL DNAse, and 1 mg/mL lysozyme, and lysed by three snap freeze-thaw cycles. The lysates were cleared by centrifugation at 19,000 x g, 50 minutes, and the supernatant was loaded into cOmplete™ His-Tag Purification Resin (Roche) pre-equilibrated with wash buffer, 25 mM HEPES, pH 7.5, 500 mM NaCl, 5% (w/v) glycerol, 0.5 mM TCEP, 5 mM imidazole. Non-specifically bound proteins were removed with wash buffer and specifically bound proteins were eluted using 25 mM HEPES, pH 7.5, 500 mM NaCl, 5% (w/v) glycerol, 0.5 mM TCEP, 250 mM imidazole. Elution fractions were analyzed using SDS-PAGE. The fractions containing BRD4 BD1 were pooled together, loaded into HiLoad® 26/600 Superdex® 75 pg (Cytiva), and eluted with 25 mM HEPES, pH 7.5, 200 mM NaCl, 0.5 mM TCEP. After SDS-PAGE analysis, fractions containing the protein of interest were pooled together and concentrated using Vivaspin 20 centrifugal concentrators with a 5 kDa cut-off (Sartorius). The concentration of BRD4 BD1 was determined spectrophotometrically at 280 nm using the calculated molar extinction coefficient. The content of residual nucleic acid was evaluated spectrophotometrically by measuring the ratio of absorptions A260/A280. *FKBP12:* For the production of the recombinant peptidyl-prolyl cis-trans isomerase FKBP12 F36V (aa 2-108), the DNA sequence coding for the target protein was codon optimized for expression in E.coli, synthesized, and cloned into the appropriate expression vector together with the N-terminal GST tag and HRV 3C cleavage site. The construct was transformed into BL21(DE3) using a standard protocol. The preculture was prepared using multiple colonies inoculated to the LB medium supplemented with 100 μg/mL kanamycin and incubated overnight at 37 °C at 200 rpm. Then it was used to seed 500 mL LB-autoinduction medium (LB-AIM, Formedium) supplemented with 100 μg/mL kanamycin and 8 g/L glycerol in 2-L Erlenmeyer flasks and grown at 30 °C shaking at 200 rpm until OD600 reached 0.6-0.8. Then the incubation temperature was lowered to 15 °C and flasks were incubated shaking 200 rpm overnight. The cell pellets were harvested by centrifugation at 8,000 x g, for 10 minutes, and stored frozen at -20 °C. Pellets were resuspended in 50 mM Tris, pH 7.2, 250 mM NaCl, 10% (w/v) glycerol, 0.5 mM TCEP, supplemented with 1 mg/mL lysozyme, 0.1 μg/mL DNase, and protease inhibitor cocktail (Sigma). The lysis was performed by three cycles of snap freeze-thaw. The lysate was cleared by centrifugation at 19,000 x g, 50 minutes, and the supernatant was loaded into Glutathione Sepharose 4B GST-tagged protein purification resin (Cytiva) and non-specifically bound proteins were removed by multiple washes with 50 mM Tris, pH 7.2, 250 mM NaCl, 10% (w/v) glycerol, 0.5 mM TCEP. The protein of interest was eluted using the same buffer supplemented with 15 mM reduced glutathione. The N-terminal GST-tag was removed by cleavage with HRV 3C protease in 1:100 (w/w) ratio overnight at 4 °C and then separated from the protein of interest using reverse affinity chromatography. The flow through containing FKBP12 F36V was loaded into HiLoad® 26/600 Superdex® 75 pg (Cytiva), and eluted with 50 mM Tris, pH 7.0, 150 mM NaCl, 1 mM TCEP, 10% (w/v) glycerol. After SDS-PAGE analysis, fractions containing FKBP12 F36V were pooled together and concentrated using Vivaspin 20 centrifugal concentrators with a 5 kDa cut-off (Sartorius). The concentration of FKBP12 F36V was determined spectrophotometrically at 280 nm using the calculated molar extinction coefficient. The content of residual nucleic acid was evaluated spectrophotometrically by measuring the ratio of absorptions A260/A280.

### Immunoprecipitation and immunoblotting

293_HFL cells were plated at 5×10^6^ cells per 75cm^2^ flask (#229341, cell treat scientific) in 15ml of complete DMEM media. After 24 hours, cells were treated with indicated compounds for 2h. After the treatments, cells were harvested and washed three times with 1xPBS. Halotag-FKBP Immunoprecipitation was performed using Halo-Trap Agarose beads (# OTA-200 Chromotek) as per manufacturer instructions with some modifications. Briefly, HFL cell pellet was resuspended in 500μL of 1X lysis buffer (#9803, Cell Signaling) for 30min at 4°C, vortex every 10 min, and then the suspension was pelleted at 14000 x g for 10 minutes at 4°C. The supernatant was precleared with 25 μL of lysis-buffer-washed binding control agarose beads (#bab-20, chromotek) rotating at 4°C for 1hr. Non-specific aggregates were removed by centrifugation at 2500 x g for 5 mins and 5% of the supernatant was used as the input. Remaining supernatant was incubated with 25 μL of lysis-buffer-washed Halo-Trap agarose beads rotating at 4°C for 1 hr. Halo-trap beads were washed thrice with 500 μL of lysis buffer and fractionated by SDS-PAGE. Immunoblotting was done using mouse-anti FKBP (#sc-136962, Santa cruz), rabbit-anti BRD4 (#13440S, Cell Signaling), mouse-anti Plk1 (#17056, Abcam) and rabbit-anti CDK9 (#2316, Cell signaling), rabbit-anti phospho-Rpb1 CTD Ser2 (#13499, Cell Signaling)

### CellTiterGlo viability assay

Cells were seeded in 25ul growth media in poly-d lysine coated black clear-bottom 384-well plates at 150 cells/well for continuous treatment or 300 cells/well for washout treatment. Following seeding, plates were spun at 300 × g for 30 seconds, then equilibrated to room temperature for 30 minutes, and moved into the incubator. 24 hours after seeding, compounds were titrated in 100% DMSO and diluted in growth medium. 25ul of the compound/medium mixture was added to cells, bringing the total volume in each well to 50ul. DMSO was used as a negative control. After treatment, plates were spun at 300 × g for 30 seconds, then cultured at 37°C with 5% CO2 for 7 days in a humidified tissue culture incubator. For wash out experiments, treatment continued for 4 hours, following which the compound/media was flicked out and wells washed with 75ul growth medium. Fresh growth medium was added, the plates spun, and cells cultured for 7 days. For competition experiments, 10ul competition compound was added 1 hour prior to treating with 15ul test compound, maintaining a total volume of 50ul. On Day 7 of treatment, cell viability was quantified with CellTiter-Glo 2.0 reagent (Promega). Plates were equilibrated to room temperature for 30 minutes, then 25ul of CellTiter-Glo 2.0 reagent was added to cells, bringing the total volume in each well to 75ul. After reagent was added, plates were spun at 3000 x g for 30 seconds then mixed on a shaker for two minutes at 500rpm and then incubated at room temperature for 10 minutes. Following incubation, plates were spun at 300 x g for 30 seconds, sealed with an optical adhesive cover, and luminescence readings were measured with an EnVision Plate Reader. Data was normalized to 0 luminescence for baseline. A four-parameter non-linear regression curve fit was applied to dose-response data in GraphPad Prism data analysis software to determine the half maximal growth inhibitory concentration (GI50) for each compound.

### Biophysical co-operativity assay

An AlphaLISA-based competition assay was developed to assess compound binding to BRD4 BD1 and cooperativity. All samples were assayed in triplicate. All incubations were performed at room temperature, and in low light condition after the addition of AlphaLISA beads (PerkinElmer). 20 nM of His-tagged BRD4 BD1 and 20 nM of biotinylated histone H4 K5/8/12/16(Ac) peptide (AnaSpec) were mixed in 20 mM HEPES pH 7.5, 100 mM NaCl, 0.5 mM TCEP, 0.1% BSA, 0.02% CHAPS, and were added into a 384-well Alpha plate (PerkinElmer). Compounds were serially diluted in the same buffer containing 1% DMSO, in the absence and presence of 8 μM of FKBP12 F36V. The compound samples were incubated for 30 min and were added to the plate. The plate was sealed and incubated for 60 min. After incubation, 10 μg/mL of streptavidin acceptor beads were added to the plate, followed by a 30 min incubation. 10 μg/mL of nickel chelate donor beads were added subsequently, followed by a final 60 min incubation. AlphaLISA signals were detected by using an EnVision plate reader. Data analysis was performed by converting raw data to percent values normalized to DMSO.

### Biophysical binding constants

BRD4-BD1, BRD4-BD2, as well as CDK1/2/4/5/6/9 and PLK1 binding dissociation constants were determined at Eurofins Discoverx.

### Intracellular accumulation

Cells were treated with indicated compounds in 6-well plates for 4h and harvested by trypsinization. Next, cell pellets were washed 3x with 1mL PBS in microfuge tubes. All the residual PBS was aspirated and pellets were weighed. Pellets weighing >10mg were used for downstream analysis. *Sample preparation:* An appropriate volume of methanol:water 1:1 (v/v), typically with a ratio of 4 μL per mg of cell pellet, was added into each unknown cell pellet sample. The mixture was vortexed to homogenize. 20 μL of the unknown cell pellet homogenate was pipetted out into a well in 96-well plates for analysis. After spiking in 20 μL of DMSO:acetonitrile 1:1 (v/v) and 20 μL of 200 ng/mL propranolol and 500 ng/mL diclofenac in methanol:water 1:1 (v/v) as internal standard, 150 μL of chilled acetonitrile was added to precipitate protein. Samples were vortexed and centrifuged at 3500 rpm for 10 min. The supernatant, typically 2 μL, was injected for LC-MS/MS analysis. For calibration standard sample preparation, 20 μL of unknown sample homogenate was replaced with 20 μL of blank cell pellet homogenate with the same solvent volume/cell pellet weight ratio (μL/mg) as in the unknown cell pellet homogenate preparation. 20 μL of DMSO:acetonitrile 1:1 (v/v) was replaced with 20 μL of a standard working solution in DMSO:acetonitrile 1:1 (v/v). Otherwise, the calibration standard sample preparation was the same as the unknown cell pellet sample preparation. The solvent volume/cell pellet weight ratio (μL/mg) in cell pellet homogenization, the volume of DMSO:acetonitrile 1:1 (v/v) or the volume of standard working solution in DMSO:acetonitrile 1:1 (v/v), the volume of acetonitrile added to precipitate protein, and the injection volume may be adjusted to achieve different quantification limit. The concentrations of standard working solutions were usually at 0.3, 3, 30, 300, and 3000 nM, which may by adjusted for different quantification limits and calibration ranges. *LC-MS/MS Analysis:* Reverse phase chromatography was carried out on a Shimadzu HPLC system with LC-20AD pumps and a SIL-20AC HT autosampler. Separation was performed on an Agilent Zorbax Extend-C18 column (5 μm, 80 Å, 2.1×50 mm) with 0.1% acetic acid 1 mM ammonium in acetonitrile:water 1:9 (v/v) as mobile phase A and 50 mM acetic acid in acetonitrile as mobile phase B at a total flow rate of 0.75 mL/min. In a typical gradient elution, the initial % B is 0. In one minute, % B was increased linearly to 95%. After holding at 95% B for 0.5 min, % B was dropped to 0% and the column was equilibrated with 0% B for 0.5 min to prepare for the next injection. The eluent from the HPLC column was introduced to the TurboIon Spray source attached to an AB Sciex Triple Quad 5500 mass spectrometer, which operated in positive or negative mode, depending on the nature of a test compound. Multiple reaction monitoring (MRM) was used for the detection of test compounds and internal standards (260.1/116.1 for propranolol in positive ion mode; 293.8/249.9 for diclofenac in negative ion mode). The mass spectrometer was set at unit resolution with curtain gas at 30 psi, IS (ionspray voltage) at 5000 V (−4500 V in negative ion mode), TEMP (source temperature) at 500 °C, GS1 (nebulizer gas) at 50 psi, GS2 (heating gas) at 50 psi, and CAD (collision gas) at 6. The other parameters, DP (declustering potential), EP (entrance potential), CE (collision energy), and CXP (collision cell exit potential), were compound dependent and were optimized for each compound. The DP, EP, CE, and CXP values are 66 V, 8 V, 23 eV, and 6 V, respectively, for propranolol in positive ion mode. In negative ion mode, the DP, EP, CE, and CXP values are -50 V, -10 V, -17 eV, and -13 V, respectively, for diclofenac.

### Immunofluorescence

293-derived HFL, NLS2HF, and MYRHF cells were seeded at 5×10^5^ cells per well in Nunc™ Lab-Tek™ II CC2™ Chamber (#154941, Thermo Scientific) in 500 μL complete DMEM medium. After 48 hours, cells were treated with 0.1 μM TAMRA (H-789) for 1 hr at 37 °C. Following treatment, cells were washed on the chamber twice with 1xPBS. 4% formaldehyde (#12606P, Cell Signaling) was added to fix cells for 15 min at room temperature, followed with 0.1% TritonX-100 permeabilization for 10min. After washing with 1xPBS for three times, 300nM Alexa Fluor® 647 Phalloidin and DAPI in 1xPBS were added to the cells for 30 min at room temperature. Cells were washed three times with 1xPBS, followed by adding 5uL of ProLong™ Gold. Slides were removed from the chamber and the cells covered overnight with coverslips (#7816, LabScientific). Cells were imaged on a Zeiss (Oberkochen, Germany) LSM 880 Airyscan confocal microscope.

---

**Supplementary Fig. 1.**
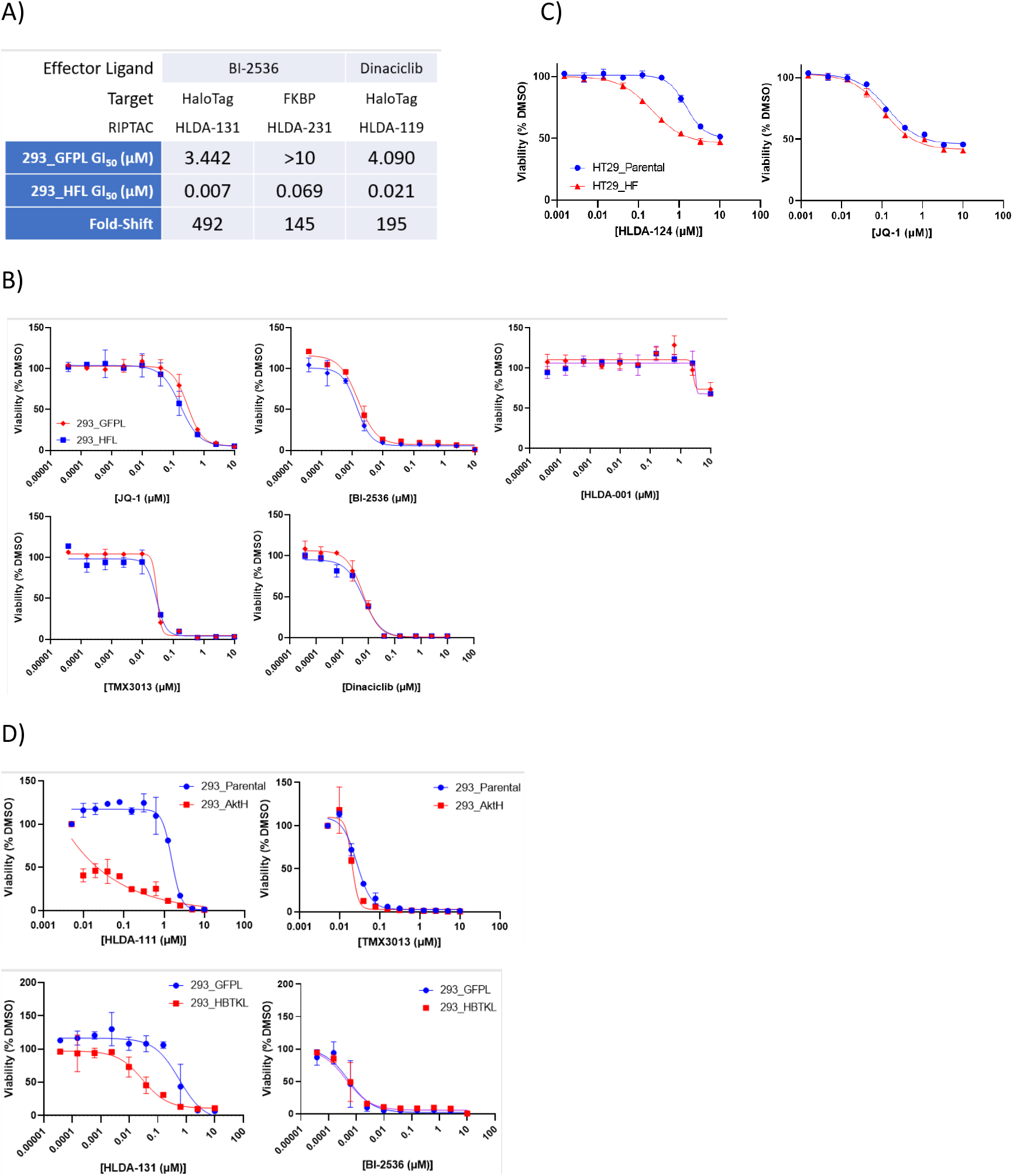
a) Differential anti-proliferative activity of RIPTACs incorporating BI-2536 and dinaciclib ELs in 293_HFL cells under 4h pulse/7 day chase conditions in a CellTiterGlo assay. b) ELs and the FKBP ligand HLDA-001 show no differential anti-proliferative activity in 293_HFL cells in a 7 day CellTiterGlo assay. c) Differential biology observed with the covalent JQ1-CA RIPTAC HLDA-124 in a 3 day CellTiterGlo assay in HT29_HFL cells that express the HaloTag-FKBP target protein. d) Differential anti-proliferative activity observed in a 5 day CellTiterGlo assay with the TMX3013-CA RIPTAC HLDA-111 in 293_AktH cells overexpressing Akt1-HaloTag but not the TMX3013 EL. Similarly, differential anti-proliferative activity observed in a 5 day CellTiterGlo assay with the BI2536-CA RIPTAC HLDA-131, but not with BI2536 alone, in the HaloTag-BTK-overexpressing 293_HBTKL cells compared to 293_GFPL cells.

**Supplementary Fig. 2.**
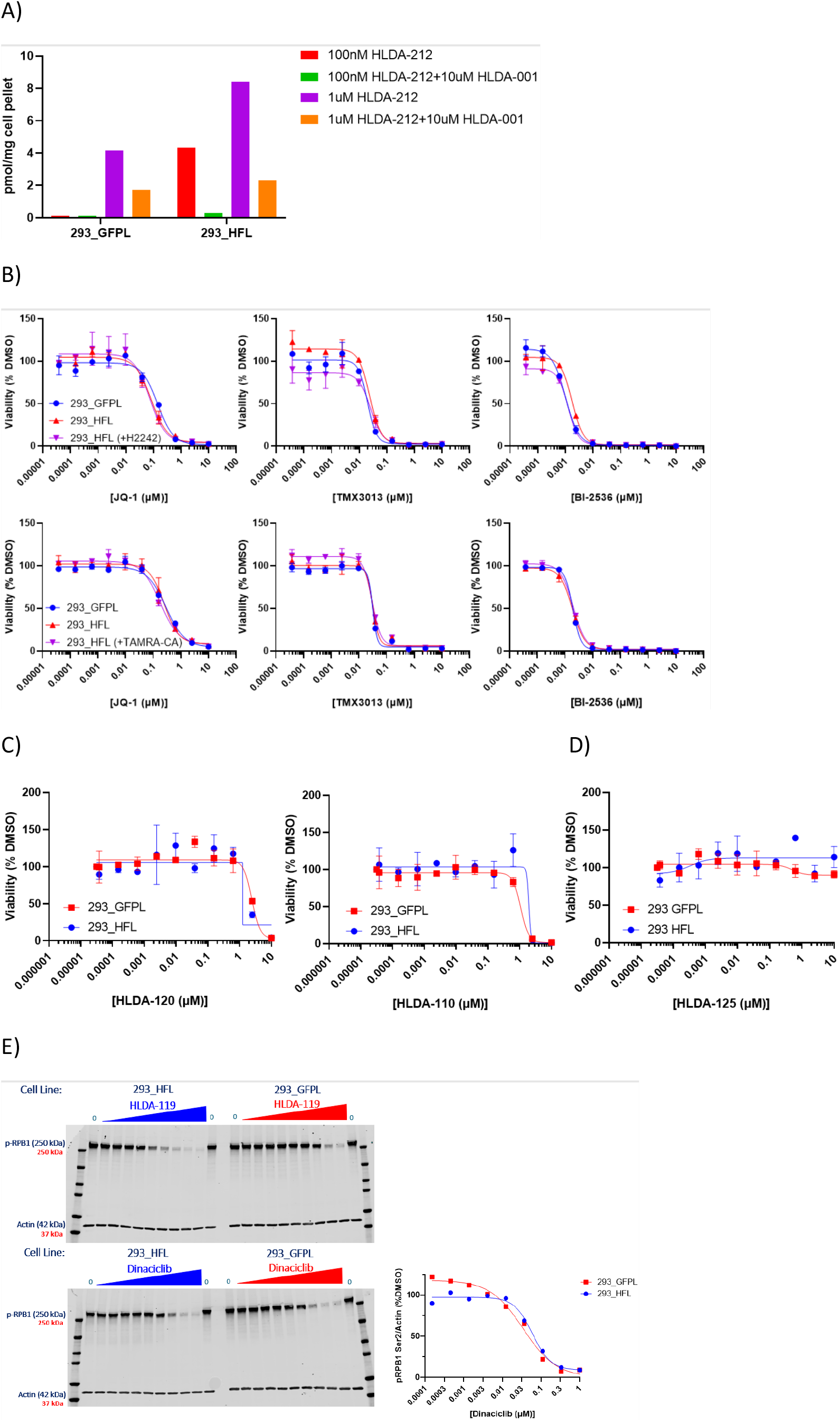
a) Pre-treatment with 10μM FKBP ligand HLDA-001 abrogates intracellular accumulation of RIPTAC HLDA-212. b) ELs JQ1, TMX3013, and BI2536 are equipotent in 293_GFPL and 293_HFL cells and show no reduction in potency upon pre-treatment of 293_HFL cells with either 10μM FKBP ligand HLDA-001 or 300nM HaloTag ligand TAMRA-CA. c) HLDA-120 and HLDA-110, des-chloro negative controls for JQ1-CA HLDA-124 and TMX3013-CA HLDA-111 respectively, are equipotent in 293_GFPL and 293_HFL cells. d) HLDA-125, a negative control RIPTAC, that cannot bind BRD2/3/4, shows no differential potency in 293_HFL cells. e) Dinaciclib is equipotent in 293_GFPL and 293_HFL cells, as determined by its effect on phospho-RPB1 CTD Ser2 levels in cells.

**Supplementary Fig. 3.**
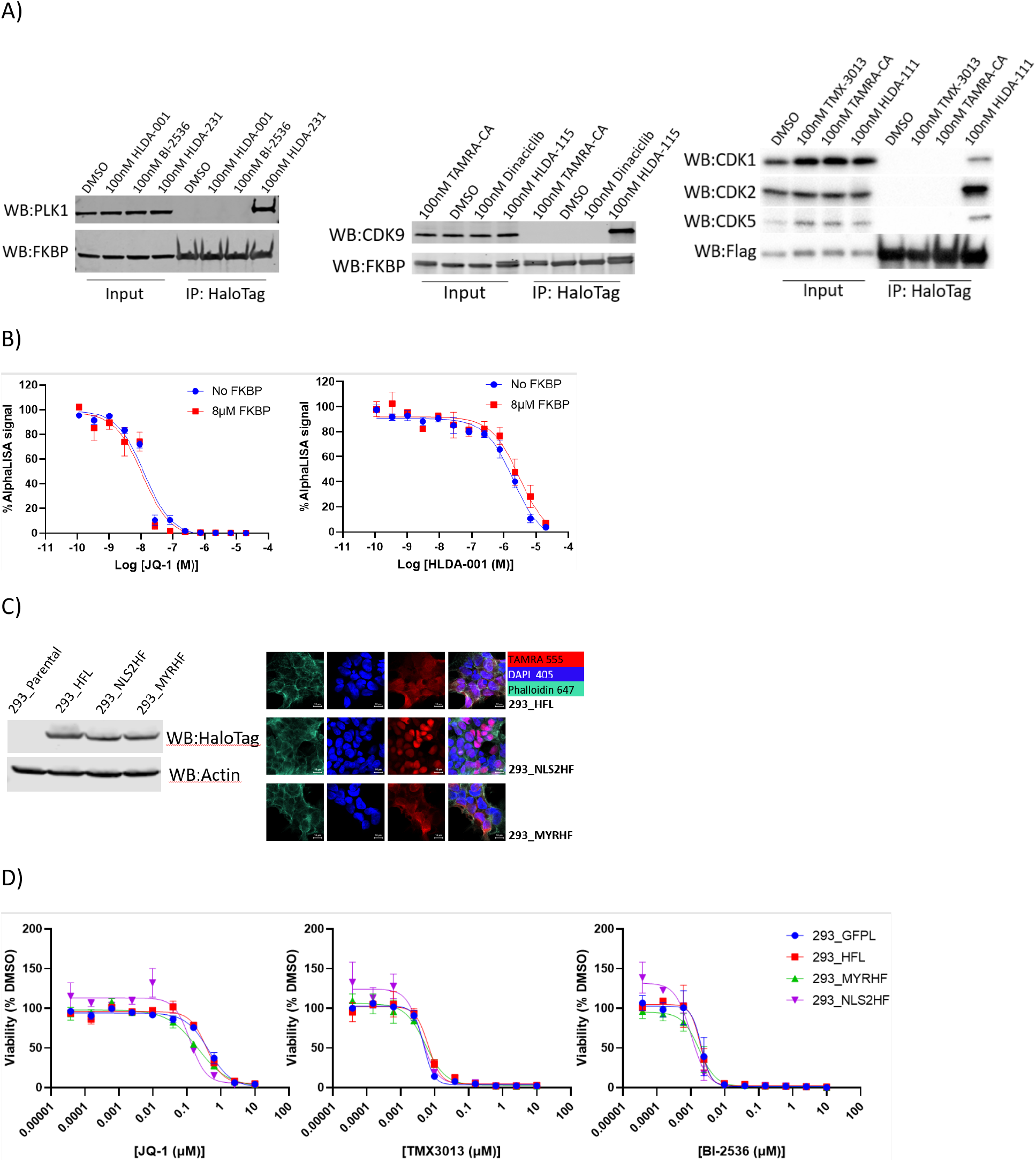
a) Ternary complex formation observed with PLK1 by treating 293_HFL cells with the BI2536 containing non-covalent RIPTAC HLDA-231, with CDK9 by treating with the dinaciclib-CA RIPTAC HLDA-115, and with CDK1/2/5 with the TMX3013-CA RIPTAC HLDA-111. No complex formation is observed with EL and TL controls. b) No positive co-operativity observed in AlphaLISA with TL HLDA-001 or EL JQ1. c) Expression levels of HaloTag-FKBP by immunoblotting and localization by confocal microscopy in 293_NLS2HF and 293_MYRHF cells compared to 293_HFL cells. d) ELs are equipotent across 293_HFL, 293_NLS2HF, and 293_MYRHF cells in a 7-day CellTiter Glo assay.

**Supplementary Fig. 4.**
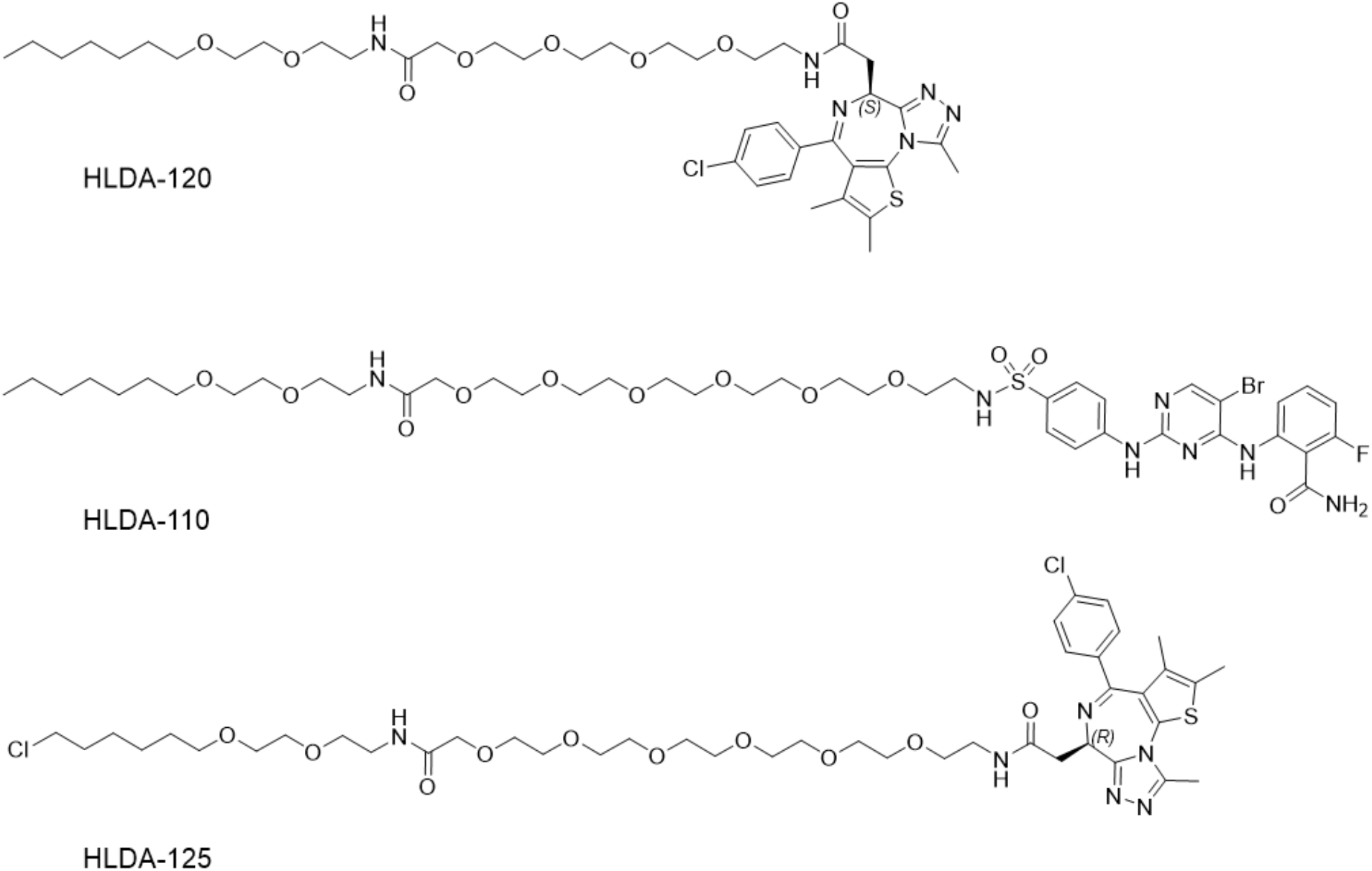
a) Structures of negative control compounds HLDA-120, HLDA-110, and HLDA-125

