## supplemental Table and Methods for "Regulated Induced Proximity Targeting Chimeras (RIPTACs): a Novel Heterobifunctional Small Molecule Therapeutic Strategy for Killing Cancer Cells Selectively"

**SUPPLEMENTARY TABLE**

| **Compound** | **TL** | **EL** | **Kinomescan/Bromoscan Data** |
| --- | --- | --- | --- |
| HLDA-002866 | chlorotag | BRD4 | BROMOScan BRD4-BD1 KD (uM) 0.0210  BROMOScan BRD4-BD2 KD (uM) 0.0086 |
| HLDA-002867 | chlorotag | BRD4 | BROMOScan BRD4-BD1 KD (uM) 0.02  BROMOScan BRD4-BD2 KD (uM) 0.0063 |
| HLDA-002864 | chlorotag | BRD4 | BROMOScan BRD4-BD1 KD (uM) 0.0285  BROMOScan BRD4-BD2 KD (uM) 0.0071 |
| HLDA-003288 | FKBP | BRD4 | BROMOScan BRD4-BD1 KD (uM) 0.11  BROMOScan BRD4-BD2 KD (uM) 0.038 |
| HLDA-003289 | FKBP | BRD4 | BROMOScan BRD4-BD1 KD (uM) 0.091  BROMOScan BRD4-BD2 KD (uM) 0.02 |
| HLDA-003368 | FKBP | BRD4 | BROMOScan BRD4-BD1 KD (uM) 0.063  BROMOScan BRD4-BD2 KD (uM) 0.015 |
| HLDA-003196 | Chlorotag | CDK | KINOMEscan CDK1 KD (uM) 0.0770 |
| HLDA-003197 | Chlorotag | CDK | KINOMEscan CDK1 KD (uM) 0.046  KINOMEscan CDK2 KD (uM) 0.0026  KINOMEscan CDK4 KD (uM) 2.3  KINOMEscan CDK6 KD (uM) 1.1  KINOMEscan CDK9 KD (uM) 0.31 |
| HLDA-002951 | Chlorotag | CDK | KINOMEscan CDK1 KD (uM) 0.018  KINOMEscan CDK2 KD (uM) 0.0019  KINOMEscan CDK4 KD (uM) 1.2  KINOMEscan CDK6 KD (uM) 0.92  KINOMEscan CDK9 KD (uM) 0.2 |
| HLDA-003798 | FKBP | CDK |  |
| HLDA-003771 | FKBP | CDK | KINOMEscan CDK1 KD (uM) 0.98  KINOMEscan CDK9 KD (uM) 9.3 |
| HLDA-003772 | FKBP | CDK |  |
| HLDA-003519 | Chlorotag | PLK1 | KINOMEscan PLK1 KD (uM) 0.0003 |
| HLDA-003520 | Chlorotag | PLK1 |  |
| HLDA-003551 | Chlorotag | PLK1 | KINOMEscan PLK1 KD (uM) 0.0004 |
| HLDA-003714 | FKBP | PLK1 | KINOMEscan PLK1 KD (uM) 0.0570 |
| HLDA-003715 | FKBP | PLK1 | KINOMEscan PLK1 KD (uM) 0.0260 |
| HLDA-003716 | FKBP | PLK1 |  |
| HLDA-003224 | None | PLK1 | KINOMEscan PLK1 KD (uM) 6.5E-5 |
| HLDA-000420 |  | BRD4 | BRD4 BIND ALPHA IC50 BD1 (uM) 0.0174  BRD4 BIND ALPHA IC50 BD2 (uM) 0.0073 |
| HLDA-003133 | None | CDK | KINOMEscan CDK1 KD (uM) 0.0016  KINOMEscan CDK9 KD (uM) 0.32 |
| HLDA-003927 | Chlorotag | CDK | KINOMEscan CDK1 KD (uM) 1.5  KINOMEscan CDK9 KD (uM) 0.0035 |
| HLDA-003938 | Chlorotag | CDK | KINOMEscan CDK9 KD (uM) 0.0054 |
| HLDA-003939 | Chlorotag | CDK | KINOMEscan CDK1 KD (uM) 1.6  KINOMEscan CDK9 KD (uM) 0.0056 |
| HLDA-003928 | Chlorotag | CDK | KINOMEscan CDK1 KD (uM) 9.2  KINOMEscan CDK9 KD (uM) 0.0099 |
| HLDA-003940 | Chlorotag | CDK | KINOMEscan CDK9 KD (uM) 0.019 |
| HLDA-003941 | Chlorotag | CDK | KINOMEscan CDK1 KD (uM) >10  KINOMEscan CDK9 KD (uM) 0.0015 |
| HLDA-003222 |  | CDK | KINOMEscan CDK1 KD (uM) 0.725  KINOMEscan CDK2 KD (uM) 0.021  KINOMEscan CDK4 KD (uM) 0.036  KINOMEscan CDK5 KD (uM) 0.0065  KINOMEscan CDK6 KD (uM) 0.013  KINOMEscan CDK9 KD (uM) 0.0004 |

**SUPPORTING METHODS**

**Chemical Synthesis:**

**A. General considerations**. Chemicals used for synthesis were purchased from commercial

sources and were used without further purification. Flash chromatography was performed on silica gel columns. 1H NMR spectra were recorded on a Bruker 400 NMR spectrometer (400 MHz for 1H and 101 MHz for 13C). The values of chemical shifts (δ) are reported in p.p.m. Coupling constants (J) are reported in Hz. LCMS were recorded on Agilent 1100 LC and Agilent G1956A. HPLC purifications were performed on a reverse-phase column using a Shimadzu HPLC system or Agilent HPLC system.

**B. Experimental Protocols.** Chloroalkanes (*Series 1-6*), FKBP (*Series 7-10*).

***SERIES 1***

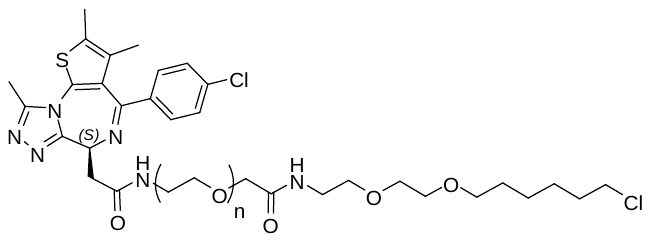

HLDA-123, n=2

HLDA-124, n=4

HLDA-121, n=6

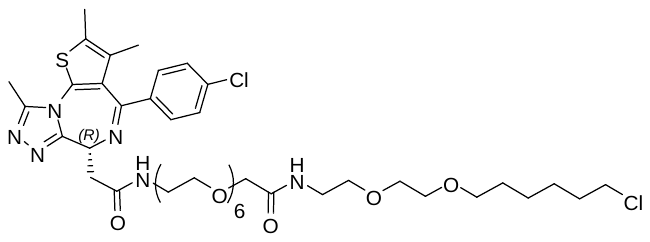

HLDA-125

The synthetic route for HLDA-123

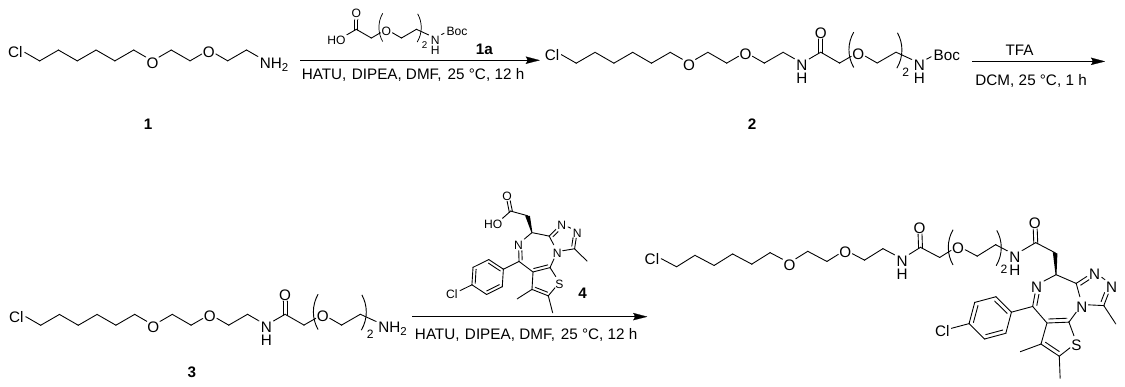

**Preparation of Compound 2**

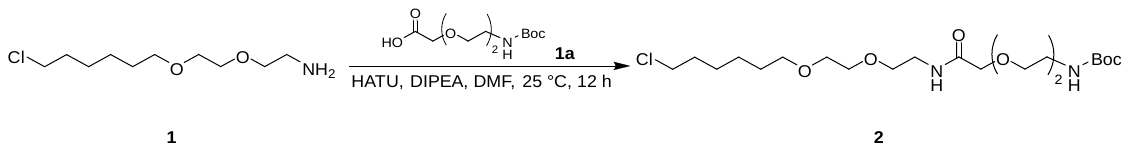

To a solution of 2-[2-[2-(*tert*-butoxycarbonylamino)ethoxy]ethoxy]acetic acid (100 mg, 380 umol, 1.0 equiv) and 2-[2-(6-chlorohexoxy)ethoxy]ethanamine (85 mg, 380 μmol, 1.0 equiv) in DMF (2 ml) was added HATU (217 mg, 570 μmol, 1.5 equiv) and DIPEA (147 mg, 1.14 mmol, 3.0 equiv). The mixture was stirred at 25 °C for 12 h. The mixture was purified by *prep*-HPLC (column: Waters Xbridge 150*25mm* 5μm; mobile phase: [water(10 mM NH_4_HCO_3_)–MeCN]; B%: 30%–60%,10 min) to afford tert-butyl *N*-[2-[2-[2-[2-[2-(6-chlorohexoxy)ethoxy]ethylamino]-2-oxo-ethoxy]ethoxy]ethyl]carbamate (150 mg, 84% yield) as a white solid.

**LC–MS:** MS (ES^+^): RT = 0.902 min, m/z = 469.2 [M + H^+^];

**Spectra:**

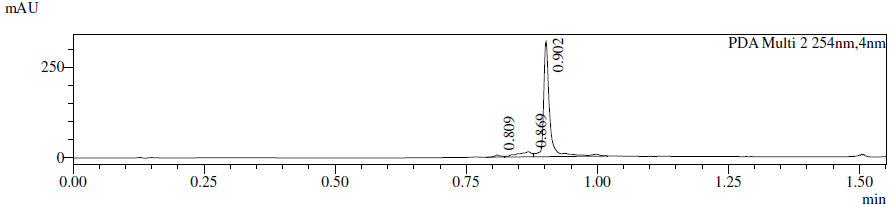

**Preparation of Compound 3**

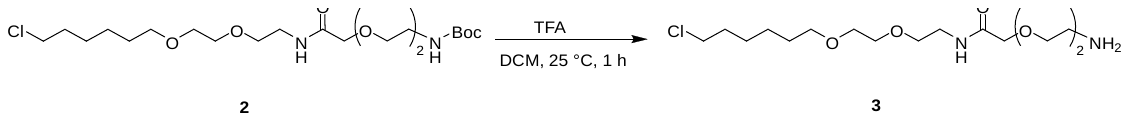

To a solution of *tert*-butyl *N*-[2-[2-[2-[2-[2-(6-chlorohexoxy)ethoxy]ethylamino]-2-oxo-ethoxy]ethoxy]

ethyl]carbamate (150 mg, 320 μmol, 1.0 equiv) in DCM (1 ml) was added TFA (0.5 ml). The mixture was stirred at 25 °C for 1 h. The reaction mixture was concentrated under reduced pressure to give 2-[2-(2-aminoethoxy)ethoxy]-*N*-[2-[2-(6-chlorohexoxy)ethoxy]ethyl]acetamide (150 mg, 97% yield, TFA salt) as a yellow oil.

**LC–MS:** MS (ES^+^): RT = 0.900 min, m/z = 369.2[M + H^+^].

**Spectra:**

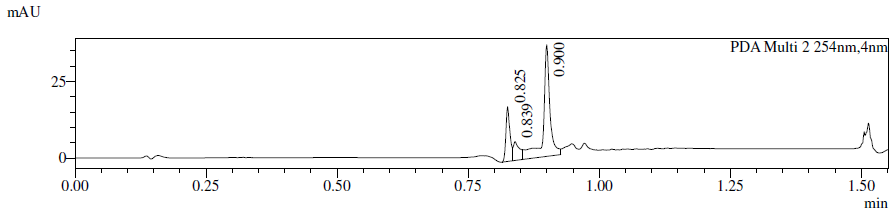

**Preparation of Compound 4**

Known compound from *J. Med. Chem*. **2022**, *65*, 6573–6592

**Preparation of HLDA-123**

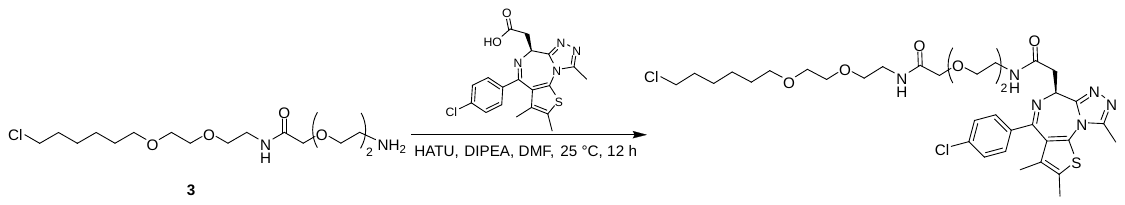

To a solution of 2-[2-(2-aminoethoxy)ethoxy]-*N*-[2-[2-(6-chlorohexoxy)ethoxy]ethyl]acetamide (150 mg, 311 umol, 1.0 equiv, TFA salt) and 2-[(9*S*)-7-(4-chlorophenyl)-4,5,13-trimethyl-3-thia-1,8,11,12-tetrazatricyclo[8.3.0.0^2,6^]trideca-2(6),4,7,10,12-pentaen-9-yl]acetic acid (125 mg, 311 μmol, 1.0 equiv) in DMF (2 ml) was added T_3_P (297 mg, 466 μmol, 50% purity, 1.5 equiv) and DIEA (200 mg, 1.55 mmol, 5.0 equiv). The mixture was stirred at 25 °C for 12 h. The reaction mixture was concentrated under reduced pressure to give a residue. The residue was purified by *prep*-HPLC (column: Waters Xbridge 150*25 mm*5 μm;mobile phase: [water(10 mM NH_4_HCO_3_)–MeCN]; B%: 44%–74%,10min) to give *N*-[2-[2-[2-[2-[2-(6-chlorohexoxy)ethoxy]ethylamino]-2-oxo-ethoxy]ethoxy]ethyl]-2-[(9*S*)-7-(4-chlorophenyl)-4,5,13-trimethyl-3-thia-1,8,11,12-tetrazatricyclo[8.3.0.0^2,6^]trideca-2(6),4,7,10,12-pentaen-9-yl]acetamide (75 mg, 32% yield) as a yellow gum.

**^1^H NMR** (400 MHz, CD_3_OD): δ 7.51–7.39 (m, 4H), 4.66 (d, 1H, *J* = 5.4 Hz), 4.02 (s, 2H), 3.72 (s, 4H), 3.68– 3.64 (m, 2H), 3.63– 3.53 (m, 8H), 3.53– 3.42 (m, 8H), 3.38– 3.31 (m, 2H), 2.71 (s, 3H), 2.47 (s, 3H), 1.80– 1.68 (m, 5H), 1.58 (t, 2H, *J* = 6.8 Hz), 1.49–1.35 (m, 4H)

**LC–MS:** MS (ES^+^): RT = 2.791 min, m/z = 751.2 [M + H^+^].

**Spectra:**

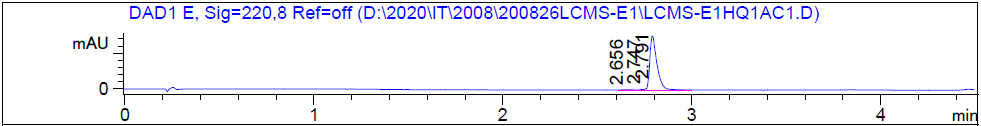

HRMS [C_35_H_48_Cl_2_N_6_O_6_S] Cal: 751.2806; Obs: 751.2776

**The synthetic route for HLDA-124**

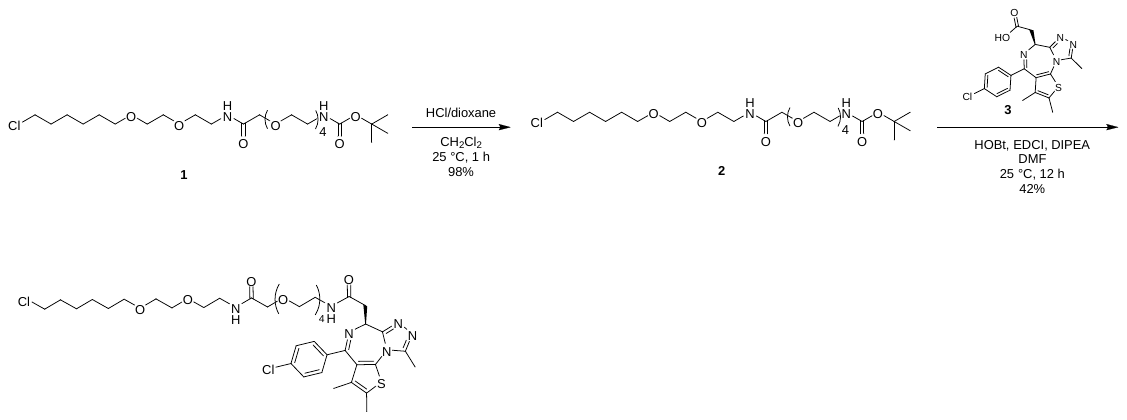

**Preparation of Compound 1**

See HLDA-113

**Preparation of Compound 2**

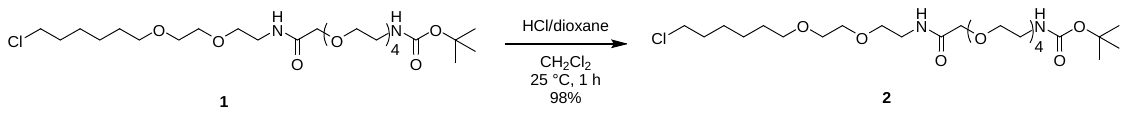

A mixture of *tert*-butyl *N*-[2-[2-[2-[2-[2-[2-[2-(6-chlorohexoxy)ethoxy]ethylamino]-2-oxo-ethoxy] ethoxy]ethoxy]ethoxy]ethyl]carbamate (230 mg, 413 μmol, 1.0 equiv) in CH_2_Cl_2_ (2 ml) and 4 M HCl/dioxane (2.3 ml) was stirred at 25 °C for 1 h. The reaction mixture was concentrated under reduced pressure to afford 2-[2-[2-[2-(2-aminoethoxy)ethoxy]ethoxy]ethoxy]-*N*-[2-[2-(6- chlorohexoxy)ethoxy]ethyl]acetamide (200 mg, 405 μmol, 98% yield, HCl salt) as a yellow solid and carried on to the next step without further purification

**Preparation of Compound 3**

Known compound from *J. Med. Chem.* **2022**, *65*6573–6592

**Preparation of HLDA-124**

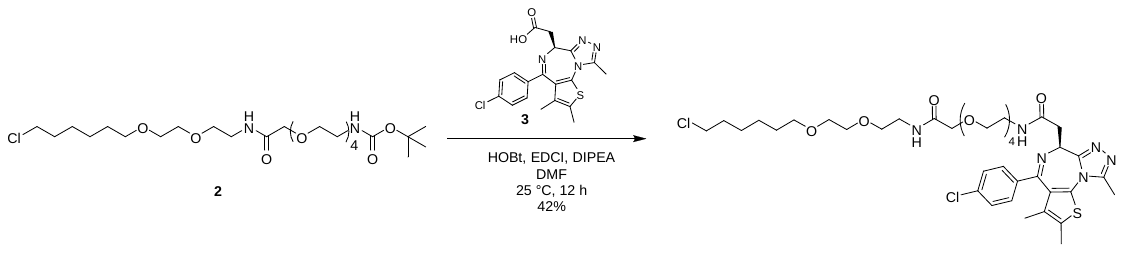

To a solution of 2-[2-[2-[2-(2-aminoethoxy)ethoxy]ethoxy]ethoxy]-*N*-[2-[2-(6-chlorohexoxy) ethoxy]ethyl]acetamide (200 mg, 0.41 mmol, 1.0 equiv, HCl salt) in DMF (2 ml) were added DIPEA (262 mg, 2.03 mmol, 0.35 ml, 5.0 equiv), HOBt (82.2 mg, 0.61 mmol, 1.5 equiv), EDCI (117 mg, 0.61 mmol, 1.5 equiv) and (9*R*)-7-(4-chlorophenyl)-4, 5,13-trimethyl-3-thia-1,8,11,12-tetrazatricyclo[8.3.0.02,6]trideca-2(6),4,7,10,12-pentaene-9-carboxylic acid (157 mg, 0.41 mmol, 1.0 equiv). The mixture was stirred at 25 °C for 12 h. The reaction mixture was diluted with water (20 mL) and the mixture was extracted with ethyl acetate (3 x 15 ml). The combined organic phase was washed with brine (10 ml), dried with anhydrous Na_2_SO_4_ and concentrated under reduced pressure. The residue was purified by *prep*-HPLC (column: Phenomenex Gemini-NX C18 75*30 mm*3um; mobile phase: [water(0.225%FA)–MeCN]; B%: 48%–78%, 7 min) to afford *N*-[2-[2-[2-[2-[2-[2-[2-(6-chlorohexoxy)ethoxy]ethylamino]-2-oxo-ethoxy]ethoxy]ethoxy]ethoxy]ethyl]-2-[(9*S*)-7-(4-chlorophenyl)-4,5,13-trimethyl-3-thia-1,8,11,12-tetrazatricyclo[8.3.0.02,6]trideca-2(6),4,7,10,12-pentaen-9-yl]acetamide (150 mg, 0.17 mmol, 42% yield) as a yellow oil.

^1^**H NMR** (400 MHz, CDCl_3_-*d*): δ 8.31–8.25 (m, 1H), 7.68–7.59 (m, 1H), 7.52–7.39 (m, 4H), 4.54–4.47 (m, 1H), 3.86 (s, 2H), 3.64–3.58 (m, 3H), 3.56 (s, 4H), 3.51–3.30 (m, 17H), 3.30–3.15 (m, 6H), 2.59 (s, 3H), 2.41 (s, 3H), 1.73–1.65 (m, 2H), 1.62 (s, 3H), 1.52–1.43 (m, 2H), 1.41–1.26 (m, 4H).

**LC–MS:** MS (ES^+^): RT = 3.260 min, m/z = 839.3 [M + H^+^].

**Spectra:**

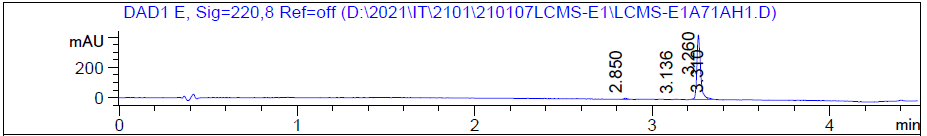

HRMS [C_39_H_56_Cl_2_N_6_O_8_S] Cal: 839.333; Obs: 839.3295

**The synthetic route for HLDA-121**

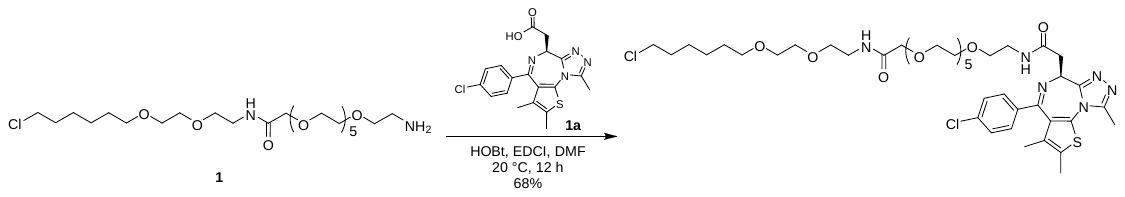

**Preparation of Compound 1**

See HLDA-133

**Preparation of Compound 1a**

Known compound from *J. Med. Chem.* **2022**, *65,* 6573–6592

**Preparation of HLDA-121**

A mixture of 2-[(9*S*)-7-(4-chlorophenyl)-4,5,13-trimethyl-3-thia-1,8,11,12-tetrazatricyclo[8.3.0.0^2,6^]trideca-2(6),4,7,10,12-pentaen-9-yl]acetic acid (100 mg, 249 umol, 1.0 equiv), 2-[2-[2-[2-[2-[2-(2-aminoethoxy)ethoxy]ethoxy]ethoxy]ethoxy]ethoxy]-*N*-[2-[2-(6-chlorohexoxy)ethoxy]ethyl]acetamide (164 mg, 249 umol, 1.0 equiv, TFA salt), HOBt (67.4 mg, 499 μmol, 2.0 equiv), EDCI (143 mg, 748 umol, 3.0 equiv) and DIEA (161 mg, 1.25 mmol, 5.0 equiv) in DMF (8 ml) was stirred at 20 °C for 12 h under N_2_. The reaction mixture was diluted with water (20 ml) and the mixture was extracted with ethyl acetate (3 x 15 ml). The combined organic phase was washed with brine (10 m), dried with anhydrous Na_2_SO_4_, filtered and concentrated under reduced pressure. The residue was purified by *prep*-HPLC (column: Phenomenex Synergi C18 150*25 mm* 10 µm; mobile phase: [water(0.225%FA)–MeCN]; B%: 48%–78%,10 min), and then *prep*-HPLC ( column: Phenomenex Synergi C18 150*25 mm* 10 µm; mobile phase: [water(0.225%FA)–MeCN]; B%: 48%–78%,10 min) to afford *N*-[2-[2-[2-[2-[2-[2-[2-[2-[2-(6-chlorohexoxy)ethoxy]ethylamino]-2-oxo-ethoxy]ethoxy]ethoxy]ethoxy]ethoxy]ethoxy]ethyl]-2-[(9*S*)-7-(4-chlorophenyl)-4,5,13-trimethyl-3-thia-1,8,11,12-tetrazatricyclo[8.3.0.0^2,6^]trideca-2(6),4,7,10,12-pentaen-9-yl]acetamide (159 mg, 170 μmol, 68% yield) as a colorless oil.

**^1^H NMR** (400 MHz, DMSO-d_6_) δ 8.28 (t, 1H, *J* = 5.2 Hz), 7.63 (t, 1H, *J* = 5.6 Hz), 7.52–7.47 (m, 2H), 7.46–7.40 (m, 2H), 4.54–4.48 (m, 1H), 3.87 (s, 2H), 3.62 (t, 2H, *J* = 6.8 Hz), 3.58–3.41 (m, 28H), 3.39–3.35 (m, 2H), 3.30–3.21 (m, 6H), 2.60 (s, 3H), 2.42 (s, 3H), 1.76-1.66 (m, 2H), 1.63 (s, 3H), 1.53–1.43 (m, 2H), 1.43–1.26 (m, 4H).

**LC–MS:** MS (ES^+^): RT = 2.897 min, m/z = 464.4 [M/2 + H^+^].

**Spectra:**

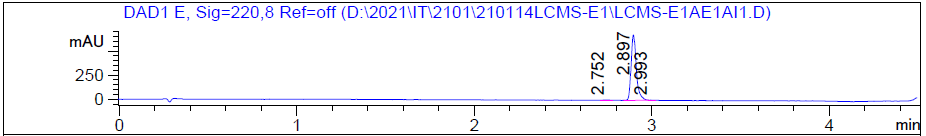

HRMS [C_43_H_64_Cl_2_N_6_O_10_S] Cal: 927.3854; Obs: 927.3826

**The synthetic route for HLDA-125**

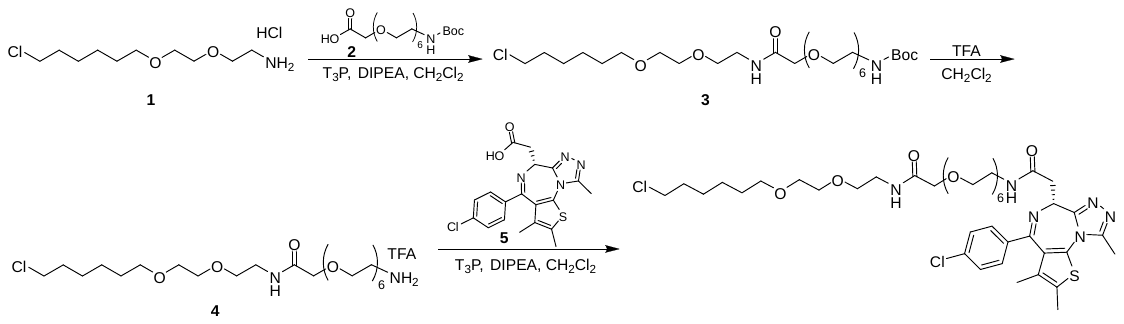

**Preparation of Compound 3**

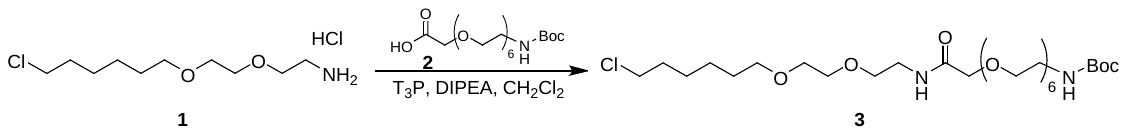

A mixture of 2-(2-((*tert*-butoxycarbonyl)amino)ethoxy)acetic acid (160 mg, 0.36 mmol, 1 equiv), 2-(2-((6-chlorohexyl)oxy)ethoxy)ethan-1-amine (94.7 mg, 0.36 mmol, 1 equiv), DIPEA (235 mg, 1.82 mmol, 0.32 ml, 5 equiv) and T_3_P (348 mg, 0.55 mmol, 0.32 ml, 50% purity, 1.5 equiv) in CH_2_Cl_2_ (5 ml) was stirred at 25 °C for 3 h. The reaction mixture was added water (20 ml) and extracted with EtOAc (2 x 20 ml). The combined organic layer was washed with brine (2 x 20 ml), dried with anhydrous Na_2_SO_4_, filtered and concentrated under reduced pressure. The residue was purified by prep-TLC (CH_2_Cl_2_/MeOH = 10/1) to afford *tert*-butyl (18-chloro-5-oxo-3,9,12-trioxa-6-azaoctadecyl)carbamate (180 mg, 0.28 mmol, 77% yield) as a yellow oil.

**^1^H NMR** (CDCl_3_, 400 MHz) δ 7.21 (brs, 1H), 4.01 (s, 2H), 3.42–3.71 (m, 34H), 3.31 (t, 2H, *J* = 5.2 Hz), 1.74–1.81 (m, 2H), 1.55–1.65 (m, 2H), 1.33–1.50 (m, 13H).

**LC–MS:** MS (ES^+^): RT = 0.926 min, m/z = 645.3 [M+H^+^].

**Spectra:**

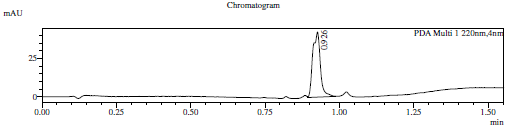

**Preparation of Compound 4**

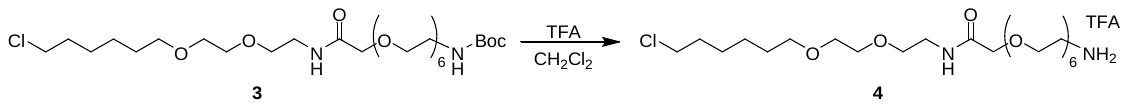

A mixture of *tert*-butyl-(18-chloro-5-oxo-3,9,12-trioxa-6-azaoctadecyl)carbamate (170 mg, 0.26 mmol, 1 equiv) in TFA (1.5 ml) and CH_2_Cl_2_ (3 ml) was stirred at 25 °C for 0.5 h. The reaction mixture was concentrated under reduced pressure to afford 2-(2-aminoethoxy)-*N*-(2-(2-((6-chlorohexyl)oxy)ethoxy)ethyl)acetamide (170 mg, 0.26 mmol, 98% yield) as a yellow oil.

**^1^H NMR** (CDCl_3_, 400 MHz) δ 7.68–7.33 (m, 4H), 4.12 (s, 2H), 3.83–3.78 (m, 2H), 3.75–3.71 (m, 2H), 3.66–3.49 (m, 30H), 3.26–3.02 (m, 2H), 1.82–1.74 (m, 2H), 1.65–1.55 (m, 2H), 1.50–1.42 (m, 2H), 1.40–1.32 (m, 2H).

**Spectra:**

**Preparation of compound 5**

Known compound from *J. Med. Chem.* **2022**, *65,* 6573–6592

**Preparation of HLDA-125**

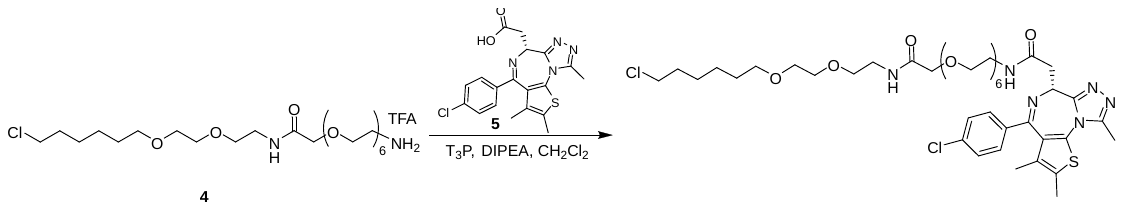

A mixture of (*R*)-2-(4-(4-chlorophenyl)-2,3,9-trimethyl-6*H*-thieno[3,2-f][1,2,4]triazolo[4,3-a][1,4]diazepin-6-yl)acetic acid (40.0 mg, 100 μmol, 1 *equiv*), 2-(2-aminoethoxy)-*N*-(2-(2-((6-chlorohexyl)oxy)ethoxy)ethyl)acetamide (65.8 mg, 100 μmol, 1 equiv), T_3_P (95.2 mg, 150 μmol, 89.0 μL, 50% purity, 1.5 equiv) and DIPEA (64.5 mg, 499 umol, 86.9 uL, 5 equiv) in CH_2_Cl_2_ (3 ml) was stirred at 25 °C for 16 h. The reaction mixture was added water (15 ml) and extracted with CH_2_Cl_2_ (2 x 15 ml). The combined organic layer was washed with brine (15 ml), dried with anhydrous Na_2_SO_4_, filtered and concentrated under reduced pressure. The residue was purified by prep-HPLC (column: Phenomenex Gemini-NX C18 75*30 mm*3 μm;mobile phase: [water(10mM NH_4_HCO_3_)–-MeCN];B%: 35%–65%, 8 min) to afford compound(*R*)-*N*-(18-chloro-5-oxo-3,9,12-trioxa-6-azaoctadecyl)-2-(4-(4-chlorophenyl)-2,3,9-trimethyl-6*H*-thieno[3,2-f][1,2,4]triazolo[4,3-a][1,4]diazepin-6-yl)acetamide (16.0 mg, 16.3 umol, 16% yield) as a yellow gum.

**^1^H NMR** (DMSO-*d*_6_, 400 MHz) δ 7.45–7.39 (m, 2H), 7.37–7.31 (m, 2H), 7.24–7.12 (m, 1H), 7.05–6.90 (m, 1H), 4.67 (t, 1H, *J* = 6.8 Hz), 3.99 (2H, s), 3.65–3.69 (m, 19H), 3.63–3.36 (m, 19H), 2.68 (s, 3H), 2.41 (s, 3H), 1.74–1.81 (m, 2H), 1.68 (s, 3H), 1.65–1.55 (m, 2H), 1.41–1.50 (m, 2H), 1.33-1.41 (m, 2H).

**LC**–**MS:** MS (ES^+^): RT = 2.799 min, m/z = 927.3 [M + H^+^];

**Spectra:**

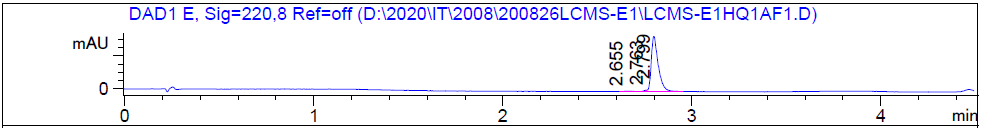

HRMS [C_43_H_64_Cl_2_N_6_O_10_S] Cal: 927.3854; Obs: 927.382

***SERIES 2***

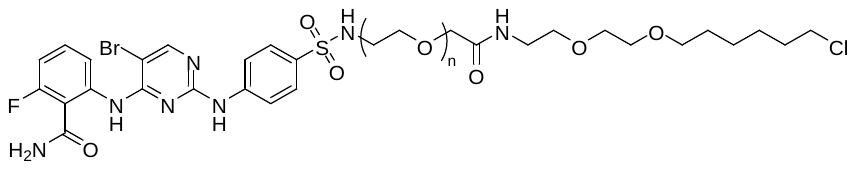

HLDA-112, n=2

HLDA-113, n=4

HLDA-111, n=6

**The synthetic route for HLDA-112**

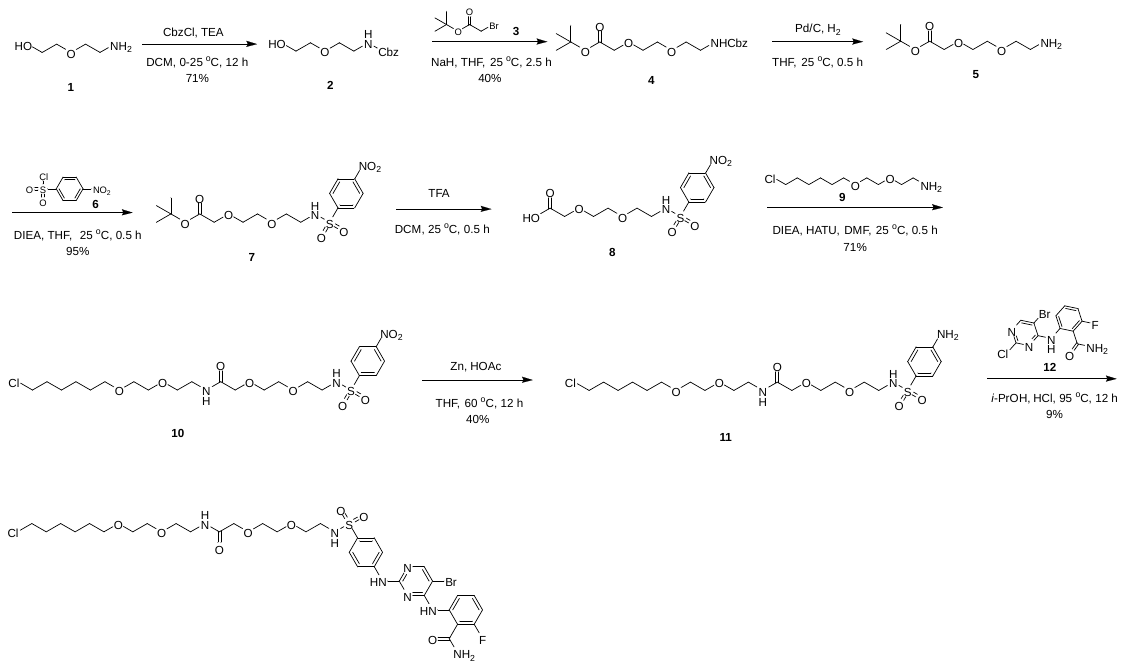

**Preparation of compound 2**

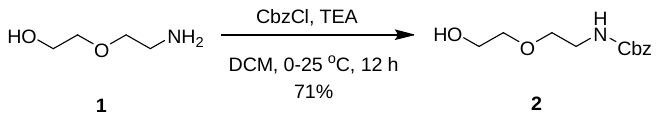

To a solution of 2-(2-aminoethoxy)ethanol (5.00 g, 47.5 mmol, 4.76 ml, 1.0 equiv) in CH_2_Cl_2_ (30 ml) was added CbzCl (9.74 g, 57.0 mmol, 8.11 ml, 1.2 equiv) and TEA (6.26 g, 61.8 mmol, 8.61 ml, 1.3 equiv) at 0 °C, and it was stirred at 25 °C for 12 h. To the reaction mixture was added water (50 ml) and the mixture was extracted with EtOAc (50 ml). The combined organic phase was washed with brine (3 x 50 ml), dried over anhydrous Na_2_SO_4_, filtered and concentrated under reduced pressure. The crude product was purified by column chromatography on silica gel (petroleum ether: EtOAc = 5:1 to 0:1) to give benzyl *N*-[2-(2-hydroxyethoxy)ethyl]carbamate (8.09 g, 33.8 mmol, 71% yield) as a colorless oil.

**LC–MS:** MS (ES^+^): RT = 0.664 min, m/z = 240.2 [M + H^+^];

**Spectra:**

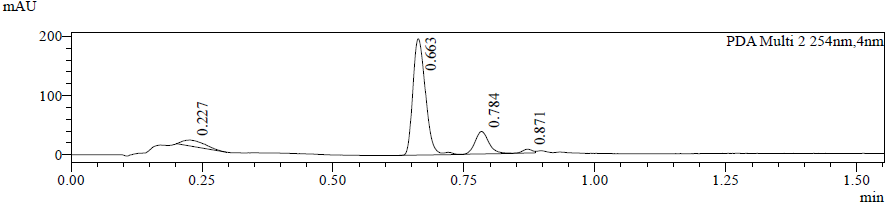

**Preparation of compound 4**

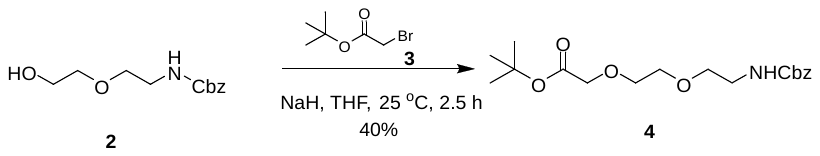

To a solution of benzyl *N*-[2-(2-hydroxyethoxy)ethyl]carbamate (8.09 g, 33.8 mmol, 1.0 equiv) in THF (40 ml) was added NaH (1.35 g, 33.8 mmol, 60% purity, 1.0 equiv) at 0 °C, then *tert*-butyl 2-bromoacetate (6.60 g, 33.8 mmol, 5.00 ml, 1.0 equiv) was added, and it was stirred at 0 °C for 30 min, then at 25 °C for 2 h. To the reaction mixture was added water (200 ml) and the mixture was extracted with EtOAc (2 x 100 ml). The combined organic phase was washed with brine (3 x 100 ml), dried over anhydrous Na_2_SO_4_, filtered and concentrated under reduced pressure. The crude product was purified by column chromatography on silica gel (petroleum ether: ethyl acetate = 3:1 to 1:1) to give *tert*-butyl 2-[2-[2-(benzyloxycarbonylamino)ethoxy]ethoxy]acetate (4.80 g, 13.5 mmol, 40% yield) as a colorless oil.

**LC–MS:** MS (ES^+^): RT = 0.865 min, m/z = 354.1 [M + H^+^];

**Preparation of compound 5**

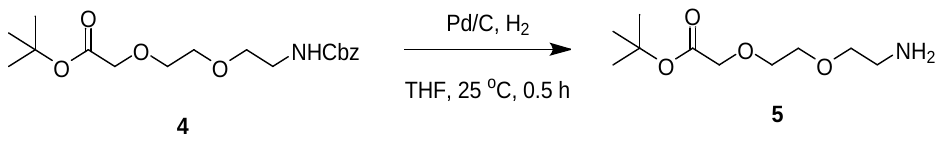

To a solution of *tert*-butyl 2-[2-[2-(benzyloxycarbonylamino)ethoxy]ethoxy]acetate (4.80 g, 13.5 mmol, 1.0 equiv) in THF (100 ml) was added Pd/C (1.2 g, 10% purity). The suspension was degassed under vacuum and purged with H_2_. The mixture was stirred under (15 psi) at 25 °C for 2 h. The reaction mixture was filtered and concentrated to give *tert*-butyl 2-[2-(2-aminoethoxy)ethoxy]acetate (2.90 g, 13.2 mmol) as a yellow oil and used directly in the next step.

**Preparation of compound 7**

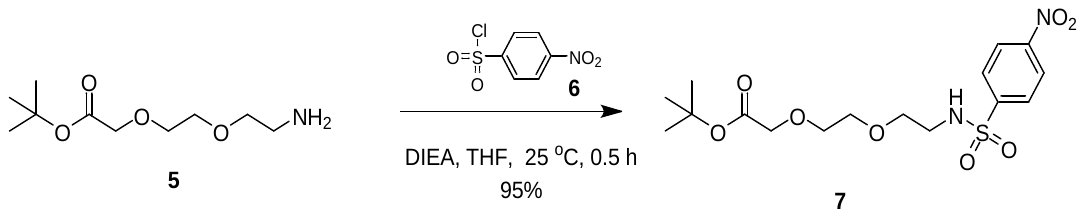

To a solution of *tert*-butyl 2-[2-(2-aminoethoxy)ethoxy]acetate (296 mg, 1.35 mmol, 1.0 equiv), DIEA (524 mg, 4.06 mmol, 707 μl, 3.0 equiv) in THF (3 ml) was added 4-nitrobenzenesulfonyl chloride (300 mg, 1.35 mmol, 1.0 equiv), and it was stirred at 25 °C for 0.5 h. H_2_O (0.1 ml) was added to quench this reaction and concentrated to afford crude product. The crude product was purified by column chromatography on silica gel (Petroleum ether: EtOAc = 5:1 to 2:1 ) to give *tert*-butyl 2-[2-[2-[(4-nitrophenyl)sulfonylamino]ethoxy]ethoxy]acetate (520 mg, 1.29 mmol, 95% yield) as a colorless oil.

**LC-MS:** MS (ES^+^): RT = 0.899 min, m/z = 303.1 [M – 100 + H^+^];

**Spectra:**

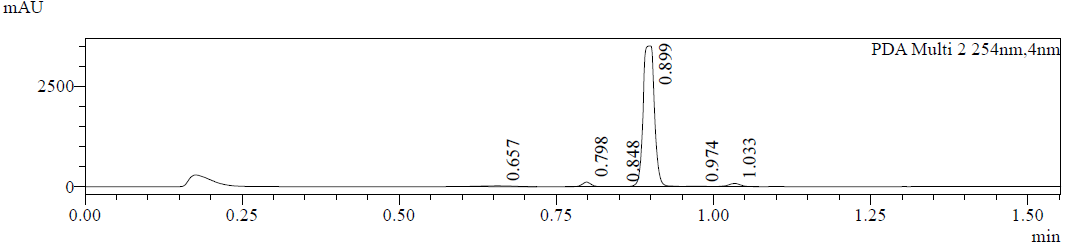

**Preparation of compound 8**

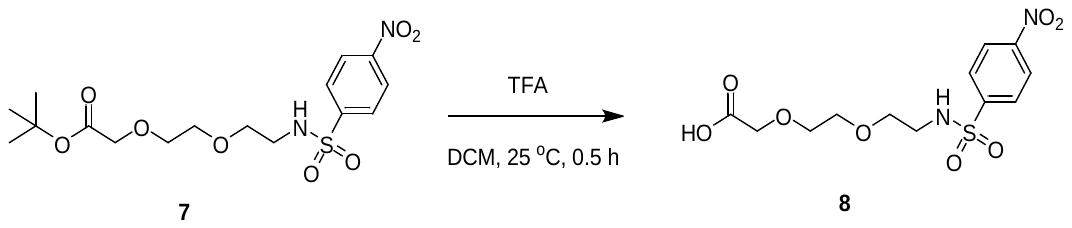

To a solution of tert-butyl 2-[2-[2-[(4-nitrophenyl)sulfonylamino]ethoxy]ethoxy]acetate (520 mg, 1.29 mmol, 1.0 equiv) in DCM (2 ml) was added TFA (2 ml), and it was stirred at 25 °C for 0.5 h. The reaction mixture was concentrated to give 2-[2-[2-[(4-nitrophenyl)sulfonylamino]ethoxy]ethoxy]acetic acid (440 mg, 1.26 mmol) as a yellow oil and it was used directly in the next step.

**Preparation of compound 10**

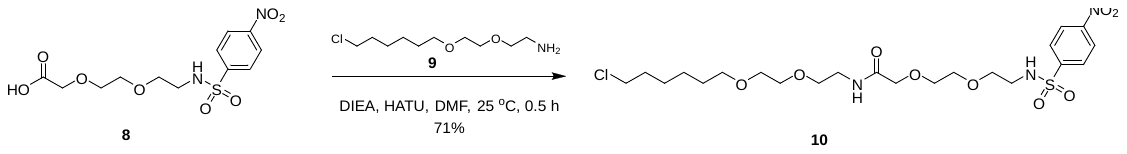

To a solution of 2-[2-[2-[(4-nitrophenyl)sulfonylamino]ethoxy]ethoxy]acetic acid (220 mg, 631 μmol, 1.0 equiv), 2-[2-(6-chlorohexoxy)ethoxy]ethanamine (164 mg, 631 μmol, 1.0 equiv, HCl salt) and DIEA (408 mg, 3.16 mmol, 550 μL, 5.0 equiv) in DMF (3 ml) was added HATU (240 mg, 631 μmol, 1.0 equiv), and it was stirred at 25 °C for 0.5 h. To the reaction mixture was added water (50 ml) and the mixture was extracted with EtOAc (50 ml). The combined organic phase was washed with brine (3 x 50 ml), dried over anhydrous Na_2_SO_4_, filtered and concentrated under reduced pressure. The residue was purified by *prep*-TLC on silica gel (petroleum ether : EtOAc = 0 :1 ) to give *N*-[2-[2-(6-chlorohexoxy)ethoxy]ethyl]-2-[2-[2-[(4-nitrophenyl)sulfonylamino]ethoxy]ethoxy]acetamide (250 mg, 451 μmol, 71% yield) as a yellow oil.

**LC–MS:** MS (ES^+^): RT = 0.946 min, m/z = 554.4 [M + H^+^];

**Spectra:**

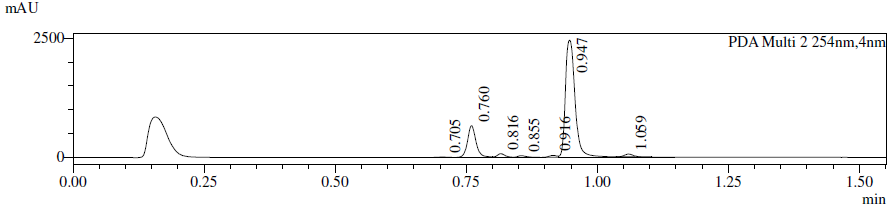

**Preparation of compound 11**

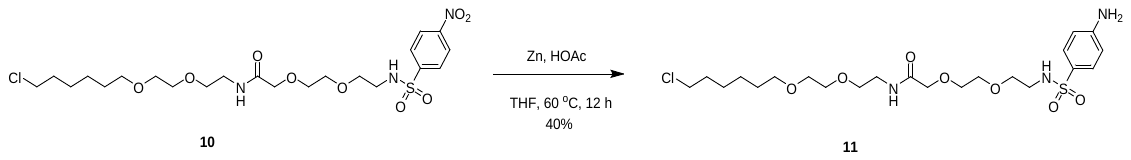

To a solution of *N*-[2-[2-(6-chlorohexoxy)ethoxy]ethyl]-2-[2-[2-[(4-nitrophenyl)sulfonylamino]ethoxy]

ethoxy]acetamide (250 mg, 451 μmol, 1.0 equiv) in THF (3 ml) and HOAc (0.6 ml) was added Zn (295 mg, 4.51 mmol, 10.0 equiv), and then it was stirred at 60 °C for 12 h. The pH was adjusted to 7–8 by sat.NaHCO_3_, and water (50 ml) was added. After extracted with EtOAc (50 ml), the organic phase was washed with brine (3 x 50 ml), dried over anhydrous Na_2_SO_4_, filtered and concentrated under reduced pressurre. The residue was purified by *prep*-TLC on silica gel (EtOAc) to give 2-[2-[2-[(4-aminophenyl)sulfonylamino]

ethoxy]ethoxy]-*N*-[2-[2-(6-chlorohexoxy)ethoxy]ethyl]acetamide (95.0 mg, 181 μmol, 40% yield) as a yellow oil.

**LC–MS:** MS (ES^+^): RT = 0.616 min, m/z = 524.3 [M + H^+^];

**Spectra:**

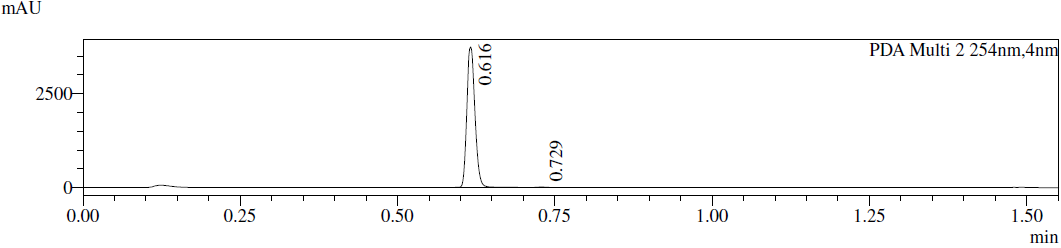

**Preparation of HLDA-112**

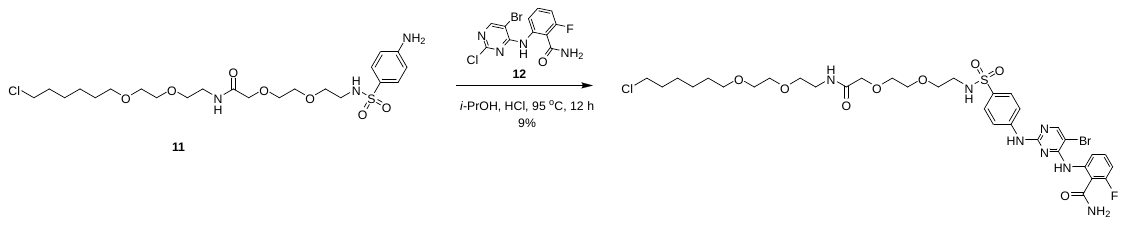

To a solution of 2-[2-[2-[(4-aminophenyl)sulfonylamino]ethoxy]ethoxy]-*N*-[2-[2-(6-chlorohexoxy)ethoxy]ethyl]acetamide (95.0 mg, 181 μmol, 1.0 equiv), 2-[(5-bromo-2-chloro-pyrimidin-4-yl)amino]-6-fluoro-benzamide (75.1 mg, 217 μmol, 1.2 equiv) in *i*-PrOH (2 ml) was added HCl (17.8 mg, 181 μmol, 17.5 μl, 37% purity, 1.0 equiv), and the mixture was stirred at 95 °C for 1 h. After this time H_2_O (0.1 ml) was added. The reaction was purified by *prep*-HPLC (column: 3_Phenomenex Luna C18 75*30 mm*3 μm; mobile phase: [water(0.05%HCl)–ACN]; B%: 44%–64%, 6.5 min) and *prep*-HPLC (column: Waters Xbridge 150*25 mm*5 μm; mobile phase: [water(10 mM NH_4_HCO_3_)–MeCN]; B%: 42%–72%, 8 min) to afford 2-[[5-bromo-2-[4-[2-[2-[2-[2-[2-(6-chlorohexoxy)ethoxy]ethylamino]-2-oxo-ethoxy]ethoxy]ethylsulfamoyl]anilino]pyrimidin-4-yl]amino]-6-fluoro-benzamide (13.9 mg, 16.6 μmol, 9% yield) as a white solid.

**^1^H NMR** (400 MHz, CD_3_OD) δ 8.37 (d, *J* = 8.3 Hz, 1H), 8.28 (s, 1H), 7.85 (d, *J* = 8.8 Hz, 2H), 7.72 (d, *J* = 8.8 Hz, 2H), 7.52 (dt, *J* = 6.4, 8.4 Hz, 1H), 7.00 (dd, *J* = 8.4, 10.4 Hz, 1H), 3.95 (s, 2H), 3.62 (dd, *J* = 2.6, 5.9 Hz, 2H), 3.59–3.49 (m, 12H), 3.45–3.38 (m, 4H), 3.07 (t, *J* = 5.6 Hz, 2H), 1.80 - 1.66 (m, 2H), 1.60–1.49 (m, 2H), 1.47– 1.26 (m, 4H)

**LC–MS:** MS (ES^+^): RT = 2.719 min, m/z = 834.2 [M + H^+^];

**Spectra:**

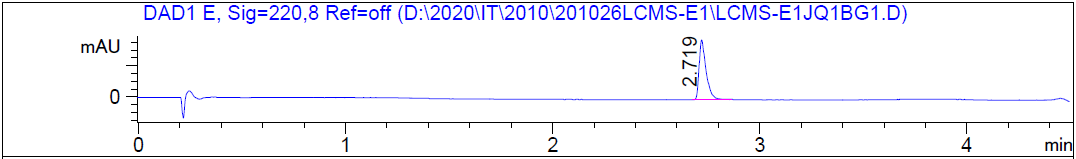

HRMS [C_33_H_44_BrClFN_7_O_8_S] Cal: 832.1901; Obs: 832.187

**The synthetic route for HLDA-113**

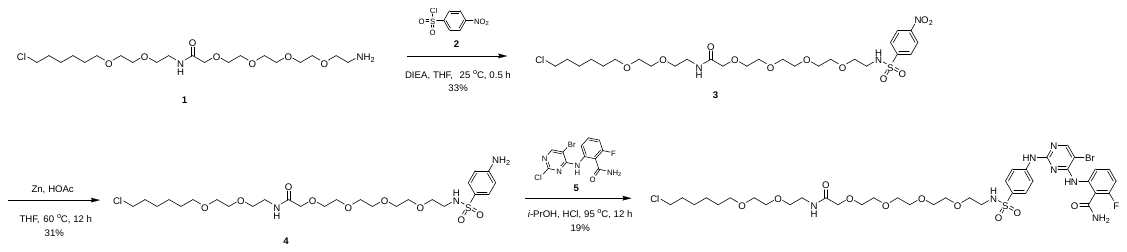

**Preparation of compound 3**

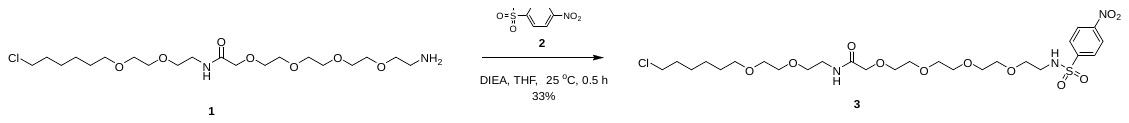

To a solution of 2-[2-[2-[2-(2-aminoethoxy)ethoxy]ethoxy]ethoxy]-*N*-[2-[2-(6-chlorohexoxy)ethoxy]ethyl]acetamide (400 mg, 700 μmol, 1.0 equiv, TFA salt), DIEA (271 mg, 2.10 mmol, 366 μl, 3.0 equiv) in THF (4 ml) was added 4-nitrobenzenesulfonyl chloride (232 mg, 1.05 mmol, 1.5 equiv), and it was stirred at 25 °C for 0.5 h. H_2_O (0.1ml) was added to quench this reaction and concentrated to afford crude product. The residue was purified by *prep*-TLC on silica gel (CH_2_Cl_2_ : MeOH = 10 :1) to give *N*-[2-[2-(6-chlorohexoxy)ethoxy]ethyl]-2-[2-[2-[2-[2-[(4-nitrophenyl)sulfonylamino]ethoxy]ethoxy]ethoxy]ethoxy]acetamide (150 mg, 233 μmol, 33 % yield) as a yellow oil.

**LC–MS:** MS (ES^+^): RT = 0.952 min, m/z = 642.5 [M + H^+^];

**Preparation of compound 4**

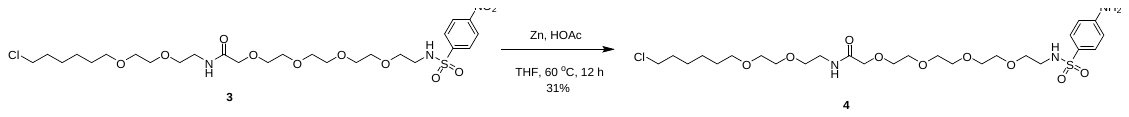

To a solution of *N*-[2-[2-(6-chlorohexoxy)ethoxy]ethyl]-2-[2-[2-[2-[2-[(4-nitrophenyl)sulfonylamino]ethoxy]ethoxy]ethoxy]ethoxy]acetamide (150 mg, 233 μmol, 1.0 equiv) in THF (2.5 ml) and HOAc (0.5 ml) was added Zn (152 mg, 2.34 mmol, 10.0 equiv), and then it was stirred at 60 °C for 12 h. After this time pH was adjust to 7–8 by sat. NaHCO_3_ water (50 ml) was added. After extracted with EtOAc (50 ml), the organic phase was washed with brine (3 x 50 ml), dried over anhydrous Na_2_SO_4_, filtered and concentrated under reduced pressure. The residue was purified by *prep*-TLC on silica gel (CH_2_Cl_2_ : MeOH = 10:1) to give 2-[2-[2-[2-[2-[(4-aminophenyl)sulfonylamino]ethoxy]ethoxy]ethoxy]ethoxy]-*N*-[2-[2-(6-chlorohexoxy)

ethoxy]ethyl]acetamide (45 mg, 73.51 μmol, 31% yield) as a yellow oil.

**LC–MS:** MS (ES^+^): RT = 0.631 min, m/z = 612.4 [M + H^+^];

**Spectra:**

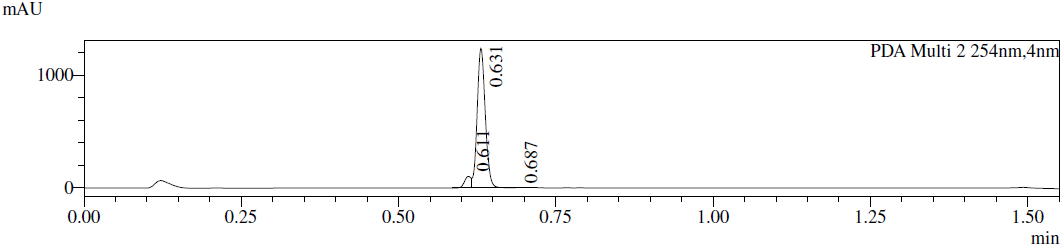

**Preparation of HLDA-113**

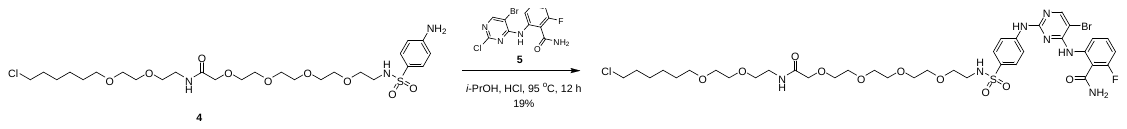

To a solution of 2-[2-[2-[2-[2-[(4-aminophenyl)sulfonylamino]ethoxy]ethoxy]ethoxy]ethoxy]-*N*-[2-[2-(6-chlorohexoxy)ethoxy]ethyl]acetamide (45 mg, 73 μmol, 1.0 equiv), 2-[(5-bromo-2-chloro-pyrimidin-4-yl)amino]-6-fluoro-benzamide (30.48 mg, 88.21 μmol, 1.2 equiv) in *i*-PrOH (2 ml) was added HCl (7.24 mg, 73.5 μmol, 7.10 μl, 37% purity, 1.0 equiv), and it was stirred at 95 °C for 12 h. After this time H_2_O (0.1 ml) was added to quench this reaction. This reaction was purified by *prep*-HPLC (column: 3_Phenomenex Luna C18 75*30 mm*3 μm; mobile phase: [water(0.05% HCl)–ACN];B%: 44%-64%,6.5min) and *prep*-HPLC (column: Waters Xbridge 150*25mm*5 μm; mobile phase: [water(10 mM NH_4_HCO_3_)–MeCN]; B%: 38%–68%, 8 min) to afford 2-[[5-bromo-2-[4-[2-[2-[2-[2-[2-[2-[2-(6-chlorohexoxy)ethoxy]ethylamino]-2-oxo-ethoxy]ethoxy]

ethoxy]ethoxy]ethylsulfamoyl]anilino]pyrimidin-4-yl]amino]-6-fluoro-benzamide (13.14 mg, 13.69 μmol, 19% yield,) as an off-white gum.

**^1^H NMR** (400 MHz, CD_3_OD) δ 8.37 (d, *J* = 8.2 Hz, 1H), 8.26 (s, 1H), 7.83 (d, *J* = 8.8 Hz, 2H), 7.71 (d, *J* = 8.8 Hz, 2H), 7.56– 7.46 (m, 1H), 6.99 (dd, *J* = 8.4, 10.6 Hz, 1H), 3.97 (s, 2H), 3.66–3.38 (m, 26H), 3.07– 3.02 (m, 2H), 1.77– 1.68 (m, 2H), 1.55 (quin, *J* = 6.9 Hz, 2H), 1.47– 1.28 (m, 4H)

**LC–MS:** MS (ES^+^): RT = 2.731 min, m/z = 922.2 [M + H^+^];

**Spectra:**

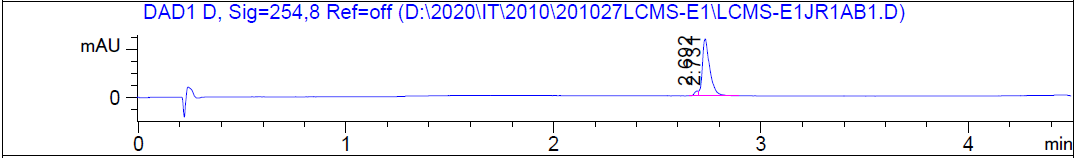

HRMS [C_37_H_52_BrClFN_7_O_10_S] Cal: 920.2425; Obs: 920.2391

**The synthetic route for HLDA-111**

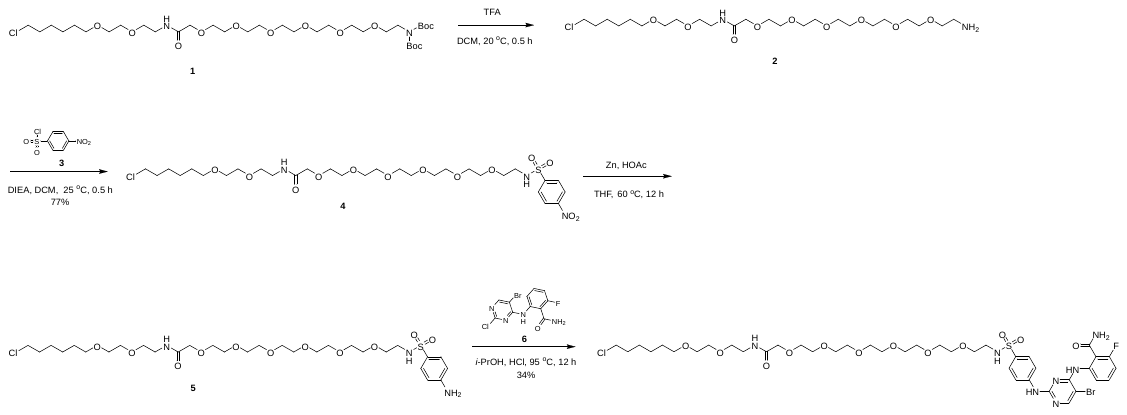

**Preparation of compound 2**

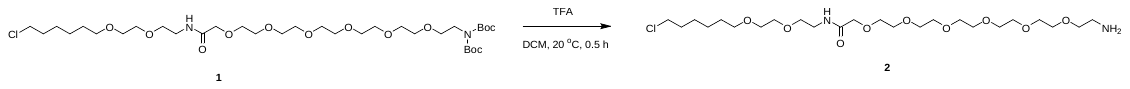

To a solution of tert-butyl *N*-tert-butoxycarbonyl-N-[2-[2-[2-[2-[2-[2-[2-[2-[2-(6-chlorohexoxy)ethoxy]ethylamino]-2-oxo-ethoxy]ethoxy]ethoxy]ethoxy]ethoxy]ethoxy]ethyl]carbamate (577 mg, 774 μmol, 1.0 equiv) in CH_2_Cl_2_ (2 ml) was added TFA (1 ml).The mixture was stirred at 20 °C for 0.5 h. Analysis by TLC demonstrated the formation of the desired product. The reaction mixture was concentrated under reduced pressure to give 2-[2-[2-[2-[2-[2-(2-aminoethoxy)ethoxy]ethoxy]ethoxy]ethoxy]ethoxy]-*N*-[2-[2-(6-chlorohexoxy)ethoxy]ethyl]acetamide (505 mg, 766 μmol, crude, TFA salt) as a yellow oil and it was used in the next step without further purification.

**Preparation of compound 4**

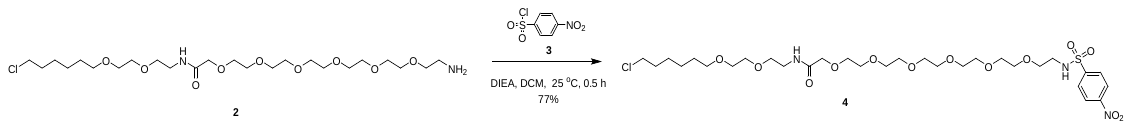

To a solution of 2-[2-[2-[2-[2-[2-(2-aminoethoxy)ethoxy]ethoxy]ethoxy]ethoxy]ethoxy]-*N*-[2-[2-(6-chlorohexoxy)ethoxy]ethyl]acetamide (505 mg, 766 μmol, 1.0 equiv, TFA salt) and 4-nitrobenzenesulfonyl chloride (339 mg, 1.53 mmol, 2.0 equiv) in CH_2_Cl­_2_ (10 ml) was added DIEA (495 mg, 3.83 mmol, 667 μl, 5.0 equiv). The mixture was stirred at 20 °C for 0.5 h after which time water was added (50 ml) and the mixture was extracted with EtOAc (50 ml). The combined organic phase was washed with brine (3 x 50 ml), dried over anhydrous Na_2_SO_4_, filtered, and concentrated under reduced pressure. The crude product was purified by column chromatography on silica gel (EtOAc : MeOH = 1:0 to 10:1 ) to give *N*-[2-[2-(6-chlorohexoxy)ethoxy]ethyl]-2-[2-[2-[2-[2-[2-[2-[(4-nitrophenyl)sulfonylamino]ethoxy]ethoxy]ethoxy]ethoxy]ethoxy]ethoxy]acetamide (430 mg, 588 μmol, 77% yield) as a yellow oil.

**LC–MS:** MS (ES^+^): RT = 0.882 min, m/z = 730.1 [M + H^+^];

**Spectra:**

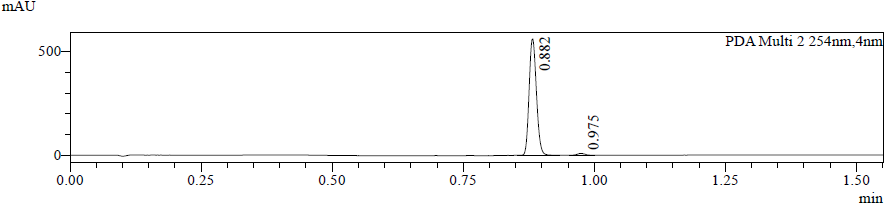

**Preparation of compound 5**

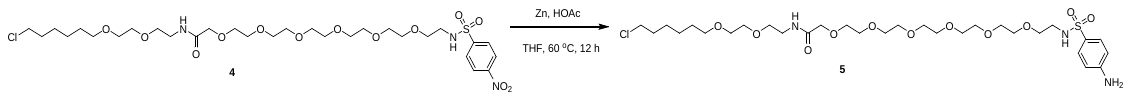

To a solution of *N*-[2-[2-(6-chlorohexoxy)ethoxy]ethyl]-2-[2-[2-[2-[2-[2-[2-[(4-nitrophenyl)sulfonylamino]ethoxy]ethoxy]ethoxy]ethoxy]ethoxy]ethoxy]acetamide (430 mg, 588 μmol, 1.0 equiv) in THF (5.5 ml) and HOAc (1.1 ml) was added Zn (1.93 g, 29.4 mmol, 50.0 equiv), and then it was stirred at 60 °C for 12 h. The pH of the mixture was adjust to 7–8 by sat. aq. NaHCO_3_ (3 ml.), To the reaction mixture was added water (50 ml) and the mixture was extracted with EtOAc (50 ml). The combined organic phase was washed with brine (3 x 50 ml), dried over anhydrous Na_2_SO_4_, filtered, and concentrated under reduced pressure to give 2-[2-[2-[2-[2-[2-[2-[(4-aminophenyl)sulfonylamino]ethoxy]ethoxy]ethoxy]ethoxy]ethoxy]ethoxy]-*N*-[2-[2-(6-chlorohexoxy)ethoxy]ethyl]acetamide (400 mg, crude) as a yellow oil and it was used by next step without further purification.

**LC–MS:** MS (ES^+^): RT = 0.881 min, m/z = 700.2 [M + H^+^];

**Spectra:**

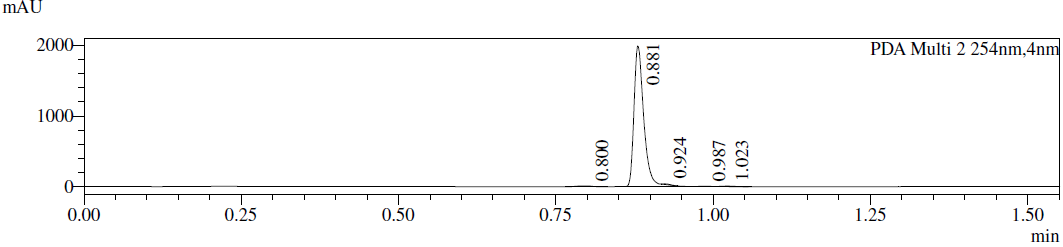

**Preparation of HLDA-111**

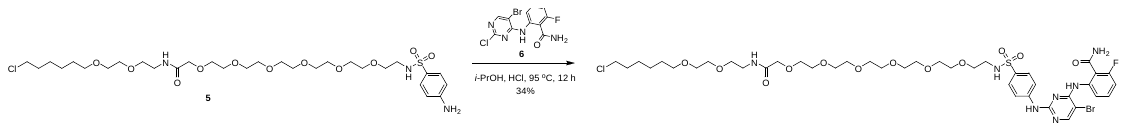

To a solution of 2-[2-[2-[2-[2-[2-[2-[(4-aminophenyl)sulfonylamino]ethoxy]ethoxy]ethoxy]ethoxy]ethoxy]ethoxy]-*N*-[2-[2-(6-chlorohexoxy)ethoxy]ethyl]acetamide (400 mg, 571 μmol, 1.0 equiv), 2-[(5-bromo-2-chloro-pyrimidin-4-yl)amino]-6-fluoro-benzamide (236 mg, 685 μmol, 1.2 equiv) in *i*-PrOH (4 ml) was added HCl (69.4 mg, 571 μmol, 68.0 ul, 30% purity, 1.0 equiv), and it was stirred at 95 °C for 12 h. The pH of the reaction mixture was adjusted to 7 by TEA (0.1 ml), then filtered. It was purified by *prep*-HPLC (column: Waters Xbridge C18 150*50 mm* 10 μm; mobile phase: [water(10 mM NH_4_HCO_3_)–MeCN]; B%: 38%–68%, min) to afford 2-[[5-bromo-2-[4-[2-[2-[2-[2-[2-[2-[2-[2-[2-(6-chlorohexoxy)ethoxy]ethylamino]-2-oxo-ethoxy]ethoxy]ethoxy]ethoxy]ethoxy]ethoxy]ethylsulfamoyl]anilino]pyrimidin-4-yl]amino]-6-fluoro-benzamide (198 mg, 194 μmol, 34% yield) as a yellow gum.

**^1^H NMR** (400 MHz, CD_3_OD-d_4_) δ 8.37 (d, *J* = 8.2 Hz, 1H), 8.29 (s, 1H), 7.85 (d, *J* = 8.8 Hz, 2H), 7.72 (d, *J* = 8.8 Hz, 2H), 7.52 (dt, *J* = 6.5, 8.2 Hz, 1H), 7.00 (dd, *J* = 8.6, 10.0 Hz, 1H), 3.97 (s, 2H), 3.69– 3.53 (m, 26H), 3.51– 3.40 (m, 8H), 3.05 (t, *J* = 5.4 Hz, 2H), 1.74 (quin, *J* = 7.0 Hz, 2H), 1.62–1.52 (m, 2H), 1.49–1.33 (m, 4H)

**LC-MS:** MS (ES^+^): RT = 3.193 min, m/z = 1010.3 [M + H^+^];

**Spectra:**

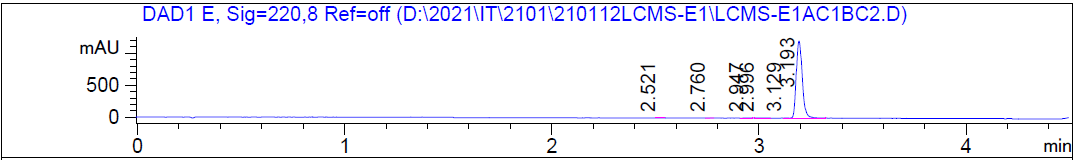

HRMS [C_41_H_60_BrClFN_7_O_12_S] Cal: 1008.2949; Obs: 1008.2916

***SERIES 3***

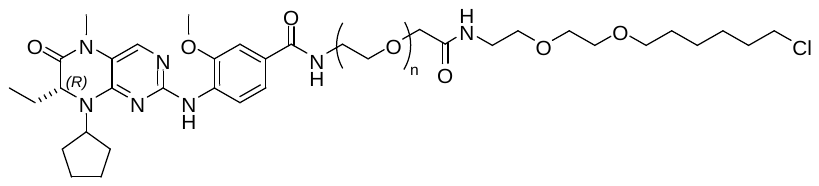

HLDA-131, n=2

HLDA-132, n=4

HLDA-133, n=6

**The synthetic route for HLDA-131**

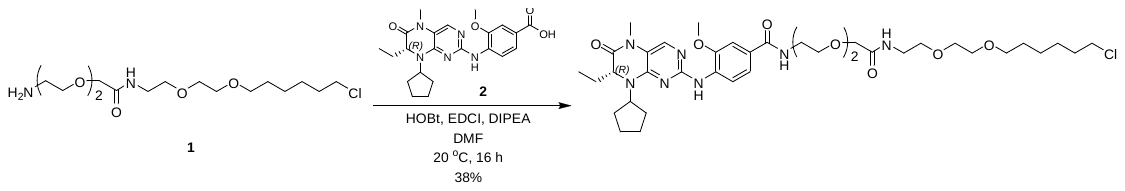

**Preparation of compound 1**

Known compound from *Angew. Chem. Int. Ed.* **2006**, *45*, 4936–4940.

**Preparation of compound 2**

Known compound from *A.C.S. Med. Chem. Lett.* **2019**, *10*, 1443–-1449

**Preparation of HLDA-131**

A mixture of 4-[(8-cyclopentyl-7-ethyl-5-methyl-6-oxo-7*H*-pteridin-2-yl)amino]-3-methoxy-benzoic acid (30.0 mg, 70.5 μmol, 1.0 equiv), 2-[2-(2-aminoethoxy)ethoxy]-*N*-[2-[2-(6-chlorohexoxy)ethoxy]ethyl]acetamide (34.3 mg, 84.6 μmol, 1.2 equiv, HCl salt), HOBt (14.3 mg, 106 μmol, 1.5 equiv), EDCI (20.3 mg, 106 μmol, 1.5 equiv) and DIPEA (45.6 mg, 353 μmol, 5.0 equiv) in DMF (1 ml) was stirred at 20 °C for 16 h. The reaction mixture was diluted with water (20 ml) and the mixture was extracted with EtOAc (2 x 20 ml). The combined organic layer was washed with brine (30 ml), dried with anhydrous Na_2_SO_4_, filtered, and concentrated under reduced pressure. The residue was purified by *prep*-HPLC (column: Phenomenex Luna C18 150*25 mm*10 μm; mobile phase: [water(0.225%FA)–ACN]; B%: 25%–55%,10 min) and *prep*-HPLC (column: Waters Xbridge 150*25 mm* 5 –m;mobile phase: [water(10 mM NH_4_HCO_3_)–MeCN] ;B%: 45%–75%, 9 min) to afford *N*-[2-[2-[2-[2-[2-(6-chlorohexoxy)ethoxy]ethylamino]-2-oxo-ethoxy]ethoxy]ethyl]-4-[(8-cyclopentyl-7-ethyl-5-methyl-6-oxo-7*H*-pteridin-2-yl)amino]-3-methoxy-benzamide (21.3 mg, 26.9 μmol, 38% yield) as a yellow gum.

**^1^H NMR** (400 MHz, DMSO-d_6_): δ 8.45–8.35 (m, 2H), 7.84 (s, 1H), 7.65–7.57 (m, 2H), 7.52–7.46 (m, 2H), 4.41–4.30 (m, 1H), 4.27–4.21 (m, 1H), 3.93 (s, 3H), 3.87 (s, 2H), 3.62-3.53 (m, 8H), 3.49–3.38 (m, 9H), 3.30–3.22 (m, 6H), 2.06–1.98 (m, 1H), 1.93–1.73 (m, 6H), 1.72–1.58 (m, 5H), 1.50–1.41 (m, 2H), 1.40–1.22 (m, 4H), 0.76 (t, 3H, *J* = 7.6 Hz).

**LC–MS:** MS (ES^+^): RT = 2.488 min, m/z = 776.3 [M + H^+^]

**Spectra:**

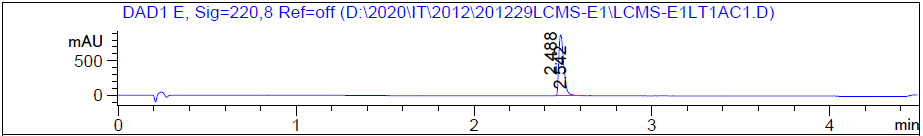

HRMS [C_38_H_58_ClN_7_O_8_] Cal: 776.4108; Obs: 776.4073

**The synthetic route for HLDA-132**

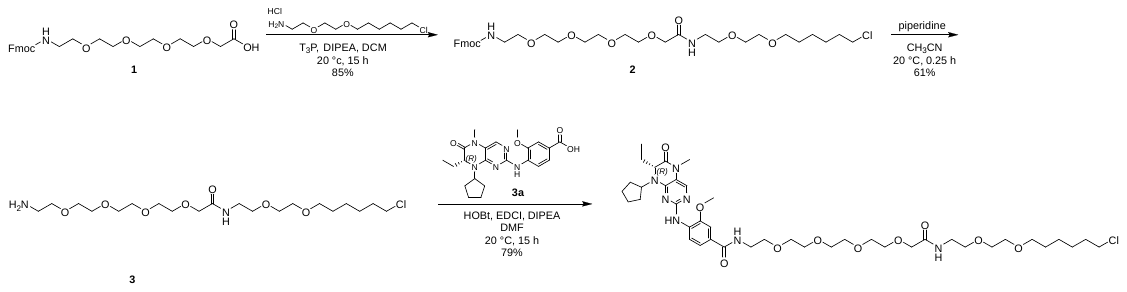

**Preparation of compound 2**

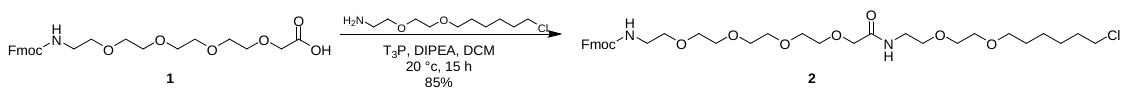

To a solution of 2-[2-[2-[2-[2-(9*H*-fluoren-9-ylmethoxycarbonylamino)ethoxy]ethoxy]ethoxy]ethoxy]acetic acid (100 mg, 0.21 mmol, 1.0 equiv) and 2-[2-(6-chlorohexoxy)ethoxy]ethanamine (55.0 mg, 0.21 mmol, 1.0 equiv, HCl salt) in CH_2_Cl_2_ (2 ml) were added T_3_P (202 mg, 0.32 mmol, 50% purity, 1.5 equiv) and DIPEA (109 mg, 0.84 mmol, 4.0 equiv). The mixture was stirred at 20 °C for 15 h. The reaction mixture was quenched with water (5 ml) and extracted with ethyl acetate (3 x 10 ml). The combined organic phase was washed with brine (15 ml), dried over anhydrous Na_2_SO_4_, filtered, and concentrated under reduced pressure. The residue was purified by *prep*-TLC on silica gel (EtOAc /petroleum ether = 1/1) to afford 9*H*-fluoren-9-ylmethyl *N*-[2-[2-[2-[2-[2-[2-[2-(6-chlorohexoxy)ethoxy]ethylamino]-2-oxoethoxy]ethoxy]ethoxy]ethoxy]ethyl]carbamate (122 mg, 0.18 mmol, 85% yield) as a yellow oil.

**LC–MS:** MS (ES^+^): RT = 0.764 min, m/z = 679.1 [M + H^+^];

**Spectra:**

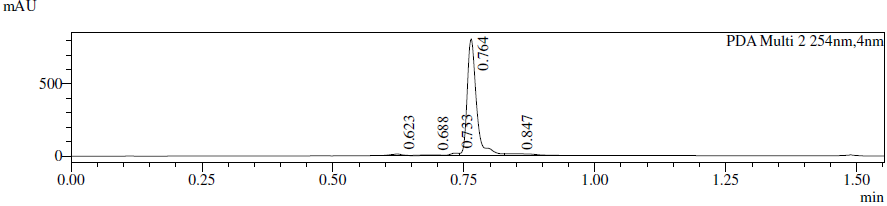

**Preparation of compound 3**

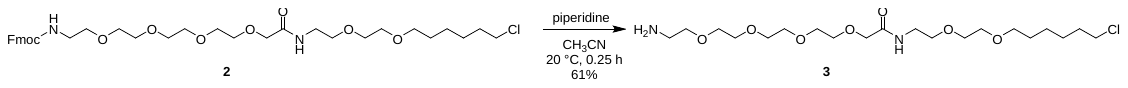

To a solution of 9*H*-fluoren-9-ylmethyl *N*-[2-[2-[2-[2-[2-[2-[2-(6-chlorohexoxy)ethoxy]ethylamino]-2-oxoethoxy]ethoxy]ethoxy]ethoxy]ethyl]carbamate (102 mg, 0.15 mmol, 1.0 equiv) in CH_3_CN (4.6 ml) was added piperidine (440 mg, 5.16 mmol, 34.4 equiv). The mixture was stirred at 20 °C for 0.25 h. The reaction mixture was concentrated under reduced pressure. The residue was purified by *prep*-HPLC (neutral conditions: column: Waters Xbridge 150*25 mm* 5 μm;mobile phase:[water(10 mM NH_4_HCO_3_)–ACN]; B%: 20%–50%,10 min) to afford 2-[2-[2-[2-(2-aminoethoxy)ethoxy]ethoxy]ethoxy]-*N*-[2-[2-(6-chlorohexoxy)ethoxy]ethyl]acetamide (42.0 mg, 0.09 mmol, 61% yield) as a yellow oil.

**LC–MS:** MS (ES^+^): RT = 0.783 min, m/z = 457.2 [M + H^+^];

**Spectra:**

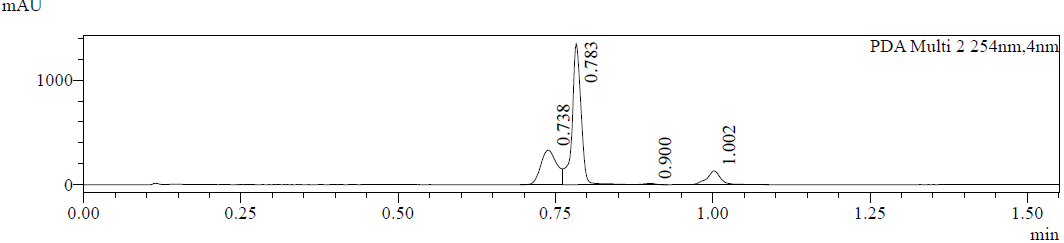

**Preparation of compound 3a**

Known compound from *A.C.S. Med. Chem. Lett.* **2019**, *10*, 1443–-1449

**Preparation of HLDA-132**

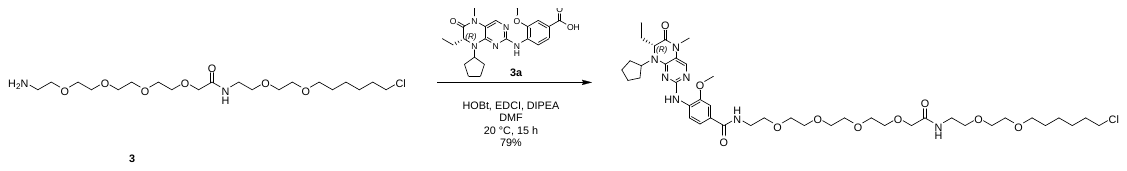

A mixture of 2-[2-[2-[2-(2-aminoethoxy)ethoxy]ethoxy]ethoxy]-*N*-[2-[2-(6-chlorohexoxy)ethoxy]ethyl]acetamide (35.0 mg, 0.08 mmol, 1.0 equiv), 4-[[(7*R*)-8-cyclopentyl-7-ethyl-5-methyl-6-oxo-7*H*-pteridin-2-yl]amino]-3-methoxy-benzoic acid (32.6 mg, 0.08 mmol, 1.0 equiv), EDCI (36.7 mg, 0.19 mmol, 2.5 equiv), HOBt (25.9 mg, 0.19 mmol, 2.5 equiv) and DIPEA (49.5 mg, 0.38 mmol, 5.0 equiv) in DMF (1 ml) was stirred at 20 °C for 15 h under N_2_ atmosphere. The reaction mixture was diluted with water (20 ml) and the mixture was extracted with EtOAc (3 x 15 ml). The combined organic phase was washed with brine (10 ml), dried with anhydrous Na_2_SO_4_, filtered, and the filtrate was concentrated under reduced pressure. The residue was purified by *prep*-HPLC (neutral condition: column: Waters Xbridge 150*25 mm* 5 μm; mobile phase:[water(10 mM NH_4_HCO_3_)–MeCN]; B%: 45%–75%, 9 min) to afford *N*-[2-[2-[2-[2-[2-[2-[2-(6-chlorohexoxy)ethoxy]ethylamino]-2-oxo-ethoxy]ethoxy]ethoxy]ethoxy]ethyl]-4-[[(7*R*)-8-cyclopentyl-7-ethyl-5-methyl-6-oxo-7*H*-pteridin-2-yl]amino]-3-methoxy-benzamide (53.5 mg, 0.06 mmol, 79% yield) as a yellow gum.

**^1^H NMR** (400 MHz, DMSO-d_6_) δ 8.45–8.36 (m, 2H), 7.84 (s, 1H), 7.64–7.57 (m, 2H), 7.52–7.45 (m, 2H), 4.35 (t, 1H, *J* = 8.0 Hz), 4.24 (dd, 1H, *J* = 3.6, 7.6 Hz), 3.93 (s, 3H), 3.86 (s, 2H), 3.60 (t, 2H, *J* = 6.8 Hz), 3.57–3.53 (m, 10H), 3.54–3.48 (m, 4H), 3.49–3.47 (m, 2H), 3.46–3.40 (m, 6H), 3.37–3.33 (m, 2H), 3.27–3.23 (m, 5H), 2.03–1.98 (m, 1H), 1.93–1.85 (m, 2H), 1.83–1.73 (m, 4H), 1.72–1.58 (m, 5H), 1.50–1.43 (m, 2H), 1.41–1.23 (m, 4H), 0.76 (t, 3H, *J* = 7.6 Hz).

**LC–MS:** MS (ES^+^): RT = 2.517 min, m/z = 864.4 [M + H^+^];

**Spectra:**

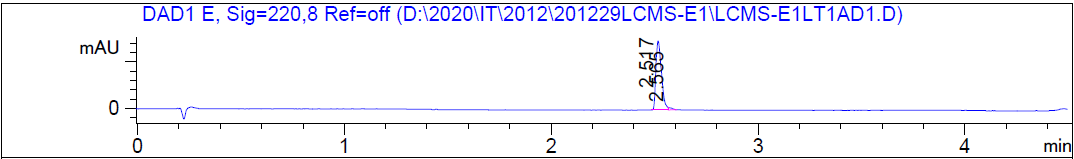

HRMS [C_42_H_66_ClN_7_O_10_] Cal: 864.4632; Obs: 864.4593

**The synthetic route for HLDA-133**

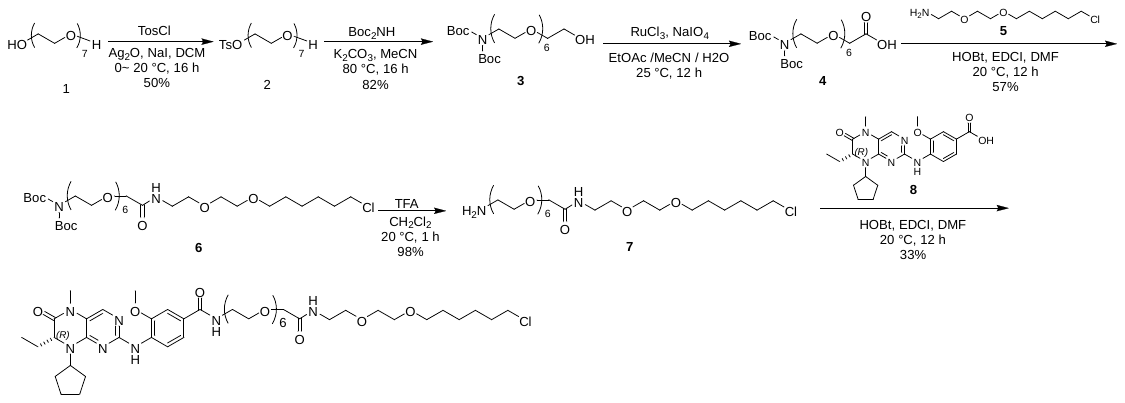

**Preparation of compound 2**

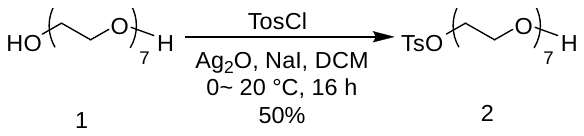

To a stirred solution of 2-[2-[2-[2-[2-[2-(2-hydroxyethoxy)ethoxy]ethoxy]ethoxy]ethoxy]ethoxy]ethanol (10.0 g, 30.6 mmol, 1.0 equiv) in CH_2_Cl_2_ (200 ml) were added Ag_2_O (10.7 g, 45.9 mmol, 1.5 equiv), NaI (5.05 g, 33.7 mmol, 1.1 equiv) and TosCl (6.43 g, 33.7 mmol, 1.1 equiv) at 0 °C. The reaction mixture was warmed to 20 °C and stirred for 16 h. The reaction mixture was filtered and the filtrate was concentrated under reduced pressure. The residue was purified by column chromatography on silica gel (CH_2_Cl_2_/MeOH = 200/1 to 10/1) to afford 2-[2-[2-[2-[2-[2-(2-hydroxyethoxy)ethoxy]ethoxy]ethoxy]ethoxy]ethoxy]ethyl 4-methylbenzenesulfonate (7.35 g, 15.3 mmol, 50% yield) as a colorless oil.

**^1^H NMR** (400 MHz, CDCl_3_) δ 7.82 (d, 2H, *J* = 8.0 Hz), 7.37 (d, 2H, *J* = 8.0 Hz), 4.18 (t, 2H, *J* = 4.8 Hz), 3.78–3.61 (m, 27H), 2.47 (s, 3H).

**Preparation of compound 3**

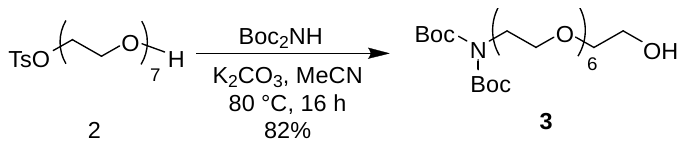

A mixture of 2-[2-[2-[2-[2-[2-(2-hydroxyethoxy)ethoxy]ethoxy]ethoxy]ethoxy]ethoxy]ethyl 4-methylbenzenesulfonate (7.35 g, 15.3 mmol, 1.0 equiv), *tert*-butyl-*N*-*tert*-butoxycarbonylcarbamate (6.65 g, 30.6 mmol, 2.0 equiv) and K_2_CO_3_ (6.34 g, 45.9 mmol, 3.0 equiv) in MeCN (120 ml) was heated to 80 °C and stirred for 16 h. The reaction mixture was filtered and the filtrate was concentrated under reduced pressure. The residue was purified by column chromatography on silica gel (CH_2_Cl_2_/MeOH = 100/1 to 50/1) to afford *tert*-butyl-*N*-*tert*-butoxycarbonyl-*N*-[2-[2-[2-[2-[2-[2-(2-hydroxyethoxy)ethoxy]ethoxy]ethoxy]ethoxy]ethoxy]ethyl]carbamate (6.60 g, 12.6 mmol, 82% yield) as a colorless oil.

**^1^H NMR** (400 MHz, CDCl_3_) δ 3.83–3.78 (m, 2H), 3.76–3.73 (m, 2H), 3.71–3.61 (m, 24H), 1.52 (s, 18H).

**Preparation of compound 4**

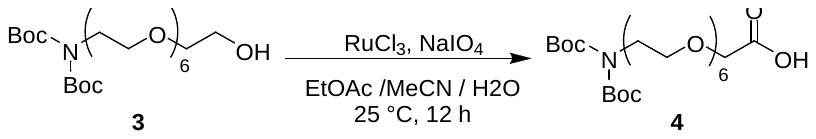

To a solution of *tert*-butyl *N-tert*-butoxycarbonyl-*N*-[2-[2-[2-[2-[2-[2-(2-hydroxyethoxy)ethoxy]ethoxy]ethoxy]ethoxy]ethoxy]ethyl]carbamate (1.00 g, 1.90 mmol, 1.0 equiv) in EtOAc (5 ml) and MeCN (5 ml) and H_2_O (5 ml) were added RuCl_3_ (39.5 mg, 190 μmol, 0.1 equiv) and NaIO_4_ (1.63 g, 7.61 mmol, 4.0 equiv) under N_2_ atmosphere. The mixture was stirred at 25 °C for 12 h. To the reaction mixture was added water (20 ml). The mixture was filtered and the filtrate was extracted with ethyl acetate (3 x 15 ml). The combined organic phase was washed with brine (18 ml), dried over anhydrous Na_2_SO_4_, filtered, and concentrated under reduced pressure. The residue was purified by column chromatography on silica gel (CH_2_Cl_2_/MeOH = 100/1 to 30/1) to afford 2-[2-[2-[2-[2-[2-[2-[bis(*tert*-butoxycarbonyl)amino]ethoxy]ethoxy]ethoxy]ethoxy]ethoxy]ethoxy]acetic acid (625 mg, crude) as a yellow oil.

**^1^H NMR** (400 MHz, CDCl3) δ 4.19 (s, 2H), 3.83–3.77 (m, 4H), 3.76–3.59 (m, 20H), 1.52 (s, 18H).

**Preparation of compound 6**

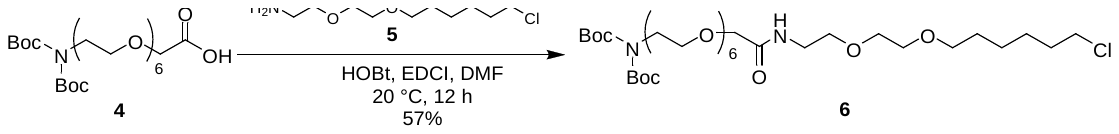

A mixture of 2-[2-[2-[2-[2-[2-[2-[bis(tert-butoxycarbonyl)amino]ethoxy]ethoxy]ethoxy]ethoxy]ethoxy]ethoxy]acetic acid (1.00 g, 1.85 mmol, 1.0 equiv), 2-[2-(6-chlorohexoxy)ethoxy]ethanamine (482 mg, 1.85 mmol, 1.0 equiv, HCl), HOBt (376 mg, 2.78 mmol, 1.5 equiv), EDCI (710 mg, 3.71 mmol, 2 equiv) and DIEA (1.20 g, 9.27 mmol, 5.0 equiv) in DMF (20 ml) was stirred at 20 °C for 12 h under N_2_. The reaction mixture was diluted with brine (35 ml) and the mixture was extracted with EtOAc (3 x 25 ml). The combined organic phase was washed with brine (30 ml), dried with anhydrous Na_2_SO_4_, filtered, and concentrated under reduced pressure. The residue was purified by column chromatography on silica gel (CH_2_Cl_2_/MeOH= 100/1 to 30/1) and then *prep*-HPLC (column: Phenomenex luna C18 150*40 mm* 15 μm; mobile phase: [water(0.225%FA)–MeCN]; B%: 50%–80%, 10 min) to afford *tert*-butyl *N-tert*-butoxycarbonyl-*N*-[2-[2-[2-[2-[2-[2-[2-[2-[2-(6-chlorohexoxy)ethoxy]ethylamino]-2-oxo-ethoxy]ethoxy]ethoxy]ethoxy]ethoxy]ethoxy]ethyl]carbamate (790 mg, 1.06 mmol, 57% yield) as a colorless oil.

**^1^H NMR** (400 MHz, CDCl_3_) δ 7.20–7.12 (m, 1H), 4.02 (s, 2H), 3.83–3.78 (m, 2H), 3.75–3.45 (m, 36H), 1.87–1.74 (m, 2H), 1.65–1.60 (m, 2H), 1.52 (s, 18H), 1.44–1.36 (m, 2H).

**Preparation of compound 7**

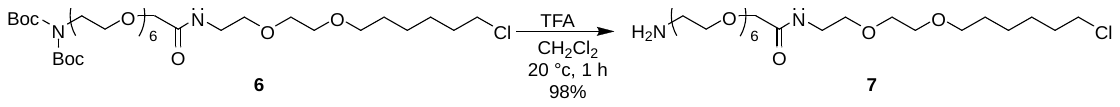

A mixture of *tert*-butyl *N-tert*-butoxycarbonyl-N-[2-[2-[2-[2-[2-[2-[2-[2-[2-(6-chlorohexoxy)ethoxy]ethylamino]-2-oxo-ethoxy]ethoxy]ethoxy]ethoxy]ethoxy]ethoxy]ethyl]carbamate (90.0 mg, 121 μmol, 1.0 equiv) in CH_2_Cl_2_ (1 ml) and TFA (0.5 ml) was stirred at 20 °C for 1 h. The reaction mixture was concentrated under reduced pressure to afford 2-[2-[2-[2-[2-[2-(2-aminoethoxy)ethoxy]ethoxy]ethoxy]ethoxy]ethoxy]-*N*-[2-[2-(6-chlorohexoxy)ethoxy]ethyl]acetamide (78.0 mg, 118 μmol, 98% yield, TFA salt) as a yellow oil.

**Preparation of compound 8**

Known compound from *ACS Med. Chem. Lett.* **2019**, *10*, 1443–1449

**Preparation of HLDA-133**

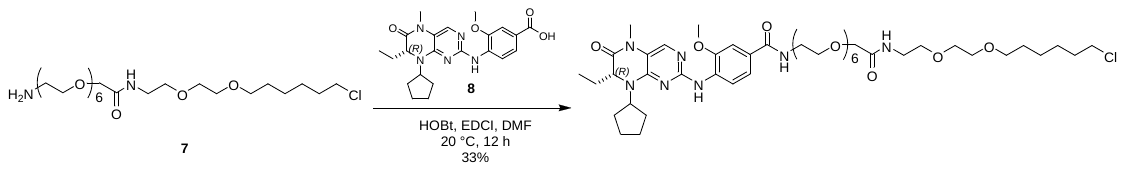

A mixture of 2-[2-[2-[2-[2-[2-(2-aminoethoxy)ethoxy]ethoxy]ethoxy]ethoxy]ethoxy]-*N*-[2-[2-(6-chlorohexoxy)ethoxy]ethyl]acetamide (74.4 mg, 113 μmol, 1.2 equiv, TFA), 4-[[(7*R*)-8-cyclopentyl-7-ethyl-5-methyl-6-oxo-7*H*-pteridin-2-yl]amino]-3-methoxy-benzoic acid (40.0 mg, 94.0 μmol, 1.0 equiv), HOBt (25.4 mg, 188 μmol, 2.0 equiv), EDCI (54.1 mg, 282 μmol, 3.0 equiv) and DIEA (72.9 mg, 564 μmol, 6.0 equiv) in DMF (2.5 ml) was stirred at 20 °C for 12 h under N_2_ atmosphere. The reaction mixture was diluted with water (20 ml) and the mixture was extracted with EtOAc (3 x 15 ml). The combined organic phase was washed with brine (10 ml), dried with anhydrous Na_2_SO_4_, filtered and concentrated under reduced pressure. The residue was purified by column chromatography on silica gel (CH_2_Cl_2_/MeOH = 10/1), *prep*-HPLC (column: Phenomenex Gemini-NX C18 75*30 mm*3 μm;mobile phase: [water(0.225%FA)–MeCN]; B%: 28%–58%, 7 min) and then *prep*-HPLC (column: Waters Xbridge 150*25 mm* 5 μm;mobile phase: [water(10 mM NH_4_HCO_3_)–MeCN]; B%: 46%–76%, 9 min) to afford *N*-[2-[2-[2-[2-[2-[2-[2-[2-[2-(6-chlorohexoxy)ethoxy]ethylamino]-2-oxo-ethoxy]ethoxy]ethoxy]ethoxy]ethoxy]ethoxy]ethyl]-4-[[(7*R*)-8-cyclopentyl-7-ethyl-5-methyl-6-oxo-7*H*-pteridin-2-yl]amino]-3-methoxy-benzamide (30.0 mg, 31.2 μmol, 33% yield) as a yellow oil.

**^1^H NMR** (400 MHz, DMSO-d_6_) δ 8.47–8.38 (m, 2H), 7.85 (s, 1H), 7.67–7.58 (m, 2H), 7.54–7.45 (m, 2H), 4.40–4.31 (m, 1H), 4.28–4.22 (m, 1H), 3.94 (s, 3H), 3.87 (s, 2H), 3.63–3.60 (m, 3H), 3.55–3.51 (m, 13H), 3.46–3.44 (m, 2H), 3.44–3.41 (m, 4H), 3.37–3.32 (m, 11H), 3.29–3.23 (m, 5H), 2.07–1.87 (m, 4H), 1.84–1.75 (m, 4H), 1.72–1.60 (m, 5H), 1.52–1.45 (m, 2H), 1.40–1.34 (m, 2H), 1.33–1.26 (m, 2H), 0.77 (t, 3H, *J* = 7.2 Hz).

**LCMS:** MS (ES^+^): RT = 3.459 min, m/z = 476.8 [M/2 + H^+^];

**Spectra:**

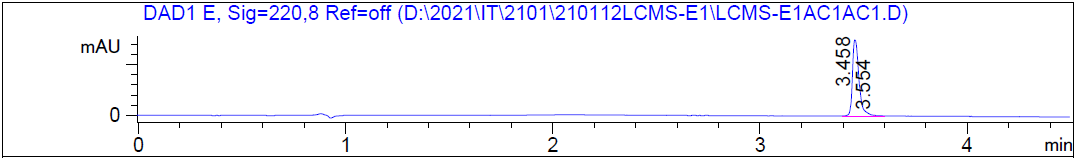

HRMS [C_46_H_74_ClN_7_O_12_] Cal: 952.5157; Obs: 952.512

***SERIES 4***

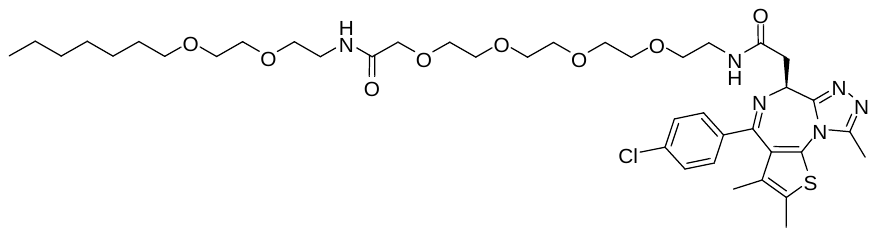

HLDA-120

**The synthetic route for HLDA-120**

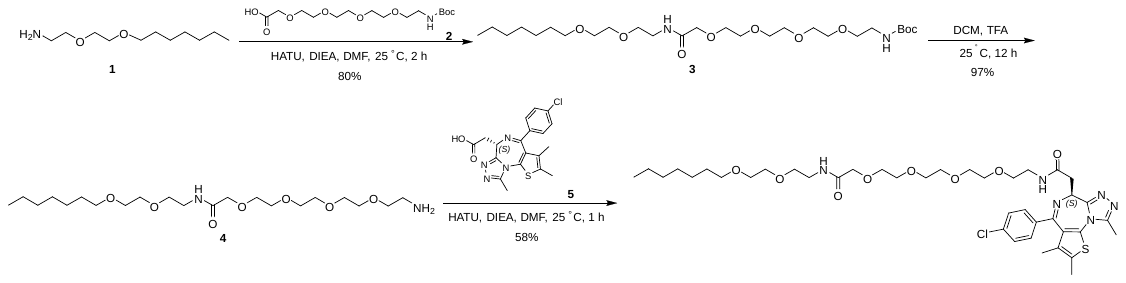

**Preparation of compound 1**

See HLDA-110

**Preparation of compound 2**

Known compound from *Inorganica Chim. Acta*, **2011**, *365*38– 48

**Preparation of compound 3**

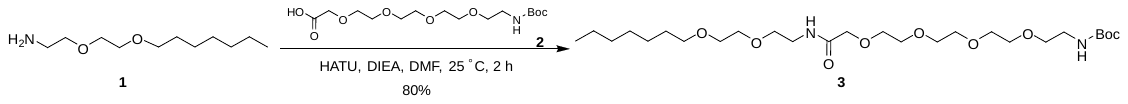

To a solution of 2-(2-heptoxyethoxy)ethanamine (310 mg, 977 μmol, 1.0 equiv, TFA salt) in DMF (3 ml) was added HATU (446 mg, 1.17 mmol, 1.2 equiv), 2-[2-[2-[2-[2-(*tert*-butoxycarbonylamino)ethoxy]ethoxy]ethoxy]ethoxy]acetic acid (343 mg, 977 μmol, 1.0 equiv) and DIEA (379 mg, 2.93 mmol, 3.0 equiv). The mixture was stirred at 25 °C for 2 h. The mixture was concentrated. The residue was purified by *prep*-HPLC (column: Waters Xbridge 150*25 mm * 5 μm; mobile phase: [water (10 mM NH_4_HCO_3_)–MeCN]; B%: 38%68%, 10 min) to give *tert*-butyl *N*-[2-[2-[2-[2-[2-[2-(2-heptoxyethoxy)ethylamino]-2-oxo-ethoxy]ethoxy]ethoxy]ethoxy]ethyl]carbamate (420 mg, 80% yield).

**LC-MS:** MS (ES^+^): RT = 1.025 min, m/z = 537.3 [M + H^+^];

**Spectra:**

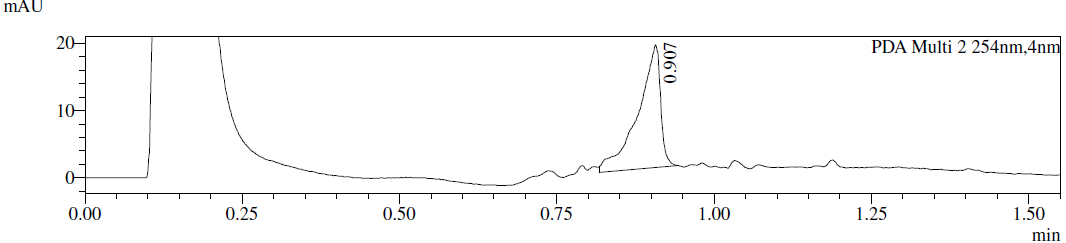

**Preparation of compound 4**

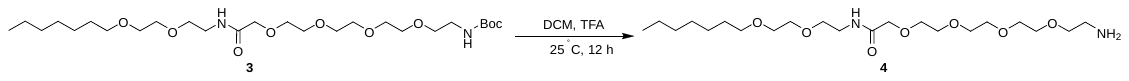

To a solution of *tert*-butyl *N*-[2-[2-[2-[2-[2-[2-(2-heptoxyethoxy)ethylamino]-2-oxoethoxy]ethoxy]ethoxy]ethoxy]ethyl]carbamate (100 mg, 186 μmol, 1.0 equiv) in DCM (1.5 ml) was added TFA (770 mg, 6.75 mmol, 0.5 ml, 36.2 equiv). The mixture was stirred at 25 °C for 12 h. The mixture was concentrated to give 2-[2-[2-[2-(2-aminoethoxy)ethoxy]ethoxy]ethoxy]-*N*-[2-(2-heptoxyethoxy)ethyl]acetamide (100 mg, 97% yield) as a colorless oil.

**LC–MS:** MS (ES^+^): RT = 0.518 min, m/z = 437.1 [M + H^+^];

**Preparation of HLDA-120**

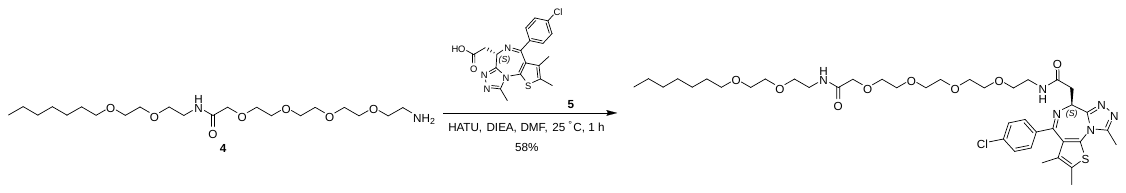

To a solution of 2-[(9*S*)-7-(4-chlorophenyl)-4,5,13-trimethyl-3-thia-1,8,11,12-tetrazatricyclo[8.3.0.02,6]trideca-2(6),4,7,10,12-pentaen-9-yl]acetic acid (73 mg, 181 μmol, 1.0 equiv) in DMF (1.5 ml) was added HATU (83 mg, 218 μmol, 1.2 equiv), 2-[2-[2-[2-(2-aminoethoxy)ethoxy]ethoxy]ethoxy]-*N*-[2-(2-heptoxyethoxy)ethyl]acetamide (100 mg, 182 μmol, 1.0 equiv, TFA salt) and DIEA (117 mg, 908 μmol, 5.0 equiv). The mixture was stirred at 25 °C for 1 h. The mixture was concentrated to give a residue. The residue was purified by *prep*-HPLC (column: Unisil 3-100 C_18_ Ultra 150 * 50 mm * 3 μm; mobile phase: [water (0.225%FA)–MeCN]; B%: 52%–82%, 10 min) to give 2-[(9*S*)-7-(4-chlorophenyl)-4,5,13-trimethyl-3-thia-1,8,11,12-tetrazatricyclo[8.3.0.02,6]trideca-2(6),4,7,10,12-pentaen-9-yl]-*N*-[2-[2-[2-[2-[2-[2-(2-heptoxyethoxy)ethylamino]-2-oxo-ethoxy]ethoxy]ethoxy]ethoxy]ethyl]acetamide (87 mg, 58% yield) as a yellow gum.

**^1^H NMR** (400 MHz, MeOD-d_4_): δ 7.54–7.36 (m, 4H), 4.63 (m, *J* = 5.2, 9.2 Hz, 1H), 3.97 (s, 2H), 3.67 (s, 12H), 3.63–3.58 (m, 4H), 3.57–3.53 (m, 4H), 3.49-3.40 (m, 7H), 3.28 (d, *J* = 5.2 Hz, 1H), 2.70 (s, 3H), 2.45 (s, 3H), 1.71 (s, 3H), 1.60–1.50 (m, 2H), 1.30 (d, *J* = 5.6 Hz, 8H), 1.00–0.81 (m, 3H)

**LC-MS:** MS (ES^+^): RT = 2.868 min, m/z = 820.8 [M + H^+^];

**Spectra:**

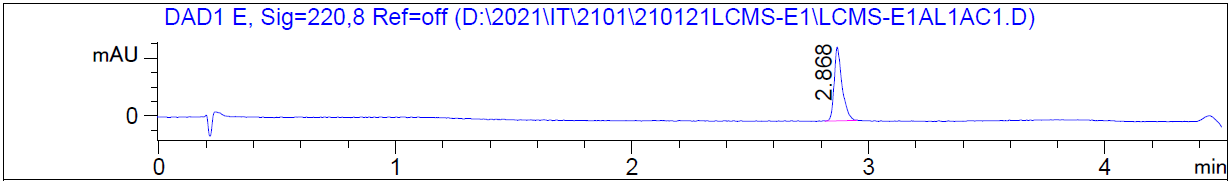

HRMS [C_40_H_59_ClN_6_O_8_S] Cal: 819.3876; Obs: 819.3846

***SERIES 5***

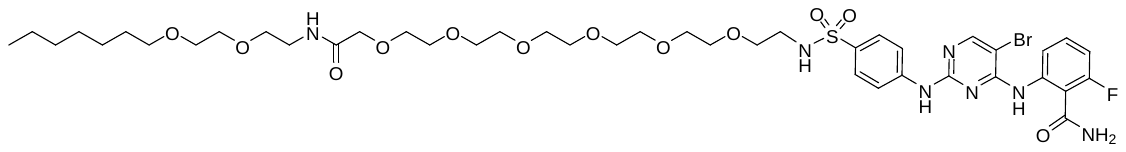

HLDA-110

**The synthetic route for HLDA-110**

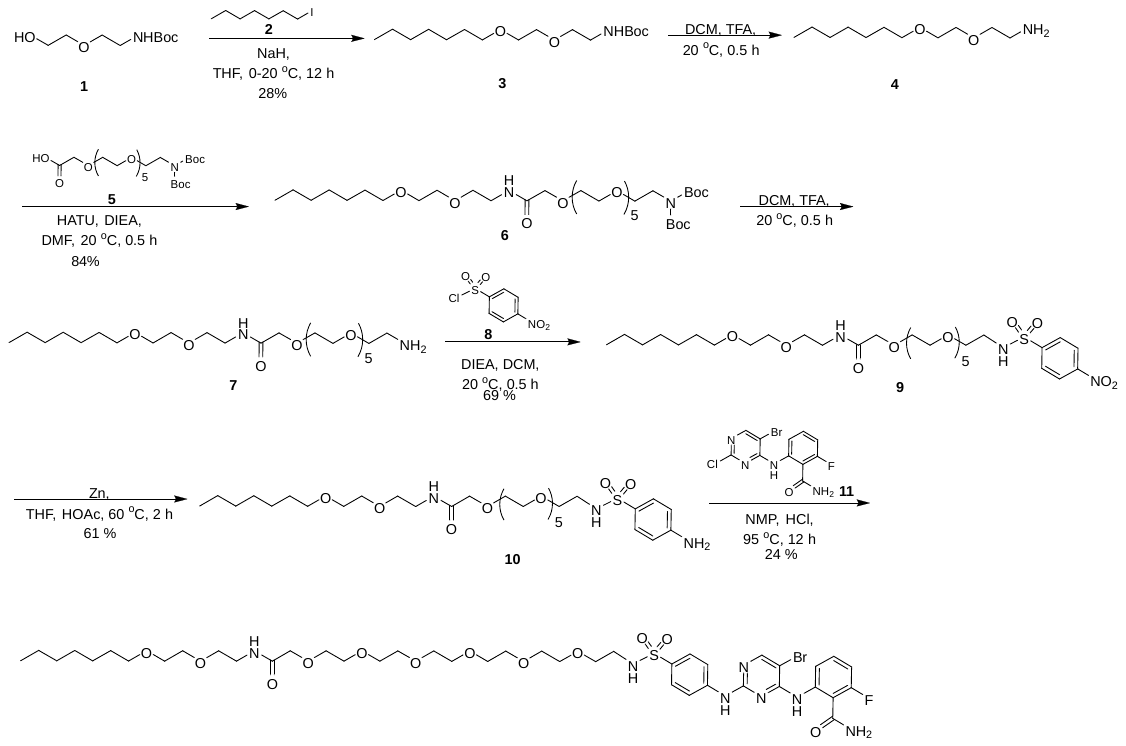

**Preparation of compound 3**

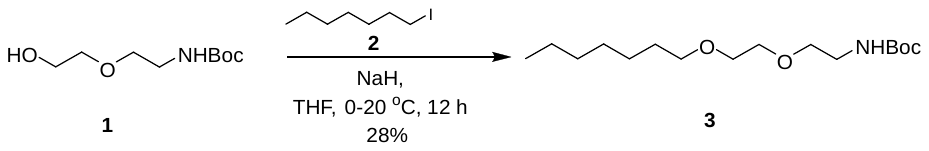

To a solution of *tert*-butyl (2-(2-hydroxyethoxy)ethyl)carbamate (5.0 g, 24.36 mmol, 1.0 equiv) in THF (50 ml) was added NaH (2.44 g, 60.90 mmol, 60% purity, 2.5 equiv) at 0 °C. The mixture was stirred at 0 °C for 30 min. Then 1-iodoheptane (8.26 g, 36.54 mmol, 1.5 equiv) in THF (10 ml) was added to the reaction mixture. The mixture was stirred at 20 °C for 11.5 h. The reaction mixture was quenched by addition of 40 ml of sat. aq. NH_4_Cl at 0°C, and then diluted with H_2_O (50 ml), and extracted with EtOAc (2 x 100 ml). The combined organic layers were washed with brine (100 ml), dried over Na_2_SO_4_, filtered, and concentrated under reduced pressure to give a residue. The residue was purified by column chromatography (SiO_2_, Petroleum ether/Ethyl acetate = 10/1 to 5/1). *tert*-butyl (2-(2-(heptyloxy)ethoxy)ethyl)carbamate (2.1 g, 6.92 mmol, 28% yield) was obtained as a yellow oil.

**^1^HNMR** (400 MHz, CDCl_3_): δ 5.14– 4.91 (m, 1H), 3.65– 3.51 (m, 6H), 3.46 (t, *J* = 6.8 Hz, 2H), 3.33 (m, 2H), 1.64– 1.58 (m, 2H), 1.48– 1.41 (s, 9H), 1.38– 1.22 (m, 8H), 0.96– 0.82 (m, 3H)

**Preparation of compound 4**

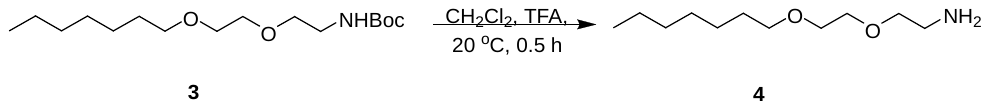

To a solution of *tert*-butyl (2-(2-(heptyloxy)ethoxy)ethyl)carbamate (200 mg, 659 umol, 1.0 equiv) in CH_2_Cl_2_ (3 ml) was added TFA (1 ml). The mixture was stirred at 20 °C for 0.5 h. The reaction mixture was concentrated under reduced pressure to give a residue. 2-(2-(heptyloxy)ethoxy)ethan-1-amine (200 mg, crude, TFA salt) was obtained as a yellow oil and was carried on to the next step without further purification.

**Preparation of compound 6**

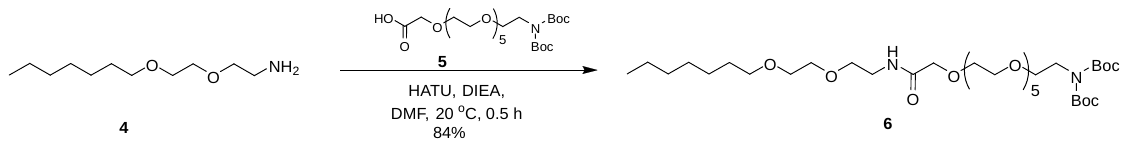

To a solution of 2-(2-(heptyloxy)ethoxy)ethan-1-amine (200 mg, crude, TFA salt) and 5-(*tert*-butoxycarbonyl)-2,2-dimethyl-4-oxo-3,8,11-trioxa-5-azatridecan-13-oic acid (230 mg, 426 μmol, 1.0 equiv) in DMF (2 ml) was added DIEA (296 mg, 2.30 mmol, 0.4 ml, 5.4 equiv) and HATU (324 mg, 852 μmol, 2.0 equiv). The mixture was stirred at 20 °C for 0.5 h. The reaction mixture was quenched by addition H_2_O (0.1 ml), and then concentrated under reduced pressure to give a residue. The residue was purified by *prep*-HPLC (column: Waters Atlantis T_3_ 150*30 mm* 5μm; mobile phase: [water(0.225%FA)–MeCN];B%: 58%–88%, 10 min). *tert*-butyl-(*tert*-butoxycarbonyl)(8-oxo-3,6,12,15-tetraoxa-9-azadocosyl)carbamate (260 mg, 358 μmol, 84% yield) was obtained as a yellow oil.

**^1^H NMR** (400 MHz, CDCl_3_): δ 8.03 (s, 1H), 7.25– 7.14 (m, 1H), 4.02 (s, 2H), 3.82– 3.77 (m, 2H), 3.68– 3.57 (m, 28H), 3.53– 3.48 (m, 4H), 3.45 (t, *J* = 6.8 Hz, 2H), 1.62– 1.56 (m, 2H), 1.51 (s, 18H), 1.35– 1.27 (m, 8H), 0.94– 0.84 (m, 3H)

**Preparation of compound 7**

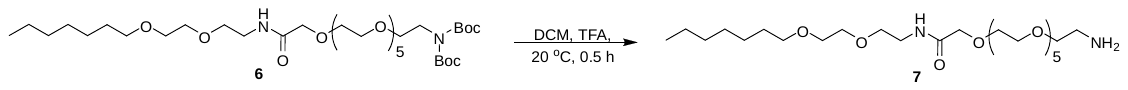

To a solution of *tert*-butyl-(*tert*-butoxycarbonyl)(8-oxo-3,6,12,15-tetraoxa-9-azadocosyl)carbamate (260 mg, 358 µmol, 1.0 equiv) in DCM (3 ml) was added TFA (1 ml). The reaction mixture was concentrated under reduced pressure to give a residue. 2-(2-(2-Aminoethoxy)ethoxy)-*N*-(2-(2-(heptyloxy)ethoxy)ethyl)acetamide (220 mg, crude, TFA salt) was obtained as a yellow oil and used in the next step without further purification.

**Preparation of compound 9**

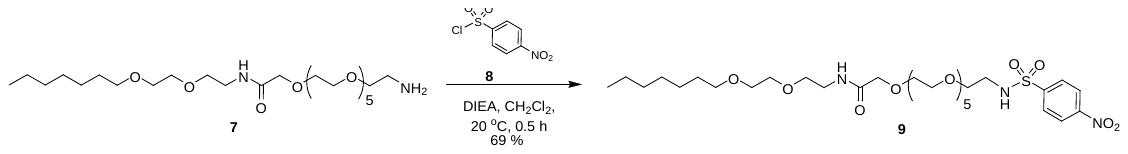

To a solution of 2-(2-(2-aminoethoxy)ethoxy)-*N*-(2-(2-(heptyloxy)ethoxy)ethyl)acetamide (220 mg, crude, TFA salt) in DCM (5 ml) was added DIEA (222 mg, 1.72 mmol, 0.3 ml, 5.0 equiv) and 4-nitrobenzenesulfonyl chloride (114 mg, 516 µmol, 1.5 equiv). The mixture was stirred at 20 °C for 0.5 h. The reaction mixture was concentrated under reduced pressure to give a residue. The residue was purified by *prep*-HPLC (column: 3_Phenomenex Luna C18 75*30 mm*3 µm; mobile phase: [water (0.05%HCl)–MeCN]; B%: 50%–70%,6.5 min). *N*-(2-(2-(Heptyloxy)ethoxy)ethyl)-2-(2-(2-((4-nitrophenyl)sulfonamido)ethoxy)ethoxy)acetamide (170 mg, 239 µmol, 69% yield) was obtained as a yellow oil.

**^1^HNMR** (400 MHz, CD_3_OD): δ 8.36 (d, *J* = 8.7 Hz, 2H), 8.10 (d, *J* = 8.7 Hz, 2H), 7.23– 7.13 (m, 1H), 6.33– 6.08 (m, 1H), 4.02 (s, 2H), 3.74– 3.43 (m, 32H), 3.20 (m, 2H), 1.69–1.52 (m, 2H), 1.36– 1.24 (m, 8H), 0.97– 0.78 (m, 3H)

**LC-MS:** MS (ES^+^): RT = 0.975 min, m/z = 710.2 [M - 55]

**Spectra:**

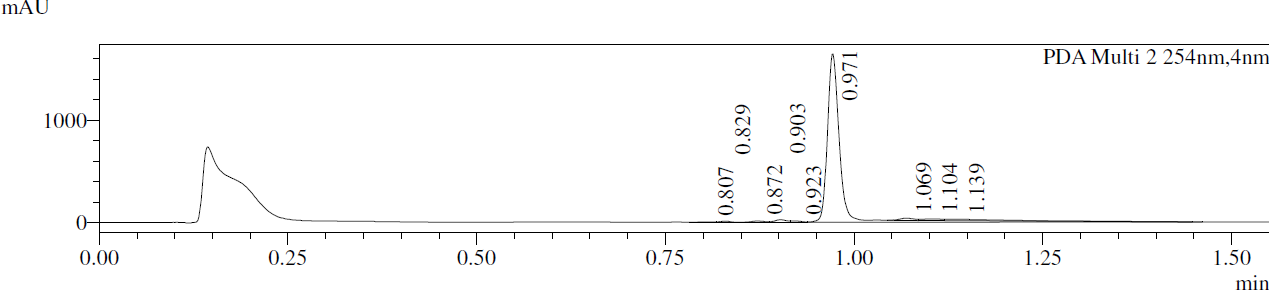

**Preparation of compound 10**

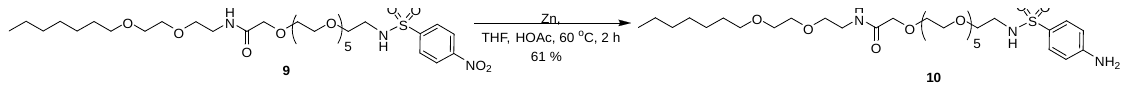
A mixture of *N*-(2-(2-(heptyloxy)ethoxy)ethyl)-2-(2-(2-((4-nitrophenyl)sulfonamido)ethoxy)ethoxy)acetamide (170 mg, 239 µmol, 1.0 equiv), Zn (668 mg, 11.97 mmol, 50 equiv) in THF (5 ml) and AcOH (1 ml) was stirred at 60 °C for 2 h. The reaction mixture was adjusted with aq. sat. NaHCO_3_ to pH = 7–8, and then the mixture was filtered and concentrated under reduced pressure to give a residue. The residue was purified by *prep*-TLC (SiO_2_, CH_2_Cl_2_: MeOH = 10:1). 2-(2-(2-((4-Aminophenyl)sulfonamido)ethoxy)ethoxy)-*N*-(2-(2-(heptyloxy)ethoxy)ethyl)acetamide (100 mg, 147 µmol, 61% yield) was obtained as a colorless oil.

**LC-MS:** MS (ES^+^): RT = 0.933 min, m/z = 680.4 [M + H^+^]

**Spectra:**

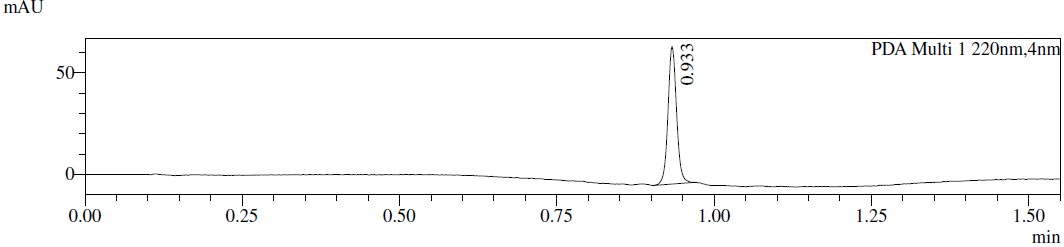

**Preparation of compound 11**

Known compound from *Angew. Chem., Int. Ed*. **2020**, *59*, 13865û13870

**Preparation of HLDA-110**

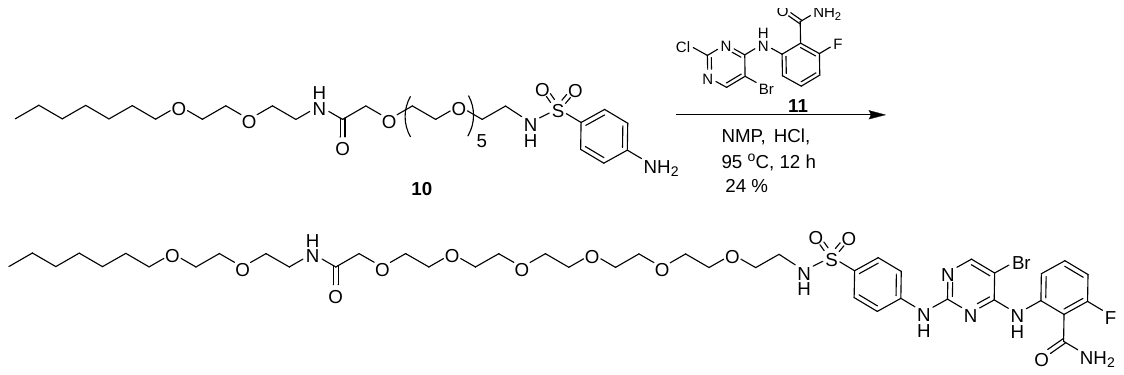

To a solution of 2-(2-(2-((4-aminophenyl)sulfonamido)ethoxy)ethoxy)-*N*-(2-(2-(heptyloxy)ethoxy)ethyl)acetamide (100 mg, 147 µmol, 1.0 equiv) and 2-((5-bromo-2-chloropyrimidin-4-yl)amino)-6-fluorobenzamide (51 mg, 147 µmol, 1.0 equiv) in NMP (3 ml) was added HCl (12 M, 12 µl, 1.0 equiv). The mixture was stirred at 95 °C for 12 h. The reaction mixture was filtered and concentrated under reduced pressure to give a residue. The residue was purified by *prep*-HPLC (column: 3_Phenomenex Luna C18 75*30 mm*3 µm; mobile phase: [water(0.05%HCl)–MeCN]; B%: 53%–73%, 6.5 min) to give desired compound 20-(4-((5-bromo-4-((2-carbamoyl-3-fluorophenyl)amino)pyrimidin-2-yl)amino)phenylsulfonamido)-*N*-(2-(2-(heptyloxy)ethoxy)ethyl)-3,6,9,12,15,18-hexaoxaicosan-1-amide (38 mg, 36 umol, 24% yield, 96% purity, HCl salt) as a yellow gum.

**^1^HNMR** (400 MHz, CD_3_OD): δ 8.35 (s, 1H), 7.95 (d, *J* = 8.4 Hz, 1H), 7.83 (d, *J* = 8.6 Hz, 2H), 7.64 (d, *J* = 8.8 Hz, 2H), 7.57– 7.43 (m, *J* = 6.3, 8.3, 8.3 Hz, 1H), 7.28– 7.14 (m, 1H), 3.98 (s, 2H), 3.67– 3.54 (m, 24H), 3.51– 3.40 (m, 8H), 3.06 (t, *J* = 5.4 Hz, 2H), 1.62– 1.47 (m, 2H), 1.36– 1.25 (m, 8H), 0.89 (m, *J* = 6.8 Hz, 3H)

**LC-MS:** MS (ES^+^): RT = 2.462 min, m/z = 987.7, 989.6 [M + H^+^];

**Spectra:**

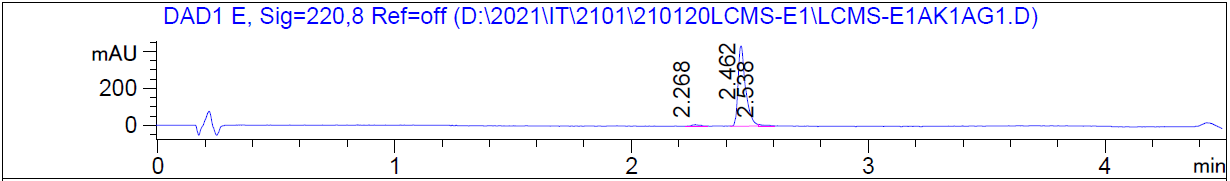

***SERIES 6***

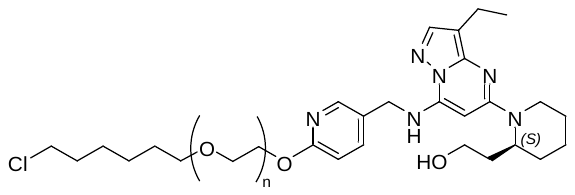

HLDA-117, n=7

HLDA-118, n=9

HLDA-119, n=11

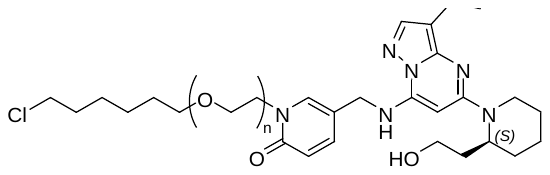

HLDA-114, n=7

HLDA-115, n=9

HLDA-116, n=11

**The synthetic route for HLDA-117**

Part 1:

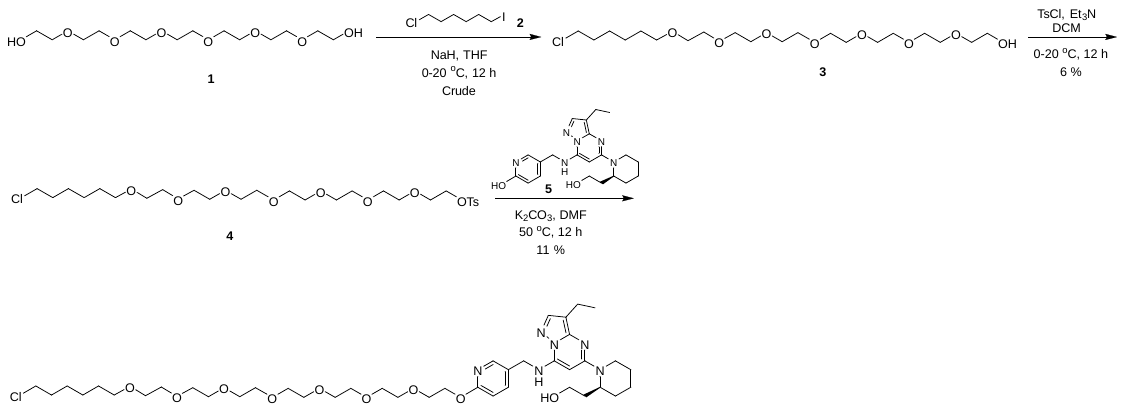

Part 2:

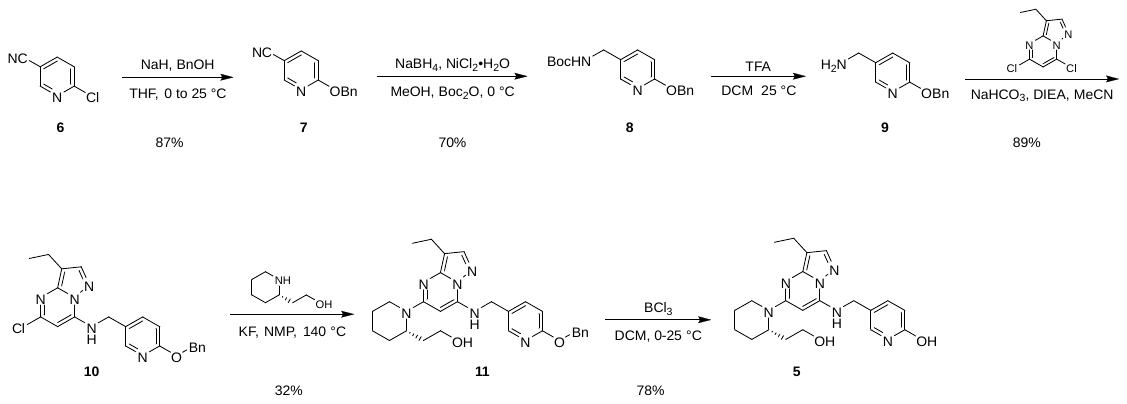

**Preparation of compound 3**

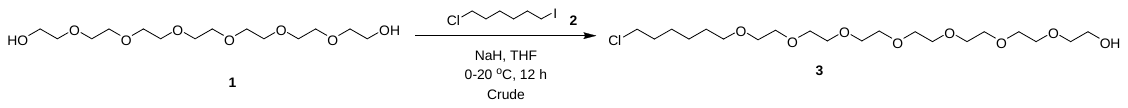

To a solution of 2-[2-[2-[2-[2-[2-(2-hydroxyethoxy)ethoxy]ethoxy]ethoxy]ethoxy]ethoxy]ethanol (2 g, 6.1 mmol, 1.0 equiv) and 1-chloro-6-iodo-hexane (1.5 g, 6.1 mmol, 1.0 equiv) in THF (30 ml) was added NaH (367 mg, 9.2 mmol, 60% purity, 1.5 equiv) at 0 °C. The mixture was stirred at 20 °C for 12 h. The reaction mixture filtered and concentrated under reduced pressure to give a residue. The residue was used for next step directly. Compound 2-[2-[2-[2-[2-[2-[2-(6-chlorohexoxy)ethoxy]ethoxy]ethoxy]ethoxy]ethoxy]ethoxy]ethanol (2.7 g) was obtained as a colorless oil and carried on the the next step without further purification

**LC–MS:** MS (ES^+^): RT = 0.875 min, m/z = 445.2 [M + H^+^];

**Spectra:**

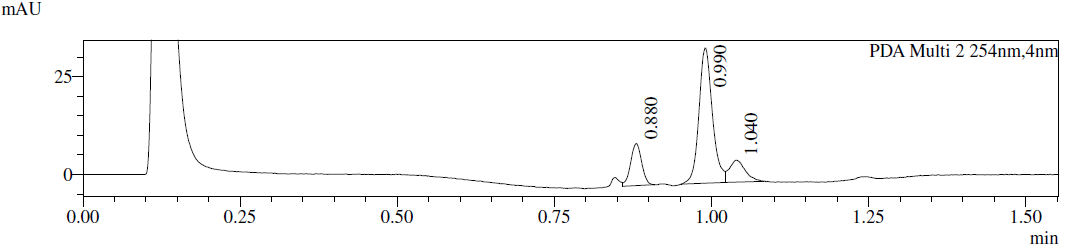

**Preparation of compound 4**

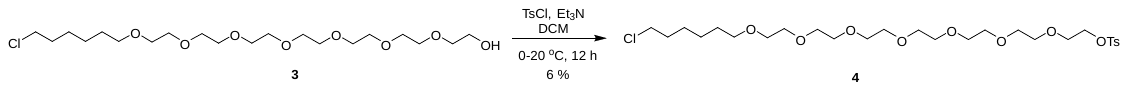

To a solution of 2-[2-[2-[2-[2-[2-[2-(6-chlorohexoxy)ethoxy]ethoxy]ethoxy]ethoxy]ethoxy]ethoxy]ethanol (2.7 g, 6.0 mmol, 1.0 equiv) in CH_2_Cl_2_ (20 ml) was added Et_3_N (4.3 g, 42 mmol, 5.9 mL, 7.0 equiv) and 4-methylbenzenesulfonyl chloride (3.4 g, 18 mmol, 3.0 equiv) at 0 °C. The mixture was stirred at 20 °C for 12 h. The reaction mixture filtered and concentrated under reduced pressure to give a residue. The residue was purified by prep-HPLC (column: Waters Xbridge C18 150*50mm* 10 µm;mobile phase: [water(10mM NH_4_HCO_3_)–MeCN]; B%: 44%–74%, 11 min) to give compound 2-[2-[2-[2-[2-[2-[2-(6-chlorohexoxy)ethoxy]ethoxy]ethoxy]ethoxy]ethoxy]ethoxy]ethyl 4-methylbenzenesulfonate (240 mg, 6% yield).a as a colorless oil.

**^1^H NMR** (400 MHz, CDCl_3_): δ =7.82 (d, *J* = 8.0 Hz, 2H), 7.37 (d, *J* = 8.0 Hz, 2H), 3.71– 3.50 (m, 30H), 2.47 (s, 3H), 1.82– 1.75 (m, 4H), 1.66– 1.56 (m, 2H), 1.49– 1.32 (m, 4H).

**Preparation of HLDA-117**

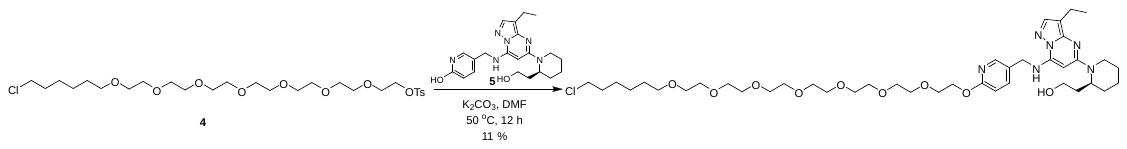

To a solution of 2-[2-[2-[2-[2-[2-[2-(6-chlorohexoxy)ethoxy]ethoxy]ethoxy]ethoxy]ethoxy]ethoxy]ethyl 4-methylbenzenesulfonate (47 mg, 78 µmol, 1.5 equiv) and 5-[[[3-ethyl-5-[(2*S*)-2-(2-hydroxyethyl)-1-piperidyl]pyrazolo[1,5-a]pyrimidin-7-yl]amino]methyl]pyridin-2-ol (20 mg, 50 µmol, 1.0 equiv) in DMF (0.5 ml) was added K_2_CO_3_ (14 mg, 105 µmol, 2.0 equiv). The mixture was stirred at 50 °C for 12 h. The reaction mixture filtered and concentrated under reduced pressure to give a residue. The residue was purified by prep-HPLC (column: Waters Xbridge 150*25 mm* 5 µm; mobile phase: [water(10mM NH_4_HCO_3_–MeCN]; B%: 52%–82%, 9 min) to give 2-[(2*S*)-1-[7-[[6-[2-[2-[2-[2-[2-[2-[2-(6-chlorohexoxy)ethoxy]ethoxy]ethoxy]ethoxy]ethoxy]ethoxy]ethoxy]-3-pyridyl]methylamino]-3-ethyl-pyrazolo[1,5-a]pyrimidin-5-yl]-2-piperidyl]ethanol (5 mg, 11% yield) as a white solid.

**Spectra:**

**LC-MS:** MS (ES^+^): RT = 2.645 min, m/z = 823.4 [M + H+];

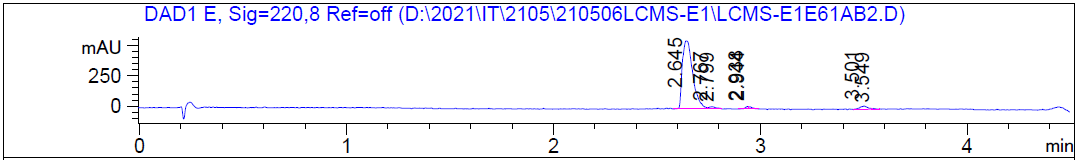

**^1^H NMR** (400 MHz, CDCl_3_): δ = 8.16 (s, 1H), 7.70– 7.55 (m, 2H), 6.82 (d, *J* = 8.4 Hz, 1H), 6.38 (s, 1H), 5.30 (s, 1H), 5.08 (s, 1H), 4.49– 4.45 (m, 4H), 3.93–3.81 (m, 2H), 3.74– 3.64 (m, 23H), 3.60–3.52 (m, 4H), 3.47 (t, *J* = 6.8 Hz, 2H), 3.41– 3.29 (m, 1H), 3.12–2.99 (m, 1H), 2.63– 2.60 (m, 2H), 2.16– 2.05 (m, 1H), 1.81– 1.24 (m, 20H).

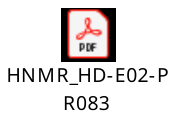

HRMS [C_41_H_67_ClN_6_O_9_] Cal: 823.4731; Obs: 823.47

**Preparation of compound 7**

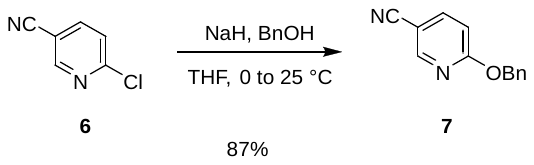

To a solution of BnOH (3.0 g, 28 mmol, 1.3 equiv) in THF (60 ml) was added NaH (1.3 g, 32 mmol, 60% purity, 1.5 equiv) at 0 °C. The mixture was stirred at 25 °C for 0.5 h and 6-chloropyridine-3-carbonitrile (3 g, 22 mmol, 1.0 equiv) was added. The mixture was stirred at 25 °C for 0.5 h, and poured into EtOAc (200 ml). The mixture was stirred for 0.2 h and filtered. The filtrate was concentrated to give 6-benzyloxypyridine-3-carbonitrile (4 g, 88% yield) as a yellow solid.

**LC–MS:** MS (ES^+^): RT = 0.852 min, m/z = 211.1 [M + H^+^];

**Spectra:**

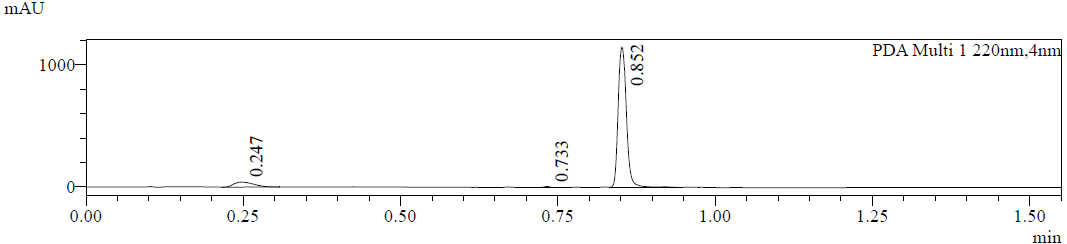

**Preparation of compound 8**

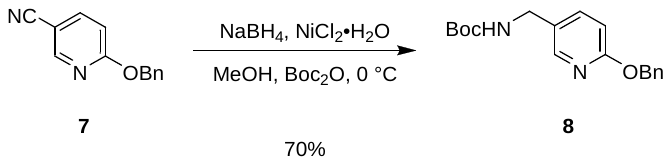

To a mixture of 6-benzyloxypyridine-3-carbonitrile (4.0 g, 19 mmol, 1.0 equiv), NiCl_2_•6H_2_O (904 mg, 3.81 mmol, 0.2 equiv) and Boc_2_O (8.3 g, 38 mmol, 2.0 equiv) in MeOH (60 ml) was added NaBH_4_ (1.83 g, 48 mmol, 2.5 equiv) in small portions at 0 °C. The mixture was stirred at 0 °C for 1 h and concentrated. The residue was poured into EtOAc (200 ml), washed with water (200 ml), brine (200 ml), dried over by Na_2_SO_4_, filtered, and the filtrate was concentrated under reduced pressure. The residue was purified by silica gel chromatography (petroleum ether: EtOAc = 1:0 to 10:1) to give *tert*-butyl-*N*-[(6-benzyloxy-3-pyridyl)methyl]carbamate (4.2 g, 70% yield) as a yellow solid.

**^1^H NMR** (400 MHz, CDCl_3_): δ 8.08 (d, *J* = 2.4 Hz, 1H) 7.57 (d, *J* = 7.2 Hz, 1H), 7.43–7.51 (m, 2H), 7.29–7.43 (m, 4H), 6.80 (d, *J* = 8.4 Hz, 1H), 5.38 (s, 2H), 4.26 (d, *J* = 5.2 Hz, 2H), 1.47 (s, 9H).

**LC–MS:** MS (ES^+^): RT = 0.879 min, m/z = 315.1 [M +H^+^];

**Preparation of compound 9**

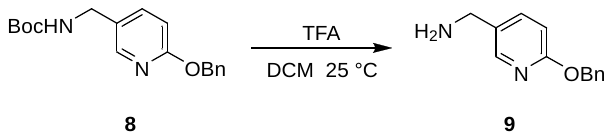

To a solution of tert-butyl *N*-[(6-benzyloxy-3-pyridyl)methyl]carbamate (11 g, 35 mmol, 1.0 equiv) in CH_2_Cl_2_ (70 ml) was added TFA (30.8 g, 270 mmol, 7.7 equiv). The mixture was stirred at 25 °C for 0.5 h. The mixture was concentrated to give crude (6-benzyloxy-3-pyridyl)methanamine (11.4 g, 34.73 mmol, crude, TFA salt) as a yellow oil. The crude product was carried on to the next step without further purification.

**Preparation of compound 10**

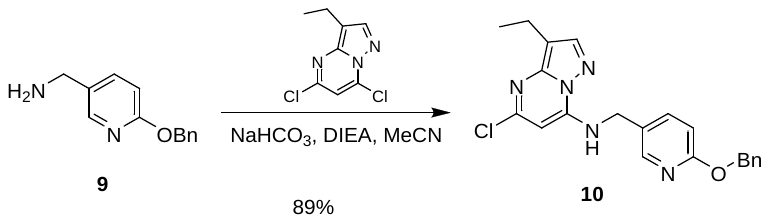

To a solution of (6-benzyloxy-3-pyridyl)methanamine (11.5 g, 35 mmol, 1.5 equiv, TFA salt) and 5,7-dichloro-3-ethyl-pyrazolo[1,5-a]pyrimidine (5.0 g, 23 mmol, 1.0 equiv) in MeCN (100 ml) was added NaHCO_3_ (5.9 g, 70 mmol, 3.0 equiv) and DIEA (3.0 g, 23 mmol, 1.0 equiv). The mixture was stirred at 80 °C for 12 h. The mixture was filtered and concentrated. The residue was purified by column: Kromasil Eternity XT 250*80 mm*10 µm; mobile phase: [water(10 mM NH_4_HCO_3_)–MeCN]; B%: 60%–90%, 20 min to give *N*-[(6-benzyloxy-3-pyridyl)methyl]-5-chloro-3-ethyl-pyrazolo[1,5-a]pyrimidin-7-amine (8.2 g, 89% yield) as a brown solid.

**LC-MS:** MS (ES^+^): RT = 1.040 min, m/z = 394.1 [M + H^+^];

**Preparation of compound 11**

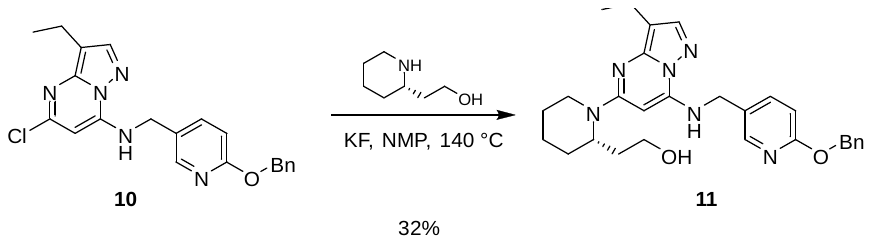

To a solution of *N*-[(6-benzyloxy-3-pyridyl)methyl]-5-chloro-3-ethyl-pyrazolo[1,5-a]pyrimidin-7-amine (2.0 g, 5 mmol, 1.0 equiv) and 2-[(2*S*)-2-piperidyl]ethanol (984 mg, 7.6 mmol, 1.5 equiv) in NMP (2 ml) was added KF (1.48 g, 25 mmol, 5.0 equiv). The mixture was heated to 140 °C and stirred under N_2_ atmosphere for 12 h. The solution was poured into EtOAc (200 ml), washed with water (100 ml), brine (100 ml), dried over by Na_2_SO_4_. The combined organic layer was concentrated under reduced pressure and the residue was purified by silica gel chromatography (petroleum ether : EtOAc = 20:1 to 1:1) to give 2-[(2*S*)-1-[7-[(6-benzyloxy-3-pyridyl)methylamino]-3-ethyl-pyrazolo[1,5-a]pyrimidin-5-yl]-2-piperidyl]ethanol (0.8 g, 32% yield) as a yellow oil.

**LC-MS:** MS (ES^+^): RT = 0.818 min, m/z = 487.2 [M +H^+^];

**Preparation of compound 5**

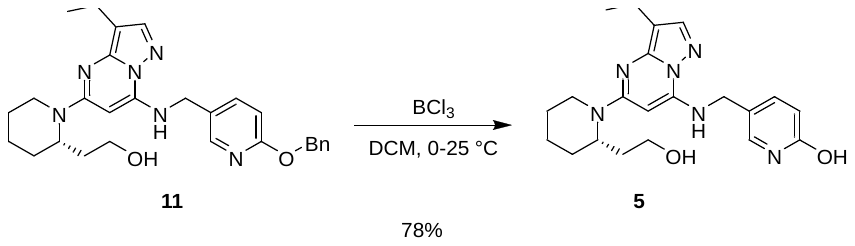

To a solution of 2-[(2*S*)-1-[7-[(6-benzyloxy-3-pyridyl)methylamino]-3-ethyl-pyrazolo[1,5-a]pyrimidin-5-yl]-2-piperidyl]ethanol (580 mg, 1.2 mmol, 1.0 equiv) in CH_2_Cl_2_ (10 ml) was added BCl_3_ (1 M, 10 mL, 1.0 equiv) at 0 °C. The mixture was stirred at 25 °C for 1 h and quenched with saturated sodium bicarbonate (30 mll) and ammonium hydroxide (3 ml) at 0 °C. The mixture was stirred at 25 °C for 1 h and extracted with CH_2_Cl_2_ (3 x 30 ml). The combined organic layers were washed with brine (50 ml), dried over sodium sulfate, filtered, and concentrated. The residue was purified by silica column chromatography on silica gel (CH_2_Cl_2_ : MeOH from 50/1 to 5/1) to give 5-[[[3-ethyl-5-[(*2S*)-2-(2-hydroxyethyl)-1-piperidyl]pyrazolo[1,5-a]pyrimidin-7-yl]amino]methyl]pyridin-2-ol (375 mg, 78% yield) as an off-white solid.

**^1^H NMR** (400 MHz, DMSO-*d*_6_): δ 11.43 (s, 1H), 7.71 (t, *J* = 6.4 Hz, 1H), 7.64 (s, 1H), 7.53 (dd, *J* = 2.4, 9.6 Hz, 1H), 7.46 (d, *J* = 2.4 Hz, 1H), 6.30 (d, *J* = 9.6 Hz, 1H), 5.59 (s, 1H), 4.72 (t, *J* = 5.4 Hz, 1H), 4.61 (s, 1H), 4.34–4.19 (m, 3H), 3.43–3.35 (m, 2H), 2.84 (t, *J* = 12.4 Hz, 1H), 2.49–2.44 (m, 2H), 1.89–1.76 (m, 1H), 1.70–1.54 (m, 6H), 1.43–1.30 (m, 1H), 1.17 (t, *J* = 7.6 Hz, 3H)

**LC-MS:** MS (ES^+^): RT = 2.213 min, m/z = 397.2 [M + H^+^];

**Spectra:**

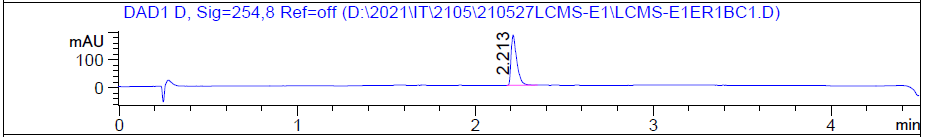

**The synthetic route for HLDA-118**

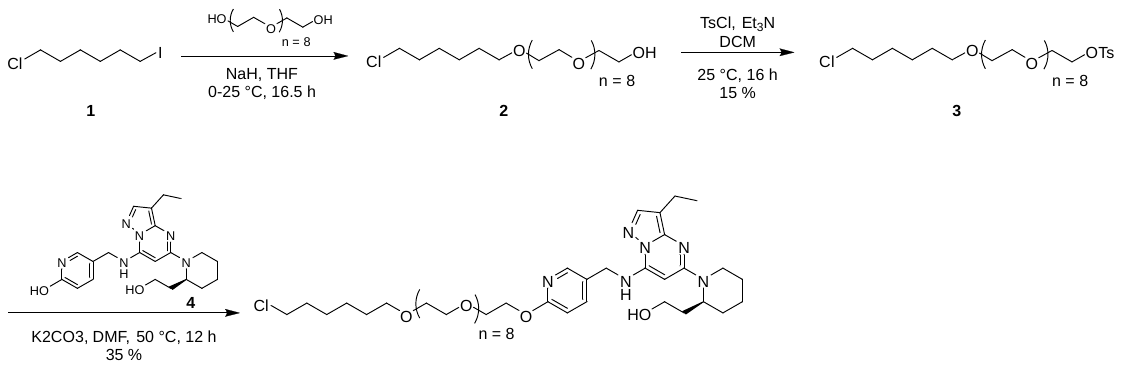

**Preparation of 2**

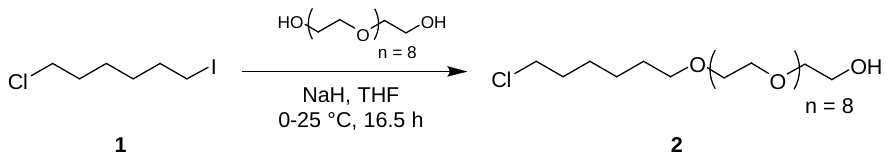

To a solution of 2-[2-[2-[2-[2-[2-[2-[2-(2-hydroxyethoxy)ethoxy]ethoxy]ethoxy]ethoxy]ethoxy]ethoxy]ethoxy]ethanol (2 g, 4.83 mmol, 1.0 equiv) in THF (20 ml) was added NaH (289 mg, 7.24 mmol, 60 % purity, 1.5 equiv) at 0 °C, the mixture was stirred at 25 °C for 0.5 h, then 1-chloro-6-iodo-hexane (1.55 g, 6.27 mmol, 1.3 equiv) was added. The mixture was stirred at 25 °C for 16 h and quenched with HCl/dioxane (1 M, 8 ml). The mixture was concentrated to give 2-[2-[2-[2-[2-[2-[2-[2-[2-(6-chlorohexoxy)ethoxy]ethoxy]ethoxy]ethoxy]ethoxy]ethoxy]ethoxy]ethoxy]ethanol (2.57 g,). The crude material was used in the next step without further purification

**LC–MS:** MS (ES^+^): RT = 0.848 min, m/z = 550.3 [M + H_2_O];

**Preparation of 3**

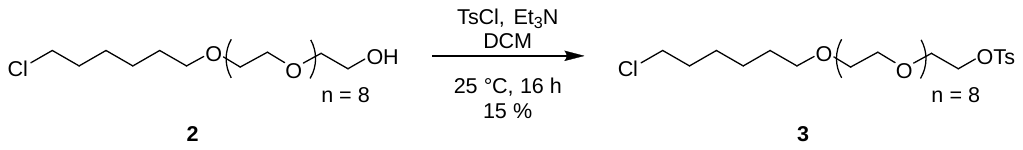

To a solution of 2-[2-[2-[2-[2-[2-[2-[2-[2-(6-chlorohexoxy)ethoxy]ethoxy]ethoxy]ethoxy]ethoxy]ethoxy]ethoxy]ethoxy]ethanol (2.57 g, 4.82 mmol, 1.0 equiv) in CH_2_Cl_2_ (20 ml) was added TEA (2.44 g, 24.1 mmol, 5.0 equiv) and 4-methylbenzenesulfonyl chloride (2.76 g, 14.4 mmol, 3.0 equiv). The reaction mixture was stirred at 25 °C for 16 h. The mixture was diluted with water (30 ml) and extracted with CH_2_Cl_2_ (3 x 30 ml). The combined organic layers were dried over Na_2_SO_4_, filtered, and the filtrate was concentratedunder reduced pressure. The residue was purified by *prep*-HPLC (column: Waters Xbridge C18 150*50 mm* 10 µm; mobile phase: [water (10 mM NH_4_HCO_3_)–MeCN]; B%: 42%–72%, 11 min) to give 2-[2-[2-[2-[2-[2-[2-[2-[2-(6-chlorohexoxy)ethoxy]ethoxy]ethoxy]ethoxy]ethoxy]ethoxy]ethoxy]ethoxy]ethyl 4-methylbenzenesulfonate (500 mg, 15 % yield).

**^1^H NMR** (400 MHz, CDCl_3_): δ 7.80 (d, *J* = 8.4 Hz, 2H), 7.35 (d, *J* = 8.4 Hz, 2H), 4.20– 4.12 (m, 2H), 3.71– 3.62 (m, 28H), 3.61– 3.57 (m, 6H), 3.56– 3.51 (m, 2H), 3.48– 3.43 (m, 2H), 2.45 (s, 3H), 1.82– 1.73 (m, 2H), 1.65– 1.54 (m, 2H), 1.51– 1.32 (m, 4H).

**LC–MS:** MS (ES^+^): RT = 0.933 min, m/z = 704.3 [M + H_2_O];

**Preparation of compound 4**

See HLDA-117

**Preparation of HLDA-118**

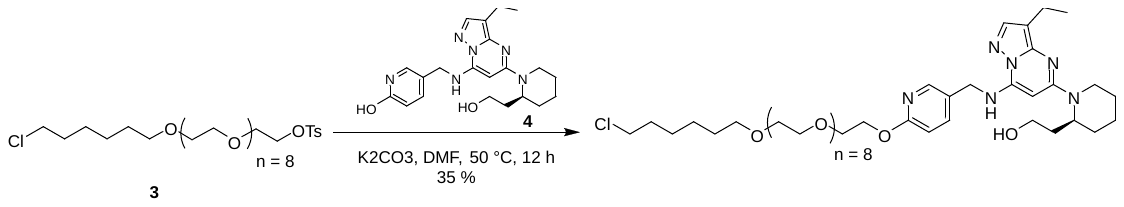

To a solution of 5-[[[3-ethyl-5-[(2S)-2-(2-hydroxyethyl)-1-piperidyl]pyrazolo[1,5-a]pyrimidin-7-yl]amino]methyl]pyridin-2-ol (50 mg, 126 µmol, 1.0 equiv) and 2-[2-[2-[2-[2-[2-[2-[2-[2-(6-chlorohexoxy)ethoxy]ethoxy]ethoxy]ethoxy]ethoxy]ethoxy]ethoxy]ethoxy]ethyl 4-methylbenzenesulfonate (130 mg, 189 µmol, 1.5 equiv) in DMF (1 ml) was added K_2_CO_3_ (34 mg, 252 µmol, 2.0 equiv), the mixture was stirred at 50 °C for 12 h. The mixture was filtered and concentrated. The residue was purified by prep-HPLC (column: Waters Xbridge 150*25 mm* 5 µm;mobile phase: [water(10 mM NH_4_HCO_3_)–MeCN]; B%: 55%–85%, 9 min) to give 2-[(2*S*)-1-[7-[[6-[2-[2-[2-[2-[2-[2-[2-[2-[2-(6-chlorohexoxy)ethoxy]ethoxy]ethoxy]ethoxy]ethoxy]ethoxy]ethoxy]ethoxy]ethoxy]-3-pyridyl]methylamino]-3-ethyl-pyrazolo[1,5-a]pyrimidin-5-yl]-2-piperidyl]ethanol (40 mg, 35 % yield).

**^1^H NMR** (400 MHz, MeOD): δ 8.23– 8.14 (m, 1H), 7.81– 7.73 (m, 1H), 7.68– 7.59 (m, 1H), 6.86– 6.79 (m, 1H), 5.57– 5.49 (m, 1H), 4.54 (s, 2H), 4.46– 4.36 (m, 2H), 4.09– 3.97 (m, 1H), 3.87– 3.80 (m, 2H), 3.70– 3.41 (m, 39H), 3.07– 2.94 (m, 1H), 2.60– 2.51 (m, 2H), 2.14 –2.05 (m, 1H), 1.79– 1.62 (m, 8H), 1.61– 1.49 (m, 3H), 1.47– 1.32 (m, 4H), 1.23 (t, *J* = 7.6 Hz, 3H).

**LC–MS:** MS (ES^+^): RT = 2.986 min, m/z = 911.5 [M + H^+^];

**Spectra:**

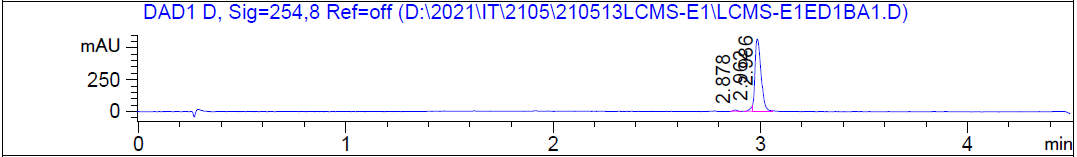

HRMS [C_45_H_75_ClN_6_O_11_] Cal: 911.5255; Obs: 911.5217

**The synthetic route for HLDA-119**

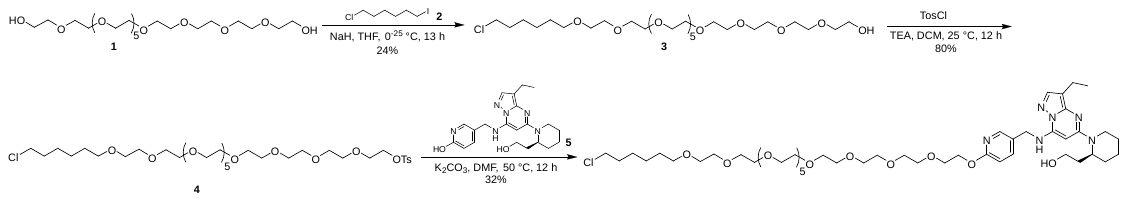

**Preparation of compound 2**

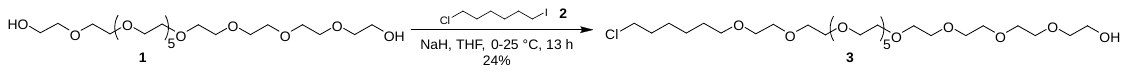

To a solution of 2-[2-[2-[2-[2-[2-[2-[2-[2-[2-(2-hydroxyethoxy)ethoxy]ethoxy]ethoxy]ethoxy]ethoxy]

ethoxy]ethoxy]ethoxy]ethoxy]ethanol (2.0 g, 3.98 mmol, 1.0 equiv) in THF (40 ml) was added NaH (239 mg, 5.97 mmol, 60% purity, 1.5 equiv) at 0 °C and stirred at 25 °C for 1 h. Then 1-chloro-6-iodo-hexane (980 mg, 3.98 mmol, 1.0 equiv) was added and the mixture was stirred at 25 °C for 12 h. The reaction mixture was quenched by addition of NH_4_Cl (5 ml) and then diluted with H_2_O (30 ml) and extracted with EtOAc (3 x 30 ml). The combined organic layers were washed with brine (2 x 30 mL), dried over Na_2_SO_4_, filtered and concentrated under reduced pressure to give a residue. The residue was purified by column chromatography (SiO_2_, **CH_2_Cl_2_: MeOH** from **1/0** to 10**/1**) to give 2-[2-[2-[2-[2-[2-[2-[2-[2-[2-[2-(6-chlorohexoxy)ethoxy]ethoxy]ethoxy]ethoxy]ethoxy]ethoxy]ethoxy]ethoxy]ethoxy]ethoxy]ethanol (600 mg, 24% yield) as a colorless oil.

**Preparation of compound 3**

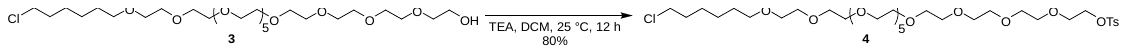

To a solution of 2-[2-[2-[2-[2-[2-[2-[2-[2-[2-[2-(6-chlorohexoxy)ethoxy]ethoxy]ethoxy]ethoxy]ethoxy]ethoxy]ethoxy]ethoxy]ethoxy]ethoxy]ethanol (600 mg, 966 µmol, 1.0 equiv) and TEA (293 mg, 2.90 mmol, 3.0 equiv) in CH_2_Cl_2_ (5 ml) was added 4-methylbenzenesulfonyl chloride (368 mg, 1.93 mmol, 2.0 equiv) at 0 °C. The mixture was stirred at 25 °C for 12 h. The reaction mixture was diluted with H_2_O (30 ml) and extracted with EtOAc (3 x 20 ml). The combined organic layers were washed with brine (2 x 20 ml), dried over Na_2_SO_4_, filtered, and concentrated under reduced pressure to give a residue. The residue was purified by column chromatography (SiO_2_, **DCM: MeOH** from **1/0** to 10**/1**) to give 2-[2-[2-[2-[2-[2-[2-[2-[2-[2-[2-(6-chlorohexoxy)ethoxy]ethoxy]ethoxy]ethoxy]ethoxy]ethoxy]ethoxy]ethoxy]ethoxy]ethoxy]ethyl 4-methylbenzenesulfonate (600 mg, 80% yield) as a yellow oil.

**LC–MS:** MS (ES^+^): RT = 0.973 min, m/z = 775.4[M + H^+^];

**Preparation of compound 5**

See HLDA-117

**Preparation of HLDA-119**

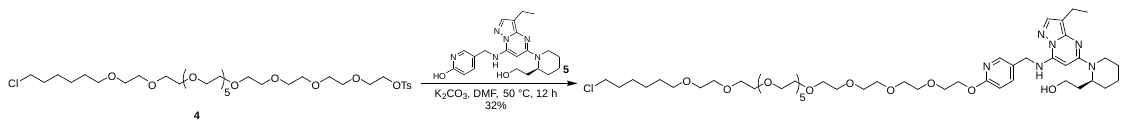

To a solution of 2-[2-[2-[2-[2-[2-[2-[2-[2-[2-[2-(6-chlorohexoxy)ethoxy]ethoxy]ethoxy]ethoxy]ethoxy]ethoxy]ethoxy]ethoxy]ethoxy]ethoxy]ethyl 4-methylbenzenesulfonate (117 mg, 151 µmol, 1.2 equiv) and5-[[[3-ethyl-5-[(2*S*)-2-(2-hydroxyethyl)-1-piperidyl]pyrazolo[1,5-a]pyrimidin-7-yl]amino]methyl]pyridin-2-ol (50 mg, 126 µmol, 1.0 equiv) in DMF (2 ml) was added K_2_CO_3_ (35 mg, 2525 µmol, 2.0 equiv). The mixture was stirred at 50 °C for 12 h. The reaction mixture was filtered and concentrated under reduced pressure to give a residue. The residue was purified by *prep*-HPLC (column: Waters Xbridge 150*25 mm*5 µm; mobile phase: [water(NH_4_HCO_3_)–MeCN]; B%: 58%–88%, 9 min) to give 2-[(2*S*)-1-[7-[[6-[2-[2-[2-[2-[2-[2-[2-[2-[2-[2-[2-(6-chlorohexoxy)ethoxy]ethoxy]ethoxy]ethoxy]ethoxy]ethoxy]ethoxy]ethoxy]ethoxy]ethoxy]ethoxy]-3-pyridyl]methylamino]-3-ethyl-pyrazolo[1,5-a]pyrimidin-5-yl]-2-piperidyl]ethanol (42 mg, 32% yield) as a brown gum.

**^1^H NMR** (400 MHz, CDCl_3_): δ 8.12 (d, 1H, *J* = 2.2 Hz), 7.61 (s, 1H), 7.59 (d, 1H, *J* = 8.6 Hz), 6.78 (d, 1H, *J* = 8.4 Hz), 6.38 (t, 1H, *J* = 5.4 Hz), 5.27 (s, 1H), 5.10–5.01 (m, 1H), 4.49– 4.39 (m, 4H), 3.86– 3.77 (m, 2H), 3.71– 3.68 (m, 2H), 3.68– 3.59 (m, 38H), 3.58– 3.54 (m, 3H), 3.51 (t, 2H, *J* = 6.8 Hz), 3.44 (t, 2H, *J* = 6.7 Hz), 3.31 (t, 1H, *J* = 11.8 Hz), 3.07– 2.95 (m, 1H), 2.58 (d, 2H, *J* = 2.9, 7.5 Hz), 1.82– 1.31 (m, 16H), 1.23 (t, 3H, *J* = 7.6 Hz)

**LC-MS:** MS (ES^+^): RT = 2.992 min, m/z = 999.6 [M + H^+^];

**Spectra:**

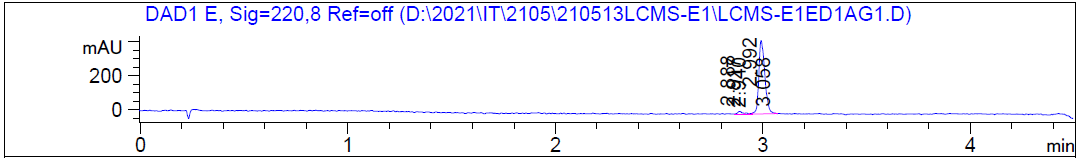

HRMS [C_49_H_83_ClN_6_O_13_] Cal: 358.1624; Obs: 358.1614

**The synthetic route for HLDA-114**

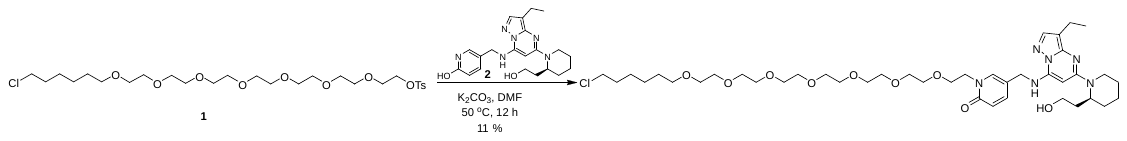

**Preparation of compound 1**

See HLDA-117

**Preparation of compound 2**

See HLDA-117

**Preparation of HLDA-114**

To a solution of 2-[2-[2-[2-[2-[2-[2-(6-chlorohexoxy)ethoxy]ethoxy]ethoxy]ethoxy]ethoxy]ethoxy]ethyl 4-methylbenzenesulfonate (47 mg, 78 µmol, 1.5 equiv) and 5-[[[3-ethyl-5-[(2*S*)-2-(2-hydroxyethyl)-1-piperidyl]pyrazolo[1,5-a]pyrimidin-7-yl]amino]methyl]pyridin-2-ol (20 mg, 50 µmol, 1.0 equiv) in DMF (0.5 ml) was added K_2_CO_3_ (14 mg, 105 µmol, 2.0 equiv). The mixture was stirred at 50 °C for 12 h. The reaction mixture filtered and concentrated under reduced pressure to give a residue. The residue was purified by prep-HPLC (column: Waters Xbridge 150*25 mm* 5 µm; mobile phase: [water(10 mM NH_4_HCO_3_)–MeCN]; B%: 52%–82%, 9 min) to give 2-[(2S)-1-[7-[[6-[2-[2-[2-[2-[2-[2-[2-(6-chlorohexoxy)ethoxy]ethoxy]ethoxy]ethoxy]ethoxy]ethoxy]ethoxy]-3-pyridyl]methylamino]-3-ethyl-pyrazolo[1,5-a]pyrimidin-5-yl]-2-piperidyl]ethanol (5 mg, 11% yield) as a white solid.

**Spectra:**

**LC-MS:** MS (ES^+^): RT = 2.558 min, m/z = 823.4 [M + H+]; M

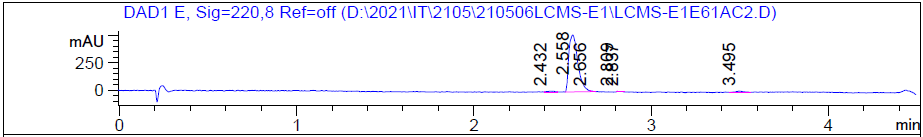

**^1^H NMR** (400 MHz, CDCl_3_): δ 7.64 (s, 1H), 7.52 (d, *J* = 2.0 Hz, 1H), 7.39– 7.36 (m, 1H), 6.61 (d, *J* = 9.2 Hz, 1H), 6.35 (s, 1H), 5.33 (s, 1H), 5.19– 4.99 (m, 1H), 4.29 (d, *J* = 5.2 Hz, 2H), 4.14 (t, *J* = 4.8 Hz, 2H), 3.82– 3.71 (m, 3H), 3.68– 3.42 (m, 29H), 3.40– 3.32 (m, 1H), 3.11– 2.98 (m, 1H), 2.65– 2.58 (m, 2H), 2.10 (d, *J* = 2.0 Hz, 1H), 1.83– 1.23 (m, 19H).

HRMS [C_41_H_67_ClN_6_O_9_] Cal: 823.4731; Obs: 823.4695

**The synthetic route for HLDA-115**

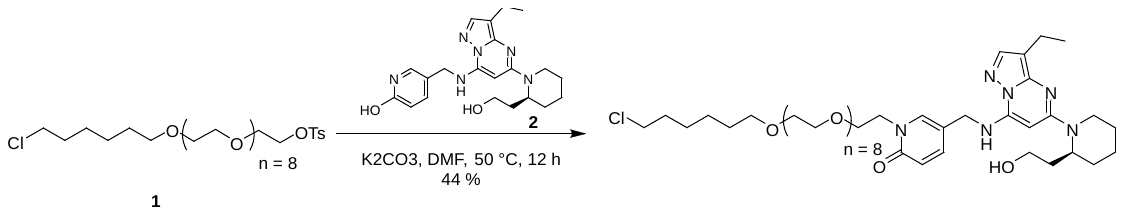

**Preparation of compound 1**

See HLDA-118

**Preparation of compound 2**

See HLDA-117

**Preparation of HLDA-115**

To a solution of 5-[[[3-ethyl-5-[(2*S*)-2-(2-hydroxyethyl)-1-piperidyl]pyrazolo[1,5-a]pyrimidin-7-yl]amino]methyl]pyridin-2-ol (50 mg, 126 µmol, 1.0 equiv) and 2-[2-[2-[2-[2-[2-[2-[2-[2-(6-chlorohexoxy)ethoxy]ethoxy]ethoxy]ethoxy]ethoxy]ethoxy]ethoxy]ethoxy]ethyl 4-methylbenzenesulfonate (130 mg, 189 µmol, 1.5 equiv) in DMF (1 ml) was added K_2_CO_3_ (34 mg, 252 µmol, 2.0 equiv), the mixture was stirred at 50 °C for 12 h. The mixture was filtered and concentrated to give a residue. The residue was purified by *prep*-HPLC (column: Waters Xbridge 150*25 mm* 5 µm;mobile phase: [water(10 mM NH_4_HCO_3_)-MeCN]; B%: 55%–85%, 9 min) to give 1-[2-[2-[2-[2-[2-[2-[2-[2-[2-(6-chlorohexoxy)ethoxy]ethoxy]ethoxy]ethoxy]ethoxy]ethoxy]ethoxy]ethoxy]ethyl]-5-[[[3-ethyl-5-[(2*S*)-2-(2-hydroxyethyl)-1-piperidyl]pyrazolo[1,5-a]pyrimidin-7-yl]amino]methyl]pyridin-2-one (50 mg, 44 % yield).

**^1^H NMR** (400 MHz, MeOD): δ 7.80– 7.69 (m, 2H), 7.66– 7.59 (m, 1H), 6.60– 6.53 (m, 1H), 5.63 (s, 1H), 4.47 (s, 2H), 4.21– 4.14 (m, 2H), 4.10– 4.00 (m, 1H), 3.76 (t, *J* = 4.8 Hz, 2H), 3.67–3.51 (m, 34H), 3.50– 3.47 (m, 5H), 3.20– 3.08 (m, 1H), 2.63– 2.54 (m, 2H), 2.21– 2.10 (m, 1H), 1.83– 1.67 (m, 8H), 1.63– 1.52 (m, 3H), 1.49– 1.34 (m, 4H), 1.25 (t, *J* = 7.6 Hz, 3H).

**LC-MS:** MS (ES^+^): RT = 2.906 min, m/z = 911.5 [M + H^+^];

**Spectra:**

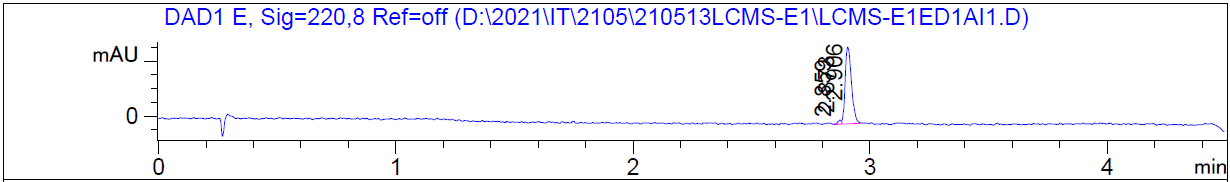

HRMS [C_45_H_75_ClN_6_O_11_] Cal: 911.5255; Obs: 911.5215

**The synthetic route for HLDA-116**

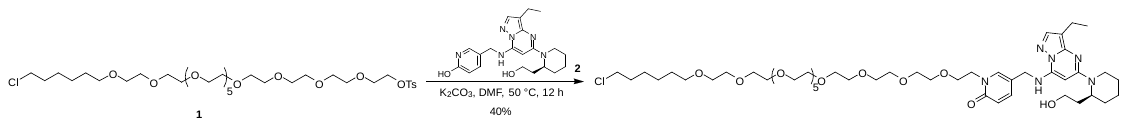

**Preparation of compound 1**

See HLDA-119

**Preparation of compound 2**

See HLDA-117

**Preparation of HLDA-116**

To a solution of 2-[2-[2-[2-[2-[2-[2-[2-[2-[2-[2-(6-chlorohexoxy)ethoxy]ethoxy]ethoxy]ethoxy]ethoxy]ethoxy]ethoxy]ethoxy]ethoxy]ethoxy]ethyl 4-methylbenzenesulfonate (117 mg, 151 µmol, 1.2 equiv) and5-[[[3-ethyl-5-[(2*S*)-2-(2-hydroxyethyl)-1-piperidyl]pyrazolo[1,5-a]pyrimidin-7-yl]amino]methyl]pyridin-2-ol (50 mg, 126 µmol, 1.0 equiv) in DMF (2 ml)was added K_2_CO_3_ (35 mg, 253 µmol, 2.0 equiv). The mixture was stirred at 50 °C for 12 h. The reaction mixture was filtered and concentrated under reduced pressure to give a residue. The residue was purified by *prep*-HPLC (column: Waters Xbridge 150*25 mm*5 µm; mobile phase: [water(NH_4_HCO_3_)–ACN]; B%: 58%–88%, 9 min) to give 1-[2-[2-[2-[2-[2-[2-[2-[2-[2-[2-[2-(6-chlorohexoxy)ethoxy]ethoxy]ethoxy]ethoxy]ethoxy]ethoxy]ethoxy]ethoxy]ethoxy]ethoxy]ethyl]-5-[[[3-ethyl-5-[(2*S*)-2-(2-hydroxyethyl)-1-piperidyl]pyrazolo[1,5-a]pyrimidin-7-yl]amino]methyl]pyridin-2-one (51 mg, 40% yield) as a colorless oil.

**H NMR** (400 MHz, CDCl_3_): δ 7.62 (s, 1H), 7.51 (d, 1H, *J* = 2.0 Hz), 7.36 (d, 1H, *J* = 9.4 Hz), 6.57 (d, 1H, *J* = 9.4 Hz), 6.37 (s, 1H), 5.30 (s, 1H), 5.12 - 5.00 (m, 1H), 4.26 (d, 2H, *J* = 5.4 Hz), 4.12 (t, 2H, *J* = 4.8 Hz), 3.83– 3.70 (m, 4H), 3.64– 3.59 (m, 36H), 3.55 (s, 4H), 3.53– 3.49 (m, 3H), 3.44 (t, 2H, *J* = 6.8 Hz), 3.37– 3.29 (m, 1H), 3.09– 2.98 (m, 1H), 2.58 (d, 2H, *J* = 7.6 Hz), 1.81– 1.51 (m, 12H), 1.48– 1.41 (m, 2H), 1.40– 1.33 (m, 2H), 1.23 (t, 3H, *J* = 7.6 Hz)

**LC–MS:** MS (ES^+^): RT = 2.909 min, m/z = 999.5 [M + H^+^];

**Spectra:**

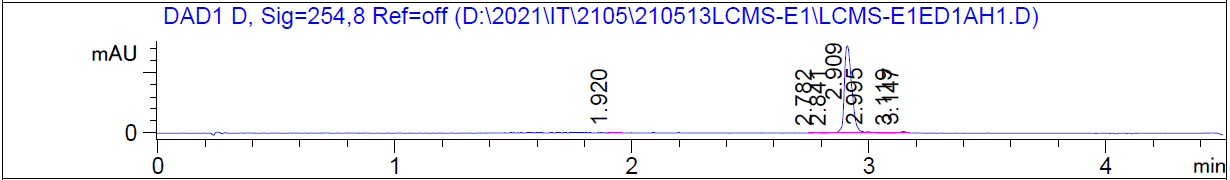

HRMS [C_49_H_83_ClN_6_O_13_] Cal: 999.5779; Obs: 999.5736

***SERIES 7***

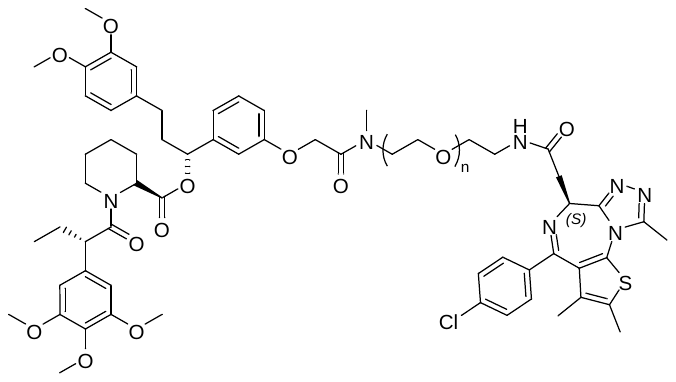

HLDA-221, n=2

HLDA-222, n=4

HLDA-223, n=6

**The synthetic route for HLDA-221**

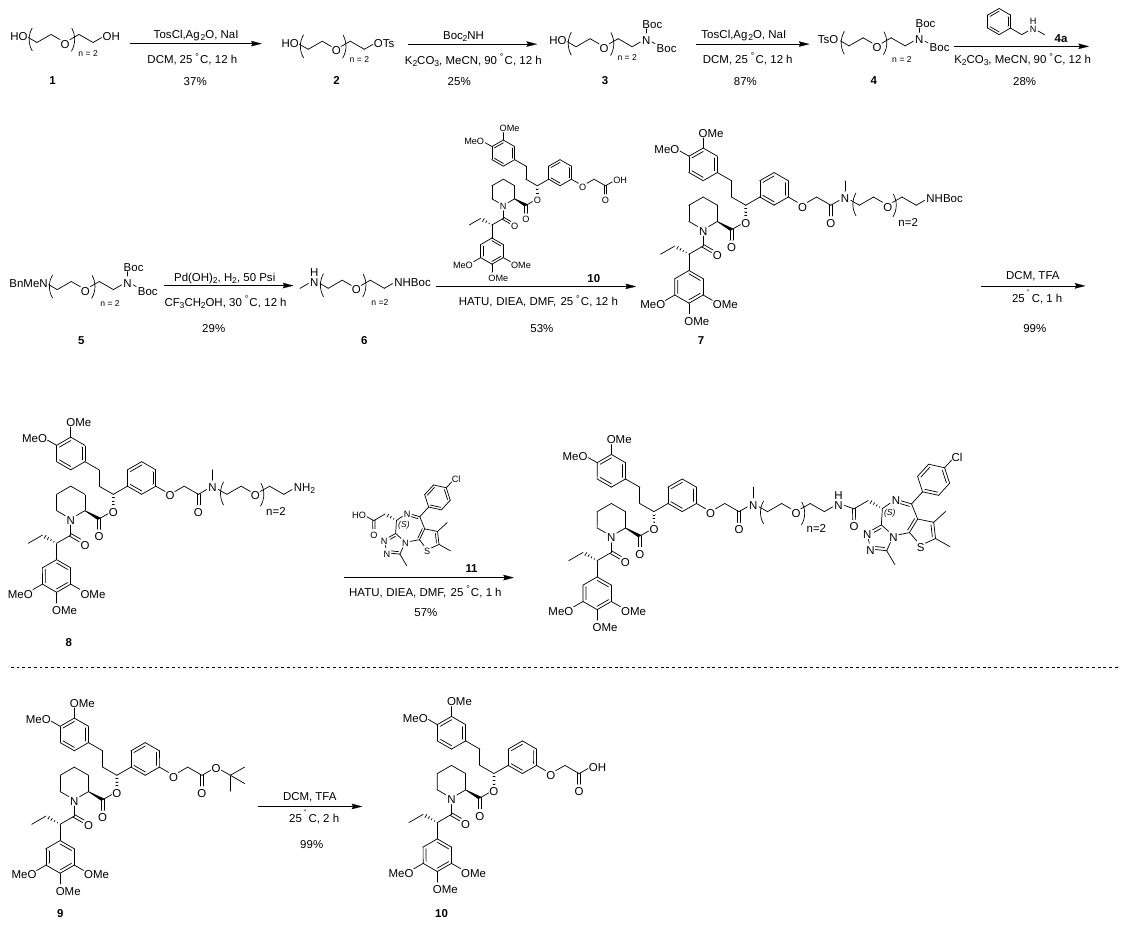

**Preparation of compound 2**

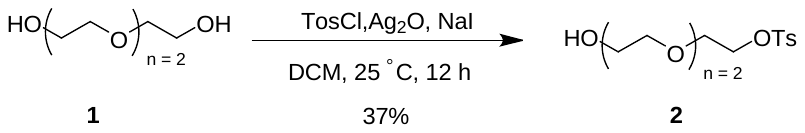

To a solution of 2-[2-(2-hydroxyethoxy)ethoxy]ethanol (20.0 g, 133 mmol, 17.9 mL, 1.0 equiv) and 4-methylbenzenesulfonyl chloride (30.5 g, 160 mmol, 1.2 equiv) in CH_2_Cl_2_ (150 ml) was added NaI (22 g, 147 mmol, 1.1 equiv) and Ag_2_O (46.3 g, 200 mmol, 1.5 equiv). The mixture was stirred at 25 °C for 12 h. The mixture was filtered and concentrated. The residue was purified by silica column chromatography on silica gel (Petroleum ether: EtOAc from 20/1 to 1/2) to give 2-[2-(2-hydroxyethoxy)ethoxy]ethyl 4-methylbenzenesulfonate (15 g, 37% yield).

**^1^H NMR** (400 MHz, CDCl_3_): δ 7.74 (d, *J* = 8.4 Hz, 2H), 7.27 (s, 2H), 4.17-4.06 (m, 2H), 3.70–3.60 (m, 4H), 3.53–3.47 (m, 2H), 2.69 (s, 1H), 2.39 (s, 3H).

**LC–MS:** MS (ES^+^): RT = 0.783 min, m/z = 305.1 [M + H^+^];

**Spectra:**

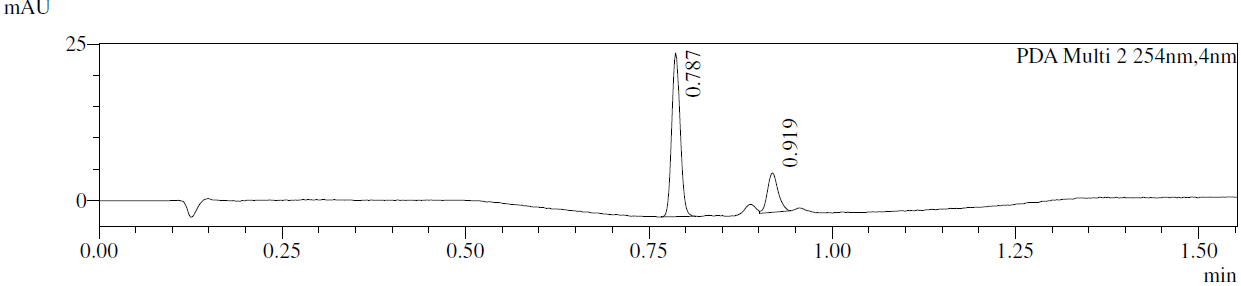

**Preparation of compound 3**

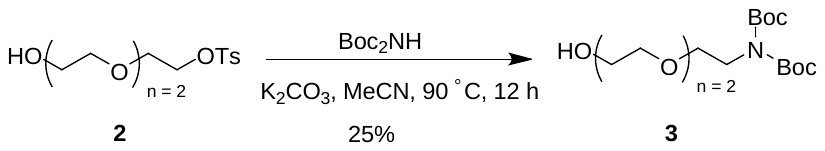

To a solution of 2-[2-(2-hydroxyethoxy)ethoxy]ethyl 4-methylbenzenesulfonate (5.68 g, 18.7 mmol, 1.0 equiv) in MeCN (50 ml) and *tert*-butyl *N*-*tert*-butoxycarbonylcarbamate (5.27 g, 24.3 mmol, 1.3 equiv) was added K_2_CO_3_ (6.45 g, 46.7 mmol, 2.5 equiv). The mixture was stirred at 90 °C for 12 h. The mixture was filtered and concentrated. The residue was purified by silica column chromatography on silica gel (Petroleum ether: EtOAc from 20/1 to 1/2) to give *tert*-butyl *N-tert*-butoxycarbonyl-*N*-[2-[2-(2-hydroxyethoxy)ethoxy]ethyl]carbamate (1.6 g, 25% yield).

**^1^H NMR** (400 MHz, CDCl_3_): δ 3.82–3.77 (m, 2H), 3.74–3.67 (m, 2H), 3.66–3.54 (m, 8H), 2.55 (s, 1H), 1.52–1.48 (m, 18H)

**Preparation of compound 4**

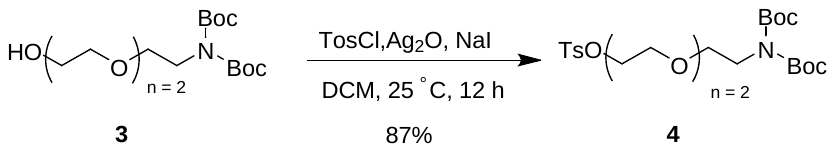

To a solution of *tert*-butyl *N-tert*-butoxycarbonyl-*N*-[2-[2-(2-hydroxyethoxy)ethoxy]ethyl]carbamate (1.60 g, 4.58 mmol, 1.0 equiv) and 4-methylbenzenesulfonyl chloride (1.05 g, 5.49 mmol, 1.2 equiv) in CH_2_Cl_2_ (10 ml) was added Ag_2_O (1.27 g, 5.49 mmol, 1.2 equiv) and NaI (0.76 g, 5.04 mmol, 1.1 equiv). The mixture was stirred at 25 °C for 12 h. The mixture was filtered and concentrated under reduced pressure. The residue was purified by silica column chromatography on silica gel (Petroleum ether: EtOAc from 20/1 to 10/1) to give 2-[2-[2-[bis(*tert*-butoxycarbonyl)amino]ethoxy]ethoxy]ethyl 4-methylbenzenesulfonate (2 g, 87% yield) as a colorless oil.

**LC–MS:** MS (ES^+^): RT = 0.675 min, m/z = 304.1 [M – 200^+^];

**Spectra:**

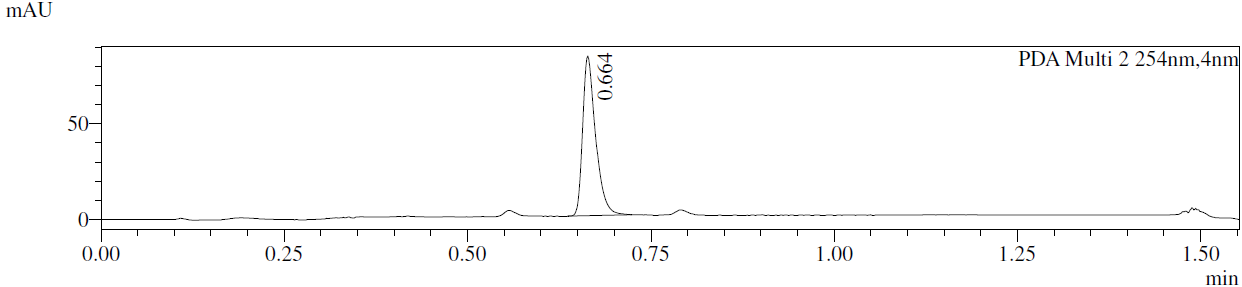

**Preparation of compound 5**

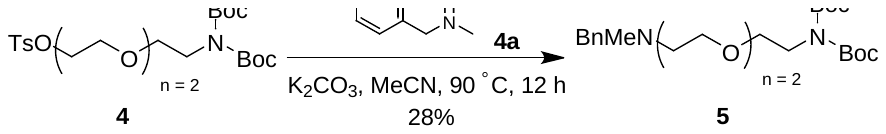

To a solution of 2-[2-[2-[bis(*tert*-butoxycarbonyl)amino]ethoxy]ethoxy]ethyl 4-methylbenzenesulfonate (2 g, 3.97 mmol, 1.0 equiv) and *N*-methyl-1-phenyl-methanamine (0.48 g, 3.97 mmol, 0.5 ml, 1.0 equiv) in MeCN (20 ml) was added K_2_CO_3_ (0.82 g, 5.96 mmol, 1.5 equiv). The mixture was stirred at 90 °C for 12 h. The mixture was filtered and concentrated under reduced pressure. The residue was purified by silica column chromatography on silica gel (Petroleum ether: EtOAc from 20/1 to 1/1) to give *tert*-butyl-*N*-[2-[2-[2-[benzyl(methyl)amino]ethoxy]ethoxy]ethyl]-*N-tert*-butoxycarbonyl-carbamate (0.5 g, 28% yield) as a colorless oil.

**LC–MS:** MS (ES^+^): RT = 0.992 min, m/z = 353.1 [M -100^+^];

**Spectra:**

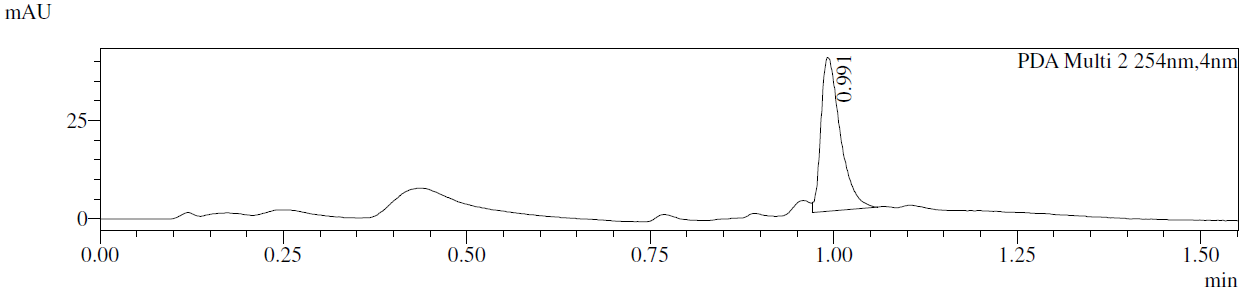

**Preparation of compound 6**

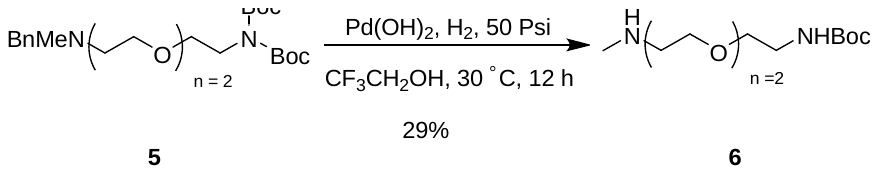

To a solution of *tert*-butyl-*N*-[2-[2-[2-[benzyl(methyl)amino]ethoxy]ethoxy]ethyl]-*N-tert*-butoxycarbonyl-carbamate (0.5 g, 1.1 mmol, 1.0 equiv) in TFE (10 ml) was added Pd(OH)_2_/C (50 mg, 10% purity) under N_2_ atmosphere. The mixture was stirred under H_2_ (50 psi) at 30 °C for 12 h. The mixture was filtered and concentrated under reduced pressure. The residue was purified by *prep*-HPLC (column: Waters Xbridge 150*25 mm* 5 µm; mobile phase: [water (10 mM NH_4_HCO_3_)–MeCN]; B%: 22%–52%, 10 min) to give *tert*-butyl *N*-[2-[2-[2-(methylamino)ethoxy]ethoxy]ethyl]carbamate (85 mg, 29% yield).

**^1^H NMR** (400 MHz, CDCl_3_): δ 3.64–3.50 (m, 8H), 3.31 (d, *J* = 4.8 Hz, 2H), 2.85–2.74 (m, 2H), 2.46 (s, 3H), 1.44 (s, 9H).

**Preparation of compound 7**

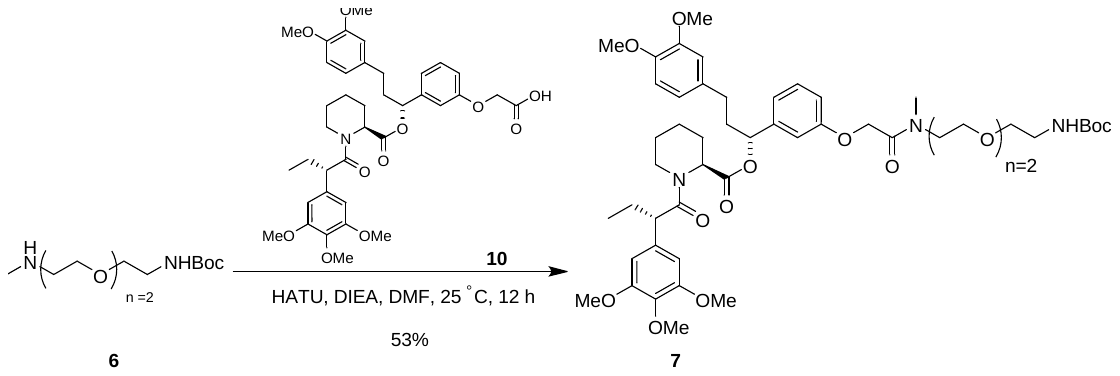

To a solution of 2-[3-[(1*R*)-3-(3,4-dimethoxyphenyl)-1-[(2*S*)-1-[(2*S*)-2-(3,4,5-trimethoxyphenyl)butanoyl]piperidine-2-

carbonyl]oxy-propyl]phenoxy]acetic acid (225 mg, 324 µmol, 1.0 equiv) in DMF (2 ml) was added tert-butyl *N*-[2-[2-[2-(methylamino)ethoxy]ethoxy]ethyl]carbamate (85 mg, 324 µmol, 1.0 equiv), HATU (148 mg, 389 µmol, 1.2 equiv) and DIEA (84 mg, 648 µmol, 2.0 equiv). The mixture was stirred at 25 °C for 12 h. The mixture was concentrated. The residue was purified by *prep*-HPLC (column: Waters Xbridge 150*25 mm* 5 µm; mobile phase: [water (10 mM NH_4_HCO_3_)–MeCN]; B%: 52%–82%, 9 min) to give [1-[3-[2-[2-[2-[2-(*tert*-butoxycarbonylamino)ethoxy]ethoxy]ethyl-methyl-amino]-2-oxo-ethoxy]phenyl]-3-(3,4-dimethoxyphenyl)propyl] (2*S*)-1-[(2*S*)-2-(3,4,5-trimethoxyphenyl)butanoyl]piperidine-2-carboxylate (160 mg, 53% yield) as a colorless oil.

**LC–MS:** MS (ES^+^): RT = 1.067 min, m/z = 955.6 [M + 18^+^];

**Spectra:**

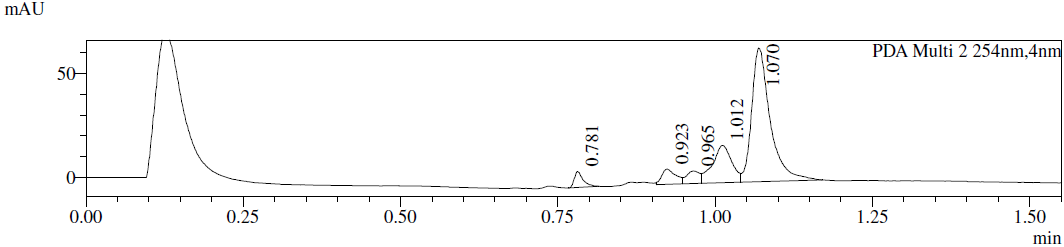

**Preparation of compound 8**

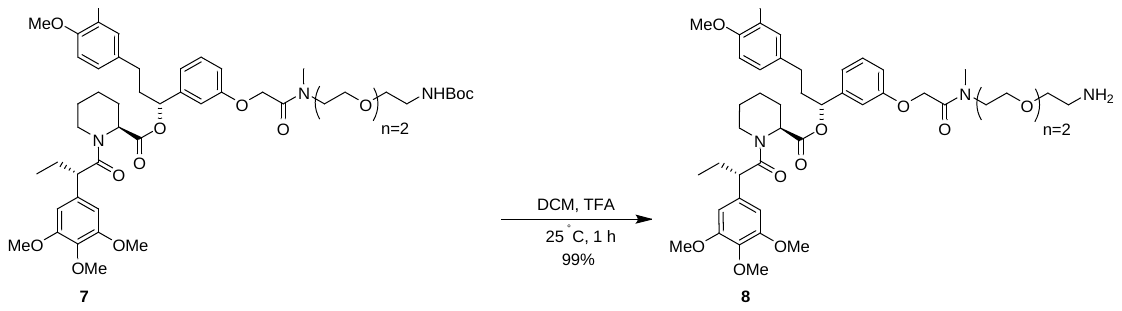

To a solution of [1-[3-[2-[2-[2-[2-(*tert*-butoxycarbonylamino)ethoxy]ethoxy]ethyl-methyl-amino]-2-oxo-ethoxy]phenyl]-3-(3,4-dimethoxyphenyl)propyl] (2*S*)-1-[(2*S*)-2-(3,4,5-trimethoxyphenyl)butanoyl]piperidine-2-carboxylate (80 mg, 85 µmol, 1.0 equiv) in CH_2_Cl_2_ (1 ml) was added TFA (0.5 ml). The mixture was stirred at 25 °C for 1 h. The mixture was concentrated to give [1-[3-[2-[2-[2-(2-aminoethoxy)ethoxy]ethyl-methyl-amino]-2-oxo-ethoxy]phenyl]-3-(3,4-dimethoxyphenyl)propyl] (2*S*)-1-[(2*S*)-2-(3,4,5-trimethoxyphenyl)butanoyl]piperidine-2-carboxylate (80 mg, 99% yield, TFA salt) as a brown oil.

**LC–MS:** MS (ES^+^): RT = 1.004 min, m/z = 838.5 [M + H^+^];

**Spectra:**

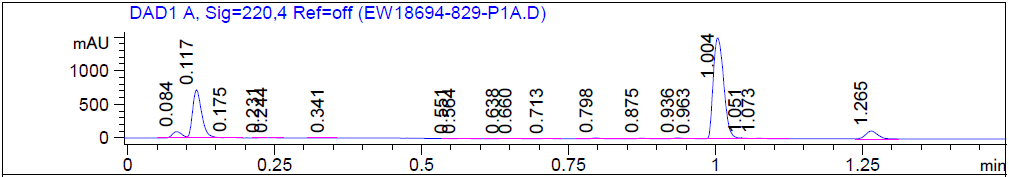

**Preparation of compound 9**

Known compound from *J. Med. Chem.* 2019, 62, 5191–-5216

**Preparation of compound 10**

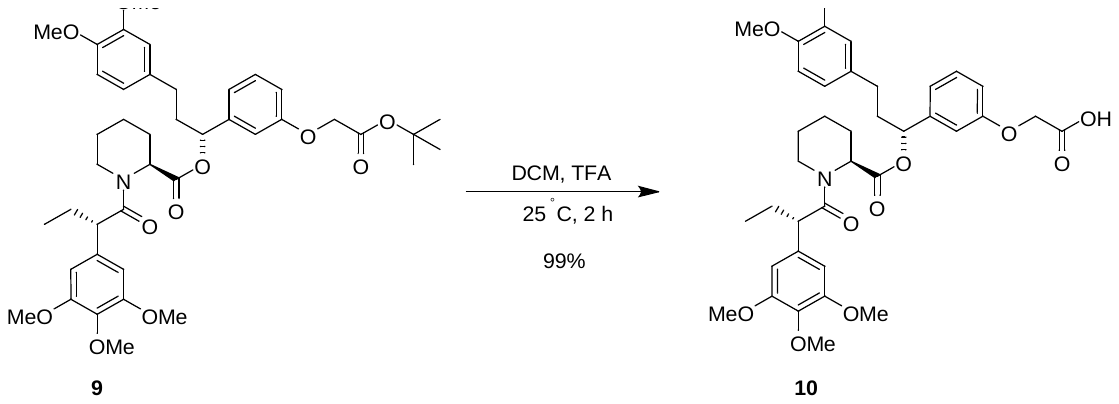

To a solution of [(1*R*)-1-[3-(2-tert-butoxy-2-oxo-ethoxy)phenyl]-3-(3,4-dimethoxyphenyl)propyl] (2*S*)-1-[(2*S*)-2-(3,4,5-trimethoxyphenyl)butanoyl]piperidine-2-carboxylate (250 mg, 333 µmol, 1.0 equiv) in CH_2_Cl_2_ (2 ml) was added TFA (1 ml). The mixture was stirred at 25 °C for 2 h. The mixture was concentrated to give 2-[3-[(1*R*)-3-(3,4-dimethoxyphenyl)-1-[(2*S*)-1-[(2*S*)-2-(3,4,5-trimethoxyphenyl)butanoyl]piperidine-2-carbonyl]oxypropyl] phenoxy]acetic acid (230 mg, 99% yield).

**LC-MS:** MS (ES^+^): RT = 0.742 min, m/z = 694.2 [M + H^+^];

**Spectra:**

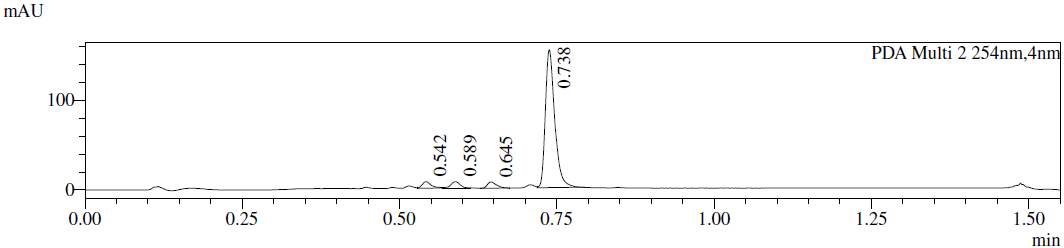

**Preparation of compound 11**

Known compound from *J. Med. Chem.* **2022**, *65*, 6573–6592

**Preparation of HLDA-221**

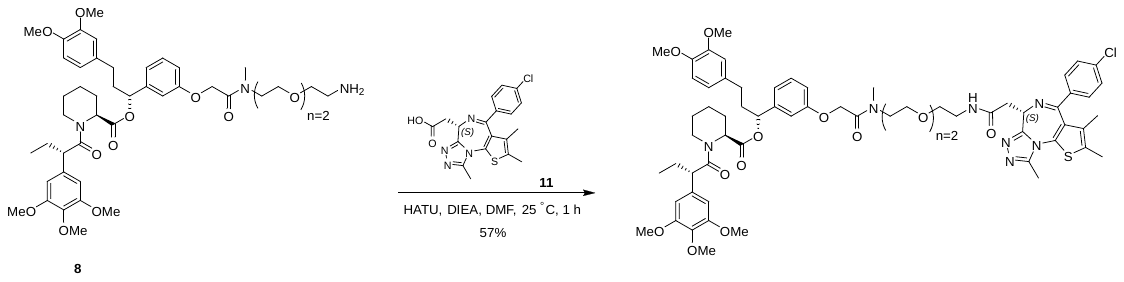

To a solution of 2-[(9*S*)-7-(4-chlorophenyl)-4,5,13-trimethyl-3-thia-1,8,11,12-tetrazatricyclo[8.3.0.02,6]trideca-2(6),4,7,10,12-pentaen-9-yl]acetic acid (34 mg, 84 µmol, 1.0 equiv) in DMF (1 ml) was added HATU (38 mg, 101 µmol, 1.2 equiv), [1-[3-[2-[2-[2-(2-aminoethoxy)ethoxy]ethyl-methyl-amino]-2-oxo-ethoxy]phenyl]-3-(3,4-dimethoxyphenyl)propyl] (2*S*)-1-[(2*S*)-2-(3,4,5-trimethoxyphenyl)butanoyl]piperidine-2-carboxylate (80 mg, 84 µmol, 1.0 equiv, TFA salt) and DIEA (33 mg, 252 µmol, 3.0 equiv). The mixture was stirred at 25 °C for 1 h. The mixture was concentrated. The residue was purified by *prep*-HPLC (column: Waters Xbridge 150*25 mm* 5 µm; mobile phase: [water (10 mM NH_4_HCO_3_)-MeCN]; B%: 53%–83%, 8 min) to give [1-[3-[2-[2-[2-[2-[[2-[(9*S*)-7-(4-chlorophenyl)-4,5,13-trimethyl-3-thia-1,8,11,12-tetrazatricyclo[8.3.0.02,6]trideca-2(6),4,7,10,12-pentaen-9-yl]acetyl]amino]ethoxy]ethoxy]ethyl-methyl-amino]-2-oxo-ethoxy]phenyl]-3-(3,4-dimethoxyphenyl)propyl](2*S*)-1-[(2*S*)-2-(3,4,5-trimethoxyphenyl)butanoyl]piperidine-2-carboxylate (58 mg, 57% yield).

**^1^H NMR** (400 MHz, DMSO-d_6_): δ 8.31–8.19 (m, 1H), 7.51–7.39 (m, 4H), 7.31–7.10 (m, 1H), 6.94–6.79 (m, 2H), 6.79–6.65 (m, 3H), 6.65–6.48 (m, 4H), 5.84–5.45 (m, 1H), 5.30–5.02 (m, 1H), 4.89–4.80 (m, 2H), 4.50 (t, *J* = 6.8 Hz, 1H), 4.00 (d, *J* = 10.4 Hz, 1H), 3.86 (t, *J* = 7.2 Hz, 1H), 3.74–3.74 (m, 1H), 3.77–3.67 (m, 8H), 3.64–3.59 (m, 2H), 3.58–3.53 (m, 9H), 3.53–3.47 (m, 4H), 3.46–3.39 (m, 3H), 3.32–3.16 (m, 6H), 3.06–3.04 (m, 1H), 2.85 (s, 1H), 2.69–2.58 (m, 3H), 2.40–2.32 (m, 3H), 2.16 (d, *J* = 14.4 Hz, 1H), 1.95–1.87 (m, 2H), 1.71–1.49 (m, 7H), 1.46–1.34 (m, 1H), 1.22–1.11 (m, 1H), 0.84–0.74 (m, 3H).

**LC-MS:** MS (ES^+^): RT = 2.435 min, m/z = 1221.6 [M + H^+^];

**Spectra:**

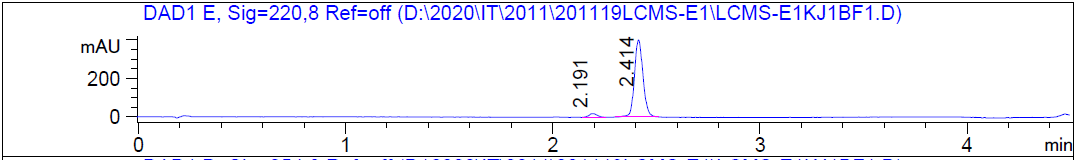

HRMS [C_64_H_78_ClN_7_O_13_S] Cal: 1220.514; Obs: 1220.5098

**The synthetic route for** **HLDA-222**

Part 1:

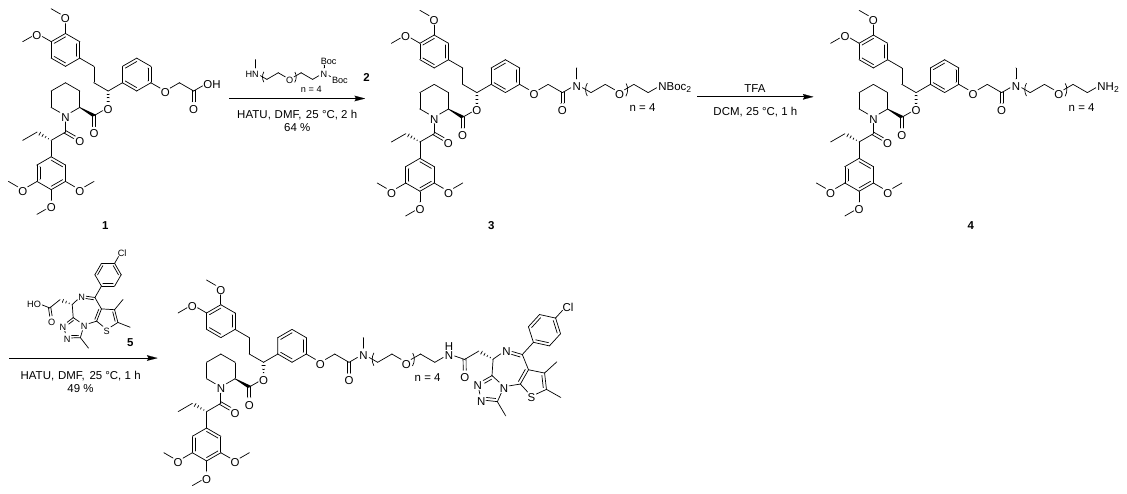

Part 2:

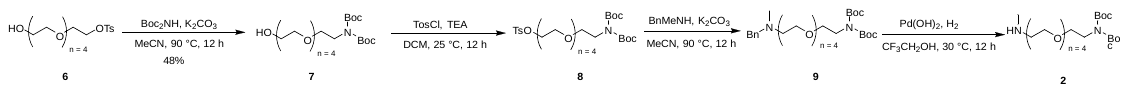

**Preparation of compound 1**

Known compound from US9096844, 2015, B2

**Preparation of 3**

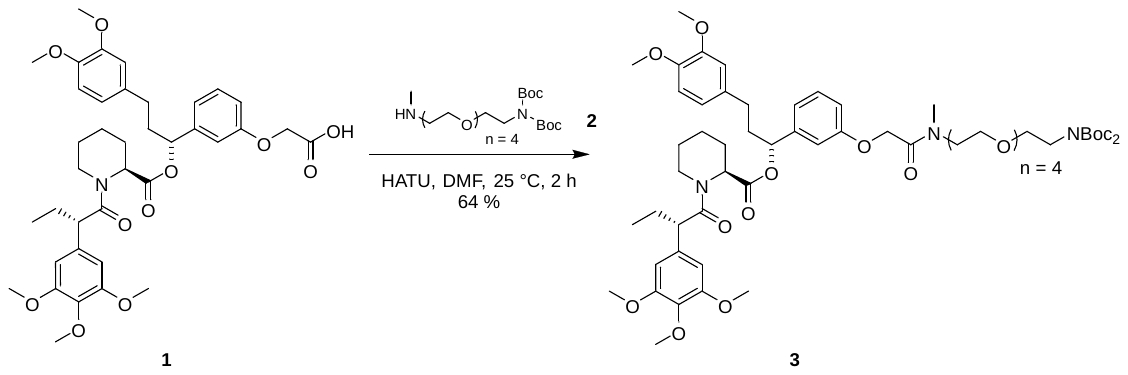

To a solution of 2-[3-[3-(3,4-dimethoxyphenyl)-1-[1-[(2*S*)-2-(3,4,5-trimethoxyphenyl)butanoyl]piperidine-2-carbonyl]oxy-propyl]phenoxy]acetic acid (185 mg, 267 µmol, 1.0 equiv) and *tert*-butyl *N-tert*-butoxycarbonyl-*N*-[2-[2-[2-[2-[2-(methylamino)ethoxy]ethoxy]ethoxy]ethoxy]ethyl]carbamate (120 mg, 267 µmol, 1.0 equiv) in DMF (2.0 ml) was added DIEA (138 mg, 1.07 mmol, 4.0 equiv) and HATU (122 mg, 320 µmol, 1.2 equiv). The mixture was stirred at 25 °C for 2 h. The mixture was purified by prep-HPLC (column: Waters Xbridge 150*25 mm* 5 µm; mobile phase: [water(10 mM NH_4_HCO_3_)–MeCN]; B%: 62%–92%, 8 min) to give [1-[3-[2-[2-[2-[2-[2-[2-[bis(*tert*-butoxycarbonyl)amino]ethoxy]ethoxy]ethoxy]ethoxy]ethyl-methyl-amino]-2-oxo-ethoxy]phenyl]-3-(3,4-dimethoxyphenyl)propyl]1-[(2*S*)-2-(3,4,5-trimethoxyphenyl)butanoyl]piperidine-2-carboxylate (200 mg, 64 % yield).

**LC–MS:** MS (ES^+^): RT = 1.153 min, m/z = 926.5 [M – 200];

**Spectra:**

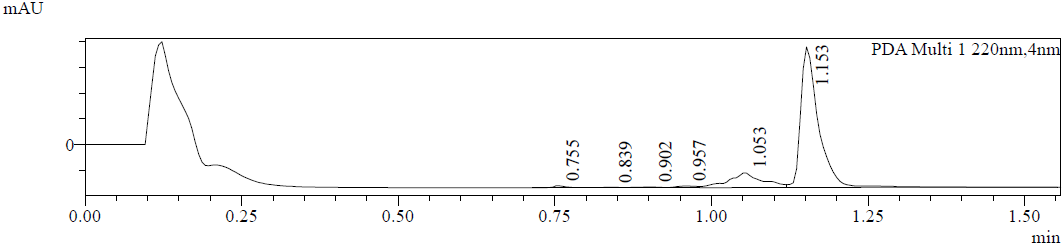

**Preparation of 4**

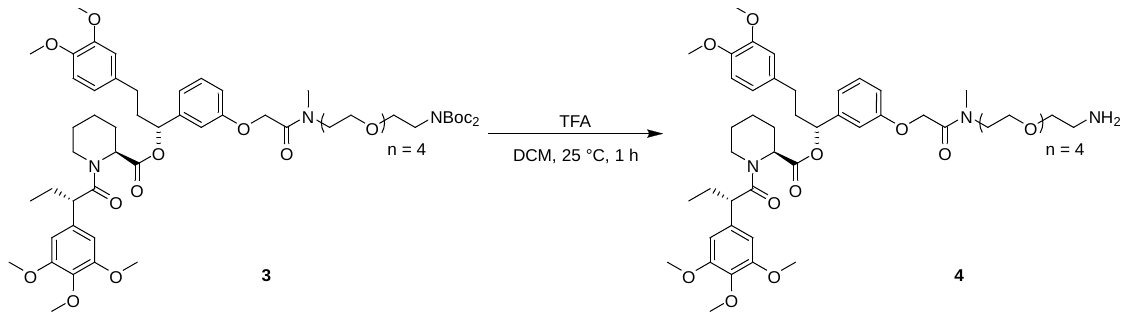

To a solution of [(1*R*)-1-[3-[2-[2-[2-[2-[2-[2-[bis(*tert*-butoxycarbonyl)amino]ethoxy]ethoxy]ethoxy]ethoxy]ethyl-methyl-amino]-2-oxo-ethoxy]phenyl]-3-(3,4-dimethoxyphenyl)propyl](2*S*)-1-[(2*S*)-2-(3,4,5-trimethoxyphenyl)butanoyl]piperidine-2-carboxylate (100 mg, 88 µmol, 1.0 equiv) in CH_2_Cl_2_ (2.0 ml) was added TFA (1.54 g, 13.5 mmol, 1.0 ml, 152.1 equiv). The mixture was stirred at 25 °C for 1 h. The mixture was concentrated to give the crude product [(1*R*)-1-[3-[2-[2-[2-[2-[2-(2-aminoethoxy)ethoxy]ethoxy]ethoxy]ethyl-methyl-amino]-2-oxo-ethoxy]phenyl]-3-(3,4-dimethoxyphenyl)propyl](2*S*)-1-[(2*S*)-2-(3,4,5-trimethoxyphenyl)butanoyl]piperidine-2-carboxylate (92 mg, crude, TFA salt).

**LC-MS:** MS (ES^+^): RT = 0.797 min, m/z = 926.4 [M + H^+^];

**Spectra:**

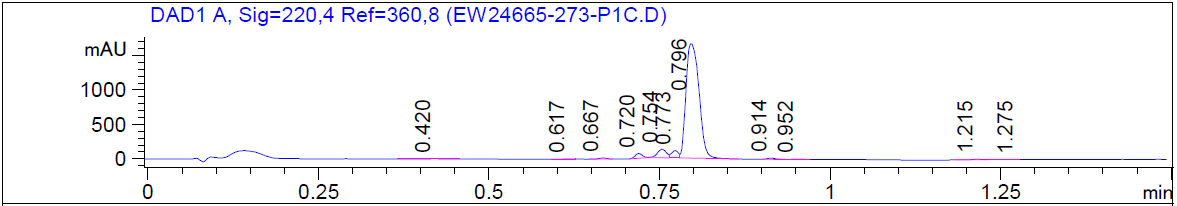

**Preparation of HLDA-222**

To a solution of [(1*R*)-1-[3-[2-[2-[2-[2-[2-(2-aminoethoxy)ethoxy]ethoxy]ethoxy]ethyl-methyl-amino]- 2-oxo-ethoxy]phenyl]-3-(3,4-dimethoxyphenyl)propyl] (2*S*)-1-[(2*S*)-2-(3,4,5-trimethoxyphenyl)butanoyl] piperidine-2-carboxylate (92 mg, 88 µmol, 1.0 equiv, TFA salt) and 2-[(9*S*)-7-(4-chlorophenyl)-4,5,13-trimethyl-3-thia-1,8,11,12-tetrazatricyclo[8.3.0.0^2,6^]trideca-2(6),4,7,10,12-pentaen-9-yl]acetic acid (35 mg, 88 µmol, 1.0 equiv) in DMF (2.0 ml) was added DIEA (57 mg, 442 µmol, 77 µl, 5.0 equiv) and HATU (40 mg, 106 µmol, 1.2 equiv).  The mixture was stirred at 25 °C for 1 h. The mixture was purified by *prep*-HPLC (column: Phenomenex Gemini-NX C18 75*30 mm*3 µm;mobile phase: [water(10mM NH_4_HCO_3_)–MeCN]; B%: 50%–80%, 8 min) to give [(1*R*)-1-[3-[2-[2-[2-[2-[2-[2-[[2-[(9*S*)-7-(4-chlorophenyl)-4,5,13-trimethyl-3-thia-1,8,11,12-tetrazatricyclo[8.3.0.0^2,6^]trideca-2(6),4,7,10,12-pentaen-9-yl]acetyl]amino]ethoxy]ethoxy]ethoxy]ethoxy]ethyl-methyl-amino]-2-oxo-ethoxy]phenyl]-3-(3,4-dimethoxyphenyl)propyl](2*S*)-1-[(2*S*)-2-(3,4,5-trimethoxyphenyl)butanoyl]piperidine-2-carboxylate (57 mg, 49 % yield).

**^1^H NMR** (400 MHz, MeOD): δ 7.49–7.36 (m, 4 H), 7.22– 7.08 (m, 1 H), 6.96– 6.65 (m, 5 H), 6.64– 6.43 (m, 3 H), 5.85– 5.52 (m, 1 H), 5.46– 5.32 (m, 1 H), 4.92–4.89 (m, 1 H), 4.84–4.77 (m, 1 H), 4.62 (dd, *J*=9.2, 5.2 Hz, 1 H), 4.54– 4.03 (m, 1 H), 3.89– 3.73 (m, 9 H), 3.71–3.51 (m, 26 H), 3.50– 3.40 (m, 3 H), 3.14– 3.10 (m, 1 H), 3.04– 2.81 (m, 2 H), 2.81– 2.61 (m, 4 H), 2.61–2.35 (m, 5 H), 2.32– 2.21 (m, 1 H), 2.09– 1.80 (m, 3 H), 1.78– 1.41 (m, 8 H), 1.30– 1.16 (m, 1 H), 0.91– 0.80 (m, 3 H).

**LC-MS:** MS (ES^+^): RT = 2.858 min, m/z = 1308.5 [M + H^+^];

**Spectra:**

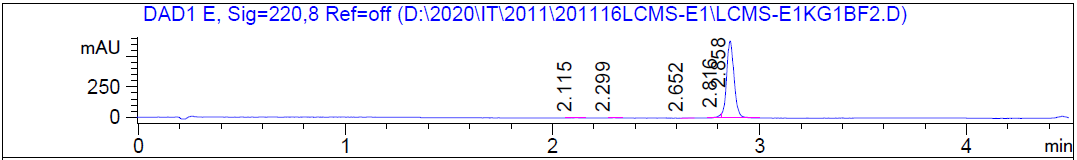

HRMS [C_68_H_86_ClN_7_O_15_S] Cal: 1308.5664; Obs: 1308.5623

**Preparation of compound 7**

To a solution of 2-[2-[2-[2-(2-hydroxyethoxy)ethoxy]ethoxy]ethoxy]ethyl 4-methylbenzenesulfonate (15 g, 38 mmol, 1.0 equiv) and *tert*-butyl *N-tert*-butoxycarbonylcarbamate (8.3 g, 38 mmol, 1.0 equiv) in MeCN (200 ml) was added K_2_CO_3_ (10 g, 76 mmol, 2.0 equiv). The mixture was stirred at 90 °C for 12 h.  The mixture was purified by *prep*-HPLC (column: Kromasil Eternity XT 250*80 mm*10 µm;mobile phase: [water(10 mM NH_4_HCO_3_)–MeCN]; B%: 25%–55%, 20 min) to give the desired product *tert*-butyl *N-tert*-butoxycarbonyl-*N*-[2-[2-[2-[2-(2-hydroxyethoxy)ethoxy]ethoxy]ethoxy]ethyl]carbamate (8.0 g, 18 mmol, 48% yield) as a yellow oil.

**LC–MS:** MS (ES^+^): RT = 0.656 min, m/z = 238.3 [M + H^+^-200];

**Preparation of compound 8**

To a solution of *tert*-butyl-*N-tert*-butoxycarbonyl-*N*-[2-[2-[2-[2-(2-hydroxyethoxy)ethoxy]ethoxy]ethoxy]ethyl]carbamate (5.5 g, 13 mmol, 1.0 equiv) in CH_2_Cl_2_ (20 ml) were added 4-methylbenzenesulfonyl chloride (3.6 g, 19 mmol, 1.5 equiv) and TEA (3.8 g, 38 mmol, 3.0 equiv). The mixture was stirred at 25 °C for 12 h. The mixture was purified by silica gel column chromatography (CH_2_Cl_2_ : MeOH = 100/1 to 30/1) to give desired product 2-[2-[2-[2-[2-[bis(*tert*-butoxycarbonyl)amino]ethoxy]ethoxy]ethoxy]ethoxy]ethyl-4-methylbenzenesulfonate (6.0 g, 10 mmol) as a yellow oil.

**LC-MS:** MS (ES^+^): RT = 1.037 min, m/z = 392.0 [M + H^+^-200];

**Spectra:**

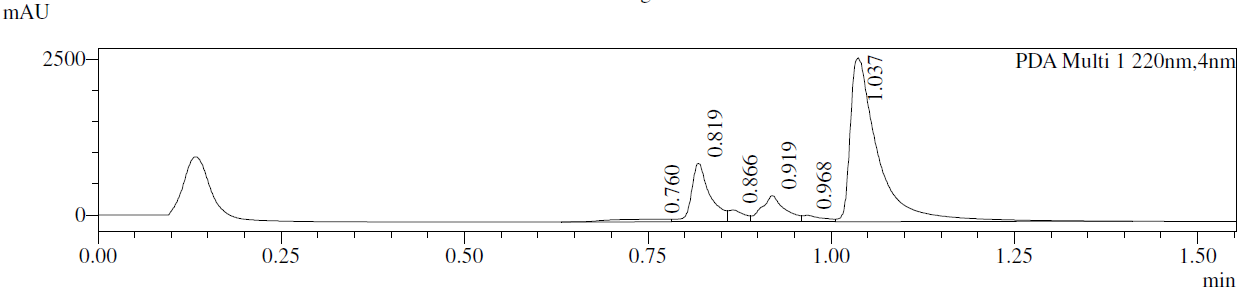

**Preparation of 9**

To a solution of 2-[2-[2-[2-[2-[bis(*tert*-butoxycarbonyl)amino]ethoxy]ethoxy]ethoxy]ethoxy]ethyl-4-methylbenzenesulfonate (9.0 g, 15 mmol, 1.0 equiv) in MeCN (100 ml) were added *N*-methyl-1-phenyl-methanamine (1.8 g, 15 mmol, 1.0 equiv), K_2_CO_3_ (2.3 g, 17 mmol, 1.1 equiv). The mixture was stirred at 90 °C for 12 h. The mixture was purified by silica column chromatography on silica gel (CH_2_Cl_2_ : MeOH = 100/1 to 30/1) to give desired product *tert*-butyl-*N*-[2-[2-[2-[2-[2-[benzyl(methyl)amino]ethoxy]ethoxy]ethoxy]ethoxy]ethyl]-*N-tert*-butoxycarbonyl-carbamate (8.0 g, 15 mmol) as a yellow oil.

**LC–MS:** MS (ES^+^): RT = 0.843 min, m/z = 541.5 [M + H^+^];

**Spectra:**

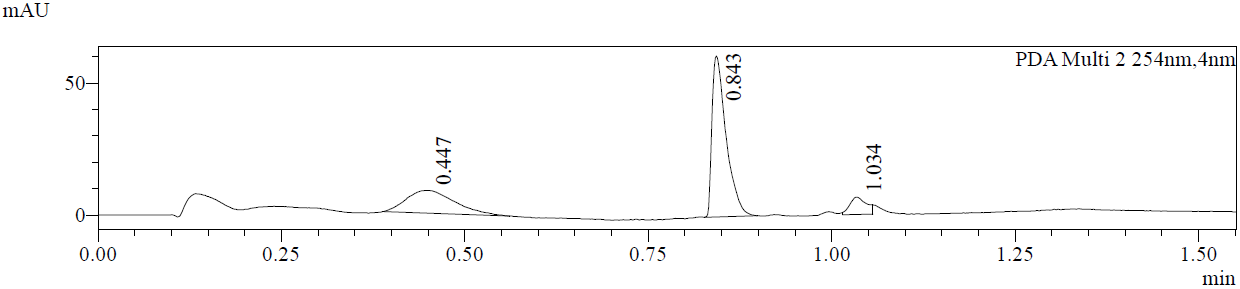

**Preparation of 2**

To a solution of *tert*-butyl-*N*-[2-[2-[2-[2-[2-[benzyl(methyl)amino]ethoxy]ethoxy]ethoxy]ethoxy]ethyl]-*N-tert*-butoxycarbonyl-carbamate (8.0 g, 15 mmol, 1.0 equiv) in CF_3_CH_2_OH (80 ml) was added Pd(OH)_2_ (0.8 g, 0.6 mmol, 10% purity). The mixture solution was purged 3 times with H_2_. The mixture was stirred at 30 °C for 12 h under H_2_ atmosphere (50 psi). The reaction mixture was filtered. The filtrate was concentrated under reduced pressure to give a desired product *tert*-butyl-*N-tert*-butoxycarbonyl-*N*-[2-[2-[2-[2-[2-(methylamino)ethoxy]ethoxy]ethoxy]ethoxy]ethyl]carbamate (6.0 g, 13 mmol) as a yellow oil.

**LC-MS:** MS (ES^+^): RT = 0.788 min, m/z = 451.5 [M + H^+^];

**The synthetic route for HLDA-223**

Part 1:

Part 2:

**Preparation of compound 3a**

See HLDA-221

**Preparation of compound 2**

To a solution of tert-butyl *N*-[2-[2-[2-[2-[2-[2-(2-aminoethoxy)ethoxy]ethoxy]ethoxy]ethoxy]ethoxy]ethyl]-*N*-methyl-carbamate (171 mg, 389 µmol, 1.2 equiv) in DMF (3 mL) was added HATU (148 mg, 389 µmol, 1.2 equiv), 2-[(9*S*)-7-(4-chlorophenyl)-4,5,13-trimethyl-3-thia-1,8,11,12-tetrazatricyclo[8.3.0.02,6]trideca-2(6),4,7,10,12-pentaen-9-yl]acetic acid (130 mg, 324 µmol, 1.0 equiv) and DIEA (126 mg, 973 µmol, 3.0 equiv). The mixture was stirred at 25 °C for 1 h. The mixture was filtered and concentrated under reduced pressure. The residue was purified by *prep*-HPLC (column: 3_Phenomenex Luna C_18_ 75*30 mm*3 µm; mobile phase: [water (0.1%TFA)–ACN]; B%: 50%–80%, 7 min) to give *tert*-butyl *N*-[2-[2-[2-[2-[2-[2-[2-[[2-[(9*S*)-7-(4-chlorophenyl)-4,5,13-trimethyl-3-thia-1,8,11,12-tetrazatricyclo[8.3.0.02,6]trideca-2(6),4,7,10,12-pentaen-9-yl]acetyl]amino]ethoxy]ethoxy]ethoxy]ethoxy]ethoxy]ethoxy]ethyl]-*N*-methyl-carbamate (130 mg, 49% yield) as a yellow oil.

**LC–MS:** MS (ES^+^): RT = 0.733 min, m/z = 821.3 [M + H^+^];

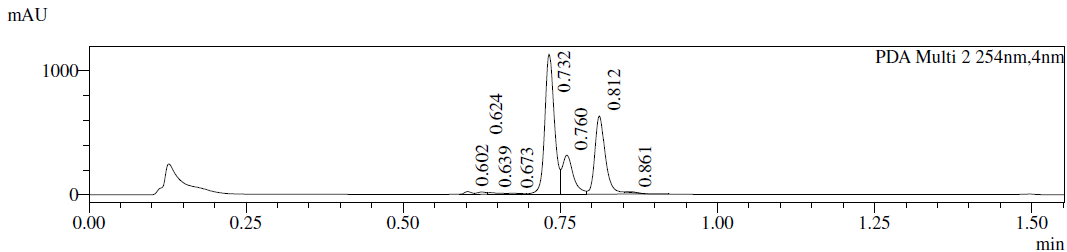

**Preparation of compound 3**

To a solution of tert-butyl *N*-[2-[2-[2-[2-[2-[2-[2-[[2-[(9*S*)-7-(4-chlorophenyl)-4,5,13-trimethyl-3-thia-1,8,11,12-tetrazatricyclo[8.3.0.02,6]trideca-2(6),4,7,10,12-pentaen-9-yl]acetyl]amino]ethoxy]ethoxy]ethoxy]ethoxy]ethoxy]ethoxy]ethyl]-*N*-methyl-carbamate (130 mg, 158 µmol, 1.0 equiv) in CH_2_Cl_2_ (1 ml) was added TFA (462 mg, 4.05 mmol, 0.3 ml, 25.6 equiv). The mixture was stirred at 25 °C for 1 h. The mixture was concentrated under reduced pressure to give 2-[(9*S*)-7-(4-chlorophenyl)-4,5,13-trimethyl-3-thia-1,8,11,12-tetrazatricyclo[8.3.0.02,6]trideca-2(6),4,7,10,12-pentaen-9-yl]-*N*-[2-[2-[2-[2-[2-[2-[2-(methylamino)ethoxy]ethoxy]ethoxy]ethoxy]ethoxy]ethoxy]ethyl]acetamide (130 mg, 98% yield, TFA salt) as a yellow oil.

**LC-MS:** MS (ES^+^): RT = 0.567 min, m/z = 721.2 [M + H^+^];

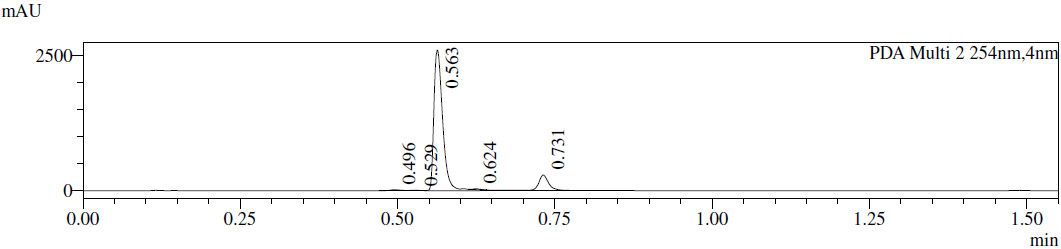

**Preparation of HLDA-223**

To a solution of 2-[3-[(1*R*)-3-(3,4-dimethoxyphenyl)-1-[(2*S*)-1-[(2*S*)-2-(3,4,5-trimethoxyphenyl)butanoyl]piperidine-2-carbonyl]oxy-propyl]phenoxy]acetic acid (108 mg, 156 µmol, 1.0 equiv) in DMF (2 ml) was added 2-[(9*S*)-7-(4-chlorophenyl)-4,5,13-trimethyl-3-thia-1,8,11,12-tetrazatricyclo[8.3.0.02,6]trideca-2(6),4,7,10,12-pentaen-9-yl]-*N*-[2-[2-[2-[2-[2-[2-[2-(methylamino)ethoxy]ethoxy]ethoxy]ethoxy]ethoxy]ethoxy]ethyl]acetamide (130 mg, 156 µmol, 1.0 equiv, TFA salt) , HATU (71 mg, 0.19 mmol, 1.2 equiv) and DIEA (60 mg, 0.47 mmol, 3.0 equiv). The mixture was stirred at 25 °C for 1 h. The mixture was filtered and concentrated. The residue was purified by *prep*-HPLC (column: 3_Phenomenex Luna C18 75*30 mm*3 µm; mobile phase: [water (0.1%TFA)–MeCN]; B%: 55%–85%, 10 min) to give [(1*R*)-1-[3-[2-[2-[2-[2-[2-[2-[2-[2-[[2-[(9*S*)-7-(4-chlorophenyl)-4,5,13-trimethyl-3-thia-1,8,11,12-

tetrazatricyclo[8.3.0.02,6]trideca-2(6),4,7,10,12-pentaen-9-yl]acetyl]amino]ethoxy]ethoxy]ethoxy]ethoxy]ethoxy]ethoxy]ethylmethyl-amino]-2-oxo-ethoxy]phenyl]-3-(3,4-dimethoxyphenyl)propyl](2*S*)-1-[(2*S*)-2-(3,4,5-trimethoxyphenyl)butanoyl]piperidine-2-carboxylate (95 mg, 43% yield,) as an off-white solid.

**^1^H NMR** (400 MHz, DMSO-d_6_): δ 8.28 (t, *J* = 5.6 Hz, 1H), 7.57–7.36 (m, 4H), 7.29–7.08 (m, 1H), 6.87–6.72 (m, 4H), 6.65–6.61 (m, 1H), 6.51 (d, *J* = 7.2 Hz, 2H), 5.87–5.44 (m, 1H), 5.33–5.04 (m, 1H), 4.87–4.79 (m, 2H), 4.50 (m, *J* = 6.4, 8.0 Hz, 1H), 4.00 (d, *J* = 13.2 Hz, 1H), 3.86 (t, *J* = 6.8 Hz, 1H), 3.74–3.68 (m, 8H), 3.63–3.42 (m, 36H), 3.33–3.17 (m, 6H), 3.03 (s, 1H), 2.84 (s, 1H), 2.59 (s, 3H), 2.40 (s, 3H), 2.20–2.01 (m, 2H), 1.94–1.87 (m, 2H), 1.68–1.51 (m, 7H), 1.37 (d, *J* = 13.2 Hz, 1H), 1.23–1.06 (m, 1H), 0.83–0.74 (m, 3H)

**LC–MS:** MS (ES^+^): RT = 2.467 min, m/z = 699.2 [1/2M + H^+^];

**Spectra:**

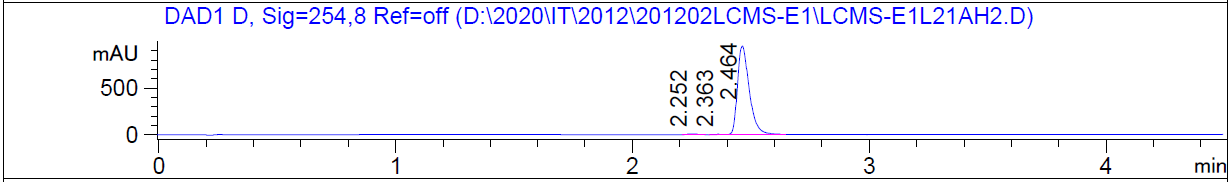

HRMS [C_72_H_94_ClN_7_O_17_S] Cal: 1396.6188; Obs: 1396.615

**Preparation of compound 5**

To a stirred solution of 2-[2-[2-[2-[2-[2-(2-hydroxyethoxy)ethoxy]ethoxy]ethoxy]ethoxy]ethoxy]ethanol (10.0 g, 30.6 mmol, 1.0 equiv) in CH_2_Cl_2_ (200 ml) at 0 °C were added Ag_2_O (10.7 g, 46.0 mmol, 1.5 equiv), NaI (5.05 g, 33.7 mmol, 1.1 equiv) and TosCl (6.13 g, 32.2 mmol, 1.1 equiv). The reaction mixture was stirred at 25 °C for 12 h. The mixture was filtered and concentrated under reduced pressure to give a residue. The residue was purified by silica column chromatography on silica gel (CH_2_Cl_2_ = 100/1 to 50/1) to give the desired product 2-[2-[2-[2-[2-[2-(2-hydroxyethoxy)ethoxy]ethoxy]ethoxy]ethoxy]ethoxy]ethyl 4-methylbenzenesulfonate (17.5 g, 36.4 mmol, 59% yield) as a colorless oil.

**LC–MS:** MS (ES^+^): RT = 0.571 min, m/z = 481.1 [M + H^+^];

**Spectra:**

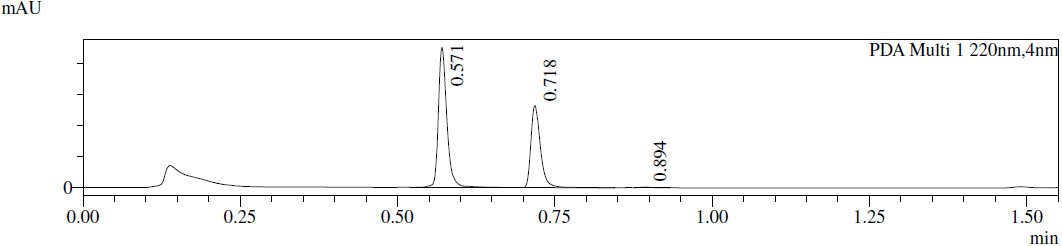

**Preparation of compound 6**

To a solution of 2-[2-[2-[2-[2-[2-(2-hydroxyethoxy)ethoxy]ethoxy]ethoxy]ethoxy]ethoxy]ethyl 4-methylbenzenesulfonate (17.5 g, 36.4 mmol, 1.0 equiv) in MeCN (150 ml) were added K_2_CO_3_ (7.55 g, 54.6 mmol, 1.5 equiv) and N-methyl-1-phenyl-methanamine (4.41 g, 36.4 mmol, 4.7 ml, 1.0 equiv). The mixture was stirred at 90 °C for 12 h. The mixture was filtered and CH_2_Cl_2_ : MeOHfrom 100/1 to 10/1) to give the crude product 2-[2-[2-[2-[2-[2-[2-[benzyl(methyl)amino]ethoxy]ethoxy]ethoxy]ethoxy]ethoxy]ethoxy]ethanol (20.0 g, crude) as a light yellow oil.

**LC–MS:** MS (ES^+^): RT = 0.842 min, m/z = 430.1 [M + H^+^];

**Spectra:**

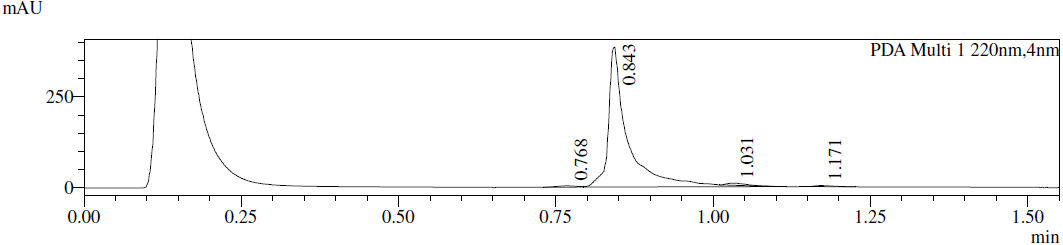

**Preparation of compound 7**

To a solution of 2-[2-[2-[2-[2-[2-[2-[benzyl(methyl)amino]ethoxy]ethoxy]ethoxy]ethoxy]ethoxy]ethoxy]ethanol (10.0 g, 23.3 mmol, 1.0 equiv) in CF_3_CH_2_OH (120 ml) were added Pd(OH)_2_/C (1.50 g, 10% purity) and Boc_2_O (10.2 g, 46.6 mmol, 10.7 ml, 2.0 equiv) under N_2_ atmosphere. The suspension was degassed under vacuum and purged with H_2_ several times. The mixture was stirred under H_2_ (50 psi) at 30 °C for 12 h. The mixture was filtered and concentrated to give the crude product 2-[2-[2-[2-[2-[2-[2-(methylamino)ethoxy]ethoxy]ethoxy]ethoxy]ethoxy]ethoxy]ethanol (15.0 g, crude) as a colorless oil.

**LC–MS:** MS (ES^+^): RT = 0.900 min, m/z = 340.0 [M + H^+^];

**Preparation of compound 8**

To a solution of 2-[2-[2-[2-[2-[2-[2-(methylamino)ethoxy]ethoxy]ethoxy]ethoxy]ethoxy]ethoxy]ethanol (15.0 g, 44.2 mmol, 1.0 equiv) in MeOH (200 ml) were added Boc_2_O (9.64 g, 44.2 mmol, 1.0 equiv). The mixture was stirred at 25 °C for 12 h. The mixture was concentrated to give a residue. The residue was purified by silica column chromatography on silica gel (CH_2_Cl_2_ = 100/1 to 20/1) to give the desired product *tert*-butyl *N*-[2-[2-[2-[2-[2-[2-(2-hydroxyethoxy)ethoxy]ethoxy]ethoxy]ethoxy]ethoxy]ethyl]-*N*-methyl-carbamate (10.0 g, 22.8 mmol, 51% yield) as a colorless oil.

**LC-MS:** MS (ES^+^): RT = 0.998 min, m/z = 457.2 [M+18^+^];

**Preparation of compound 9**

To a solution of *tert*-butyl *N*-[2-[2-[2-[2-[2-[2-(2-hydroxyethoxy)ethoxy]ethoxy]ethoxy]ethoxy]ethoxy]ethyl]-*N*-methylcarbamate (3.00 g, 6.83 mmol, 1.0 equiv) in CH_2_Cl_2_ (50 ml) were added TEA (1.38 g, 13.7 mmol, 1.9 ml, 2.0 equiv) and TosCl (1.95 g, 10.2 mmol, 1.5 equiv) at 0 °C. The mixture was stirred at 25 °C for 12 h. The mixture was concentrated to give a residue. The residue was purified by silica column chromatography on silica gel (CH_2_Cl_2_ / MeOH = 100/1 to 20/1) to give the desired product 2-[2-[2-[2-[2-[2-[2-[*tert*-butoxycarbonyl(methyl)amino]ethoxy]ethoxy]ethoxy]ethoxy]ethoxy]ethoxy]ethyl 4-methylbenzenesulfonate (3.34 g, 5.63 mmol, 82% yield) as a light yellow oil.

**Preparation of compound 10**

To a solution of 2-[2-[2-[2-[2-[2-[2-[*tert*-butoxycarbonyl(methyl)amino]ethoxy]ethoxy]ethoxy]ethoxy]ethoxy]ethoxy]ethyl 4-methylbenzenesulfonate (3.34 g, 5.63 mmol, 1.0 equiv) in DMF (20 ml) was added NaN_3_ (1.17 g, 18.0 mmol, 3.2 equiv). The mixture was stirred at 70 °C for 2 h. To the reaction mixture was added water (50 ml) and the mixture was extracted with EtOAc (3 x 50 ml). The combined organic phase was washed with brine (3 x 50 ml), dried over by Na_2_SO_4_ and filtered. The filtrate was concentrated to give the desired product *tert*-butyl *N*-[2-[2-[2-[2-[2-[2-(2-azidoethoxy)ethoxy]ethoxy]ethoxy]ethoxy]ethoxy]ethyl]-*N*-methyl-carbamate (2.50 g, 5.38 mmol, 96% yield) as a colorless oil.

**^1^H NMR** (400 MHz, CDCl_3_): δ 3.71-3.63 (m, 24H), 3.40 (d, *J* = 5.1 Hz, 4H), 2.91 (s, 3H), 1.45 (s, 9H)

**LC-MS:** MS (ES^+^): RT = 0.900 min, m/z = 465.1 [M + H^+^];

**Spectra:**

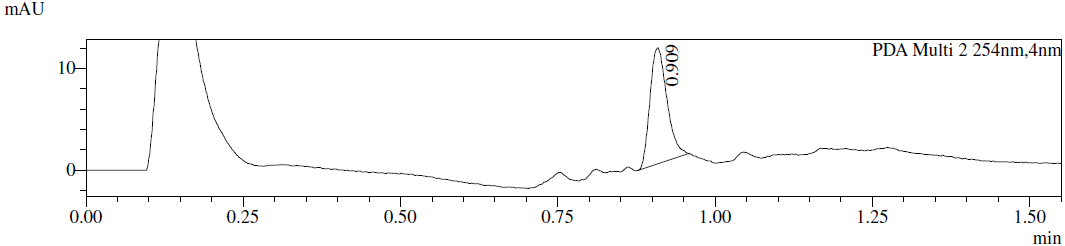

**Preparation of compound 11**

To a solution of *tert*-butyl *N*-[2-[2-[2-[2-[2-[2-(2-azidoethoxy)ethoxy]ethoxy]ethoxy]ethoxy]ethoxy]ethyl]-*N*-methyl-carbamate (1.30 g, 2.80 mmol, 1.0 equiv) in EtOH (20 ml) was added Pd/C (0.50 g, 10% purity). The mixture solution was stirred under H_2_ atmosphere (15 psi, balloon) for 0.5 h at 25 °C. The mixture was filtered and concentrated under reduced pressure to give the desired product *tert*-butyl *N*-[2-[2-[2-[2-[2-[2-(2-aminoethoxy)ethoxy]ethoxy]ethoxy]ethoxy]ethoxy]ethyl]-*N*-methyl-carbamate (1.15 g, 2.62 mmol, 94% yield) as a colorless oil.

**^1^H NMR** (400 MHz, CDCl_3_): δ 3.70-3.60 (m, 24H), 3.38 (s, 2H), 2.94–2.88 (m, 3H), 2.88–2.85 (m, 1H), 2.80 (t, *J* = 5.4 Hz, 1H), 1.45 (s, 9H)

***SERIES 8***

HLDA-231, n=2

HLDA-232, n=4

HLDA-233, n=6

**The synthetic route for HLDA-231**

**Preparation of compound 1**

known compound for *J. Med. Chem.*, **2000**, *43*, 1135

**Preparation of compound 3**

A mixture of 2-[3-[(1*R*)-3-(3,4-dimethoxyphenyl)-1-[(2*S*)-1-[(2*S*)-2-(3,4,5-trimethoxyphenyl)butanoyl]piperidine-2-carbonyl]oxy-propyl]phenoxy]acetic acid (189 mg, 253 µmol, 1.0 equiv), *tert*-butyl *N-tert*-butoxycarbonyl-*N*-[2-[2-[2-(methylamino)ethoxy]ethoxy]ethyl]carbamate (110 mg, 304 µmol, 1.0 equiv), HOBt (68.5 mg, 507 µmol, 2.0 equiv), EDCI (194 mg, 1.01 mmol, 4.0 equiv) and DIPEA (196 mg, 1.52 mmol, 6.0 equiv) in DMF (5 ml) was stirred at 20 °C for 12 h. To the reaction mixture was added 5 drops water. The residue was purified by *prep-*HPLC (column: Phenomenex Gemini-NX C18 75*30 mm*3 µm;mobile phase: [water(0.225%FA)–MeCN]; B%: 65%–95%, 7 min) to afford [(1*S*)-1-[3-[2-[2-[2-[2-[bis(*tert*-butoxycarbonyl)amino]ethoxy]ethoxy]ethyl-methyl-amino]-2-oxo-ethoxy]phenyl]-3-(3,4-dimethoxyphenyl)propyl] (2*R*)-1-[(2*R*)-2-(3,4,5-trimethoxyphenyl)butanoyl]piperidine-2-carboxylate (193 mg, 186 µmol, 73% yield) as a colorless oil.

**LC–MS:** MS (ES^+^): RT = 0.850 min, m/z = 1060.4 [M + Na^+^];

**Spectra:**

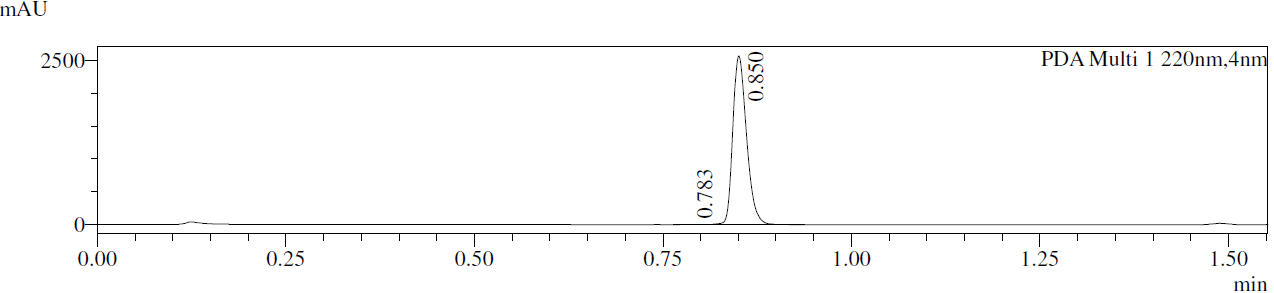

**Preparation of compound 4**

A mixture of [(1*S*)-1-[3-[2-[2-[2-[2-[bis(*tert*-butoxycarbonyl)amino]ethoxy]ethoxy]ethyl-methyl-amino]-2-oxo-ethoxy]phenyl]-3-(3,4-dimethoxyphenyl)propyl] (2*R*)-1-[(2*R*)-2-(3,4,5-trimethoxyphenyl)butanoyl]piperidine-2-carboxylate (50.0 mg, 48.2 µmol, 1.0 equiv) in TFA (1 ml) and CH_2_Cl_2_ (2 ml) was stirred at 20 °C for 1 h. The reaction mixture was concentrated under reduced pressuer to afford [(1*S*)-1-[3-[2-[2-[2-(2-aminoethoxy)ethoxy]ethyl-methyl-amino]-2-oxo-ethoxy]phenyl]-3-(3,4-dimethoxyphenyl)propyl] (2*R*)-1-[(2*R*)-2-(3,4,5-trimethoxyphenyl)butanoyl]piperidine-2-carboxylate (45.0 mg, 45.9 µmol, 95% yield, TFA salt) as a yellow oil.

**LC–MS:** MS (ES^+^): RT = 0.630 min, m/z = 838.3 [M + H^+^];

**Spectra:**

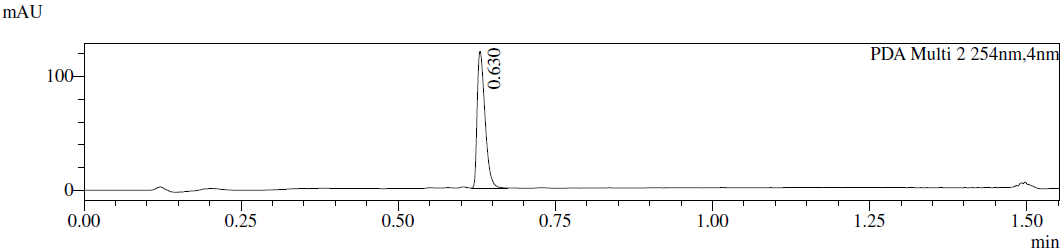

**Preparation of compound 5**

Known compound from *A.CS. Med. Chem. Lett.* **2019**, *10*, 1443–1449

**Preparation of HLDA-231**

A mixture of 4-[[(7*R*)-8-cyclopentyl-7-ethyl-5-methyl-6-oxo-7*H*-pteridin-2-yl]amino]-3-methoxy-benzoic acid (20.0 mg, 47.0 µmol, 1.0 equiv), [(1*R*)-1-[3-[2-[2-[2-(2-aminoethoxy)ethoxy]ethyl-methyl-amino]-2-oxo-ethoxy]phenyl]-3-(3,4-dimethoxyphenyl)propyl] (2*S*)-1-[(2*S*)-2-(3,4,5-trimethoxyphenyl)butanoyl]piperidine-2-carboxylate (44.8 mg, 47.0 µmol, 1.0 equiv, TFA salt), HOBt (12.7 mg, 94.0 µmol, 2.0 equiv), EDCI (27.0 mg, 141 µmol, 3.0 equiv) and DIPEA (36.5 mg, 282 µmol, 6.0 equiv) in DMF (2.5 ml) was stirred at 20 °C for 12 h. To the reaction mixture was added 2 drops of water. The mixture was purified by*prep-*HPLC (column: Phenomenex Gemini-NX C18 75*30 mm*3 µm;mobile phase: [water(0.225%FA)–MeCN]; B%: 38%–68%, 8 min) to afford [(1*R*)-1-[3-[2-[2-[2-[2-[[4-[[(7*R*)-8-cyclopentyl-7-ethyl-5-methyl-6-oxo-7*H*-pteridin-2-yl]amino]-3-methoxy-benzoyl]amino]ethoxy]ethoxy]ethyl-methyl-amino]-2-oxo-ethoxy]phenyl]-3-(3,4-dimethoxyphenyl)propyl] (2*S*)-1-[(2*S*)-2-(3,4,5-trimethoxyphenyl)butanoyl]piperidine-2-carboxylate (36.0 mg, 28.2 µmol, 60% yield) as a white solid.

**^1^H NMR** (400 MHz, DMSO-d_6_): δ 8.50–8.41 (m, 1H), 8.26–8.16 (m, 1H), 7.81 (s, 1H), 7.59–7.49 (m, 2H), 7.29–7.08 (m, 1H), 6.96–6.42 (m, 9H), 5.81–5.46 (m, 1H), 5.32–5.02 (m, 1H), 4.91–4.79 (m, 2H), 4.50–4.20 (m, 2H), 4.06–3.85 (m, 4H), 3.78–3.63 (m, 9H), 3.62–3.43 (m, 20H), 3.24 (s, 3H), 3.05–3.02 (m, 1H), 2.84 (s, 1H), 2.05–1.43 (m, 20H), 1.41–1.03 (m, 3H), 0.85–0.73 (m, 6H).

**LC–MS:** MS (ES^+^): RT = 2.076 min, m/z = 1245.7 [M + H^+^];

**Spectra:**

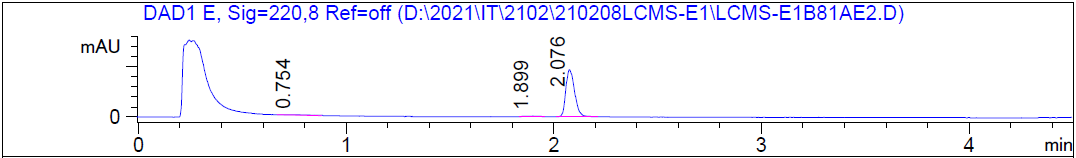

HRMS [C_67_H_88_N_8_O_15_] Cal: 1245.6442; Obs: 1245.6391

**The synthetic route for HLDA-232**

**Preparation of compound 1**

known compound for *J. Med. Chem*., **2000**, *43*, 1135

**Preparation of compound 5**

Known compound from *A.C.S. Med. Chem. Lett*. **2019**, *10*, 1443–1449

**Preparation of compound 3**

A mixture of 2-[3-[(1*R*)-3-(3,4-dimethoxyphenyl)-1-[(2*S*)-1-[(2*S*)-2-(3,4,5-trimethoxyphenyl)butanoyl]piperidine-2-carbonyl]oxy-propyl]phenoxy]acetic acid (138 mg, 199 µmol, 1.0 equiv), *tert*-butyl *N-tert*-butoxycarbonyl-*N*-[2-[2-[2-[2-[2-(methylamino)ethoxy]ethoxy]ethoxy]ethoxy]ethyl]carbamate (89.6 mg, 199 µmol, 1.0 equiv), HOBt (40.3 mg, 298 µmol, 1.5 equiv), EDCI (57.2 mg, 298 µmol, 1.5 equiv) and DIPEA (129 mg, 995 µmol, 5.0 equiv) in DMF (1.5 ml) was stirred at 20 °C for 16 h. The reaction mixture was filtered and the insoluble material was washed with DMF (0.5 ml). The combined filtrate was purified by *prep*-HPLC (column: Phenomenex luna C18 150*25 mm* 10 µm;mobile phase: [water(0.225%FA)–MeCN]; B%: 66%–96%,10 min) to afford [(1*S*)-1-[3-[2-[2-[2-[2-[2-[2-[bis(*tert*-butoxycarbonyl)amino]ethoxy]ethoxy]ethoxy]ethoxy]ethyl-methyl-amino]-2-oxo-ethoxy]phenyl]-3-(3,4-dimethoxyphenyl)propyl] (2*R*)-1-[(2*R*)-2-(3,4,5-trimethoxyphenyl)butanoyl]piperidine-2-carboxylate (150 mg, 133 µmol, 67% yield) as a yellow oil.

**^1^H NMR** (400 MHz, DMSO-d_6_): δ 7.32–7.07 (m, 1H), 6.96–6.46 (m, 8H), 5.90–5.00 (m, 2H), 4.90–4.76 (m, 2H), 4.47–3.93 (m, 1H), 3.90–3.40 (m, 37H), 3.08–2.80 (m, 3H), 2.70–2.60 (m, 1H), 2.41–2.31 (m, 1H), 2.21–2.11 (m, 1H), 2.03–1.81 (m, 3H), 1.70–1.50 (m, 4H), 1.48–1.34 (m, 18H), 1.28–0.94 (m, 2H), 0.86–0.71 (m, 3H).

**LC-MS:** MS (ES^+^): RT = 0.847 min, m/z = 926.3 [M - 200 + H^+^];

**Spectra:**

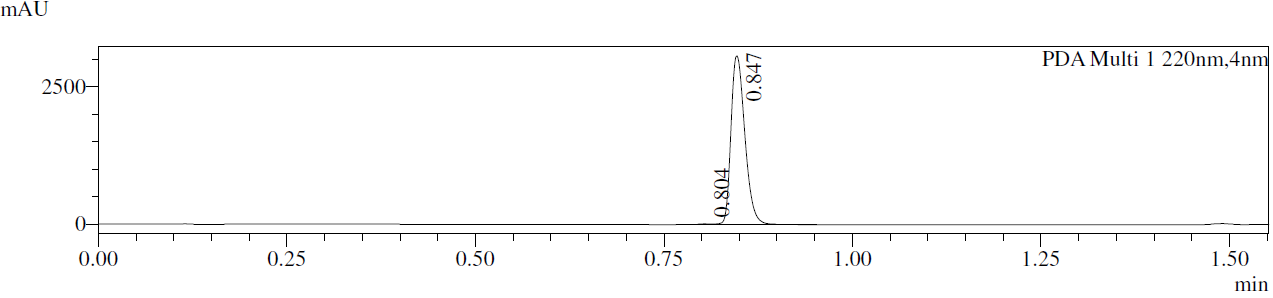

**Preparation of compound 4**

A mixture of [(1*S*)-1-[3-[2-[2-[2-[2-[2-[2-[bis(*tert*-butoxycarbonyl)amino]ethoxy]ethoxy]ethoxy]ethoxy]ethyl-methyl-amino]-2-oxo-ethoxy]phenyl]-3-(3,4-dimethoxyphenyl)propyl] (2*R*)-1-[(2*R*)-2-(3,4,5-trimethoxyphenyl)butanoyl]piperidine-2-carboxylate (150 mg, 133 µmol, 1.0 equiv) in TFA (0.5 ml) and CH_2_Cl_2_ (1 ml) was stirred at 20 °C for 0.5 h. The reaction mixture was concentrated under reduced pressure to afford [(1*S*)-1-[3-[2-[2-[2-[2-[2-(2-aminoethoxy)ethoxy]ethoxy]ethoxy]ethyl-methyl-amino]-2-oxo-ethoxy]phenyl]-3-(3,4-dimethoxyphenyl)propyl] (2*R*)-1-[(2*R*)-2-(3,4,5-trimethoxyphenyl)butanoyl]piperidine-2-carboxylate (138 mg, 133 µmol, 100% yield, TFA salt) as a yellow oil.

**LC–MS:** MS (ES^+^): RT = 0.800 min, m/z = 926.7 [M + H^+^];

**Spectra:**

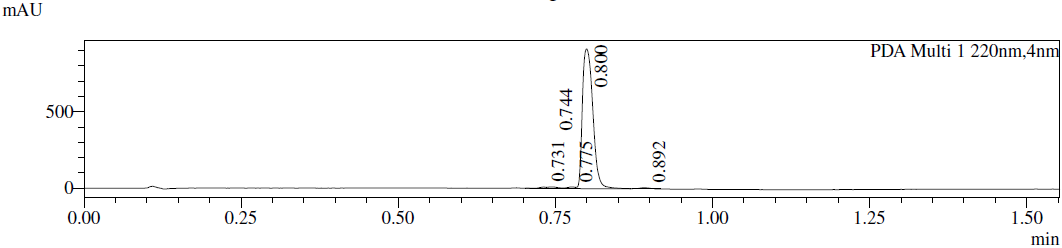

**Preparation of HLDA-232**

A mixture of 4-[[(7*R*)-8-cyclopentyl-7-ethyl-5-methyl-6-oxo-7*H*-pteridin-2-yl]amino]-3-methoxy-benzoic acid (56.0 mg, 132 µmol, 1.0 equiv), [(1*S*)-1-[3-[2-[2-[2-[2-[2-(2-aminoethoxy)ethoxy]ethoxy]ethoxy]ethyl-methyl-amino]-2-oxo-ethoxy]phenyl]-3-(3,4-dimethoxyphenyl)propyl] (2*R*)-1-[(2*R*)-2-(3,4,5-trimethoxyphenyl)butanoyl]piperidine-2-carboxylate (137 mg, 132 µmol, 1.0 equiv, TFA salt), HOBt (26.7 mg, 197 µmol, 1.5 equiv), EDCI (37.9 mg, 197 µmol, 1.5 equiv) and DIPEA (85.1 mg, 658 µmol, 5.0 equiv) in DMF (1.5 ml) was stirred at 20 °C for 16 h. The reaction mixture was filtered and the insoluble material was washed with DMF (0.5 ml). The combined filtrate was purified by *prep*-HPLC (column: Phenomenex Luna C18 150*25 mm*10 µm;mobile phase: [water(0.225%FA)–MeCN]; B%: 37%–67%, 10 min) to afford [(1*S*)-1-[3-[2-[2-[2-[2-[2-[2-[[4-[[(7*R*)-8-cyclopentyl-7-ethyl-5-methyl-6-oxo-7*H*-pteridin-2-yl]amino]-3-methoxy-benzoyl]amino]ethoxy]ethoxy]ethoxy]ethoxy]ethyl-methyl-amino]-2-oxo-ethoxy]phenyl]-3-(3,4-dimethoxyphenyl)propyl] (2*R*)-1-[(2*R*)-2-(3,4,5-trimethoxyphenyl)butanoyl]piperidine-2-carboxylate (97.0 mg, 72.7 µmol, 55% yield) as a yellow solid.

**^1^H NMR** (400 MHz, DMSO-d_6_): δ 8.50 (s, 1H), 8.14 (s, 1H), 7.81 (s, 1H), 7.57–7.48 (m, 2H), 7.31–7.09 (m, 1H), 6.95–6.71 (m, 4H), 6.69–6.48 (m, 4H), 5.81–5.02 (m, 2H), 4.89–4.79 (m, 2H), 4.38–4.20 (m, 2H), 4.05–3.82 (m, 5H), 3.76–3.54 (m, 18H), 3.53–3.36 (m, 15H), 3.22 (s, 3H), 3.04–2.81 (m, 3H), 2.68–2.58 (m, 1H), 2.47–2.31 (m, 2H), 2.20–2.11 (m, 1H), 2.00–1.77 (m, 8H), 1.75–0.95 (m, 13H), 0.84–0.72 (m, 6H).

**LC-MS:** MS (ES^+^): RT = 1.868 min, m/z = 1333.9 [M + H^+^];

**Spectra:**

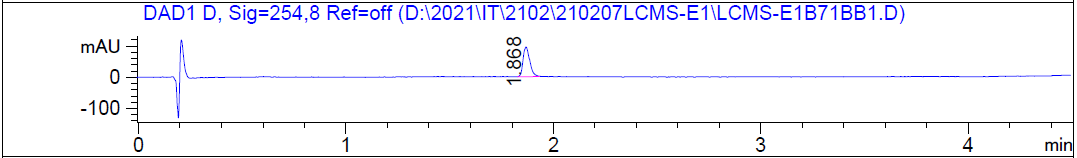

HRMS [C_71_H_96_N_8_O_17_] Cal: 1333.6966; Obs: 1333.691

**The synthetic route for HLDA-233**

**Preparation of compound 1**

known compound for *J. Med. Chem*., **2000**, *43*, 1135

**Preparation of compound 5**

Known compound from *A.C.S. Med. Chem. Lett.* **2019**, *10*, 1443–1449

**Preparation of compound 3**

A mixture of *tert*-butyl *N-tert*-butoxycarbonyl-*N*-[2-[2-[2-[2-[2-[2-[2-(methylamino)ethoxy]ethoxy]ethoxy]ethoxy]ethoxy]ethoxy]ethyl]carbamate (164 mg, 304 µmol, 1.2 equiv), 2-[3-[(1*R*)-3-(3,4-dimethoxyphenyl)-1-[(2*S*)-1-[(2*S*)-2-(3,4,5-trimethoxyphenyl)butanoyl]piperidine-2-carbonyl]oxy-propyl]phenoxy]acetic acid (189 mg, 253 µmol, 1.0 equiv), EDCI (194 mg, 1.01 mmol, 4.0 equiv), DIPEA (196 mg, 1.52 mmol, 6.0 equiv) and HOBt (68.5 mg, 507 µmol, 2.0 equiv) in DMF (5 ml) was stirred at 20 °C for 12 h. To the reaction mixture was added 5 drops of water. The residue was purified by *Prep-*HPLC (column: Phenomenex Gemini-NX C18 75*30 mm*3 µm;mobile phase: [water(0.225%FA)–MeCN]; B%: 65%–95%, 7 min) to afford [(1*S*)-1-[3-[2-[2-[2-[2-[2-[2-[2-[2-[bis(tert-butoxycarbonyl)amino]ethoxy]ethoxy]ethoxy]ethoxy]ethoxy]ethoxy]ethyl-methyl-amino]-2-oxo-ethoxy]phenyl]-3-(3,4-dimethoxyphenyl)propyl] (2*R*)-1-[(2*R*)-2-(3,4,5-trimethoxyphenyl)butanoyl]piperidine-2-carboxylate (156 mg, 128 µmol, 51% yield) as a colorless oil.

**LC-MS:** MS (ES^+^): RT = 0.847 min, m/z = 1014.4 [M – 200 + H^+^];

**Spectra:**

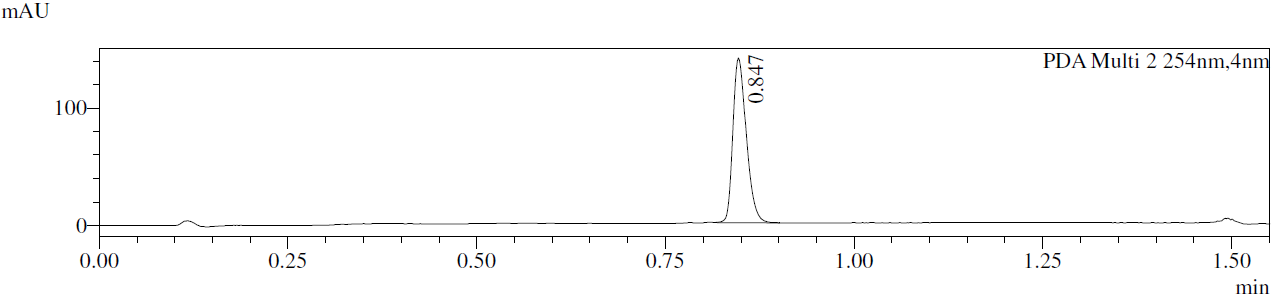

**Preparation of compound 4**

A mixture of [(1*S*)-1-[3-[2-[2-[2-[2-[2-[2-[2-[2-[bis(*tert*-butoxycarbonyl)amino]ethoxy]ethoxy]ethoxy]ethoxy]ethoxy]ethoxy]ethyl-methyl-amino]-2-oxo-ethoxy]phenyl]-3-(3,4-dimethoxyphenyl)propyl] (2*R*)-1-[(2*R*)-2-(3,4,5-trimethoxyphenyl)butanoyl]piperidine-2-carboxylate (60.0 mg, 49.4 µmol, 1.0 equiv) in TFA (1 ml) and CH_2_Cl_2_ (2 ml) was stirred at 20 °C for 1 h. The reaction mixture was concentrated under reduced pressure to afford [(1*S*)-1-[3-[2-[2-[2-[2-[2-[2-[2-(2-aminoethoxy)ethoxy]ethoxy]ethoxy]ethoxy]ethoxy]ethyl-methyl-amino]-2-oxo-ethoxy]phenyl]-3-(3,4-dimethoxyphenyl)propyl] (2*R*)-1-[(2*R*)-2-(3,4,5-trimethoxyphenyl)butanoyl]piperidine-2-carboxylate (55.0 mg, 47.8 µmol, 97% yield, TFA salt) as a yellow oil.

**LC-MS:** MS (ES^+^): RT = 0.648 min, m/z = 1014.4 [M + H^+^];

**Spectra:**

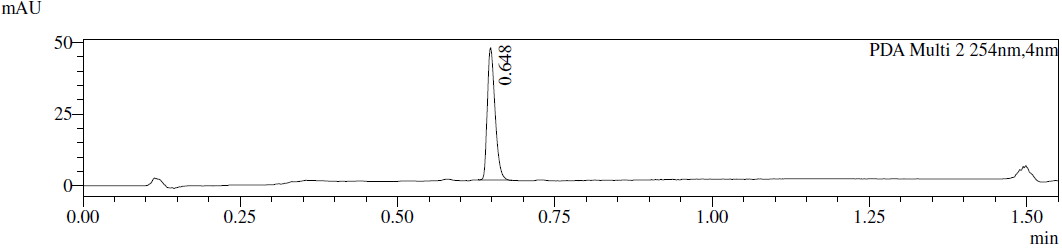

**Preparation of HLDA-233**

A mixture of 4-[[(7*R*)-8-cyclopentyl-7-ethyl-5-methyl-6-oxo-7*H*-pteridin-2-yl]amino]-3-methoxy-benzoic acid (20.0 mg, 47.0 µmol, 1.0 equiv), [(1*R*)-1-[3-[2-[2-[2-[2-[2-[2-[2-(2-aminoethoxy)ethoxy]ethoxy]ethoxy]ethoxy]ethoxy]ethyl-methyl-amino]-2-oxo-ethoxy]phenyl]-3-(3,4-dimethoxyphenyl)propyl] (2*S*)-1-[(2*S*)-2-(3,4,5-trimethoxyphenyl)butanoyl]piperidine-2-carboxylate (53.0 mg, 47.0 µmol, 1.0 equiv, TFA salt) , HOBt (12.7 mg, 94.0 µmol, 2.0 equiv) , EDCI (27.0 mg, 141 µmol, 3.0 equiv) and DIPEA (36.5 mg, 282 µmol, 6.0 equiv) in DMF (2.5 ml) was stirred at 20 °C for 12 h. To the reaction mixture was added 2 drops of water. The mixture was purified by *Prep-*HPLC (column: Phenomenex Gemini-NX C18 75*30 mm*3 µm;mobile phase: [water(0.225%FA)–MeCN]; B%: 38%–68%, 8 min) to afford [(1*R*)-1-[3-[2-[2-[2-[2-[2-[2-[2-[2-[[4-[[(7*R*)-8-cyclopentyl-7-ethyl-5-methyl-6-oxo-7*H*-pteridin-2-yl]amino]-3-methoxy-benzoyl]amino]ethoxy]ethoxy]ethoxy]ethoxy]ethoxy]ethoxy]ethyl-methyl-amino]-2-oxo-ethoxy]phenyl]-3-(3,4-dimethoxyphenyl)propyl] (2*S*)-1-[(2*S*)-2-(3,4,5-trimethoxyphenyl)butanoyl]piperidine-2-carboxylate (28.0 mg, 19.3 µmol, 41% yield, 98% purity) as a white solid.

**^1^H NMR** (400 MHz, DMSO-d_6_): δ 8.60–8.48 (m, 1H), 8.08–7.93 (m, 1H), 7.80 (s, 1H), 7.61–7.50 (m, 2H), 7.32–7.10 (m, 1H), 6.98–6.48 (m, 9H), 5.81–5.47 (m, 1H), 5.32–5.03 (m, 1H), 4.90–4.79 (m, 2H), 4.46–4.39 (m, 1H), 4.27–4.16 (m, 1H), 4.06–3.95 (m, 1H), 3.92 (s, 3H), 3.89–3.86 (m, 1H), 3.77–3.62 (m, 10H), 3.60–3.56 (m, 8H), 3.55–3.51 (m, 8H), 3.50–3.48 (m, 10H), 3.46–3.44 (m, 6H), 3.23 (s, 3H), 3.07-3.04 (m, 1H), 2.86–2.84 (m, 1H), 2.66–2.56 (m, 1H), 2.40–2.35 (m, 1H), 2.22–2.13 (m, 1H), 2.01–1.46 (m, 20H), 1.27–1.04 (m, 2H), 0.84–0.74 (m, 6H).

**LC–MS:** MS (ES^+^): RT = 2.939 min, m/z = 711.5 [M/2 + H^+^];

**Spectra:**

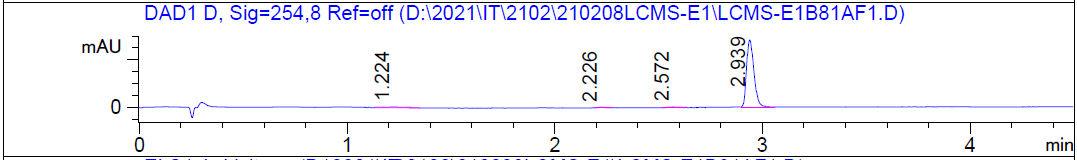

HRMS [C_75_H_104_N_8_O_19_] Cal: 1421.749; Obs: 1421.7435

***SERIES 9***

HLDA-211, n=4

HLDA-212, n=6

HLDA-213, n=8

**The synthetic route for HLDA-211**

**Preparation of compound 1**

See HLDA-222

**Preparation of compound 8**

Known compound from *Angew. Chem., Int. Ed.* **2020**¸*59*, 13865–13870

**Preparation of compound 10**

Known compound from *J. Med. Chem*., **2000**, *43*, 1135–1142

**Preparation of compound 2**

To a solution of *tert-*butyl (*tert*-butoxycarbonyl)(2-(2-hydroxyethoxy)ethyl)carbamate (2.56 g, 5.85 mmol, 1.0 equiv) in CH_2_Cl_2_ (10 ml) was added TEA (1.18 g, 11.7 mmol, 2.0 equiv) and TosCl (1.7 g, 8.8 mmol, 1.5 equiv). The mixture was stirred at 20 °C for 12 h and concentrated**.** The residue was purified by column chromatography (SiO_2_, **Petroleum ether/EtOAc = 10/1** to 1**/1**) to give 2-(2-(bis(*tert*-butoxycarbonyl)amino)ethoxy)ethyl 4-methylbenzenesulfonate (2.82 g, 81% yield) was obtained as a colorless oil.

**^1^H NMR** (400 MHz, CD_3_OD): δ 7.81 (d, *J* = 8.3 Hz, 2H), 7.35 (d, *J* = 8.1 Hz, 2H), 4.19– 4.14 (m, 2H), 3.82– 3.76 (m, 2H), 3.72– 3.66 (m, 2H), 3.64– 3.57 (m, 14H), 2.46 (s, 3H), 1.51 (s, 18H)

**LC–MS:** MS (ES^+^): RT = 0.995 min, m/z = 609.2 [M + H_3_O^+^]

**Spectra:**

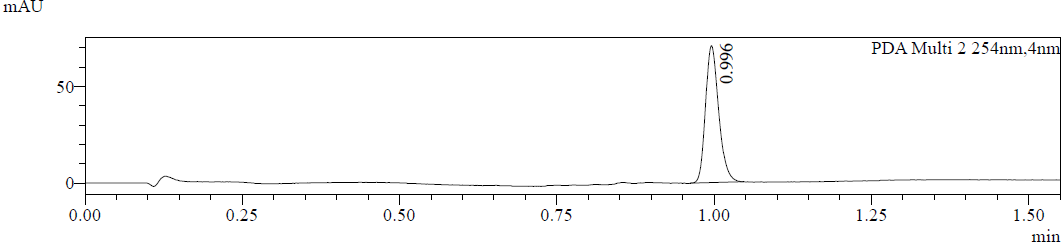

**Preparation of compound 3**

To a solution of 2-(2-(bis(*tert*-butoxycarbonyl)amino)ethoxy)ethyl 4-methylbenzenesulfonate (2.82 g, 4.77 mmol, 1.0 equiv) and *N*-methyl-1-phenyl-methanamine (866 mg, 7.15 mmol, 1.5 equiv) in MeCN (30 ml) was added K_2_CO_3_ (1.32 g, 9.53 mmol, 2.0 equiv). The mixture was stirred at 90 °C for 16 h. The mixture was filtered and concentrated. The residue was purified by column chromatography (SiO_2_, **Petroleum ether/EtOAc=0/1** to **EtOAc/MeOH = 10/1**) to give *tert*-butyl (2-(2-(benzyl(methyl)amino)ethoxy)ethyl)(tert-butoxycarbonyl)carbamate (2.4 g, 93% yield) was obtained as a colorless oil.

**^1^HNMR** (400 MHz, CD_3_OD): δ 7.43– 7.27 (m, 5H), 3.83– 3.41 (m, 20H), 2.79–2.54 (m, 2H), 2.53– 2.15 (m, 3H), 1.55–1.46 (m, 18H)

**LC–MS:** MS (ES^+^): RT = 0.864 min, m/z = 541.6 [M + H^+^]

**Spectra:**

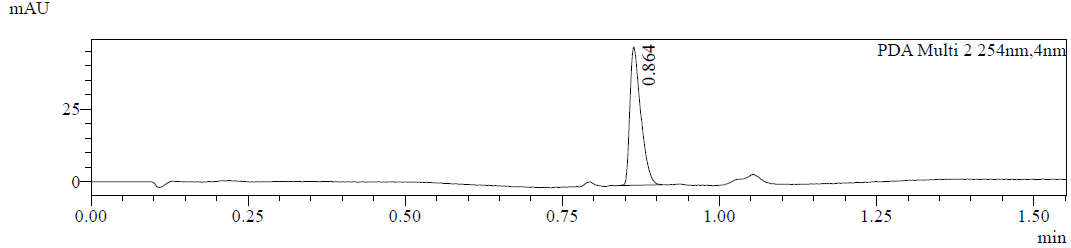

**Preparation of compound 4**

To a solution of *tert*-butyl (2-(2-(benzyl(methyl)amino)ethoxy)ethyl)(*tert*-butoxycarbonyl)carbamate (2.40 g, 4.44 mmol, 1.0 equiv) in CH_2_Cl_2_ (20 ml) was added TFA (6 ml). The mixture was stirred at 20 °C for 1 h. The mixture was concentrated to give 2-(2-aminoethoxy)-*N*-benzyl-*N*-methylethan-1-amine (2.0 g, 4.40 mmol, 99% yield, TFA salt) as a yellow oil in the next step without further purification.

**Preparation of compound 6**

To a solution of 2-(2-aminoethoxy)-*N*-benzyl-*N*-methylethan-1-amine (2.0 g, 4.4 mmol, 1.0 equiv, TFA salt) in CH_2_Cl_2_ (20 ml) was added TEA (1.11 g, 11 mmol, 2.5 equiv) and 4-nitrobenzenesulfonyl chloride (1.17 g, 5.28 mmol, 1.2 equiv). The mixture was stirred at 20 °C for 2 h. The reaction mixture was concentrated under reduced pressure. The residue was purified by *prep*-HPLC (column: Waters Xbridge C18 150*50 mm*10 µm; mobile phase: [water (10mM NH_4_HCO_3_)–MeCN]; B%: 37%–67%, 11 min) to give *N*-(2-(2-(benzyl(methyl)amino)ethoxy)ethyl)-4-nitrobenzenesulfonamide (538 mg, 23% yield) as a yellow oil.

**LC–MS:** MS (ES^+^): RT = 0.806 min, m/z = 526.4 [M + H^+^]

**Spectra:**

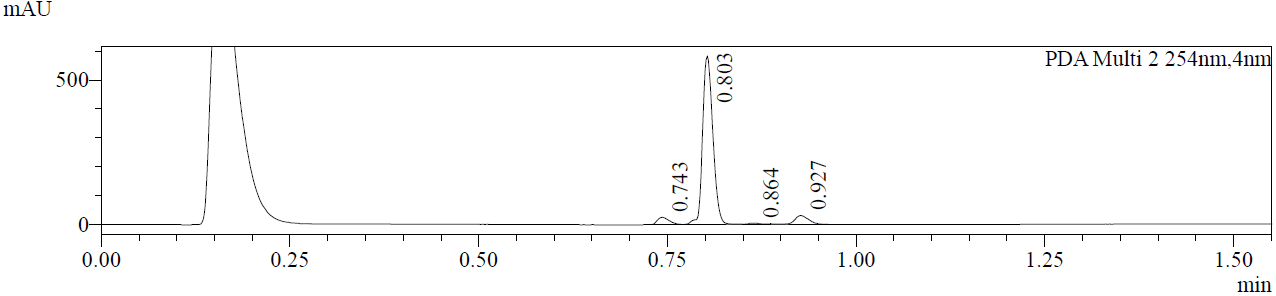

**Preparation of compound 7**

To a solution of *N*-(2-(2-(benzyl(methyl)amino)ethoxy)ethyl)-4-nitrobenzenesulfonamide (538 mg, 1.02 mmol, 1.0 equiv) in THF (10 ml) was added Pd/C (80 mg, 10% purity) under N_2_ atmosphere. The suspension was degassed and purged 3 times with H_2_. The mixture was stirred under H_2_ atmosphere (**15** psi) at 20 °C for 20 h. The mixture was filtered and concentrated under reduced pressure to afford 4-amino-*N*-(2-(2-(methylamino)ethoxy)ethyl)benzenesulfonamide (370 mg, 89% yield) as a colorless oil. The compound was used in the next step without further purification.

**LC–MS:** MS (ES^+^): RT = 0.436 min, m/z = 406.0 [M + H^+^]

**Spectra:**

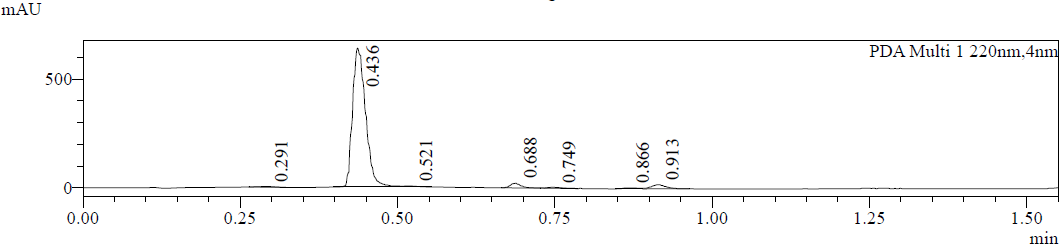

**Preparation of compound 9**

To a solution of 4-amino-*N*-(2-(2-(methylamino)ethoxy)ethyl)benzenesulfonamide (370 mg, 912 µmol, 1.0 equiv) and 2-((5-bromo-2-chloropyrimidin-4-yl)amino)-6-fluorobenzamide (378 mg, 1.09 mmol, 1.2 equiv) in NMP (2 ml) was added HCl (12 M, 380 µl, 5.0 equiv). The mixture was stirred at 95 °C for 12 h and concentrated under reduced presssure. The residue was purified by *prep*-HPLC (column: 3_Phenomenex Luna C18 75*30 mm*3 µm; mobile phase: [water (0.05%HCl)–MeCN]; B%: 20%–40%, 6.5 min) to afford 2-((5-bromo-2-((4-(*N*-(2-(2-(methylamino)ethoxy)ethyl)sulfamoyl)phenyl)amino)pyrimidin-4-yl)amino)-6-fluorobenzamide (200 mg, 31% yield, HCl salt) as a white solid.

**LC-MS:** MS (ES^+^): RT = 0.792 min, m/z = 716.0 [M + H^+^]

**Spectra:**

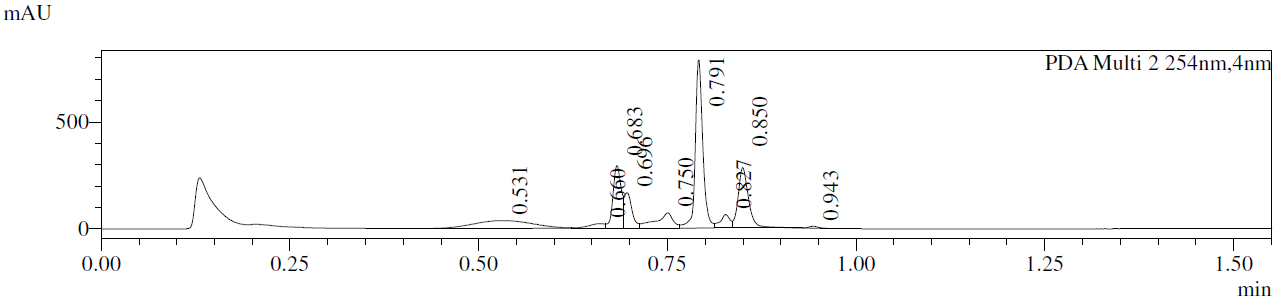

**Preparation of HLDA-211**

To a solution of 2-((5-bromo-2-((4-(*N*-(2-(2-(methylamino)ethoxy)ethyl)sulfamoyl)phenyl)amino)pyrimidin-4-yl)amino)-6-fluorobenzamide (70 mg, 93 µmol, 1.0 equiv, HCl salt) and 2-(3-((*R*)-3-(3,4-dimethoxyphenyl)-1-(((*S*)-1-((*S*)-2-(3,4,5-trimethoxyphenyl)butanoyl)piperidine-2-carbonyl)oxy)propyl)phenoxy)acetic acid (71 mg, 0.10 mmol, 1.1 equiv) in DMF (1 ml) was added HATU (53 mg, 0.14 mmol, 1.5 equiv) and DIEA (36 mg, 0.28 mmol, 3.0 equiv). The mixture was stirred at 20 °C for 0.5 h and concentrated. The residue was purified by *prep*-HPLC (column: Waters Xbridge 150*25 mm*5 µm; mobile phase: [water (10 mM NH_4_HCO_3_)–MeCN]; B%: 53%–83%, 8 min) to afford (*S*)-(*R*)-1-(3-((17-(4-((5-bromo-4-((2-carbamoyl-3-fluorophenyl)amino)pyrimidin-2-yl)amino)phenylsulfonamido)-3-methyl-2-oxo-6,9,12,15-tetraoxa-3-azaheptadecyl)oxy)phenyl)-3-(3,4-dimethoxyphenyl)propyl 1-((*S*)-2-(3,4,5-trimethoxyphenyl)butanoyl)piperidine-2-carboxylate (28 mg, 21% yield) as off-white solid.

**^1^H NMR** (400 MHz, CD_3_OD):δ 8.35 (d, *J* = 8.6 Hz, 1H), 8.25 (s, 1H), 7.82 (d, *J* = 8.9 Hz, 2H), 7.70 (d, *J* = 8.8 Hz, 2H), 7.52– 7.43 (m, 1H), 7.16– 7.09 (m, 1H), 7.01– 6.94 (m, 1H), 6.85–6.67 (m, 4H), 6.67– 6.61 (m, 1H), 6.60– 6.54 (m, 2H), 6.52– 6.43 (m, 1H), 5.60– 5.48 (m, 1H), 5.38 (m, 1H), 4.94– 4.89 (m, 2H), 4.84– 4.77 (m, 1H), 4.13– 4.01 (m, 1H), 3.89– 3.83 (m, 1H), 3.81 (s, 1H), 3.80– 3.75 (m, 6H), 3.74 (s, 1H), 3.69– 3.62 (m, 8H), 3.61– 3.39 (m, 16H), 3.15– 3.10 (m, 1H), 3.05– 2.98 (m, 2H), 2.95 (s, 1H), 2.78– 2.67 (m, 1H), 2.64– 2.33 (m, 2H), 2.31– 2.19 (m, 1H), 2.14– 1.39 (m, 8H), 1.37–1.01 (m, 2H), 0.93– 0.78 (m, 3H)

**LC–MS:** MS (ES^+^): RT = 2.723 min, m/z = 1391.4 [M + H^+^];

**Spectra:**

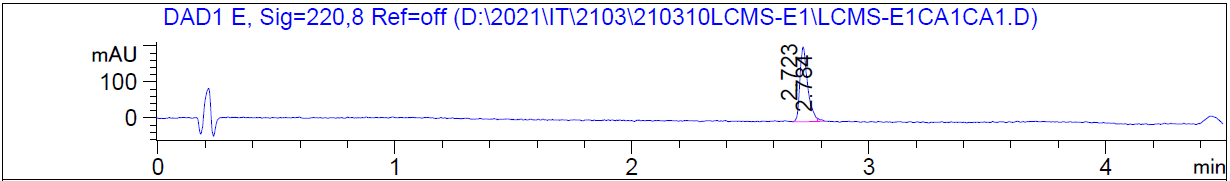

HRMS [C_66_H_82_BrFN_8_O_17_S] Cal: 1391.4739; Obs: 1391.4686

**The synthetic route for HLDA-212**

Part 1.

Part 2.

**Preparation of compound 7**

Known compound from *Angew. Chem., Int. Ed.* **2020**, *59*, 13865–13870.

**Preparation of compound 9**

Known compound from *J. Med. Chem*., **2000**, *43*, 1135 – 1142

**Preparation of compound 2**

To a solution of *tert-*butyl (2-(2-(2-bromoethoxy)ethoxy)ethyl)(methyl)carbamate (1.0 g, 1.99 mmol, 1.0 equiv) in DMF (10 ml) was added NaN_3_ (330 mg, 5.08 mmol, 2.55 equiv). The mixture was stirred at 70 °C for 12 h. The reaction mixture was poured into water (50 ml) and extracted with EtOAc (2 x 30 ml). The organic layers were washed with water (3 x 20 ml), dried over anhydrous Na_2_SO_4_, filtered, and concentrated under reduced pressure to afford *tert*-butyl (2-(2-(2-azidoethoxy)ethoxy)ethyl)(methyl)carbamate (800 mg, 1.72 mmol) was obtained as a yellow oil.

**LC–MS:** MS (ES^+^): RT = 0.712 min, m/z = 482.2 [M + 18] ^+^

**Spectra:**

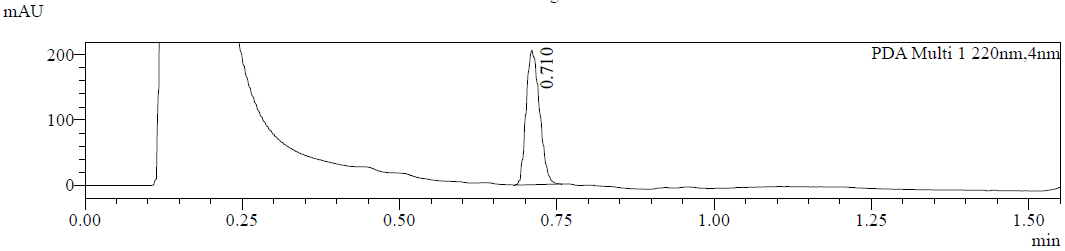

**Preparation of compound 3**

To a solution of *tert*-butyl (2-(2-(2-azidoethoxy)ethoxy)ethyl)(methyl)carbamate (800 mg, 1.72 mmol, 1.0 equiv) in THF (20 ml) was added Pd/C (0.2 g, 10% purity), and then it was degassed and purged with H_2_. The reaction mixture was stirred at 25 °C for 12 h under 15 psi pressure. After filtration, the filtrate was concentrated to afford *tert*-butyl (2-(2-(2-aminoethoxy)ethoxy)ethyl)(methyl)carbamate (800 mg, crude) as a colorless oil, and used for the next step directly without further purification.

**^1^H NMR:** (400 MHz, CDCl_3_) δ 3.77– 3.52 (m, 26H), 3.44– 3.35 (m, 2H), 2.91 (s, 3H), 1.46 (s, 9H)

**Preparation of compound 5**

To a solution of *tert*-butyl (2-(2-(2-aminoethoxy)ethoxy)ethyl)(methyl)carbamate (800 mg, 1.82 mmol, 1.0 equiv) in CH_2_Cl_2_ (10 ml) was added 4-nitrobenzenesulfonyl chloride (606 mg, 2.74 mmol, 1.5 equiv) and DIEA (1.18 g, 9.12 mmol, 1.6 ml, 5.0 equiv).The mixture was stirred at 20 °C for 1 h. The reaction mixture was quenched by 5 ml MeOH, and then it was concentrated under reduced pressure to afford crude product. The residue was purified by *prep*-HPLC (column: Waters Xbridge C18 150*50 mm*10 µm; mobile phase: [water(10 mM NH_4_HCO_3_)–MeCN]; B%: 38%–68%, 11 min) to afford *tert*-butyl methyl(2-(2-(2-((4-nitrophenyl)sulfonamido)ethoxy)ethoxy)ethyl)carbamate (240 mg, 384 µmol, 21% yield) as a yellow oil.

**^1^H NMR:** (400 MHz, CDCl_3_) δ 8.36 (d, *J* = 8.8 Hz, 2H), 8.10 (d, *J* = 8.8 Hz, 2H), 6.51– 6.03 (m, 1H), 3.71– 3.51 (m, 24H), 3.45– 3.33 (m, 2H), 3.20 (brt, *J* = 4.6 Hz, 2H), 2.91 (s, 3H), 1.46 (s, 9H)

**Preparation of compound 6**

To a solution of *tert*-butyl methyl(2-(2-(2-((4-nitrophenyl)sulfonamido)ethoxy)ethoxy)ethyl)carbamate (240 mg, 384 µmol, 1.0 equiv) in TFE (10 ml) was added Pd/C (50 mg, 10% purity), and then it was degassed and purged with H_2_. The resulting solution was stirred at 20 °C for 12 h under 15 psi pressure. The reaction mixture was filtered and the filtrate was concentrated to afford *tert*-butyl (2-(2-(2-((4-aminophenyl)sulfonamido)ethoxy)ethoxy)ethyl)(methyl)carbamate (220 mg, 370 µmol) as a colorless oil and used for the next step directly.

**LC–MS:** MS (ES^+^): RT = 0.867 min, m/z = 494.3 [M + Na] ^+^

**Spectra:**

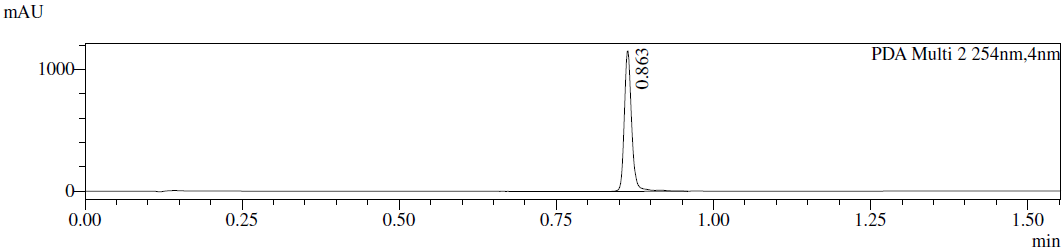

**Preparation of compound 8**

To a solution of *tert*-butyl (2-(2-(2-((4-aminophenyl)sulfonamido)ethoxy)ethoxy)ethyl)(methyl)carbamate (210 mg, 353 µmol, 1.0 equiv) and 2-((5-bromo-2-chloropyrimidin-4-yl)amino)-6-fluorobenzamide (147 mg, 425 µmol, 1.2 equi*v*) in *i*PrOH (4 ml) was added HCl (12 M, 29.47 µl, 1.0 equiv).The mixture was stirred at 95 °C for 12 h. The reaction mixture was concentrated under reduced pressure to afford crude product. The residue was purified by *prep*-HPLC (column: Phenomenex luna C18 150*40 mm*15 µm; mobile phase: [water (0.1%TFA)–MeCN]; B%: 15%–45%, 11 min) to afford 2-((5-bromo-2-((4-(*N*-(2-(2-(2-(methylamino)ethoxy)ethoxy)ethyl)sulfamoyl)phenyl)amino)pyrimidin-4-yl)amino)-6-fluorobenzamide (200 mg, 218 µmol, 61% yield, TFA salt) was obtained as a yellow gum.

**LC–MS:** MS (ES^+^): RT = 0.800 min, m/z = 804.1 [M + 2] ^+^

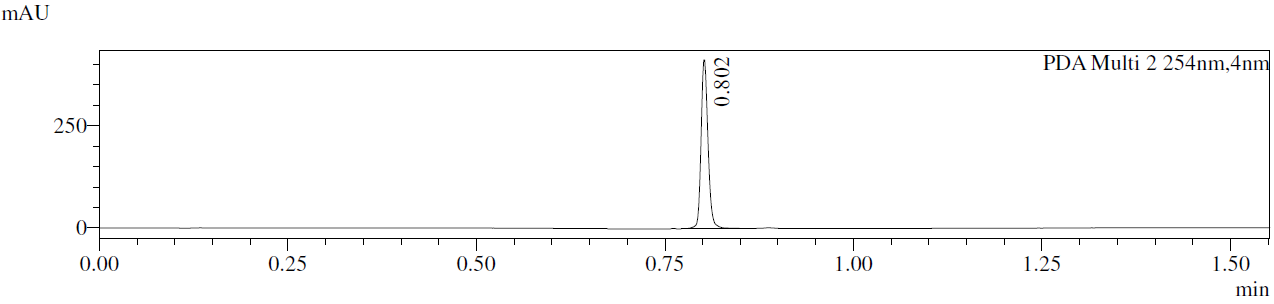

**^1^H NMR:** (400 MHz, CD_3_OD) δ 10.14 (s, 1H), 9.96 (s, 1H), 8.52 (brs, 2H), 8.39 (s, 1H), 8.24 (brd, *J* = 8.3 Hz, 1H), 8.14 (brd, *J* = 17.9 Hz, 2H), 7.85 (d, *J* = 8.8 Hz, 2H), 7.64 (d, *J* = 8.8 Hz, 2H), 7.56– 7.46 (m, 2H), 7.09 (t, *J* = 9.2 Hz, 1H), 6.88– 5.83 (m, 2H), 3.69– 3.61 (m, 2H), 3.48 (brd, *J* = 3.5 Hz, 15H), 3.46– 3.41 (m, *J* = 3.1, 5.1 Hz, 4H), 3.38 (t, *J* = 5.9 Hz, 2H), 3.14– 3.04 (m, 2H), 2.92– 2.82 (m, *J* = 5.8 Hz, 2H), 2.57 (t, *J* = 5.4 Hz, 3H)

**Preparation of HLDA-212**

To a solution of 2-(3-((*R*)-3-(3,4-dimethoxyphenyl)-1-(((*S*)-1-((*R*)-2-(3,4,5-trimethoxyphenyl)butanoyl)piperidine-2-carbonyl)oxy)propyl)phenoxy)acetic acid (80 mg, 87 µmol, 1.0 equiv, TFA salt) and 2-((5-bromo-2-((4-(*N*-(2-(2-(2-(methylamino)ethoxy)ethoxy)ethyl)sulfamoyl)phenyl)amino)pyrimidin-4-yl)amino)-6-fluorobenzamide (67 mg, 96 µmol, 1.1 equiv) in DMF (1 ml) was added HATU (49 mg, 130 µmol, 1.5 equiv) and DIEA (59 mg, 459 µmol, 80 µl, 5.2 equiv).The mixture was stirred at 20 °C for 1 h. The reaction mixture was quenched by (0.1 ml) water. The residue was purified by *prep*-HPLC (column: Phenomenex Gemini-NX C18 75*30 mm*3 µm; mobile phase: [water(10mM NH_4_HCO_3_)–MeCN]; B%: 50%–80%, 8 min) to afford (*S*)-(*R*)-1-(3-((23-(4-((5-bromo-4-((2-carbamoyl-3-fluorophenyl)amino)pyrimidin-2-yl)amino)phenylsulfonamido)-3-methyl-2-oxo-6,9,12,15,18,21-hexaoxa-3-azatricosyl)oxy)phenyl)-3-(3,4-dimethoxyphenyl)propyl 1-((*R*)-2-(3,4,5-trimethoxyphenyl)butanoyl)piperidine-2-carboxylate (42 mg, 28 µmol, 32% yield) as a white solid.

**LC–MS:** MS (ES^+^): RT = 3.355 min, m/z = 740.4 [M/2 + H^+^]

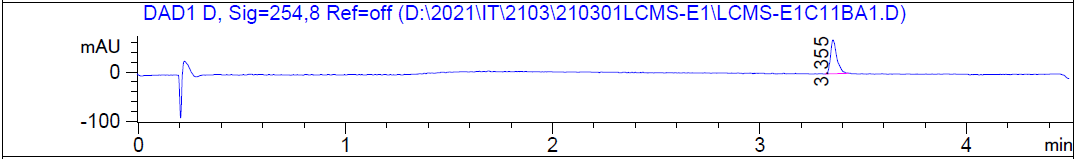

**^1^H NMR:** (400 MHz, CD_3_OD) δ 8.35 (d, *J* = 8.3 Hz, 1H), 8.26 (s, 1H), 7.83 (d, *J* = 8.7 Hz, 2H), 7.71 (d, *J* = 8.7 Hz, 2H), 7.55– 7.44 (m, 1H), 7.33– 7.08 (m, 1H), 7.04– 6.62 (m, 6H), 6.61–6.54 (m, 2H), 6.53– 6.44 (m, 1H), 5.59– 5.50 (m, 1H), 5.38 (brs, 1H), 4.96– 4.87 (m, 2H), 4.83– 4.78 (m, 1H), 4.12– 4.01 (m, 1H), 3.89– 3.62 (m, 17H), 3.62– 3.41 (m, 24H), 3.17–2.93 (m, 5H), 2.73 (brt, *J* = 12.9 Hz, 1H), 2.65– 2.33 (m, 2H), 2.27 (brd, *J* = 13.3 Hz, 1H), 2.15–1.94 (m, 2H), 1.91– 1.79 (m, 1H), 1.73 (brs, 4H), 1.42 (brs, 2H), 0.95– 0.76 (m, 3H)

HRMS [C_70_H_90_BrFN_8_O_19_S] Cal: 1479.5263; Obs: 1479.5227

**Preparation of compound 11**

To a solution of 2,2'-(ethane-1,2-diylbis(oxy))bis(ethan-1-ol) (10 g, 30.64 mmol, 1.0 equiv) in CH_2_Cl_2_ (150 ml) was added Ag_2_O (10.65 g, 45.96 mmol, 1.5 equiv), NaI (5 g, 33.36 mmol, 1.0 equiv) and TosCl (5.84 g, 30.64 mmol, 1.0 equiv). The mixture was stirred at 20 °C for 12 h. The reaction mixture was filtered and concentrated under reduced pressure to give a residue. The residue was purified by column chromatography (SiO_2_, **Petroleum ether / EtOAc = 1/3** to 0**/1 to EtOAc/MeOH = 20/1**). 2-(2-(2-hydroxyethoxy)ethoxy)ethyl 4-methylbenzenesulfonate (6.0 g, 12.49 mmol, 40% yield) was obtained as a yellow oil. (ethane-1,2-diylbis(oxy))bis(ethane-2,1-diyl) bis(4-methylbenzenesulfonate) (1.77 g, 2.79 mmol, 9% yield) was obtained as a yellow oil.

**^1^HNMR** (400 MHz, CDCl_3_): δ 7.81 (d, *J* = 8.3 Hz, 2H), 7.35 (d, *J* = 8.1 Hz, 2H), 4.19– 4.14 (m, 2H), 3.76– 3.57 (m, 26H), 2.46 (s, 3H)

**Preparation of compound 13**

A mixture of 2-(2-(2-hydroxyethoxy)ethoxy)ethyl 4-methylbenzenesulfonate (6.0 g, 12.49 mmol, 1.0 equiv), *N*-methyl-1-phenylmethanamine (3.03 g, 24.97 mmol, 3.22 ml, 2.0 equiv), K_2_CO_3_ (5.18 g, 37.46 mmol, 3.0 equiv) in MeCN (60 ml) was stirred at 80 °C for 12 h. The reaction mixture was filtered and concentrated under reduced pressure to give a residue. The residue was purified by *prep*-HPLC (column: Phenomenex luna C18 250*50 mm*10 µm;mobile phase: [water(10 mM NH_4_HCO_3_)–ACN]; B%: MeCN 30%–60 % MeCN,18 min). 2-(2-(2-(benzyl(methyl)amino)ethoxy)ethoxy)ethan-1-ol (3.7 g, 8.61 mmol, 68% yield) was obtained as a yellow oil.

**^1^H NMR** (400 MHz, CDCl_3_): δ 7.53– 7.30 (m, 5H), 3.73– 3.63 (m, 28H), 2.92– 2.61 (m, 3H), 2.54– 2.22 (m, 3H)

**LC–MS:** MS (ES^+^): RT = 0.813 min, m/z = 430.3 [M + H^+^]

**Spectra:**

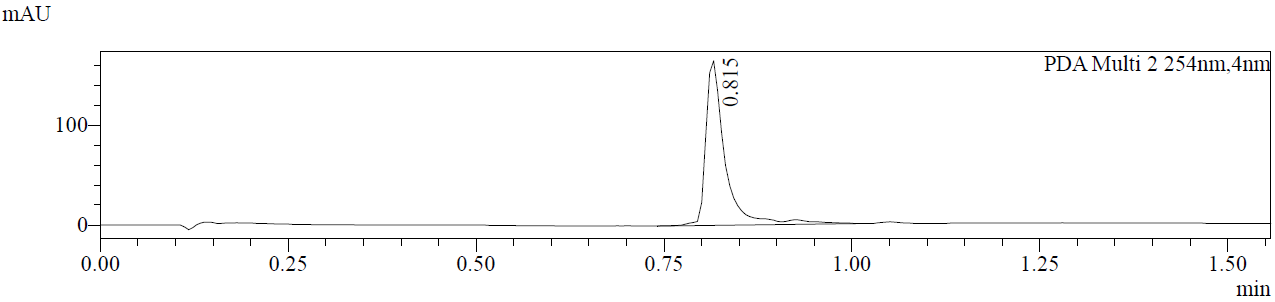

**Preparation of compound 14**

To a solution of 2-(2-(2-(benzyl(methyl)amino)ethoxy)ethoxy)ethan-1-ol (3.7 g, 8.61 mmol, 1.0 equiv) and (Boc)_2_O (2.07 g, 9.48 mmol, 2.1 ml, 1.1 equiv) in THF (60 ml) was added Pd/C (370 mg, 86 µmol, 10% purity, 0.10 equiv). The mixture was degassed and purged 3 times with H_2_, and then the mixture was stirred at 20 °C for 12 h under H_2_ atmosphere. The reaction mixture was filtered and concentrated under reduced pressure to give a residue. The product *tert*-butyl (2-(2-(2-hydroxyethoxy)ethoxy)ethyl)(methyl)carbamate (3.7 g, crude) was used into the next step without further purification.

**LC–MS:** MS (ES^+^): RT = 0.897 min, m/z = 457.3 [M + 18]

**Spectra:**

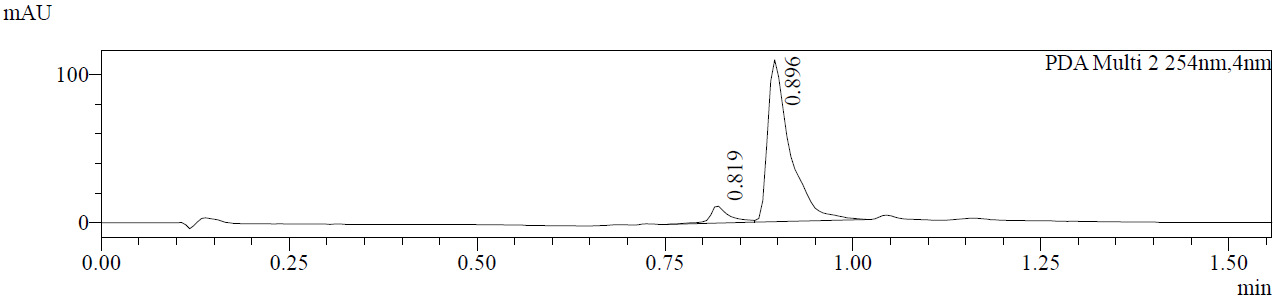

**Preparation of compound 15**

To a solution of *tert*-butyl (2-(2-(2-hydroxyethoxy)ethoxy)ethyl)(methyl)carbamate (3.7 g, 8.42 mmol, 1.0 equiv) in CH_2_Cl_2_ (40 ml) was added Et_3_N (2.54 g, 25.15 mmol, 3.5 ml, 2.9 equiv), TosCl (2.41 g, 12.63 mmol, 1.5 equiv). The mixture was stirred at 20 °C for 12 h. The reaction mixture was concentrated under reduced pressure to give a residue. The residue was purified by column chromatography (SiO_2_, EtOAc to CH_2_Cl_2_: MeOH=20/1). 2,2,5-Trimethyl-4-oxo-3,8,11-trioxa-5-azatridecan-13-yl 4-methylbenzenesulfonate (4.7 g, 7.92 mmol, 94% yield) was obtained as a yellow oil.

**^1^H NMR** (400 MHz, CDCl_3_): δ 7.81 (d, *J* = 8.3 Hz, 2H), 7.35 (d, *J* = 8.1 Hz, 2H), 4.23– 4.12 (m, 2H), 3.72– 3.55 (m, 24H), 3.39 (m, 2H), 2.91 (s, 3H), 2.46 (s, 3H), 1.46 (s, 9H)

**LC–MS:** MS (ES^+^): RT = 0.950 min, m/z = 494.2 [M - 99]

**Preparation of compound 1**

A mixture of 2,2,5-trimethyl-4-oxo-3,8,11-trioxa-5-azatridecan-13-yl 4-methylbenzenesulfonate (4.37 g, 7.36 mmol, 1.0 equiv) and LiBr (3.20 g, 36.80 mmol, 923 µl, 5.0 *equiv*) in acetone (40 ml) was stirred at 70 °C for 12 h. The reaction mixture was partitioned between H_2_O (20 ml) and EtOAc (50 ml.) The organic phase was separated, washed with brine (20 ml), dried over [Na_2_SO_4_], filtered, and concentrated under reduced pressure to give a residue. The product *tert*-butyl (2-(2-(2-bromoethoxy)ethoxy)ethyl)(methyl)carbamate (4 g, crude) was used into the next step without further purification.

**^1^HNMR** (400 MHz, CDCl_3_): δ 3.82 (t, *J* = 6.3 Hz, 2H), 3.70– 3.58 (m, 22H), 3.48 (t, *J* = 6.3 Hz, 2H), 3.43– 3.35 (m, 2H), 2.91 (s, 3H), 1.46 (s, 9H)

**The synthetic route for HLDA-213**

Part 1.

Part 2.

**Preparation of compound 7**

Known compound from *Angew. Chem., Int. Ed.* **2020**, *59*, 13865–13870.

**Preparation of compound 9**

known compound from *J. Med. Chem*., **2000**, *43*, 1135–1142

**Preparation of compound 2**

To a solution of tert-butyl *N*-[2-[2-[2-[2-[2-[2-[2-[2-(2-bromoethoxy)ethoxy]ethoxy]ethoxy]ethoxy]ethoxy]

ethoxy]ethoxy]ethyl]-*N*-methyl-carbamate (1.3 g, 2.20 mmol, 1.0 equiv) in DMF (5 ml) was added NaN_3_ (0.36 g, 5.54 mmol, 2.52 equiv), and then it was stirred at 70 °C for 12 h. The reaction mixture was poured into 50 ml water and extracted with EtOAc (2 x 20 ml). The organic layers were washed with water (3 x 20 ml), dried over anhydrous Na_2_SO_4_, filtered, and concentrated under reduced pressure to afford crude product. *tert*-butyl *N*-[2-[2-[2-[2-[2-[2-[2-[2-(2-azidoethoxy)ethoxy]ethoxy]ethoxy]ethoxy]ethoxy]ethoxy]ethoxy]ethyl]-*N*-methyl-carbamate (1.15 g, 2.08 mmol, 94.53% yield) was obtained as a yellow oil and used for the next step directly.

**LC–MS:** MS (ES^+^): RT = 0.786 min, m/z = 570.3 [M + 18] ^+^

**Spectra:**

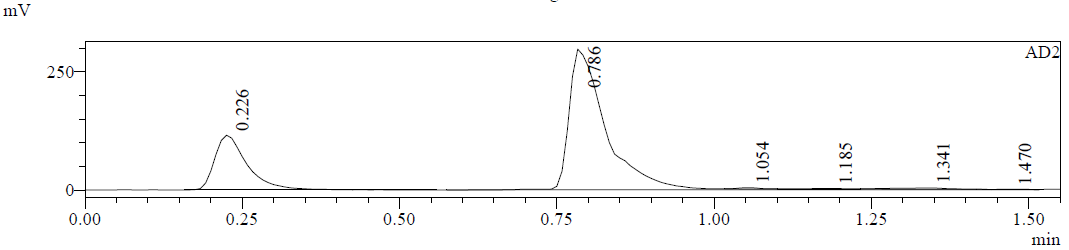

**Preparation of compound 3**

To a solution of *tert*-butyl *N*-[2-[2-[2-[2-[2-[2-[2-[2-(2-azidoethoxy)ethoxy]ethoxy]ethoxy]ethoxy]ethoxy]ethoxy]ethoxy]ethyl]-*N*-methyl-carbamate (1.15 g, 2.08 mmol, 1.0 equiv) in THF (30 ml) was added Pd/C (0.3 g, 10% purity), and then it was degassed and purged with H_2_. The reaction mixture was stirred at 25 °C for 12 h under 15 psi pressure. The reaction mixture was filtered, and the filtrate was concentrated to afford crude product. *tert*-Butyl *N*-[2-[2-[2-[2-[2-[2-[2-[2-(2-aminoethoxy)ethoxy]ethoxy]ethoxy]ethoxy]ethoxy]ethoxy]ethoxy]ethyl]-*N*-methyl-carbamate (1.05 g, 1.99 mmol, 95.81% yield) was obtained as a colorless oil and used for the next step directly.

**LC–MS:** MS (ES^+^): RT = 0.562 min, m/z = 527.3 [M + H] ^+^

**Preparation of compound 5**

To a solution of *tert*-butyl *N*-[2-[2-[2-[2-[2-[2-[2-[2-(2-aminoethoxy)ethoxy]ethoxy]ethoxy]

ethoxy]ethoxy]ethoxy]ethoxy]ethyl]-*N*-methyl-carbamate (1.05 g, 1.99 mmol, 1.0 equiv) and DIEA (515.34 mg, 3.99 mmol, 694.53 µl, 2.0 equiv) in CH_2_Cl_2_ (5 ml) was added 4-nitrobenzenesulfonyl chloride (530.21 mg, 2.39 mmol, 1.2 equiv), and then it was stirred at 20 °C for 1 h. The reaction mixture was quenched by 5 mlMeOH, and then it was concentrated to afford crude product. The residue was purified by *prep*-HPLC (column: Waters Xbridge C18 150*50 mm*10 µm;mobile phase: [water(10 mM NH_4_HCO_3_)–MeCN]; B%: 37%–67%, 11 min) to afford *tert*-butyl *N*-methyl-*N*-[2-[2-[2-[2-[2-[2-[2-[2-[2-[(4-nitrophenyl)sulfonylamino]ethoxy]ethoxy]ethoxy]ethoxy]ethoxy]ethoxy]ethoxy]ethoxy]ethyl]carbamate (0.52 g, 730.53 µmol, 36.64% yield) as a yellow oil.

**LC-MS:** MS (ES^+^): RT = 0.680 min, m/z = 612.6 [M -100 + H] ^+^

**Spectra:**

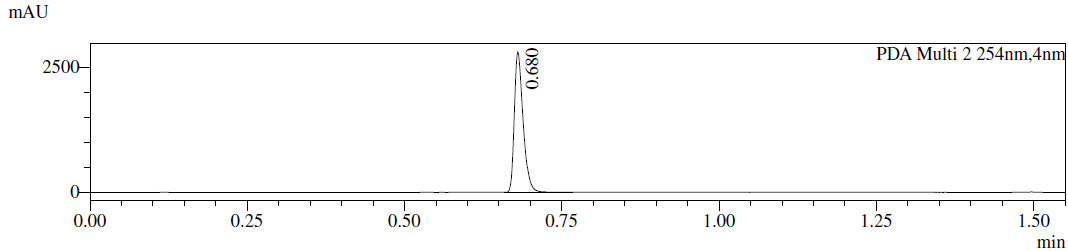

**Preparation of compound 6**

To a solution of tert-butyl *N*-methyl-*N*-[2-[2-[2-[2-[2-[2-[2-[2-[2-[(4-nitrophenyl)sulfonylamino]ethoxy]

ethoxy]ethoxy]ethoxy]ethoxy]ethoxy]ethoxy]ethoxy]ethyl]carbamate (0.26 g, 365.26 µmol, 1.0 equiv) in THF (5 ml) was added Pd/C (0.1 g, 10% purity), and then it was degassed and purged with H_2_. The resulting solution was stirred at 25 °C for 12 h under 15 psi pressure. The reaction mixture was filtered and the filtrate was concentrated under reduced pressure to afford crude product. *tert*-Butyl *N*-[2-[2-[2-[2-[2-[2-[2-[2-[2-[(4-aminophenyl)sulfonylamino]ethoxy]ethoxy]ethoxy]ethoxy]ethoxy]ethoxy]ethoxy]ethoxy]ethyl]-*N*-methyl-carbamate (0.24 g, 351.99 µmol, 96.37% yield) was obtained as a colorless oil and used for the next step directly.

**LC–MS:** MS (ES^+^): RT = 0.911 min, m/z = 699.5 [M + H_3_O] ^+^

**Spectra:**

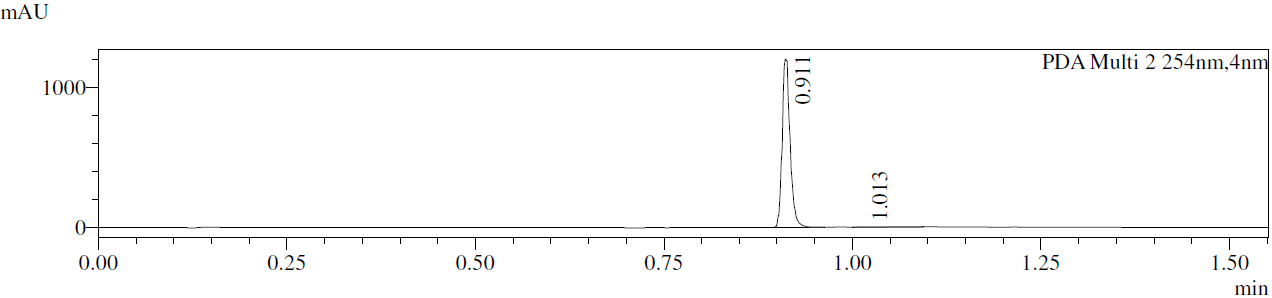

**Preparation of compound 8**

To a solution of *tert*-butyl *N*-[2-[2-[2-[2-[2-[2-[2-[2-[2-[(4-aminophenyl)sulfonylamino]ethoxy]ethoxy]

ethoxy]ethoxy]ethoxy]ethoxy]ethoxy]ethoxy]ethyl]-*N*-methyl-carbamate (150 mg, 220 µmol, 1.0 equiv) and 2-[(5-bromo-2-chloro-pyrimidin-4-yl)amino]-6-fluoro-benzamide (91.2 mg, 264 µmol, 1.2 equiv) in *i*-PrOH (3 mL) was added HCl (12 M, 27.50 µl, 1.5 equiv), and then it was stirred at 100 °C for 12 h. The reaction mixture was concentrated to afford crude product. The residue was purified by *prep*-HPLC (column: Phenomenex luna C18 150*40 mm*15 µm;mobile phase: [water(0.1%TFA)–MeCN];B%: 16%–46%,11 min) to afford 2-[[5-bromo-2-[4-[2-[2-[2-[2-[2-[2-[2-[2-[2-(methylamino)ethoxy]ethoxy]

ethoxy]ethoxy]ethoxy]ethoxy]ethoxy]ethoxy]ethylsulfamoyl]anilino]pyrimidin-4-yl]amino]-6-fluoro-benzamide (75 mg, 74.64 µmol, 33.93% yield, TFA salt) as a colorless gum.

**LC–MS:** MS (ES^+^): RT = 0.727 min, m/z = 891.7 [M + 2] ^+^

**Spectra:**

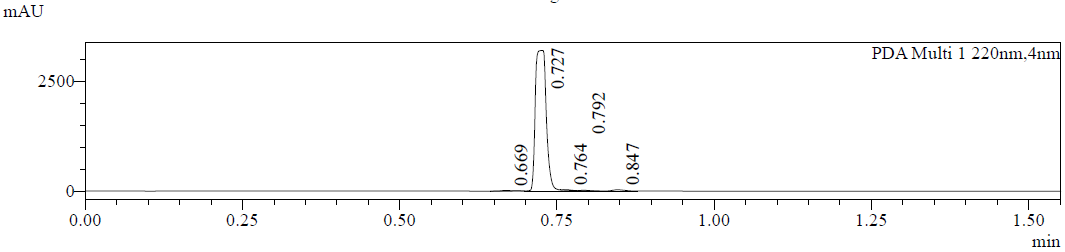

**Preparation of compound HLDA-213**

To a solution of 2-[[5-bromo-2-[4-[2-[2-[2-[2-[2-[2-[2-[2-[2-(methylamino)ethoxy]ethoxy]ethoxy]ethoxy]

ethoxy]ethoxy]ethoxy]ethoxy]ethylsulfamoyl]anilino]pyrimidin-4-yl]amino]-6-fluoro-benzamide (100 mg, 99.5 µmol, 1.0 equiv, TFA salt) and 2-[3-[(1*R*)-3-(3,4-dimethoxyphenyl)-1-[(2*S*)-1-[(2*S*)-2-(3,4,5-

trimethoxyphenyl)butanoyl]piperidine-2-carbonyl]oxy-propyl]phenoxy]acetic acid (69.1 mg, 99.5 µmol, 1.0 equiv) in DMF (2 ml) was added DIEA (38.6 mg, 299 µmol, 52.00 µl, 3.0 equiv) and HATU (45.4 mg, 119 µmol, 1.2 equiv), and then it was stirred at 25 °C for 1 h. The residue was purified by *prep*-HPLC (column: Phenomenex Gemini-NX C18 75*30 mm*3 µm;mobile phase: [water(10 mM NH_4_HCO_3_)–MeCN];B%: 44%–74%, 8 min) and (column: Phenomenex Gemini-NX C18 75*30 mm*3 µm;mobile phase: [water(0.225%FA)–MeCN]; B%: 55%–85%, 7 min) to afford [(1*R*)-1-[3-[2-[2-[2-[2-[2-[2-[2-[2-[2-[2-[[4-[[5-bromo-4-(2-carbamoyl-3-fluoro-anilino)pyrimidin-2-yl]amino]phenyl]sulfonylamino]ethoxy]ethoxy]ethoxy]ethoxy]ethoxy]ethoxy]ethoxy]ethoxy]ethyl-methyl-amino]-2-oxo-ethoxy]phenyl]-3-(3,4-dimethoxyphenyl)propyl] (2*S*)-1-[(2*S*)-2-(3,4,5-trimethoxyphenyl)butanoyl]

piperidine-2-carboxylate (22 mg, 13.64 µmol, 14% yield, FA salt) as a yellow solid.

**LC–MS:** MS (ES^+^): RT = 3.348 min, m/z = 784.4 [M/2 + H^+^]

**^1^H NMR:** (400 MHz, CD_3_OD) δ 8.35 (d, *J* = 8.3 Hz, 1H), 8.26 (s, 1H), 7.83 (d, *J* = 8.7 Hz, 2H), 7.71 (d, *J* = 8.7 Hz, 2H), 7.55– 7.44 (m, 1H), 7.33– 7.08 (m, 1H), 7.04– 6.62 (m, 6H), 6.61–6.54 (m, 2H), 6.53– 6.44 (m, 1H), 5.59– 5.50 (m, 1H), 5.38 (brs, 1H), 4.96– 4.87 (m, 2H), 4.83– 4.78 (m, 1H), 4.12– 4.01 (m, 1H), 3.89– 3.62 (m, 17H), 3.62– 3.41 (m, 32H), 3.17– 2.93 (m, 5H), 2.73 (brt, *J* = 12.9 Hz, 1H), 2.65– 2.33 (m, 2H), 2.27 (brd, *J* = 13.3 Hz, 1H), 2.15–1.94 (m, 2H), 1.91– 1.79 (m, 1H), 1.73 (brs, 4H), 1.42 (brs, 2H), 0.95– 0.76 (m, 3H)

**Spectra:**

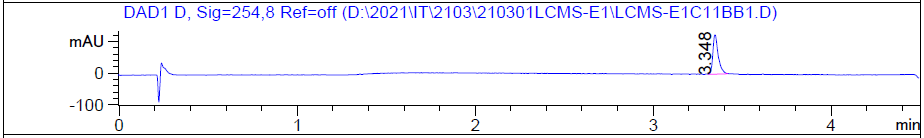

**Preparation of compound 11**

To a solution of 2-[2-[2-[2-[2-[2-[2-[2-(2-hydroxyethoxy)ethoxy]ethoxy]ethoxy]ethoxy]ethoxy]ethoxy]ethoxy]ethyl 4-methylbenzenesulfonate (4.40 g, 7.74 mmol, 1.0 equiv) in MeCN (40 ml) was added K_2_CO_3_ (3.21 g, 23.2 mmol, 3.0 equiv) and *N*-methyl-1-phenyl-methanamine (1.88 g, 15.5 mmol, 2.00 ml, 2.0 equiv), and then it was stirred at 80 °C for 12 h. The reaction mixture was filtered, and the filtrate was concentrated to afford crude product. The residue was purified by *prep*-HPLC (column: Phenomenex luna C18 250*50 mm*10 µm; mobile phase: [water(10 mM NH_4_HCO_3_)-MeCN]; B%: 30%MeCN–60% MeCN, 18 min) to afford 2-[2-[2-[2-[2-[2-[2-[2-[2-[benzyl(methyl)amino]ethoxy]ethoxy]ethoxy]ethoxy]ethoxy]ethoxy]ethoxy]ethoxy]ethanol (3.0 g, 5.8 mmol, 75% yield) as a yellow oil.

**Spectra:**

**LC–MS:** MS (ES^+^): RT = 0.868 min, m/z = 518.3 [M + H^+^]

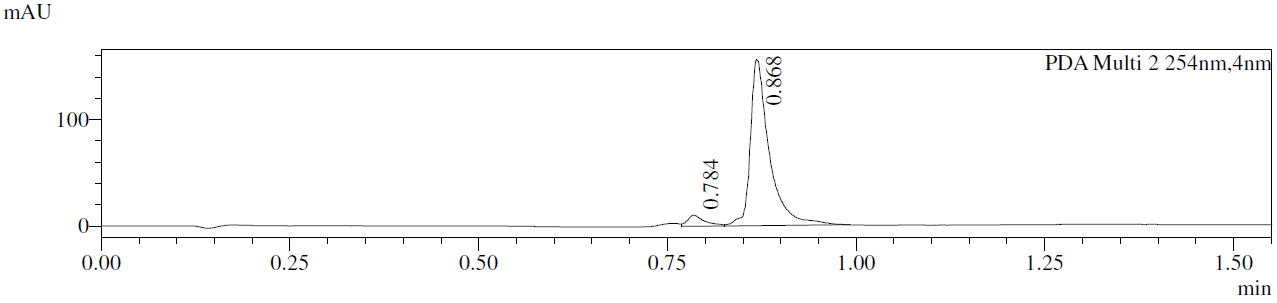

**^1^H NMR:** (400 MHz, CDCl_3_) δ 7.33– 7.22 (m, 5H), 3.72 (m, 2H), 3.69– 3.60 (m, 32H), 3.55 (s, 2H), 2.82 (br s, 1H), 2.62 (t, *J* = 6.1 Hz, 2H), 2.26 (s, 3H)

**Preparation of compound 12**

To a solution of 2-[2-[2-[2-[2-[2-[2-[2-[2-[benzyl(methyl)amino]ethoxy]ethoxy]ethoxy]ethoxy]ethoxy]ethoxy]ethoxy]ethoxy]ethanol (3.0 g, 5.8 mmol, 1.0 equiv) in THF (20 ml) was added Pd/C (600 mg, 10% purity) and Boc_2_O (1.39 g, 6.37 mmol, 1.46 ml, 1.1 equiv), and then it was degassed and purged with H_2_. The reaction mixture was stirred at 25 °C for 12 h under 15 psi pressure. The reaction mixture was filtered, and the filtrate was concentrated to afford *tert*-butyl *N*-[2-[2-[2-[2-[2-[2-[2-[2-(2-hydroxyethoxy)ethoxy]ethoxy]ethoxy]ethoxy]ethoxy]ethoxy]ethoxy]ethyl]-*N*-methyl-carbamate (3.1 g, crude) as a colorless oil and used for the next step directly.

**Spectra:**

**^1^H NMR**: (400 MHz, CDCl_3_) δ 3.75– 3.72 (m, 2H), 3.70– 3.58 (m, 32H), 3.42–3.36 (m, 2H), 2.92 (s, 3H), 1.46 (s, 9H)

**Preparation of compound 13**

To a solution of *tert*-butyl *N*-[2-[2-[2-[2-[2-[2-[2-[2-(2-hydroxyethoxy)ethoxy]ethoxy]ethoxy]ethoxy]ethoxy]ethoxy]ethoxy]ethyl]-*N*-methyl-carbamate (3.10 g, 5.88 mmol, 1.0 equiv) and Et_3_N (1.19 g, 11.8 mmol, 1.60 ml, 2.0 equiv) in CH_2_Cl_2_ (20 ml) was added TosCl (1.68 g, 8.81 mmol, 1.5 equiv), and then it was stirred at 25 °C for 12 h. The reaction mixture was concentrated to afford crude product. The residue was purified by silica chromatography (Petroleum ether: EtOAc = 1:1–0:1) to afford 2-[2-[2-[2-[2-[2-[2-[2-[2-[*tert*-butoxycarbonyl(methyl)amino]ethoxy]ethoxy]ethoxy]ethoxy]ethoxy]ethoxy]ethoxy]ethoxy]ethyl 4-methylbenzenesulfonate (3.60 g, 5.28 mmol, 90% yield) as a colorless oil.

**Spectra:**

**^1^H NMR**: (400 MHz, CDCl_3_) δ 7.80 (d, *J* = 8.4 Hz, 2H), 7.35 (d, *J* = 8.1 Hz, 2H), 4.21– 4.13 (m, 2H), 3.70 (m, 2H), 3.67– 3.55 (m, 30H), 3.39 (m, 2H), 2.91 (s, 3H), 2.45 (s, 3H), 1.46 (s, 9H)

**Preparation of compound 1**

To a solution of 2-[2-[2-[2-[2-[2-[2-[2-[2-[*tert*-butoxycarbonyl(methyl)amino]ethoxy]ethoxy]ethoxy]ethoxy]ethoxy]ethoxy]ethoxy]ethoxy]ethyl 4-methylbenzenesulfonate (3.60 g, 5.28 mmol, 1.0 equiv) in acetone (30 ml) was added LiBr (2.29 g, 26.4 mmol, 5.0 equiv), and then it was stirred at 75 °C for 12 h. The reaction mixture was poured into 100 ml water, and then it was extracted with EtOAc (3 x 30 ml). The organic layers were dried over anhydrous Na_2_SO_4_, filtered and concentrated to afford *tert*-butyl *N*-[2-[2-[2-[2-[2-[2-[2-[2-(2-bromoethoxy)ethoxy]ethoxy]ethoxy]ethoxy]ethoxy]ethoxy]ethoxy]ethyl]-*N*-methyl-carbamate (3.0 g, 5.1 mmol, 96% yield) as a yellow oil.

**Spectra:**

**^1^H NMR**: (400 MHz, CDCl_3_) δ 3.81 (t, *J* = 6.3 Hz, 2H), 3.70 - 3.55 (m, 30H), 3.48 (t, *J* = 6.3 Hz, 2H), 3.44– 3.35 (m, 2H), 2.91 (s, 3H), 1.45 (s, 9H)

***SERIES 10***

HLDA-001

The synthetic route for HLDA-001

**Preparation of compound 1**

known compound for *J. Med. Chem.* 2019, 62, 5191–5216.

**Preparation of HLDA-001**

To the solution of 2-[3-[(1*R*)-3-(3,4-dimethoxyphenyl)-1-[(2*S*)-1-[(2*S*)-2-(3,4,5-trimethoxyphenyl)butanoyl]piperidine-2-carbonyl]oxy-propyl]phenoxy]acetic acid (50 mg, 72 µmol, 1.0 equiv) and 2-methoxy-*N*-methyl-ethanamine (10 mg, 112 µmol, 12 µl, 1.6 equiv) in DMF (1 ml) was added DIEA (30 mg, 230 µmol, 0.04 ml, 3.2 equiv) and HATU (41 mg, 108 µmol, 1.5 equiv), then the solution was stirred at 25 °C for 1 h.  The solution was filtered to get the filtrate, which was purified by *prep*-HPLC (column: Phenomenex Gemini-NX C18 75*30 mm*3 µm;mobile phase: [water(0.225%FA–MeCN];B%: 48%–78%, 7 min) to get [(1*R*)-3-(3,4-dimethoxyphenyl)-1-[3-[2-[2-methoxyethyl(methyl)amino]-2-oxo-ethoxy]phenyl]propyl] (2*S*)-1-[(2*S*)-2-(3,4,5-trimethoxyphenyl)butanoyl]piperidine-2-carboxylate (20 mg, 36% yield) as a yellow solid.

**^1^H NMR** (400 MHz, CDCl_3_): δ 7.25–7.09 (m, 1H), 6.99–6.60 (m, 3H), 6.70–6.58 (m, 3H), 6.49–6.34 (m, 2H), 5.88–5.39 (m, 2H), 4.87–4.52 (m, 2H), 3.92–3.49 (m, 21H), 3.38–3.29 (m, 3H), 3.22–2.76 (m, 4H), 2.63–2.42 (m, 2H), 2.38–1.98 (m, 4H), 1.76–1.52 (m, 4H), 1.42–1.17 (m, 2H), 0.93–0.81 (m, 3H)

**LC–MS:** MS (ES^+^): RT = 2.542 min, m/z = 765.3 [M + H^+^];

**Spectra:**

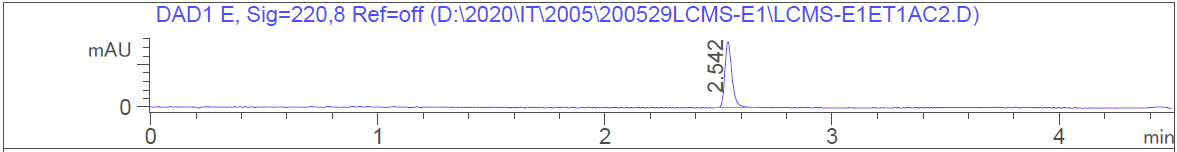

HRMS [C_42_H_56_N_2_O_11_] Cal: 765.3957; Obs: 765.3924

**NMR Spectra**

**HLDA-123**

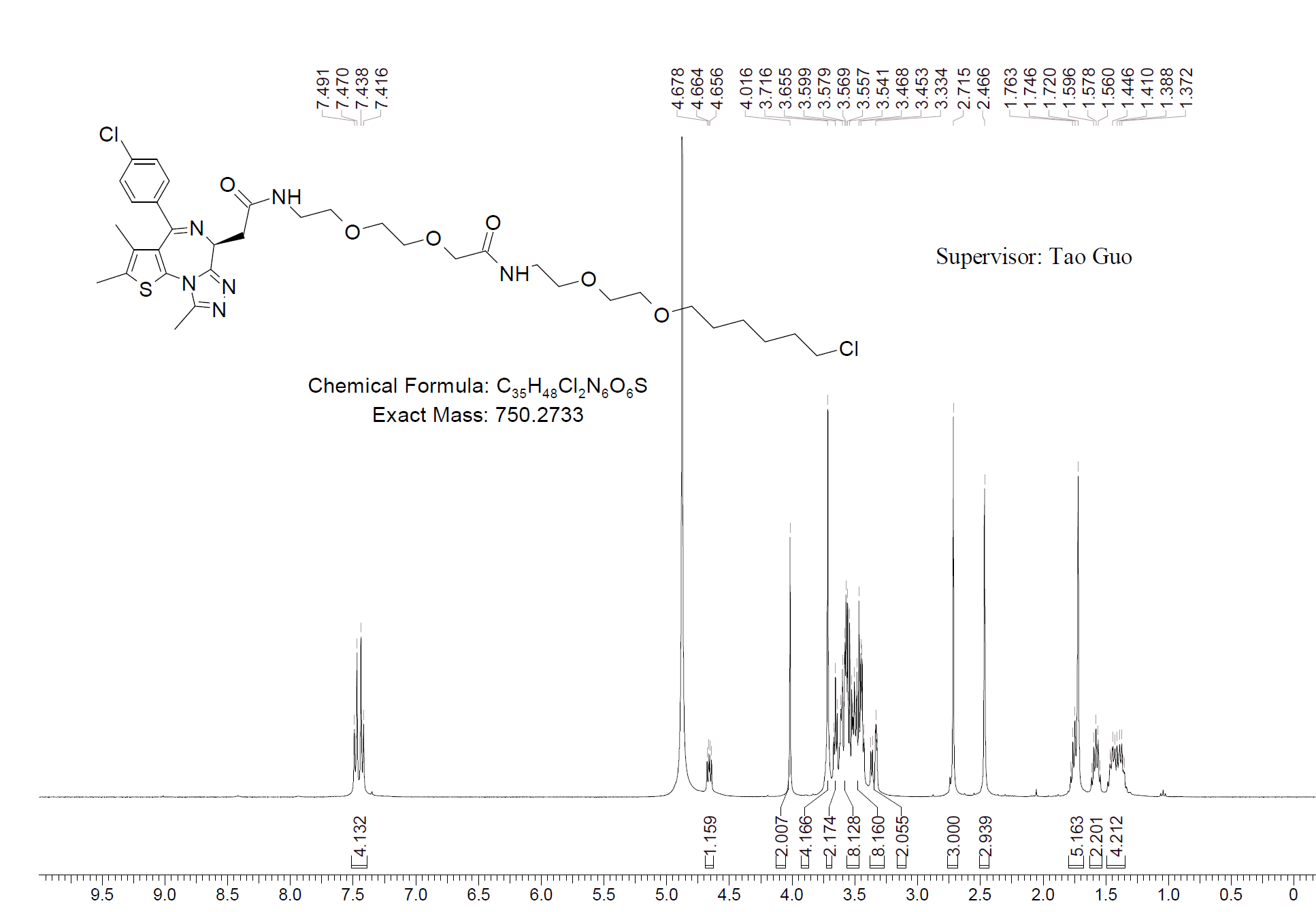

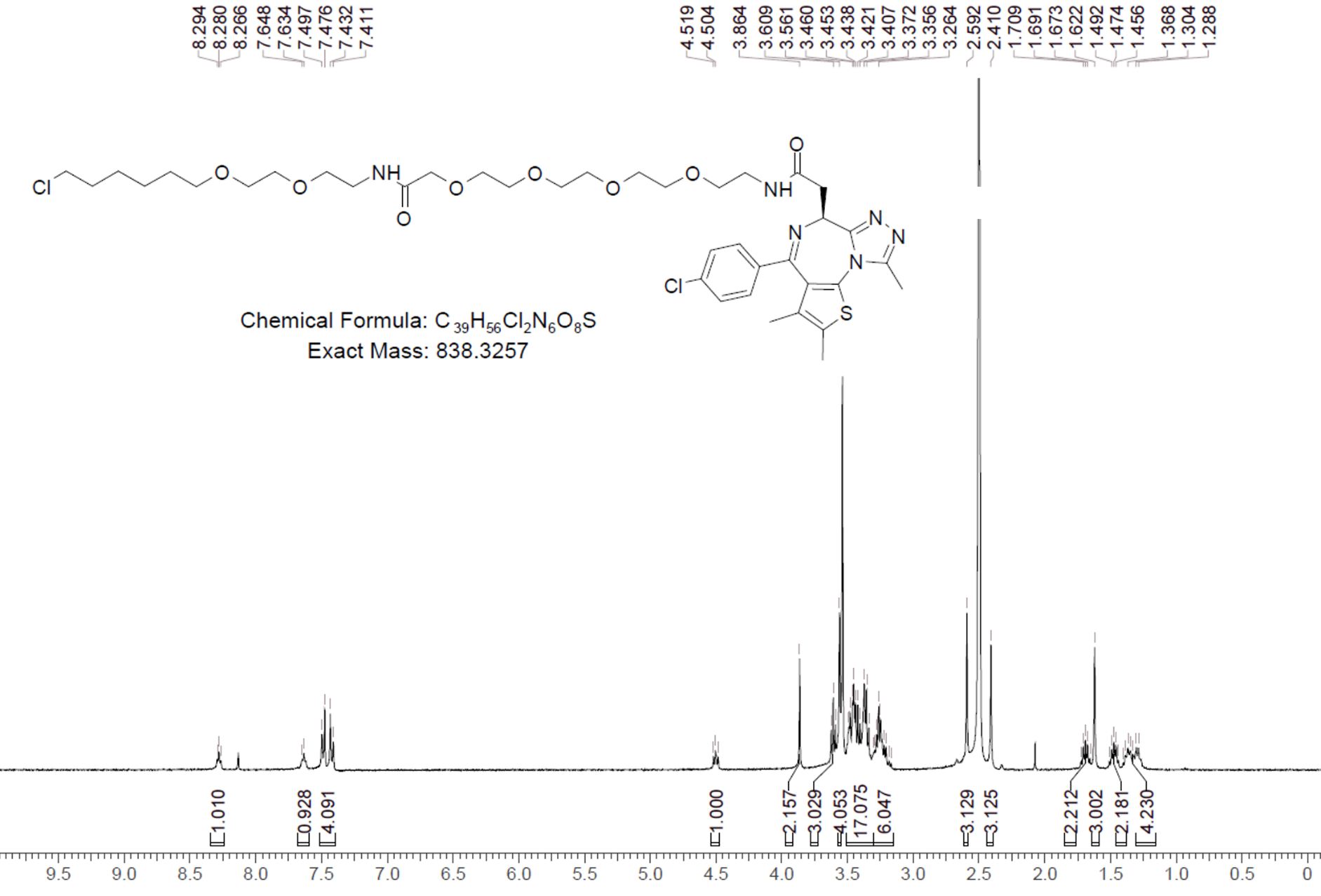
**HLDA-124**

**HLDA-121**

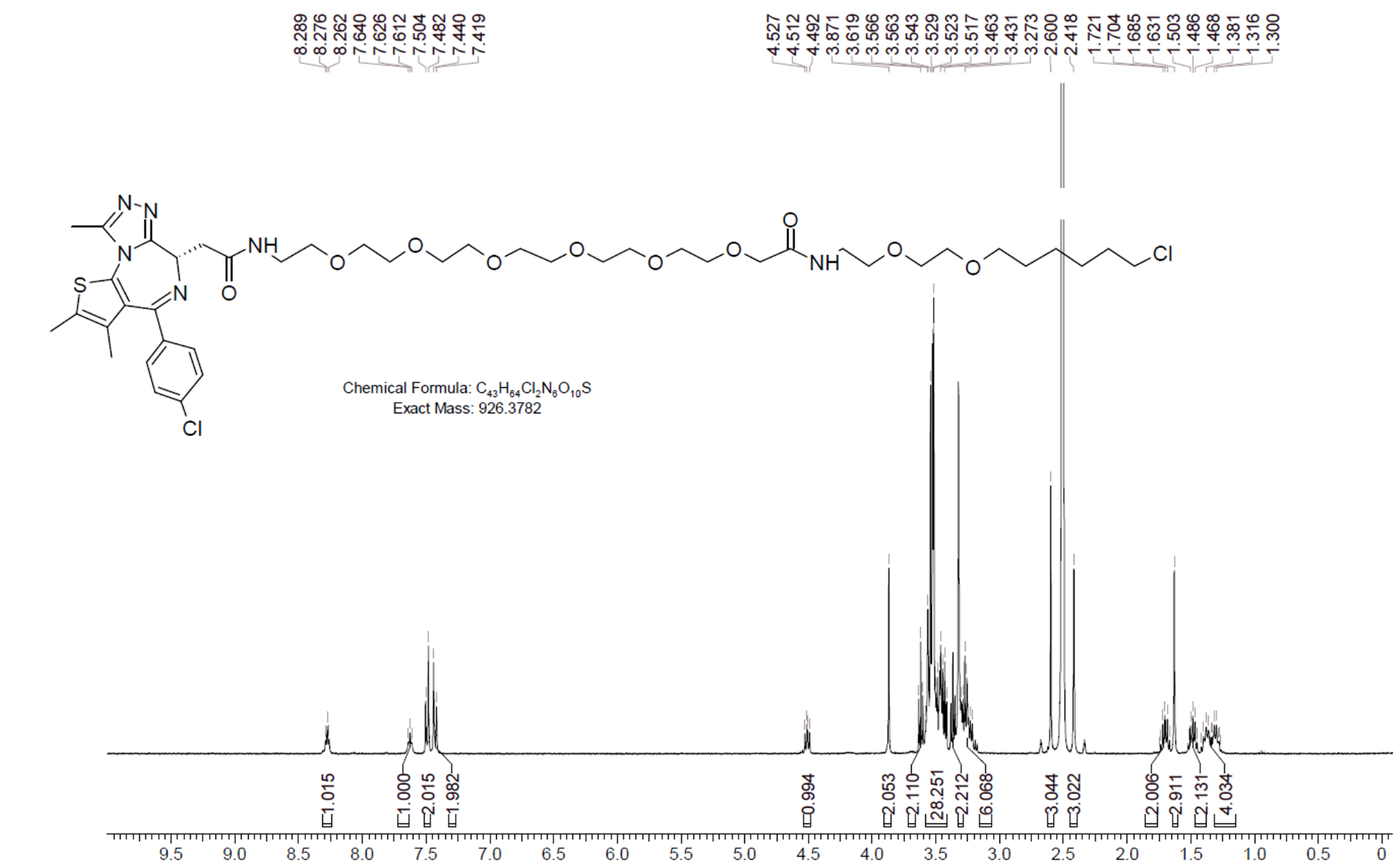

**HLDA-125 – Compound 3**

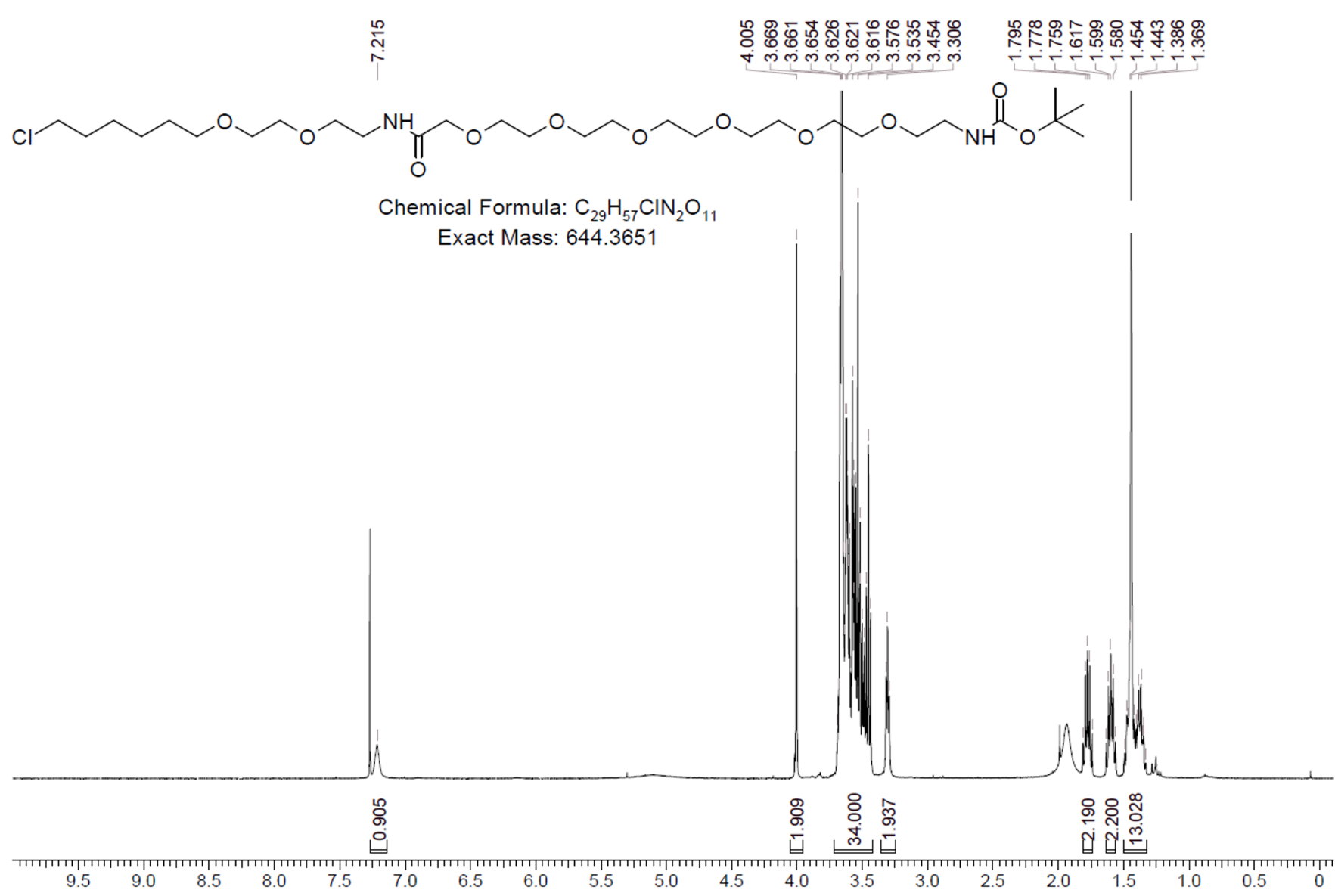

**HLDA-125 – Compound 4**

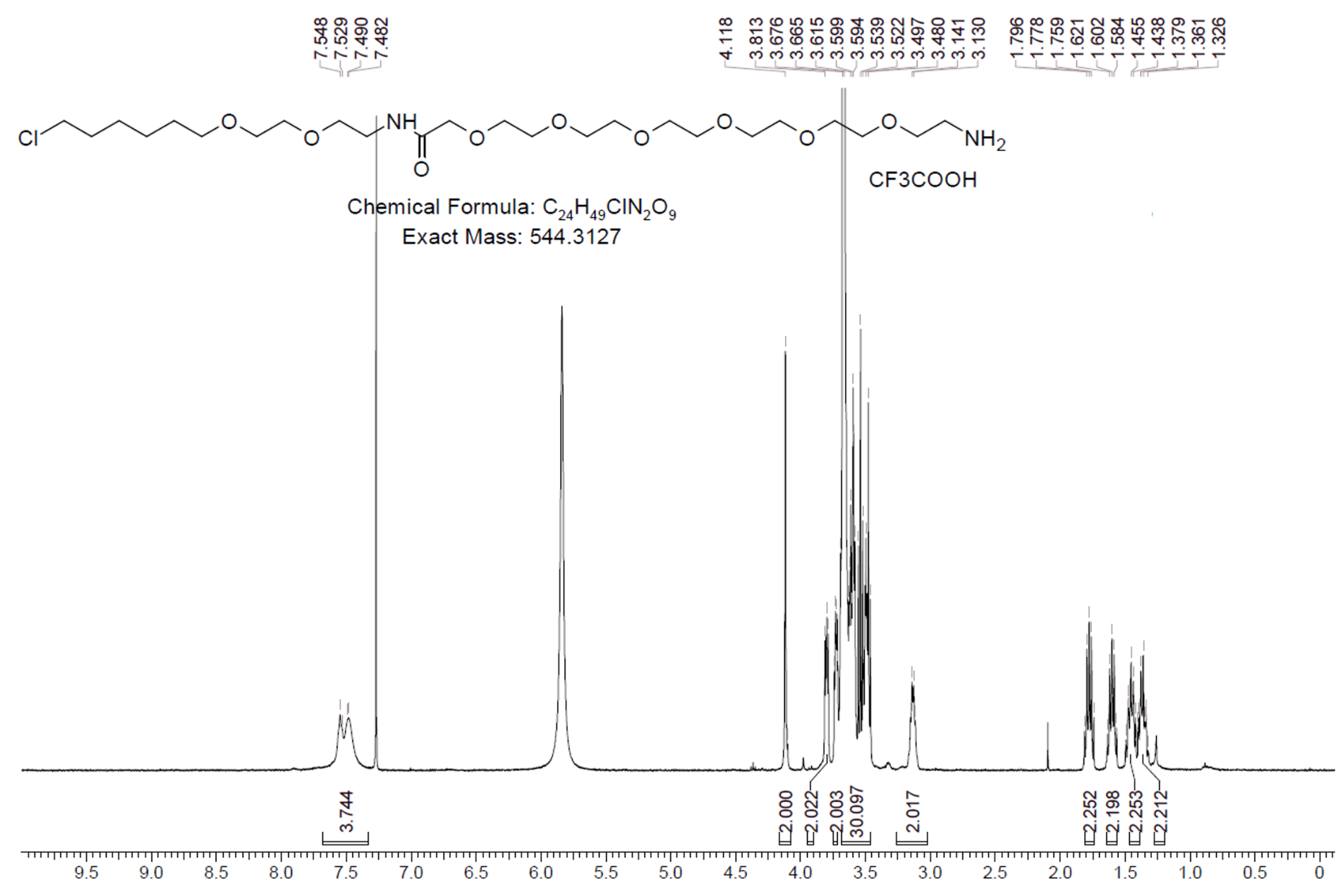

**HLDA-125**

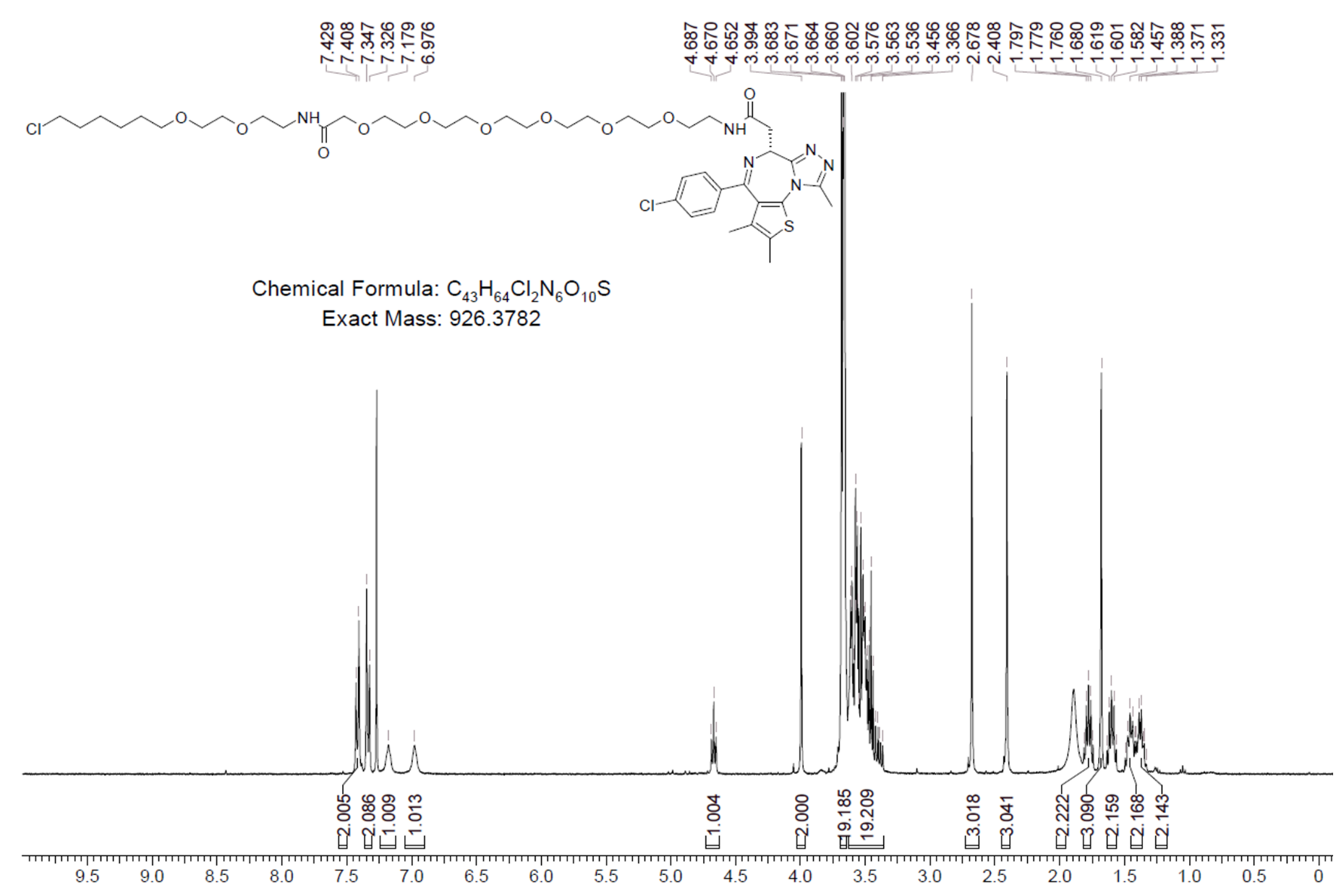

**HLDA-112**

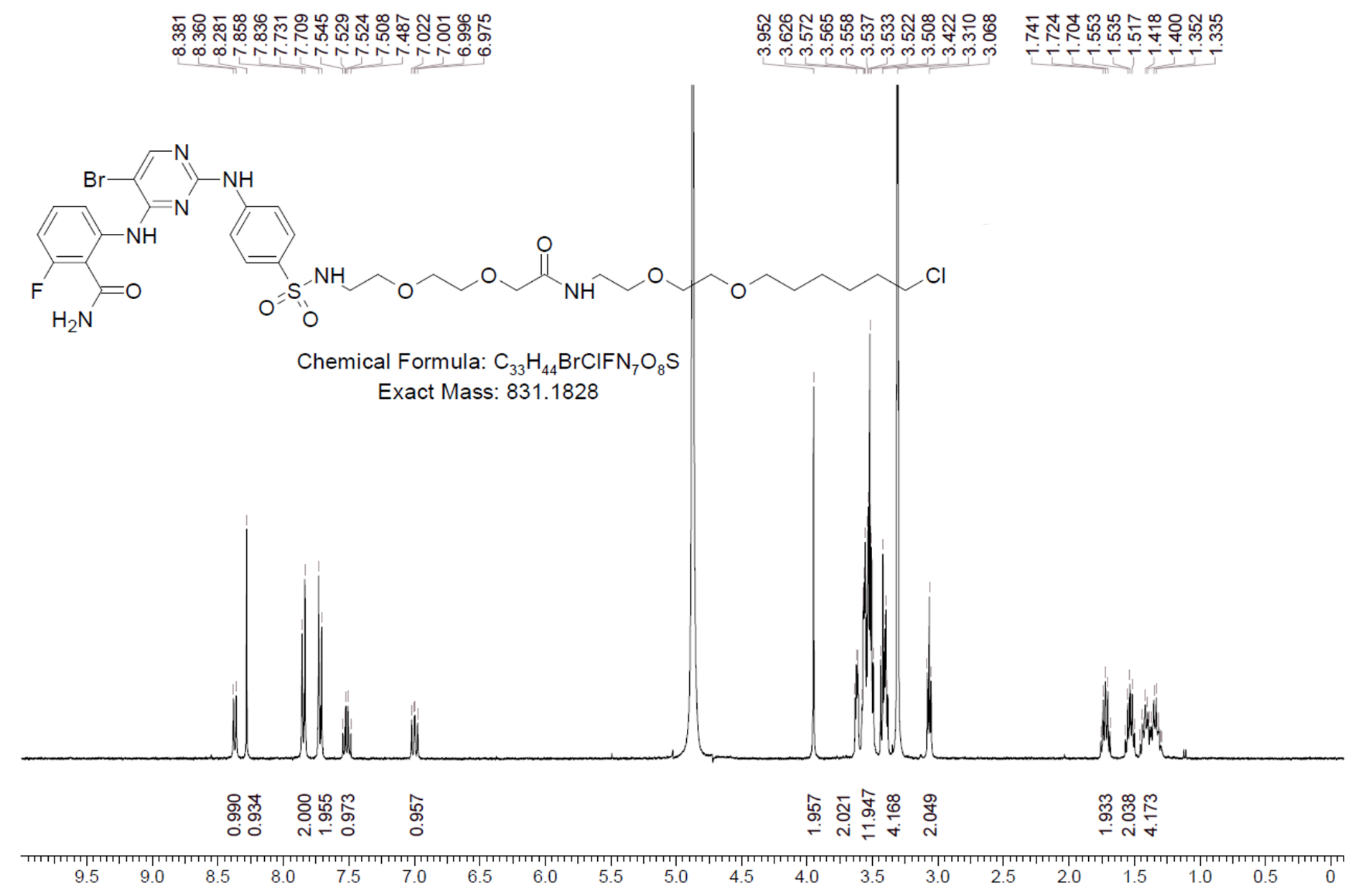

**HLDA-113**

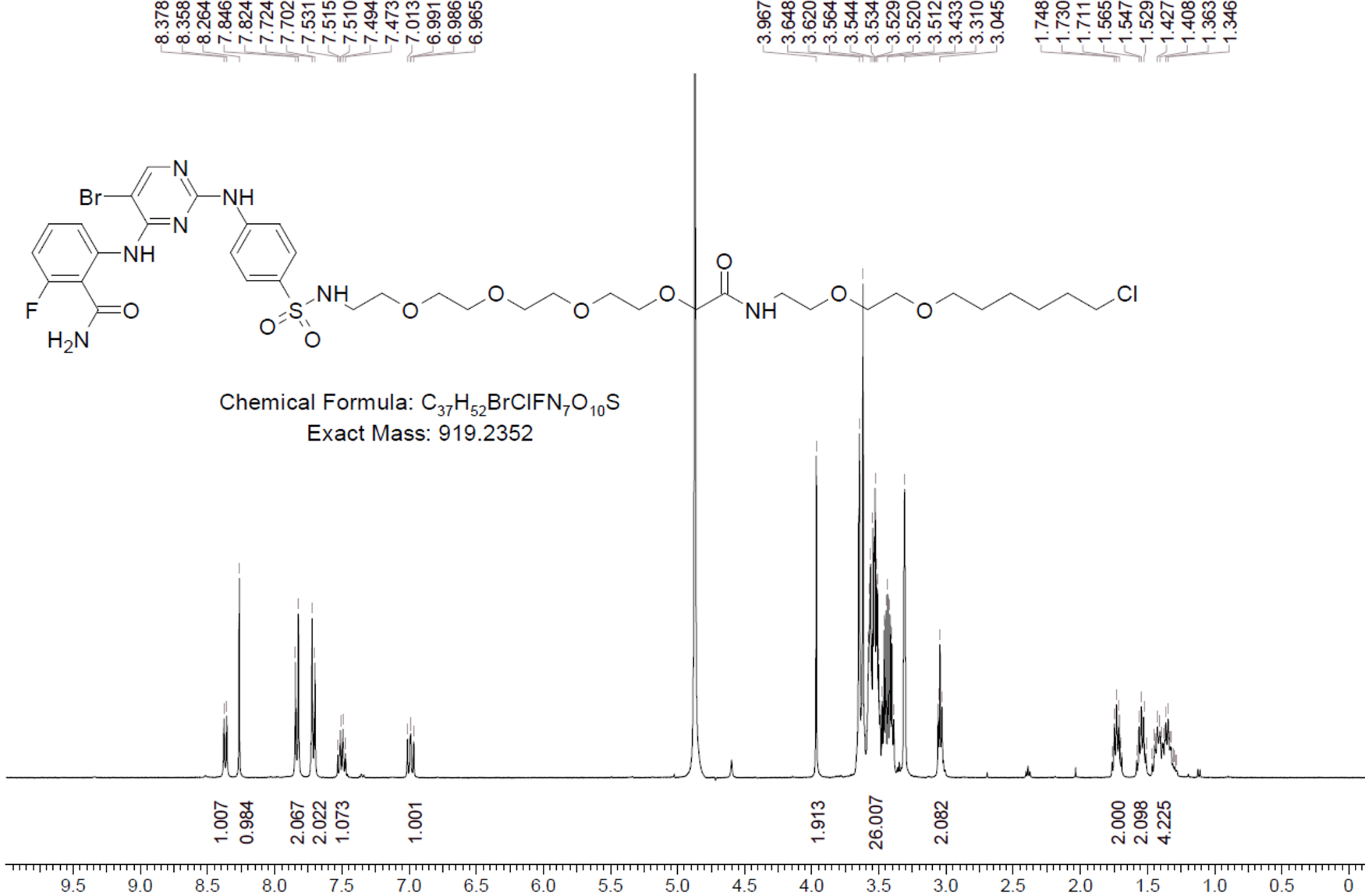

**HLDA-111**

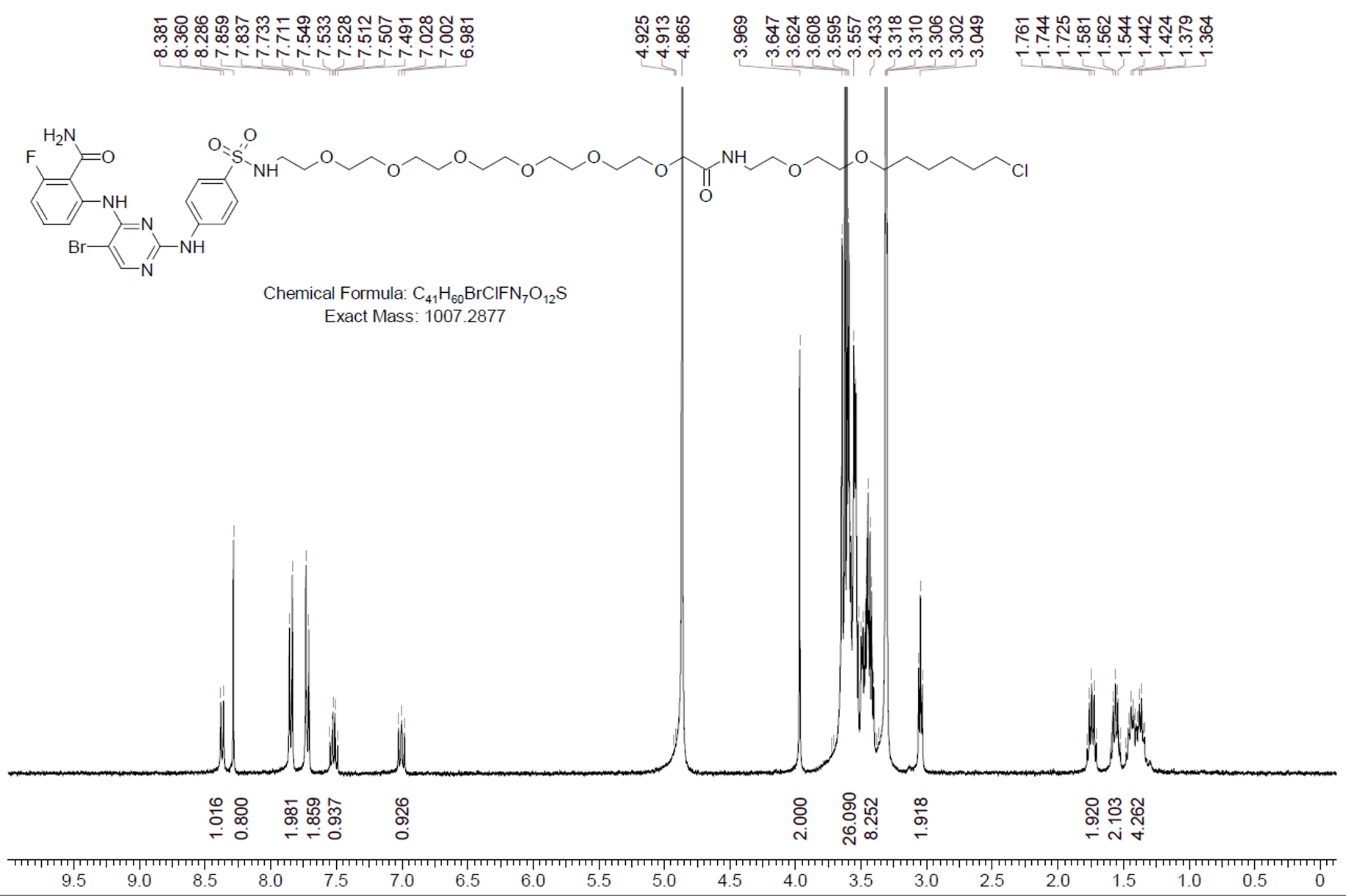

**HLDA-131**

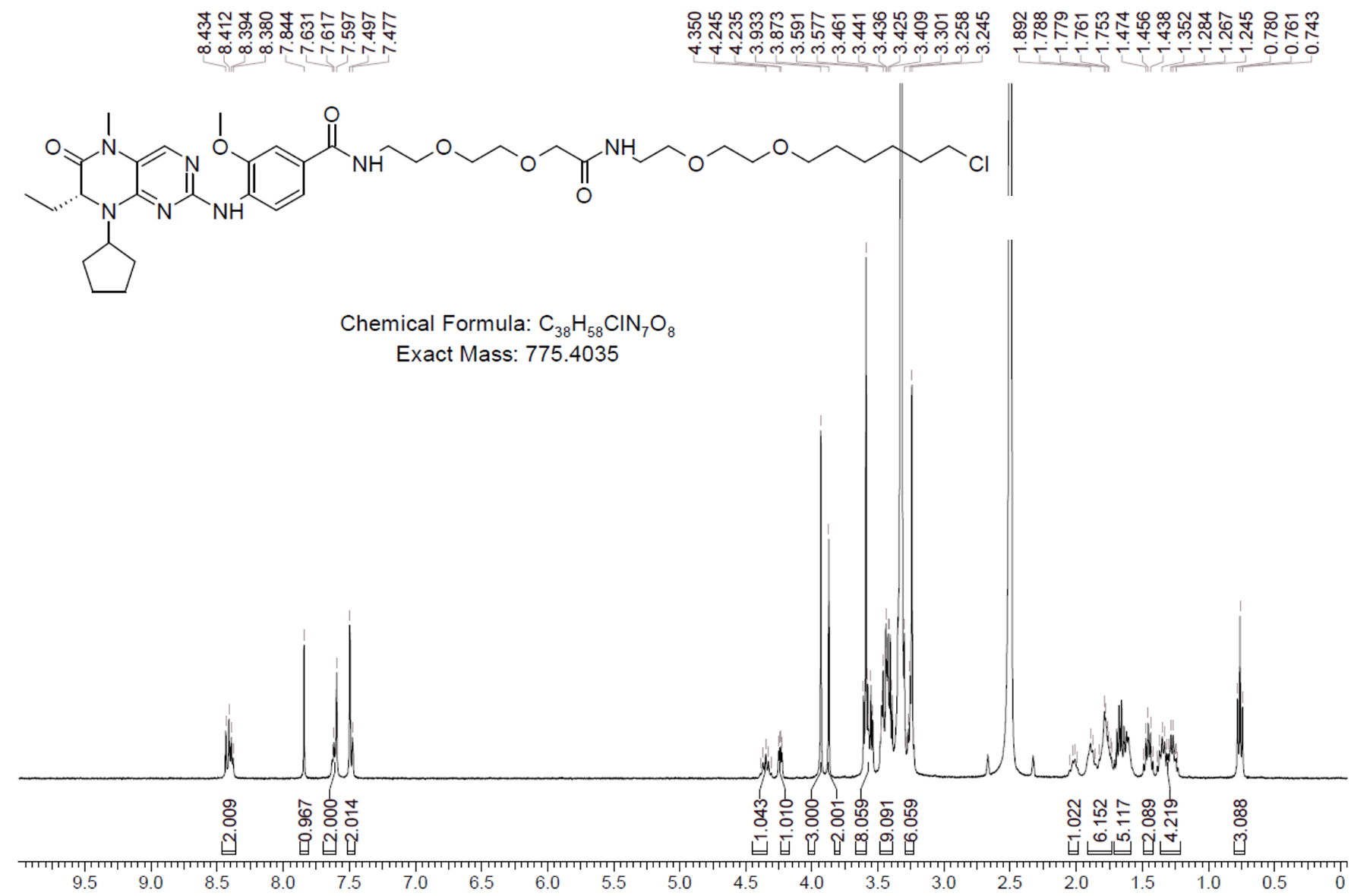

**HLDA-132**

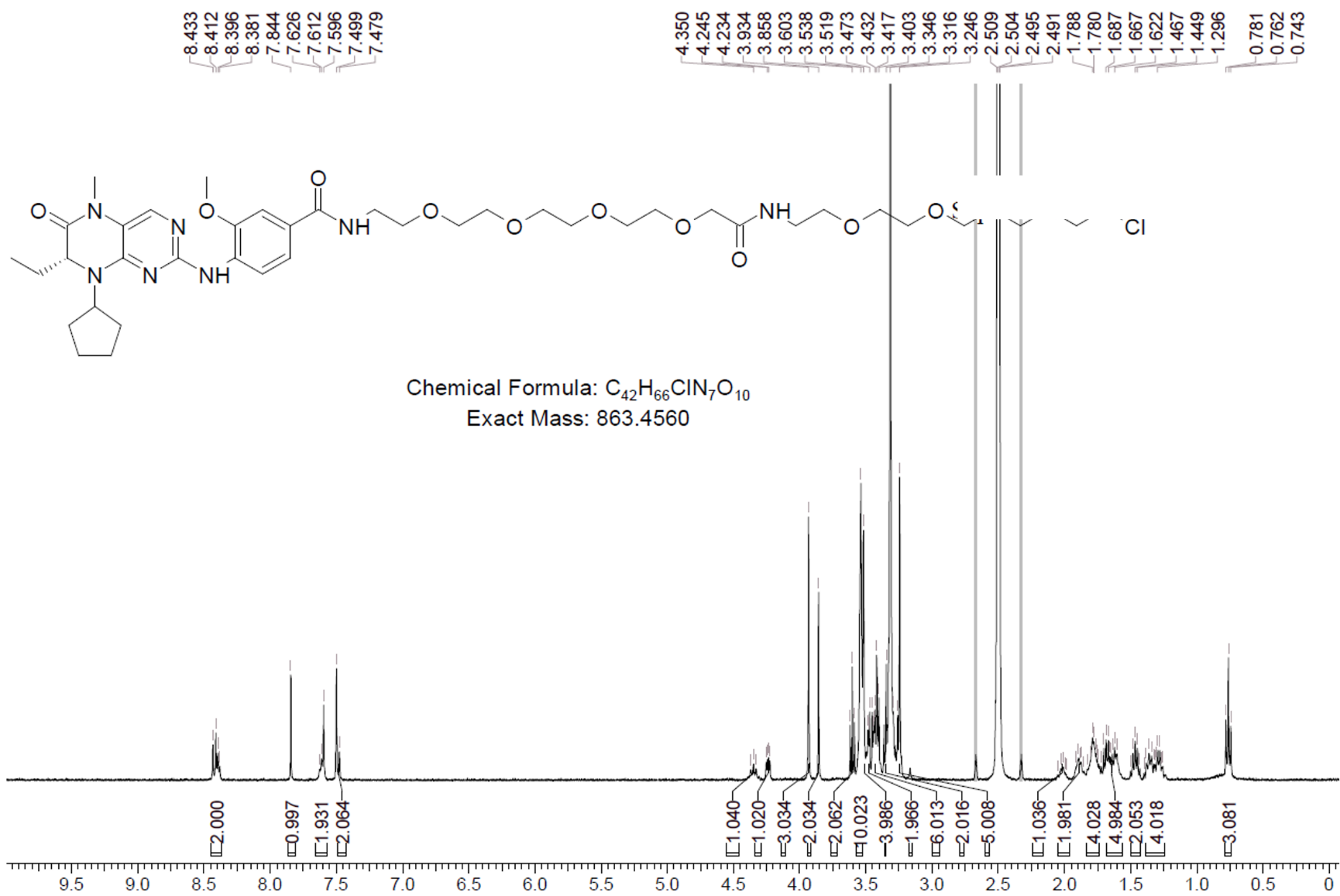

**HLDA-133 – Compound 2**

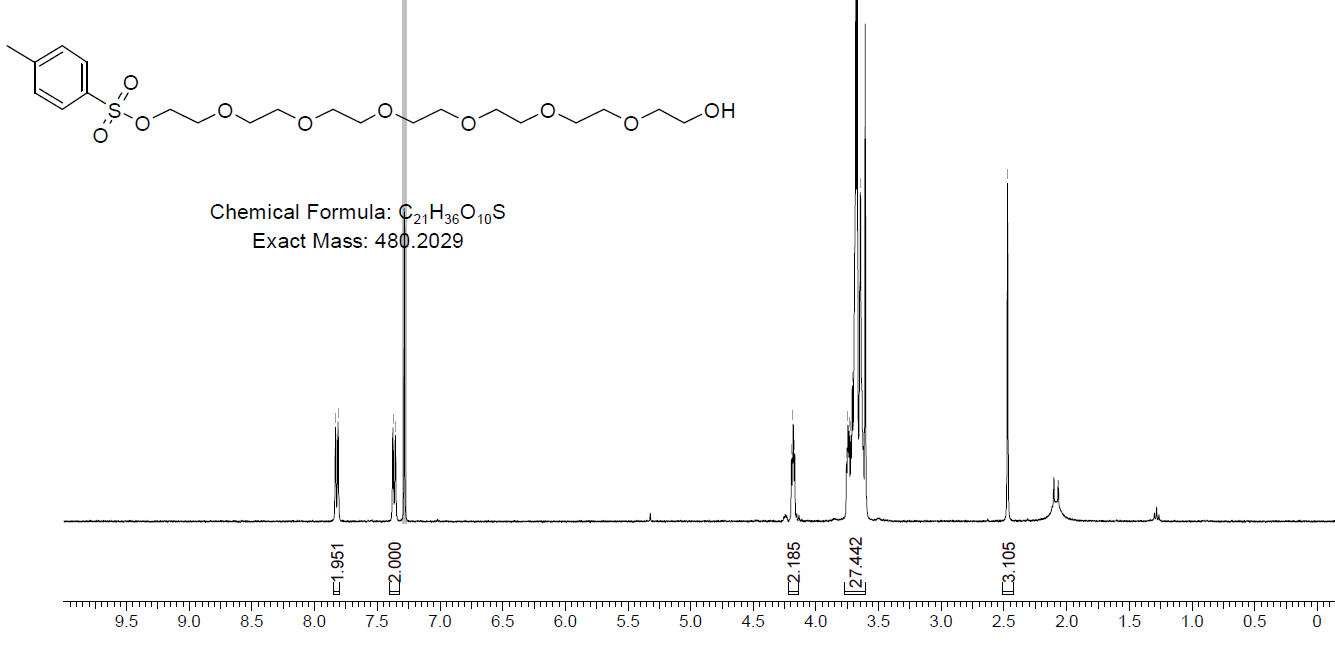

**HLDA-133 – Compound 3**

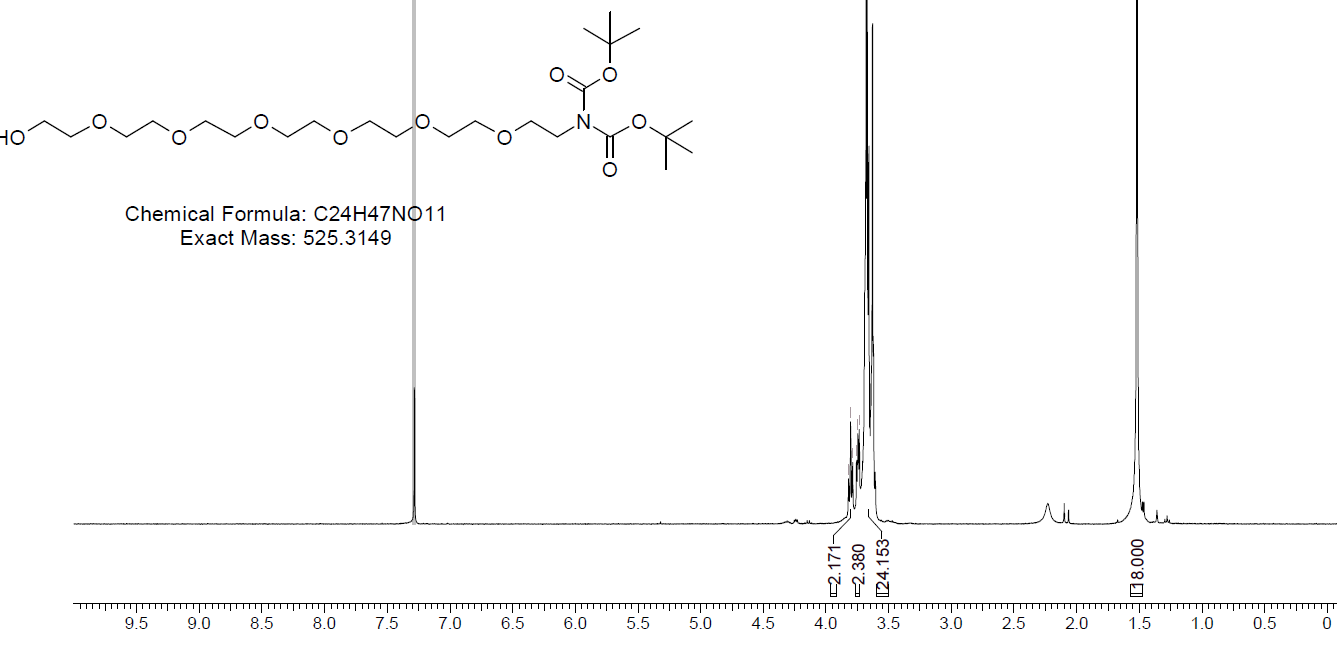

**HLDA-133 – Compound 4**

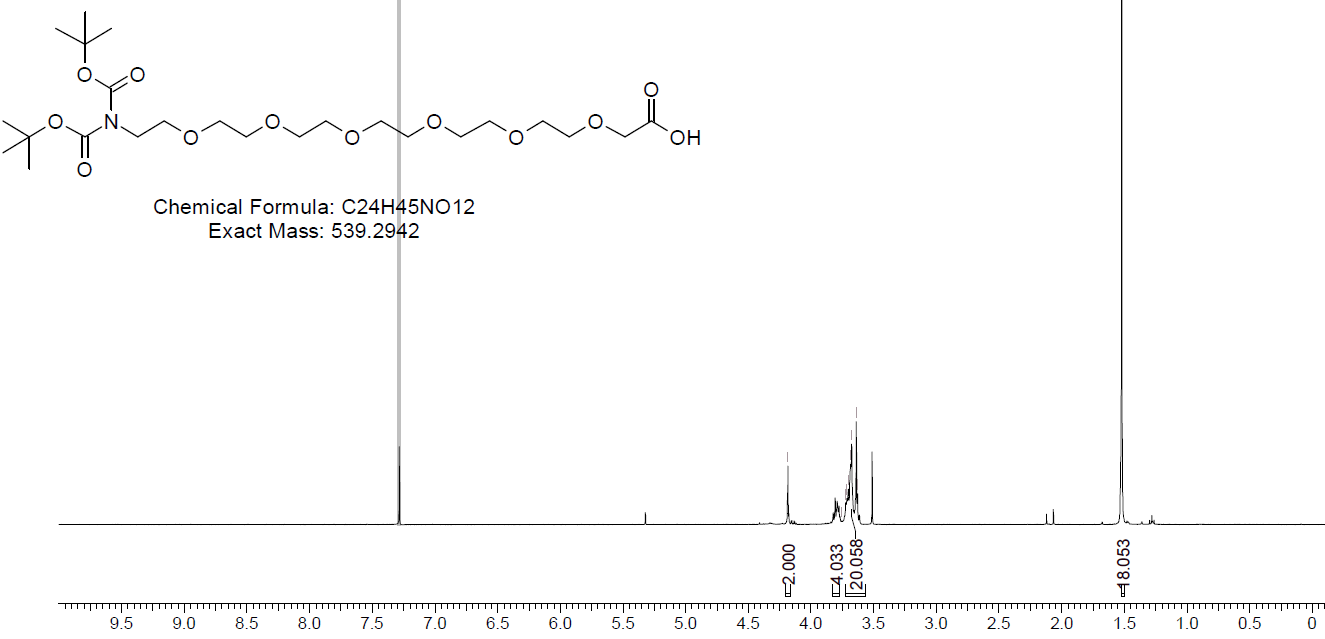

**HLDA-133 – Compound 6**

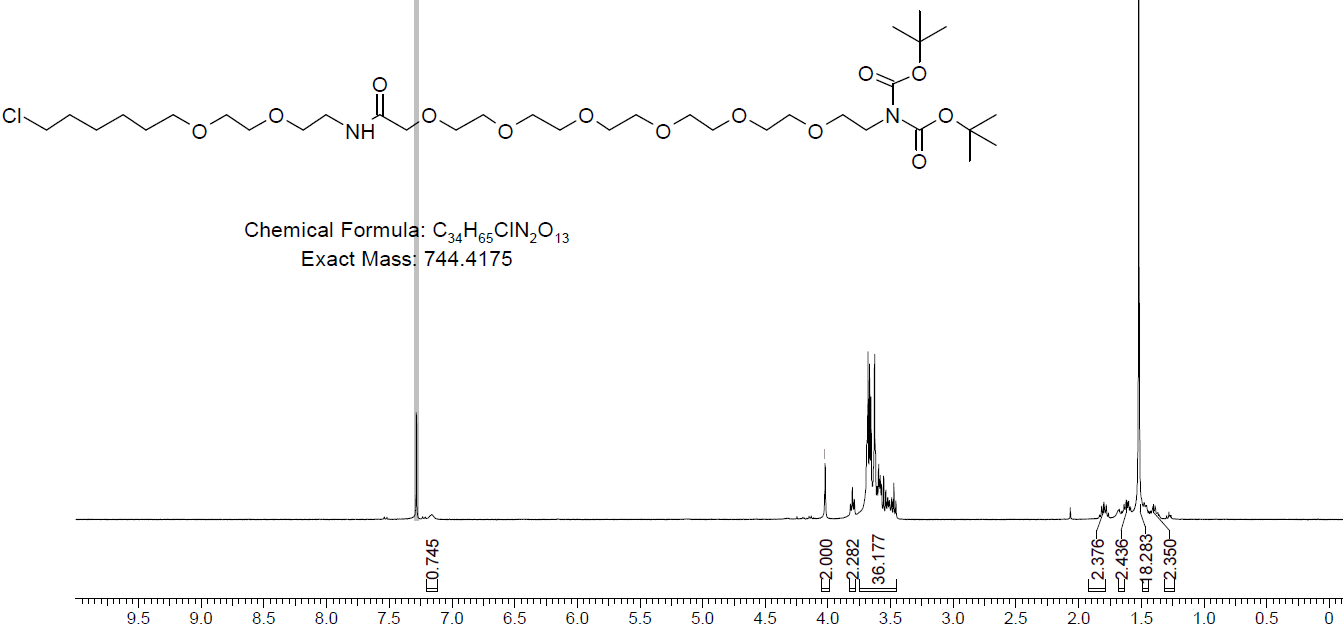

**HLDA-133**

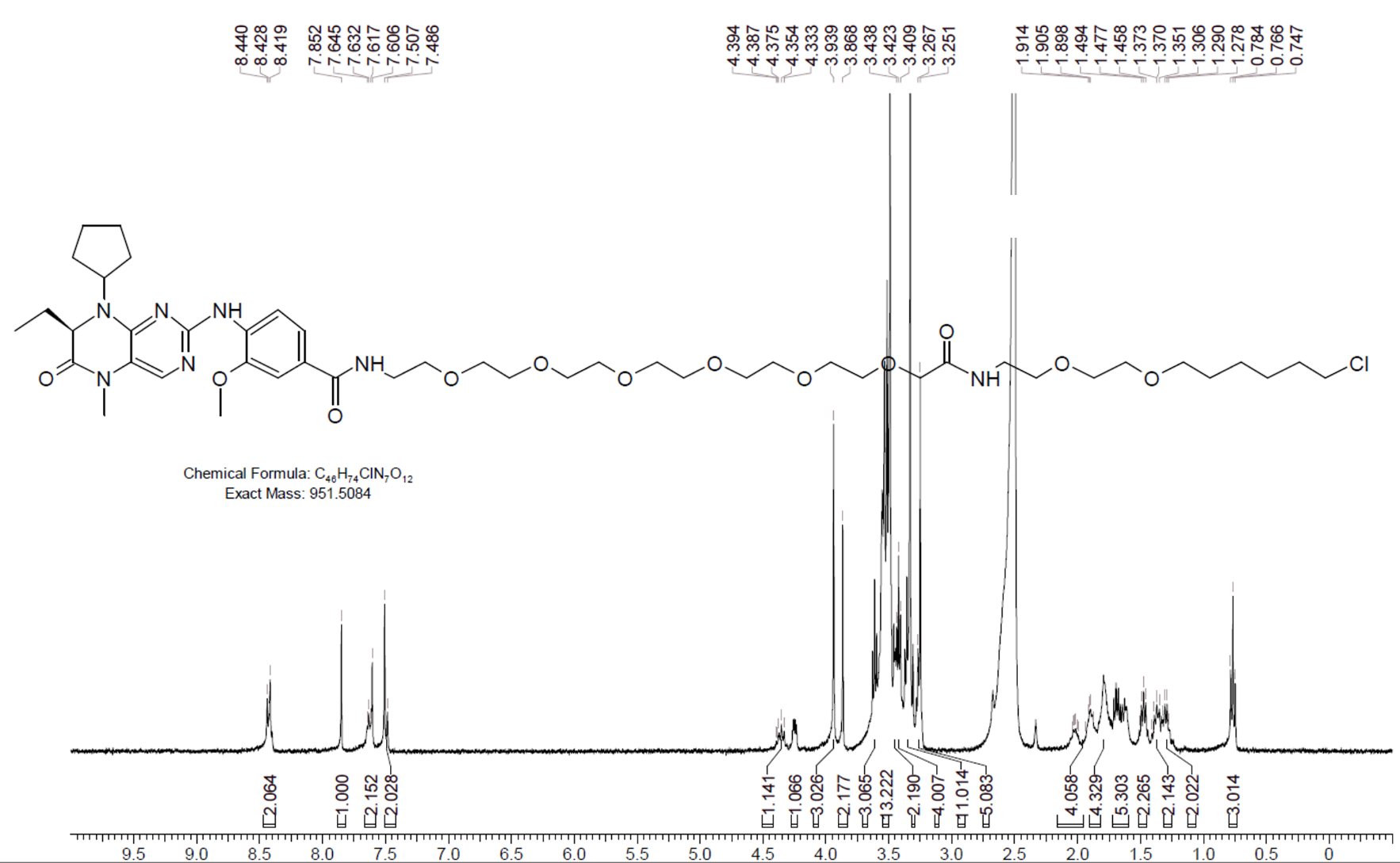

**HLDA-120**

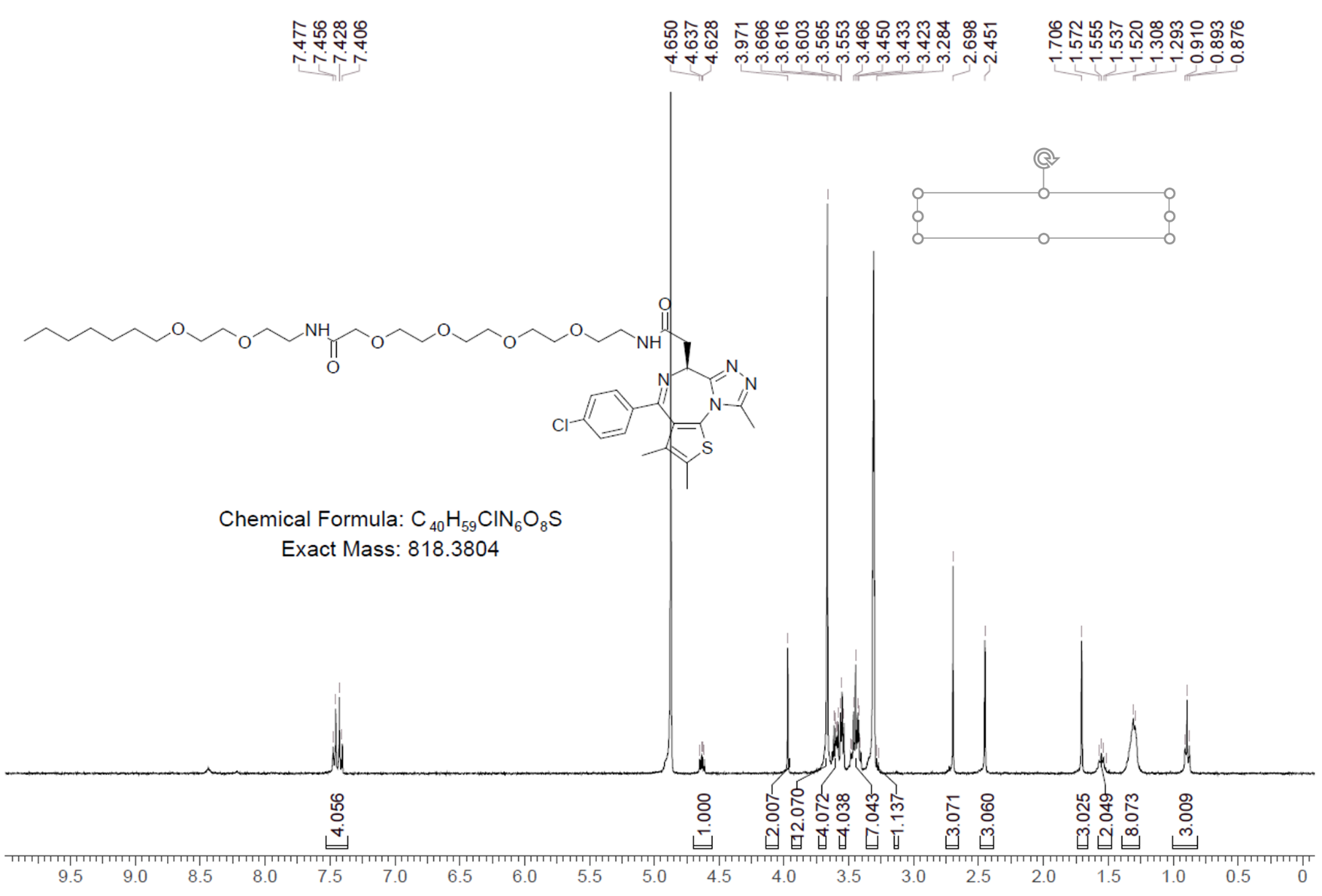

**HLDA-110 – Compound 3**

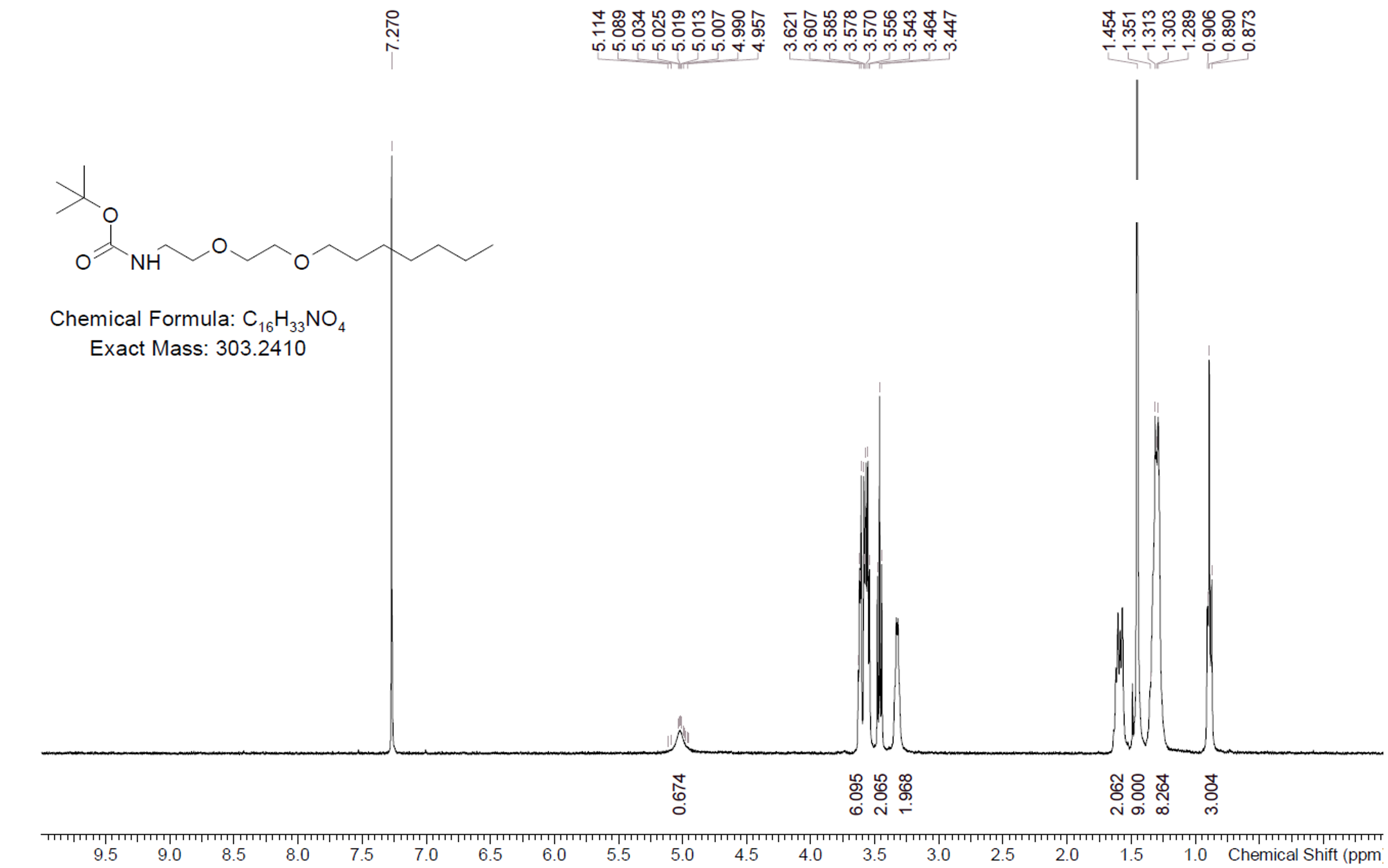

**HLDA-110 – Compound 6**

**HLDA-110 – Compound 9**

**HLDA-110**

**HLDA-117 – Compound 4**

**HLDA-117**

**HLDA-118 Intermediate 8**

**HLDA-118 Intermediate 5**

**HLDA-118 – Compound 3**

**HLDA-118**

**HLDA-119**

**HLDA-114**

**HLDA-115**

**HLDA-116**

**HLDA-221 – Compound 2**

**HLDA-221 – Compound 3**

**HLDA-221 – Compound 6**

**HLDA-221**

**HLDA-222**

**HLDA-223**

**HLDA-231**

**HLDA-232 – Compound 3**

**HLDA-232**

**HLDA-233**

**HLDA-211 – Compound 3**

**HLDA-211**

**HLDA-212 – Compound 3**

**HLDA-212 – Compound 5**

**HLDA-212 – Compound 8**

**HLDA-212 Compound 8 HSQC**

**HLDA-212 Compound 8 HMBC**

**HLDA-212 Compound 8 NOESY**

**HLDA-212**

**HLDA-213 Compound 3**

**HLDA-213 – Compound 11**

**HLDA-213 – Compound 12**

**HLDA-213 – Compound 13**

**HLDA-213 – Compound 1**

**HLDA-213**

**HLDA-001**

`
